# Evolutionary and biological insights into the paternally expressed Snord116-Ipw-Snord115 gene array at the imprinted Prader-Willi syndrome domain

**DOI:** 10.64898/2026.07.28.739784

**Authors:** Virginie Marty, Jade Hebras, Raphael Boursereau, Baptiste Grand, Julie Rivière, Sophie Le Gonidec, Ivan Milenkovic, Christophe Audouard, Pascale Mercier, Jean Personnaz, Eva Maria Novoa, Laure Verret, Jean-Philippe Pradère, Jérôme Cavaillé

## Abstract

Placental mammal-specific box C/D small nucleolar RNA (SNORD) genes within the imprinted human 15q11q13 domain have garnered increasing attention because their poorly understood roles in the brain and their potential involvement in Prader-Willi syndrome (PWS). Using two novel knockout (KO) mouse models, we demonstrate that combined deletion of Snord116 and Snord115 genes, but not the intervening Ipw ncRNA gene, leads to partially penetrant perinatal lethality (40-50%). Snord116/115-deficient neonates display postnatal growth impairment, hypoglycemia and endocrine dysregulation, including failure of the postnatal leptin surge. Despite showing no overt alterations in feeding behavior, adult Snord116/115-KO mice recapitulate most phenotypes previously reported in Snord116-KO models. RNA-seq analyses of the hypothalamus, prefrontal cortex and cerebellum reveal limited global changes. However, we observed upregulation of two of the most compelling putative RNA targets of Snord116 and Snord115 (Dgkk and Htr2c, respectively) in the postnatal hypothalamus. Nevertheless, no evidence was found to support efficient Snord115-guided ribose methylation of Htr2c mRNA. Finally, comparative analyses across 64 representative placental mammal species reveal that many PWS-associated SNORD genes, including SNORD115, display greater evolutionary changes than previously appreciated, raising questions about the functional relevance and evolutionary selective pressures that have shaped the diversification of certain family members across species. Overall, our study provides an unbiased re-assessment of the evolutionary, molecular and physiological significance of the paternally expressed Snord116-Ipw-Snord115 genomic interval and highlights the early postnatal period as a critical, yet largely underexplored, developmental window during which recently evolved SNORDs likely function as dispensable fine-tuners of gene expression.

## Introduction

Box C/D small nucleolar RNAs (SNORDs) are an evolutionarily ancient class of antisense small non-coding RNAs. Once incorporated into ribonucleoprotein particles (snoRNP), the vast majority of them guide sequence-specific 2′-O-ribose methylation of rRNA, U-snRNAs, and tRNAs, and, to a lesser extent, N4-acetylcytidine modification of rRNA [1–6]. As implied by their names, they are all characterized by the presence of two short conserved motifs, the C box and the D box, typically located at their 5’ and 3’ termini, respectively; as well as related internal variants (C′ and D′). The C/D and C′/D′ boxes together form a kink-turn (k-turn) structure that plays a pivotal role in snoRNP assembly, stability, nucleolar compartmentation and reactivity. Another essential feature of most SNORDs is the presence of one or two conserved antisense elements (AEs) located upstream of the D or D′ box. By forming transient ∼10-18 nt base-pairing interactions with complementary sequences in the target RNA, the AE-D or AE-D′ module directs the snoRNP-associated methyltransferase Fibrillarin to catalyze 2′-O-methylation at the target nucleotide, which is invariably positioned opposite the fifth nucleotide upstream of the D or D′ box [7,8].

A subset of SNORD retains the canonical features of RNA methylation guides; however, they lack identified RNA targets in vivo [9,10]. Many of these “orphan” SNORD genes are organized into large clusters whose expression is regulated by genomic imprinting, a developmentally controlled epigenetic mechanism that trigger parent-of-origin-specific silencing of one of the two parental alleles [11]. To date, imprinted SNORD genes have been identified at two evolutionarily distinct chromosomal domains: the paternally expressed SNORD64, SNORD107, SNORD108, SNORD109A/B, SNORD115 and SNORD116 genes on human chromosome 15q11-q13 (mouse chromosome 7c) and the maternally expressed SNORD113 and SNORD114 genes on human chromosome 14q32 (mouse distal chromosome 12). In contrast to canonical SNORDs, which are evolutionarily conserved, ubiquitously expressed and typically encoded by single or low-copy-number genes, imprinted SNORDs are restricted to placental mammals and are organized in large tandem arrays comprising dozens to hundreds of related gene copies [12–14]. Notably, they display strong, and in some cases exclusive, neuronal expression, suggesting tissue-specific roles, potentially mediated through interactions with mRNAs. Consistent with this hypothesis, SNORD113, SNORD115 and SNORD116 have been proposed - but not yet formally demonstrated in vivo - to target several mRNAs, notably integrin pathway mRNAs, Htr2c, Nhlh2, Fgf13 and Dgkk [12,15–18].

The imprinted domain at 15q11q13 has attracted significant interest due to its association with the Prader-Willi syndrome (PWS), a neurodevelopmental disorder that primarily affects hypothalamic function [19]. This rare disease is associated with a broad spectrum of developmental, metabolic, behavioral and endocrine abnormalities ([20–22] for a more detailed clinical picture). Briefly, it manifests in two main distinct phases: an initial perinatal stage characterized by hypotonia and poor feeding followed by hyperphagia in early childhood, which can result in life-threatening obesity if left unmanaged. PWS is caused by the simultaneous loss of approximately a dozen paternally expressed transcripts, both protein-coding and non-coding RNAs. It is therefore still debated whether PWS results from the loss of a single gene or, more likely, from the combined deficiency of multiple genes.

Despite this inherent uncertainty, a potential link between SNORD gene loss and PWS emerges from a few clinical studies that have identified paternal deletions encompassing the SNORD116-IPW-SNORD115 genomic interval, with the SNORD116 array being particularly affected [23–28]. A case of PWS caused by a microdeletion that apparently does not directly affect SNORD116 expression has, however, also been reported [29]. Additional experimental evidence comes from Snord116 knockout (Snord116-KO) mice, which exhibit growth retardation, overeating, reduced mobility and altered sleep architecture that, to some extent, mirror alterations observed in PWS [30–36]. However, Snord116-KO did not develop fully penetrant obesity, even when fed a high-fat diet (HFD) and, paradoxically, these mutants appeared relatively protected from diet-induced obesity [32,37,38]. Whether Snord116-deficient mice exhibit true hyperphagia relative to their reduced body weight remains also a matter of debate [33,34]. Thus, like other PWS mouse models, Snord116-KO mice do not recapitulate the full spectrum of PWS human clinical traits (reviewed in [39–42]).

Although substantial evidence supports an involvement of SNORD115 in the post-transcriptional regulation of the serotonin 2c receptor (Htr2c) pre-mRNA, either alternative splicing or A-to-I RNA editing [12,13,43,44], the putative regulatory function of SNORD115 remains poorly defined *in vivo*. Despite showing increased basal firing rates in both dopaminergic and serotonergic neurons and subtle alterations in monoamine levels [45], Snord115-KO mice do not exhibit significant changes in food intake, energy balance, emotional reactivity or behaviors typically associated with Htr2c signaling [46]. In addition, the in vivo impact of Snord115 loss on Htr2c post-transcriptional regulation remains modest, if detectable at all [46].

Thus, despite many efforts over the past 25 years, the precise role of SNORD116 and SNORD115 genes in brain function remains incompletely understood, both in vitro and in vivo, and key questions are still unresolved or subject to alternative interpretations. There is therefore a real need to pursue their functional dissections, both in terms of molecular mechanism of action, physiological relevance or gene-phenotype relationships underlying PWS. Given their recent emergence in placental mammals, investigating their evolutionary trajectories may also help clarify their “raison d’être,” including whether they perform redundant regulatory functions, whether their tandemly repeated organization is evolutionarily constrained and whether gene copy number has functional significance. Within this conceptual framework, we integrated phylogenetic comparative analyses with molecular and phenotypic assessments of two novel constitutive KO mouse models: one lacking the entire Snord116-Ipw-Snord115 genomic region and the other lacking only the Ipw ncRNA gene, which is located between the Snord116 and Snord115 clusters and whose functional relevance, if any, has remained elusive.

## Results

### Cross-species variation in SNORD116 and SNORD115 gene copy number

Our understanding of the evolutionary history of placental mammal-specific SNORD genes within the imprinted PWS domain relies from a relatively limited number of eutherian species [12,15,47–49]. To gain deeper insight, we conducted a comprehensive comparative analysis of all PWS locus-associated SNORD across 64 representative species from the four major superorders of placental mammals (Supplementary Table 1). As evidence of selective pressure constraints, we examined the conservation of their AE/D or AE/D′ modules, as illustrated in Figure 1A for SNORD115 and SNORD116. Given the expected redundancy among repeated SNORD sequences, functional conservation was inferred when at least one SNORD gene copy exhibited a perfect AE of ≥10 nucleotides and retained their ability to form a k-turn, as illustrated in Figure 1B.

**Figure 1-.**
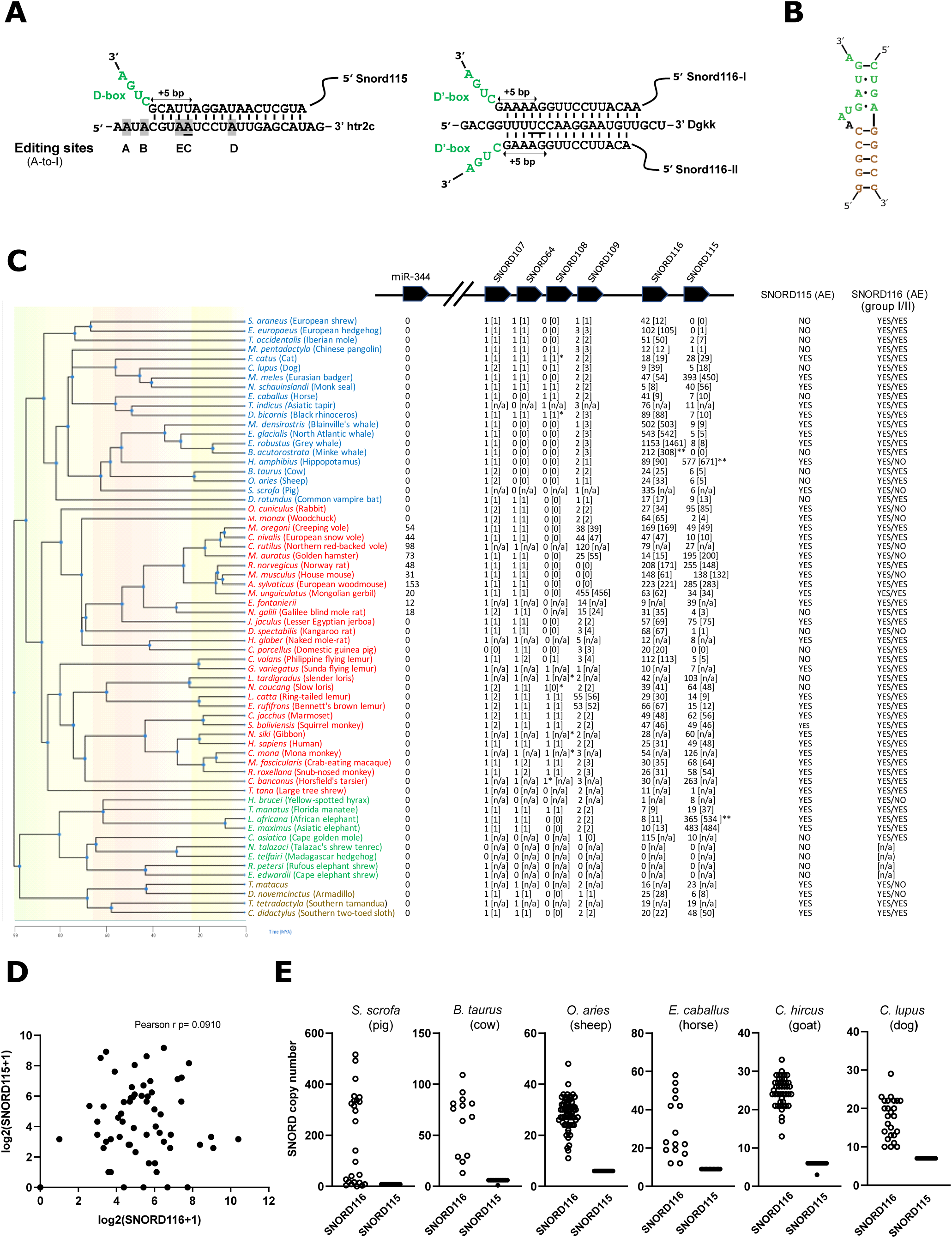
Evolutionary trajectories of PWS-associated SNORD genes in Eutherian. **(A)** Schematic representation of predicted base-pairing interactions between SNORD115 and Htr2c mRNA (left) and SNORD116 and Dgkk mRNA (right). The nucleotide predicted to undergo 2′-O-ribose methylation (i.e., the nucleotide paired to the fifth residue upstream of the D/D′ box) is underlined. The five A-to-I editing sites (A-D) within Htr2c mRNA are shown in grey. Note that one of them (the C-site) is proposed to be targeted by SNORD115-mediated ribose methylation. Owing to the presence of two major SNORD116 isoforms (group I and group II) with related antisense elements, SNORD116 potentially targets two distinct and consecutive positions within Dgkk mRNAs. **(B)** Schematic representation of a canonical k-turn motif. The C and D boxes are indicated in green, while nucleotides forming the characteristic 5′-3′ terminal stem, which is not present in all SNORDs, are shown in brown. Lowercase letters at the 5′ and 3′ ends indicate intronic nucleotides that are not always retained in the fully processed snoRNA. **(C)** Phylogenetic tree of representative placental mammals with species names color-coded as follows: Laurasiatheria (blue), Euarchontoglires (red), Afrotheria (green) and Xenarthra (brown). The estimated numbers of SNORD genes, either determined by BLAST searches or automated genome annotations (shown in brackets when available), are provided for PWS-locus-derived SNORD and the miR-344 gene family. This numbering should be regarded as relative, as it may vary slightly depending on the BLAST search parameters. Conservation of antisense elements (AE) in SNORD115 and SNORD116 predicted to target Htr2c or Dgkk mRNAs is indicated. *, SNORD sequence variants unlikely to be correctly processed; ** SNORD gene copies detected on both DNA strands, suggesting complex genomic organization or possible assembly errors; [n/a], not available. Note that tandemly repeated miR-344 genes between the Snrpn and Ndn loci can be found in many - though not all - Myomorpha species. **(D)** SNORD116 and SNORD115 gene copy numbers exhibit uncoupled evolution across mammalian species. **(E)** Estimates of SNORD116 and SNORD115 gene copy numbers in some domesticated species, as indicated above the panels (Supplementary Table 1).

As summarized in Figure 1C, BLAST searches identified PWS-associated SNORD genes in nearly all examined species, with notable exceptions in certain Afrotherian groups. Specifically, *N. talazaci* and *E. telfairi* appear to have lost all SNORD genes while *R. petersi* and *E. edwardii* retained only SNORD107, suggesting massive lineage-specific losses within Tenrecidae (tenrecs) and Macroscelidea (elephant shrews). The number of SNORD116 copies ranged from an apparent single copy in H. brucei (yellow-spotted hyrax) to 1153 copies in E. robustus (gray whale). Strikingly, whereas several hundred copies of SNORD115 were identified in phylogenetically-distant mammals like *H. amphibius* (hippopotamus), *M. meles* (Eurasian badger) and L. africana (African elephant), SNORD115 copies were not detected in *C. porcellus* (domestic guinea pig), *B. acutorostrata* (minke whale), *S. araneus* (European shrew) or E. europaeus (European hedgehog). Overall, this count of SNORD gene copies did not identify a clear relationship between the numbers of SNORD116 and SNORD115 copies within a given species (Figure 1D).

The availability of an ever-growing number of genomes for domesticated species prompted us to investigate variation in SNORD115 and SNORD116 copy numbers between breeds and among individuals. As illustrated in Figure 1E, all examined species exhibited a consistent pattern: SNORD116 copy number is variable whereas SNORD115 remains relatively stable. This variability in SNORD116 was particularly pronounced in S. scrofa, where copy numbers ranged from 516 to apparent complete absence in the Meishan breed (Supplementary Table 1). Biases inherent to genome assembly of repeated sequences may contribute to variation in SNORD116 copy number. Indeed, estimates of SNORD116 copy number in the same *Bos taurus* DNA sample varied substantially depending on the sequencing technologies used ([50]; Supplementary Figure 1A). To mitigate such assembly-related artifacts, we analyzed a set of 64 high-quality *de novo* genome assemblies representing 14 French cattle breeds, all generated using a standardized methodology by the same group and further refined through three iterative polishing steps using both PacBio CLR long-read and Illumina short-read data [51]. As shown in Supplementary Figure 1B, we confirm inter-individual heterogeneity in SNORD116 copy number. Finally, similar results were also obtained from the analysis of an additional set of 16 high-quality, haplotype-resolved genomes generated using PacBio HiFi sequencing ([52] Supplementary Figure 1C). Altogether, our findings suggest that SNORD gene copy numbers vary not only across mammalian species, as previously noticed [15,47] but also possibly among individuals, providing only limited support for the notion that the SNORD115 and SNORD116 gene arrays evolve in a tightly concerted manner [49].

### The SNORD115 gene family exhibits lineage-specific patterns of rapid evolutionary changes

We next examined mutation rates across orthologous SNORD115 genes, focusing on the AE/D module predicted to target a conserved segment of Htr2c mRNA found in all examined placental (Eutherian) mammals, but not in Prototherian (monotremes) or Metatherian (marsupials) species (Figure 1A-left, supplementary Figure 2). In phylogenetically distant eutherian species, such as *M. musculus*, *H. sapiens*, *M. meles* (Eurasian badger), *H. amphibius* (hippopotamus) or *C. didactylus* (two-toed sloth), most SNORD115 gene copies maintain base-pairing potential with Htr2c mRNA. Conversely, and rather unexpectedly, in some species, none of the SNORD115 copies retain an appropriate AE while in others, only a small fraction has preserved this essential feature of functionality (Figure 1C). This is particularly striking in the primate *C. bancanus* (tarsier) and the rodent *M. auratus* (golden hamster) in which fewer than 20 of the ∼200 SNORD115 gene copies retain the predicted ability to interact with Htr2c mRNA, in marked contrast to the patterns observed in the other rodents and primates examined (Figure 2A; supplementary Figure 3). A distinct evolutionary trajectory is observed in two Lorisiform nocturnal primates - *L. tardigradus* and *N. coucang* - in which most SNORD115 copies are highly similar to one another (Figure 2A, supplementary Figure 3), yet uniformly harbor mutations within their AE indicating that the capacity to fine-tune Htr2c signaling has been lost (Figure 2B-top).

**Figure 2-.**
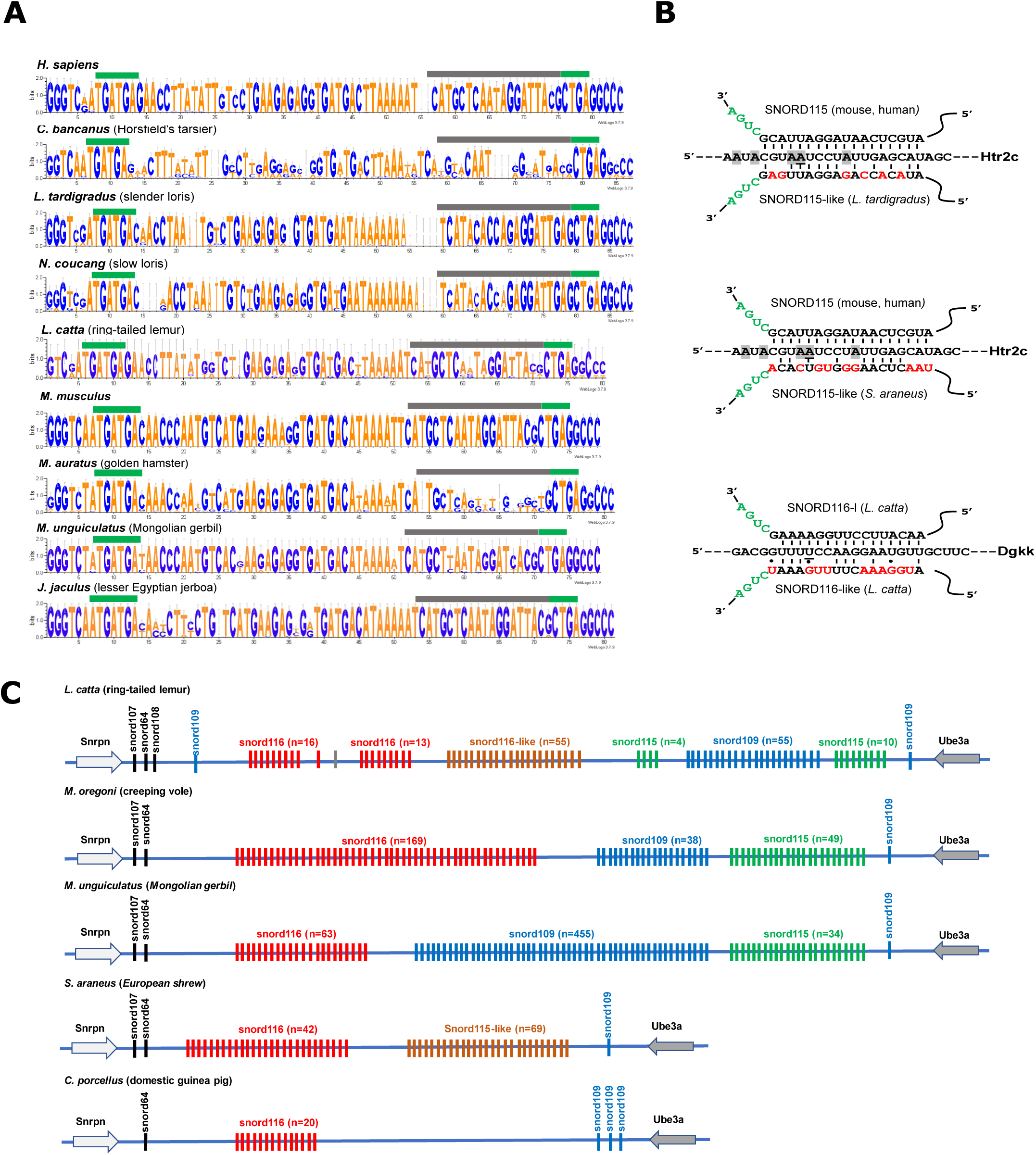
Birth and death of SNORD genes at PWS. **(A)** Sequence logo alignments of SNORD115 gene copies identified in selected primates and rodents (indicated above each alignment). The conserved C- and D-box motifs are shown as horizontal green bars while horizontal grey bars indicate the relative position of the Htr2c antisense element. **(B)** Predicted base-pairing interactions between newly-identified SNORD115-like and Htr2c mRNA (top and middle) and SNORD116-like and Dgkk mRNA (bottom). Nucleotide substitutions that abolish target recognition are highlighted in red. **(C)** Schematic representations of SNORD gene organization highlighting gene gain and loss. SNORD115 appears absent in guinea pig (C. porcellus) and highly divergent in hedgehog (S. araneus), hence its designation as SNORD115-like. In contrast, SNORD109 undergoes extensive gene amplification in the ring-tailed lemur (L. catta) and in the rodent creeping vole (M. oregoni) and M. unguiculatus (Mongolian gerbil). For clarity, the SNORD gene clusters are not drawn to scale and their relative positions are indicative only.

The apparent absence of SNORD115, or its inability to hybridize with Htr2c mRNA in some species, was unexpected. In *S. araneus* (European shrew), the absence of SNORD115 is further supported by phylogenetic evidence, since many other examined Eulipotyphla genomes also lack recognizable SNORD115 genes (supplementary Figure 4A). Interestingly, Dot Plot analysis of the Snrpn-Ube3a genomic interval *in S. araneus* revealed a cluster of 68 repeated SNORD genes between SNORD116 and Ube3a that escaped detection by BLAST because of their high sequence divergence (supplementary Figure 4A). These SNORD115-like, which are unable to target efficiency Htr2c mRNA (supplementary Figure 3; Figure 2B-middle), were present in some but not all examined Eulipotyphla species (supplementary Figure 4A). Finally, we also obtained experimental evidence consistent with the loss, or extreme divergence, of SNORD115 in the Caviidae lineage: SNORD115 was not detected in whole-brain RNA from *C. porcellus* (guinea pig), whereas SNORD116 and SNORD64, used as positive controls, were readily detected, despite slight RNA degradation during sample preparation (supplementary Figure 4B). In addition, Dot Plot analysis did not reveal repeated sequences between the SNORD116 and Ube3a genes (supplementary Figure 4B). Finally, BLAST searches failed to identify SNORD115-related sequences in two other Caviidae genomes analyzed (*C. tschudii* and *C. aperea;* not shown). Altogether, these observations indicate that SNORD115 genes are either absent in some species or have diverged to such an extent that they escape detection by Northern blotting, BLAST search and Dot-plot analyses.

### The SNORD116 gene family exhibits the highest degree of phylogenetic conservation

SNORD116 contains a conserved AE/D′ module proposed to target the Dgkk mRNA ([15]; Figure 1A). More precisely, two types of AEs, differing by a single nucleotide (referred to as group I and group II), were originally described [53] and are highly conserved across evolution, even though group I were absent in a few species (Figure 1C). Based on this set of 64 mammalian species, AE of SNORD116 appears to be under stronger evolutionary constraint than SNORD115. This does not preclude the existence of additional clusters of SNORD116-like genes. Although we did not explore this possibility in depth, certain Lemuriform primates, such as the ring-tailed lemur (*L. catta*), harbor up to 55 SNORD116-like genes. These additional, shorter SNORD116 copies exhibit substantial nucleotide divergence within their AE/D′ module, suggesting that their putative RNA targets, if any, may differ from those recognized by canonical SNORD116 (Figure 2B, bottom; Figure 2C; Supplementary Figure 3).

### The SNORD109 gene family is widespread and undergo lineage-specific gene amplification

Based on its absence in the rat and mouse, SNORD109 was originally thought to be primate-specific [12,53]. However, our analyses identified ∼ 2-5 SNORD109 copies in the vast majority of examined mammalian genomes, suggesting that its previously described loss in rodents was, in fact, restricted to the Murinae lineage (Figure 1C). In most species, SNORD109 copies contain two putative AEs, with the AE/D module more conserved than the AE/D’ module (supplementary Figures 3 and 5). Remarkably, we also found, for the first time, that SNORD109 has undergo gene amplification events, which have probably occurred on at least two separate occasions: once in Lemuriformes primates (*e.g., L. catta* and *E. rufifrons*) and once in rodents from the Cricetidae (*e.g., M. auratus, M. oregoni, C. nivali, C rutilus*) and Muridae (*e.g., M. unguiculatus, E. fontanierii, N. galili*) families (Figure 1C, Figure 2C, supplementary Figure 5). This propensity for gene amplification at PWS locus does not seem to be limited to SNORD-containing regions as the miR-344 family, located more than 2 Mb away from the SNORD arrays, also show lineage-specific expansions in Myomorpha (Figure 1C).

### The single-copy SNORD64, SNORD107 and SNORD108 genes show mild selective constraint

The single-copy SNORD genes - SNORD107, SNORD108, SNORD64 - have received less attention than repeated SNORD115 and SNORD116 genes. Unlike multi-copy SNORD genes, their sequences are anticipated to be subject to stronger selective pressure, should they perform essential function. Surprisingly, SNORD64 was identified in only 48 of the 64 species examined and did not exhibit a clearly identified conserved AE (supplementary Figure 5). SNORD107, found in 61 out of 64 species, displays only a limited stretch of 8-9 conserved nucleotides upstream of the D’-box whose functionality remain uncertain due to its rather short length (supplementary Figure 5). Finally, SNORD108 was the least conserved, detected in only 22 of the 64 species. Not only does SNORD108 lack a conserved AE, but many SNORD108 sequences also lack canonical C-box, making them unlikely to be properly incorporated into RNP complexes (supplementary Figure 5). Altogether, the frequent gene loss or sequence degeneration in essential RNA elements raises doubts about the biological importance of some single-copy SNORD genes in some species, particularly SNORD108.

### Targeted deletion of the Snord116-Ipw-Snord115 genomic interval in mice

The birth-and-death dynamics of many PWS-associated SNORD suggest they could be regarded as context-dependent fine-tuners of gene expression, rather than as indispensable regulators. Within this framework, we reasoned that murine Snord116 and Snord115 - which are genomically-linked and co-expressed - may also exert redundant functions. This hypothesis predicts that simultaneous deletion of Snord116 and Snord115 would uncover phenotypes that are absent (or attenuated) in single Snord-KO models, especially perhaps under challenging conditions that are more likely to reveal subtle defects. To test this, we generated a novel double Snord116/115-KO mouse model carrying a constitutive, ∼ 580 kb-long CRISPR-Cas9-mediated deletion spanning the entire Snord116-Ipw-Snord115 gene array (Figure 3A).

**Figure 3-.**
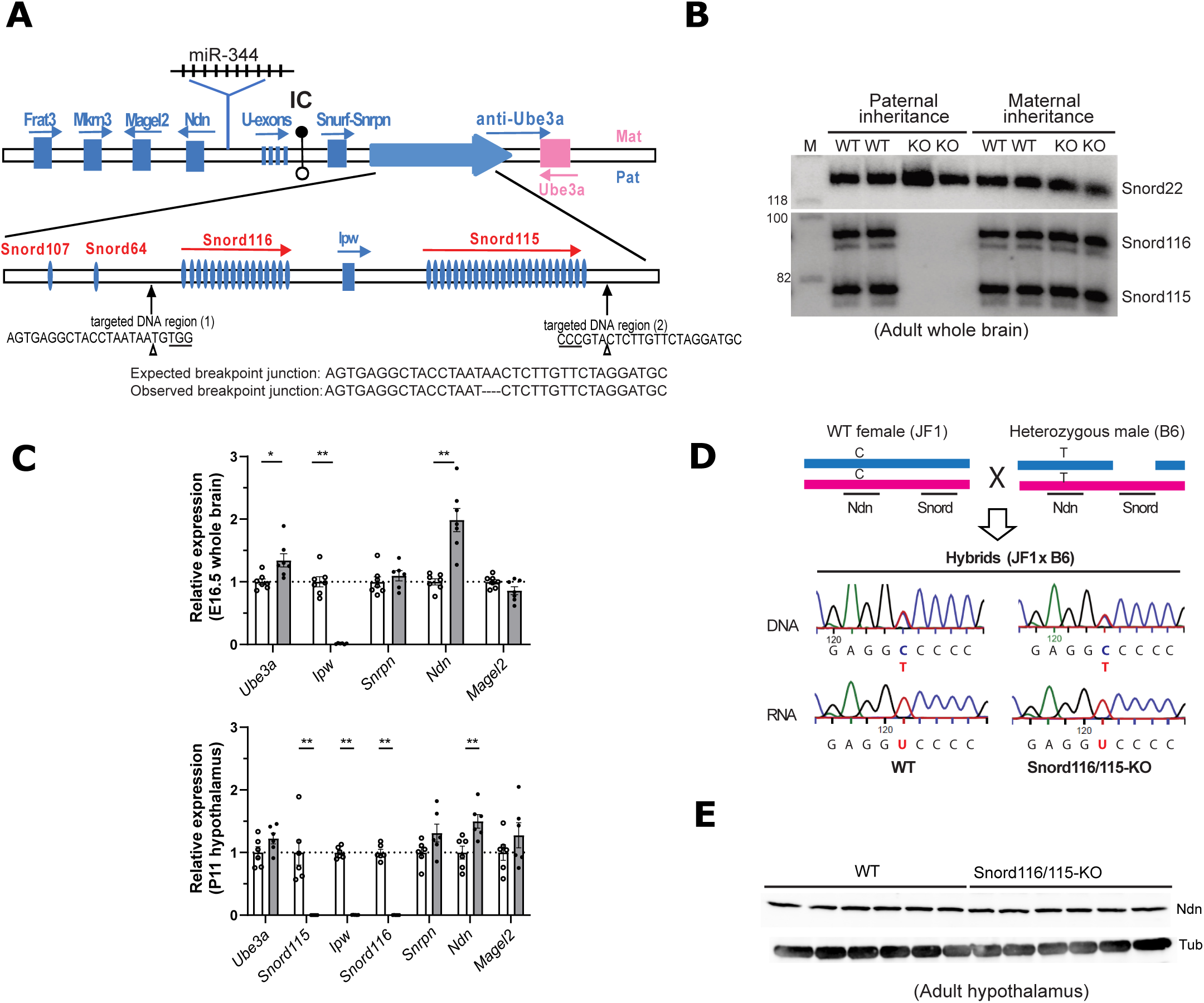
CRISPR-Cas9-mediated deletion of a ∼580 kb genomic interval encompassing the Snord116-Ipw-Snord115 locus. **(A)** Top: Schematic of the murine imprinted Snrpn domain on chromosome 7c. Major paternally and maternally expressed transcription units are shown in blue and pink, respectively. The miR-344 cluster, present in mouse but absent in human and presumed to be paternally expressed, is located between Ndn and the U-exons region. IC, imprinting center; unmethylated paternal and methylated maternal alleles are represented by white and black lollipops, respectively. Bottom: Targeted genomic regions are indicated, with predicted Cas9 cleavage sites marked by arrowheads and PAM sequences underlined. Expected and observed breakpoint junction sequences are shown. Schematic not draw to scale. **(B)** Northern blot analysis of Snord116 and Snord115 expression in adult whole brain from heterozygous mice carrying a paternally or maternally transmitted deletion, as indicated above the panel. Two heterozygous (KO) and two wild-type (WT) littermates were analyzed. Snord22 served as a loading control. M = molecular marker with size indicated in nucleotide. **(C)** Expression analysis of imprinted genes in E16.5 whole brain (top) and P11 hypothalamus (bottom), assessed by RT-qPCR in Snord116/115-KO (grey bars) and WT (white bars) mice. RNA expression levels were normalized to those of Gapdh mRNA. **(D)** SNP-based allelic expression of Ndn in progeny (hybrids JF1xB6) from a cross between a WT female (JF1 background, homozygous C allele) and a heterozygous male (C57BL/6J background, homozygous T allele). Representative Sanger electropherograms of RT-PCR (RNA) and PCR (DNA) products are shown. **(E)** Western blot analysis of Ndn protein levels in whole-cell extracts from WT and Snord116/115-KO adult whole hypothalamus. Ndn was revealed with rabbit anti-Ndn antibodies (Millipore; AB9372; 1:500) while detection of tubulin (Tub) using mouse anti-TUBA4A antibodies (Sigma; T6199; 1:10 000) served as a loading control.

As expected, paternal inheritance of the deletion resulted in the absence of Snord115 and Snord116 in adult whole brain whereas maternal inheritance did not (Figure 3B). Interestingly, the level of mRNA encoding Ndn was upregulated in both E16.5 brain and P11 hypothalamus of Snord116/115-KO mice compared to WT (Figure 3C). The expression of other surrounding imprinted genes remained largely unaffected, with the exception of Ube3A, which was slightly increased in the E16.5 brain. The observed ∼1.5-2-fold increase in Ndn expression, also found in adult brain (see below), prompted us to test the possibility that the normally silent maternal Ndn allele is reactivated in the Snord116/115-KO mice. We then assessed the allelic expression of a SNP between JF1 and C57Bl/6J mouse strains (T50C; rs261911330). As shown in Figure 3D, a cross between a WT JF1 female (C-allele) and a heterozygous B6 male (T-allele) revealed that only the paternal T allele was expressed, both in the hypothalamus of Snord116/115-KO and WT. Thus, the most likely explanation is that lack of Snord116-Ipw-Snord115 gene array results in increased transcription of the paternal Ndn allele and/or promote Ndn mRNA stability. This change in mRNA level was, however, not accompanied by any apparent increase in protein levels in whole adult hypothalamus, as judged by Western blot (Figure 3E).

### Impaired growth and decreased postnatal survival in Snord116/Snord115-KO pups

We first assessed growth and pre-weaning survival in offspring from a cross between a WT female and a heterozygous male which is expected to yield 50% WT and 50% Snord116/115-deficient pups (*i.e.,* the paternal active allele is deleted). Although the mutant pups were born at expected Mendelian ratios with normal gross morphology, including sucking behavior as evidenced by visible milk in their stomachs (not shown), only ∼ 38-56% survived to weaning (Figure 4A). This reduction in perinatal survival, occurring without any obvious warning signs, was constantly observed across two mouse facilities with distinct health statuses (conventional *vs.* SOPF) and, as expected, was not observed when the deletion was maternally inherited (Figure 4A). Closer analysis revealed that neonatal deaths occurred preferentially within the first two days post-birth. Remarkably, the lethality rate in Snord116/115-KO mice was only slightly higher than that observed in Snord116-KO mice (∼35-43%).

**Figure 4-.**
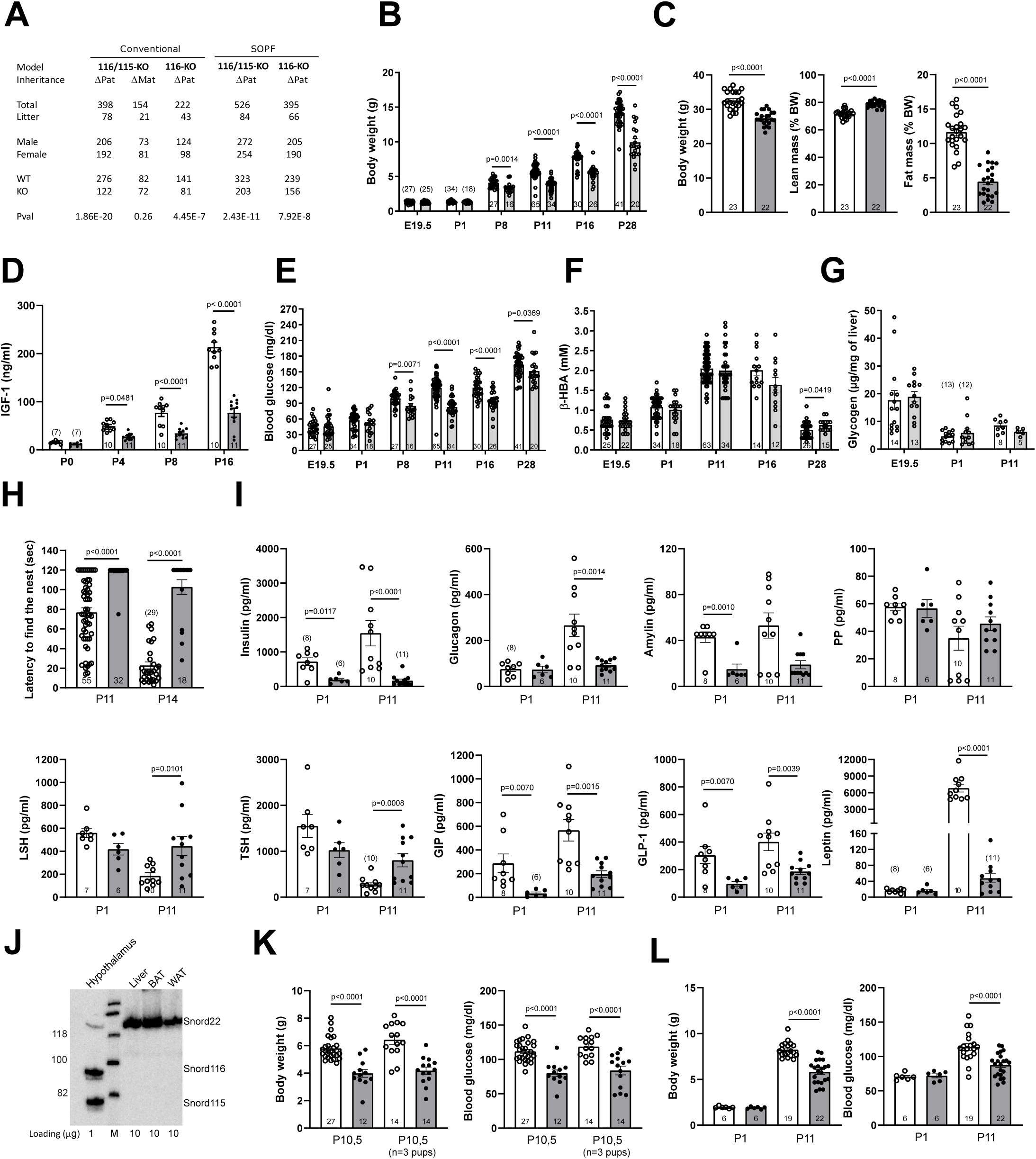
Snord116/115-KO newborn mice show reduced survival rates and metabolic and hormonal abnormalities. **(A)** A paternally, but not maternally, inherited deletion alters Mendelian ratios (WT vs. KO) at 4 weeks of age, as observed in two independent mouse housing facilities (conventional and germ-free/SOPF). **(B-C)** Snord-deficient pups exhibit growth retardation **(B)** that persists into adulthood accompanied by reduced fat mass and increased lean mass **(C)**. **(D-G)** Snord-deficient pups show reduced circulating of IGF-1 **(D)** and glucose **(E)** levels, whereas circulating β-hydroxybutyrate **(F)** and hepatic glycogen **(G)** levels remain within the normal range. **(H)** Snord116/115-KO pups exhibit reduced spontaneous locomotor activity, as evidenced by an increased latency to return to the nest when displaced away from it. (I) Snord-deficient pups display altered circulating hormone levels when assessed at P1 and P11. **(J)** Northern blot analysis showing that Snord115 and Snord116 are undetectable in peripheral P11 tissues (liver, BAT, WAT). Peripheral tissue lanes were loaded at 10× the amount used for hypothalamus (10 µg vs. 1 µg). Snord22 was used as a loading control. M = molecular marker with size indicated in nucleotide. **(K)** Reducing litter size to three pups at birth does not rescue the lower body weight (left) or the reduced blood glucose levels (right) of Snord-deficient pups. **(L)** Snord116/115-KO pups on a mixed FVB-C57BL/6J background exhibit reduced body weight (left) and lower blood glucose levels (right). **(B-L)**. The number of analyzed individuals or biological samples is indicated within each histogram or, where space is limited, in brackets above the histogram. White and grey bars represent WT and Snord116/115-KO mice, respectively.

The surviving mutant pups showed signs of growth retardation relative to their WT littermates, as evidenced by lower body weight (Figure 4B). This growth impairment, which persists into adulthood, is associated with changes in body composition with reduced fat mass and increased lean mass (Figure 4C). Perinatal growth in mouse pups prior to P10 is predominantly driven by a growth hormone (GH)-independent, IGF-1-dependent pathway [54]. Consistent with impaired IGF-1 signaling, mutant pups exhibited reduced circulating IGF-1 levels (Figure 4D). This is in agreement with previous findings from Snord116-KO models [30,32,37].

Although no differences were observed before delivery (E19.5) or P1, Snord116/115-KO pups also exhibited lower blood glucose levels, with hypoglycemia becoming more pronounced at P8 and persisting till weaning (Figure 4E). Ketogenesis appears largely unaffected, as indicated by measurements of β-hydroxybutyrate (β-HBA, Figure 4F), used as a marker of energy availability (lipid oxidation). Glycogen stores before birth were within the normal range in mutant E19.5 embryos and, as expected, were significantly reduced postnatally, indicating that glycogenolysis occurred at normal rates (Figure 4G). Finally, spontaneous activity was decreased in Snord116/115-KO pups at both P11 (eyes closed) and P14 (eyes open), relative to WT littermates (Figure 4H). It is, however, unclear whether reduced activity stems from reduced muscle tonus (*i.e.,* hypotonia) or is merely a consequence of a hypoglycemic state (*i.e.,* reduced energy availability).

### Snord116/115-deficient pups exhibit hormonal imbalances

To determine whether the simultaneous loss of Snord115 and Snord116 affects circulating hormone levels, we conducted a systematic analysis using Luminex technology, focusing on P1 and P11 (Figure 4I). Insulin levels were reduced in mutant pups compared to controls at both P1 and P11 while glucagon levels were decreased only at P11. Notably, levels of amylin (a peptide co-secreted with insulin by pancreatic β-cells) were also significantly reduced at both time points while levels of pancreatic polypeptide (PP) remained unchanged. These hormonal measurements further revealed alterations in pituitary-derived hormones: both luteinizing hormone (LSH) and thyroid-stimulating hormone (TSH) levels were increased at P11. Additionally, levels of the gastrointestinal hormones GIP and GLP-1, were decreased at both P1 and P11. Finally, Snord116/115-KO pups at P11 also failed to mount a leptin surge. Remarkably, the same alterations in hormone levels were observed in Snord116-KO pups (supplementary Figure 6). Altogether, Snord116/115-deficient pups exhibit peripheral multi-organ dysfunction, likely resulting from non-cell-autonomous defects since Snord115 and Snord116 are not expressed in the newborn liver, brown adipose tissue and white adipose tissue (Figure 4J).

### Hypoglycemia and growth retardation are unlikely to be the primary causes of perinatal lethality

To investigate whether reduced competition for nutritional resources could improve the survival of mutant pups, we artificially decreased the number of pups per litter at birth. In a blinded experimental design where only three pups were kept with their mother from P0 to P10.5, no improvement in growth (Figure 4K-left) or glucose levels (Figure 4K-right) were observed while neonatal lethality appeared rescued (14/14 mutants survived; n=15 litters analyzed). Remarkably, growth retardation (Figure 4L-left) and hypoglycemia (Figure 4L-right), but without lethality, was also found in a mixed FVB-C57Bl/6J genetic background obtained by crossing a C57Bl/6J male with a FVB female. These findings are in agreement with previous observations made with SNORD116-KO mice [31] and suggests that maternal factors linked to the FVB background and/or genetic modifiers in the hybrid background rescue perinatal survival. Such explanation was also proposed for the full PWS mouse model [55]. Thus, our observations indicate that lethality is unlikely to be solely caused by hypoglycemia and/or growth failure.

### Ipw-KO newborn mice do not manifest overt perinatal abnormalities

The ncRNA Ipw gene [56] located between Snord116 and Snord115 is part of the long RNA precursor from which Snord116 and Snord115 are processed [53]. Its function, if any, remain unknown, although a role in regulating the imprinted DLK1-DIO3 domain has been proposed using human embryonic stem cells [57]. Because exon-C of Ipw displays high level of conservation (supplementary Figure 7A) and given that this part of Ipw is removed in our Snord116/115-KO mice, as well as in Snord116-KO models and PWS patients with SNORD-centered deletions (Figure 5A), the possibility remains that loss of Ipw may contribute, at least to some aspects, to the perinatal alterations. To test this possibility, we thus generated a Ipw-KO mouse line harboring a ∼1.9 kb-long, CRISPR-Cas9-mediated deletion encompassing exon-C (supplementary Figure 7B). This does not alter the expression levels of Snord115 or Snord116, as judged by Northern blotting (Figure 5B-top) and, similarly, the expression of neighboring imprinted genes - including Ndn - remains unaffected, with only Ipw being, as expected, undetectable (Figure 5B-bottom). Mutant pups show normal survival rates (Figure 5C), with growth (Figure 5D-left) and glucose levels (Figure 5D-right) in the normal range. Although adult Ipw-KO mice were not subjected to metabolic or behavioral analyses, they appear phenotypically normal. We conclude that most, if not all, of the perinatal alterations observed in Snord116/115-KO mutants are not attributable to Ipw deficiency, at least not to the highly conserved non coding region that we deleted.

**Figure 5-.**
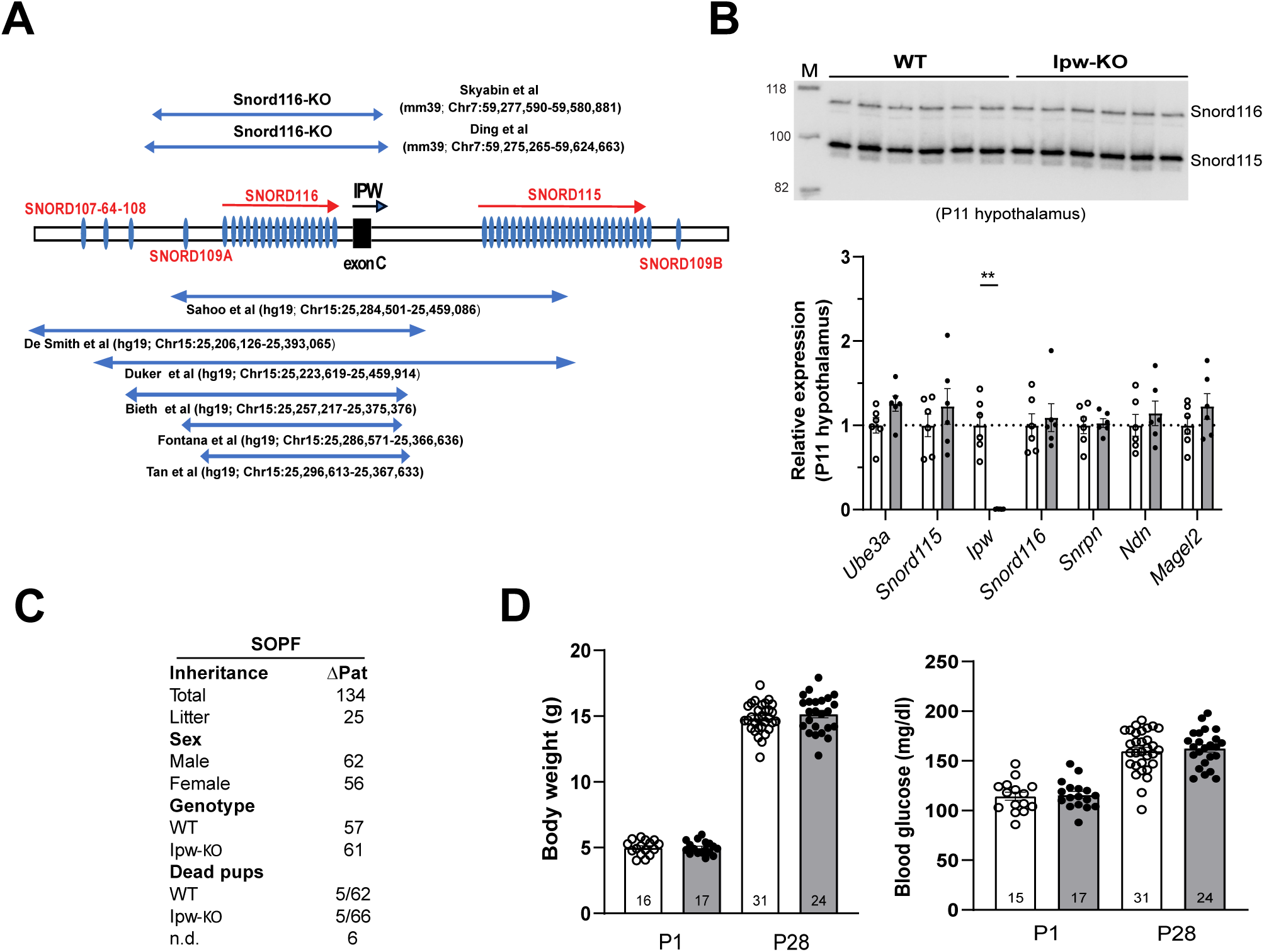
Ipw ncRNA deletion does not recapitulate the perinatal defects observed in the Snord116/115-KO model. **(A)** Schematic representation of the Snord116-Ipw-Snord115 genomic interval (not drawn to scale) showing that the conserved exon C of Ipw, denoted as a black box is deleted in the two available Snord116-KO mouse models (top) as well as in rare PWS patients carrying deletions encompassing SNORD genes (bottom), as indicated by horizontal blue bars. **(B)** Ipw loss does not affect Snord115 and Snord116 expression, nor expression of neighboring imprinted genes. Top: Northern blot analysis performed on total RNA extracted from WT or Ipw-deficient P11 hypothalamus. Bottom: mRNA levels were assessed by RT-qPCR. RNA expression levels were normalized to those of Gapdh mRNA. **(C)** Paternally inherited deletion of Ipw (ΔPat) does not alter Mendelian ratios at weaning in germ-free housing facilities (SOPF). n.d.: not determined due to maternal cannibalism. **(D)** Ipw loss does not lead to reduced body weight (left) or altered blood glucose levels (right). White and grey bars represent WT and Ipw-KO mice, respectively.

### Adult Snord116/115-KO mice do not exhibit a clear increase in food intake, nor do they display an obesity-like phenotype

Using indirect calorimetry, the energy balance of adult Snord116/115-deficient mice was assessed under two metabolically challenging conditions: exposure to a HFD and rapid temperature shifts from 22°C to 30°C, followed by a decrease from 30°C to 10°C. As shown in Figure 6A-left, when fed a HFD, both 3-4 months-old WT and mutant male mice (cohort 1) exhibited, as expected, increased body weight compared to a chow diet (ND). However, body weight gain was significantly less pronounced in mutants (Figure 6A-right), indicating a relative resistance to HFD-induced obesity. Such counterintuitive finding was previously described for Snord116-KO mice [32,37,38]. As anticipated from the nocturnal feeding behavior of rodents, food intake was higher during the dark phase in both genotypes and, consistent with caloric intake being adjusted in response to the higher energy content of a HFD, food intake was also reduced in both genotypes under HFD (Figure 6B). Thus, homeostatic response to increase calories appears intact in Snord116/115-KO mice. Remarkably, no overt increase in food intake was noticed in Snord116/115-KO mice when normalized for body weight (Figure 6B). This prompted us to examine two additional cohorts of adult males, aged 2 months (cohort 2) and 7-9 months (cohort 3). Again, these additional analyses did not reveal increase proportionated food intake (Figure 6G for cohort 2 and Supplementary Figure 8 for cohort 3). To increase statistical power, we then pooled measurements from cohorts 1, 2 and 3 (Supplementary Figure 9). As shown in Figure 6C-left, Snord116/115-KO mice displayed a slight but significant increase in food intake. However, this effect was no longer apparent without normalization for body weight (Figure 6C-right), warranting caution in its interpretation. Indeed, the appropriate method for normalizing energy intake has been debated for decades, including in recent studies on Snord116-KO mice [33]. Analysis of covariance (ANCOVA), as recommended for modeling energy balance [58], revealed that body weight rather than genotype affected food intake (Ancova genotype, p=0.087; weight p<0.001; Supplementary Figure 9). We consider it reasonable to assume that, under our experimental conditions, Snord116/115-KO mice do not develop robust hyperphagia.

**Figure 6-.**
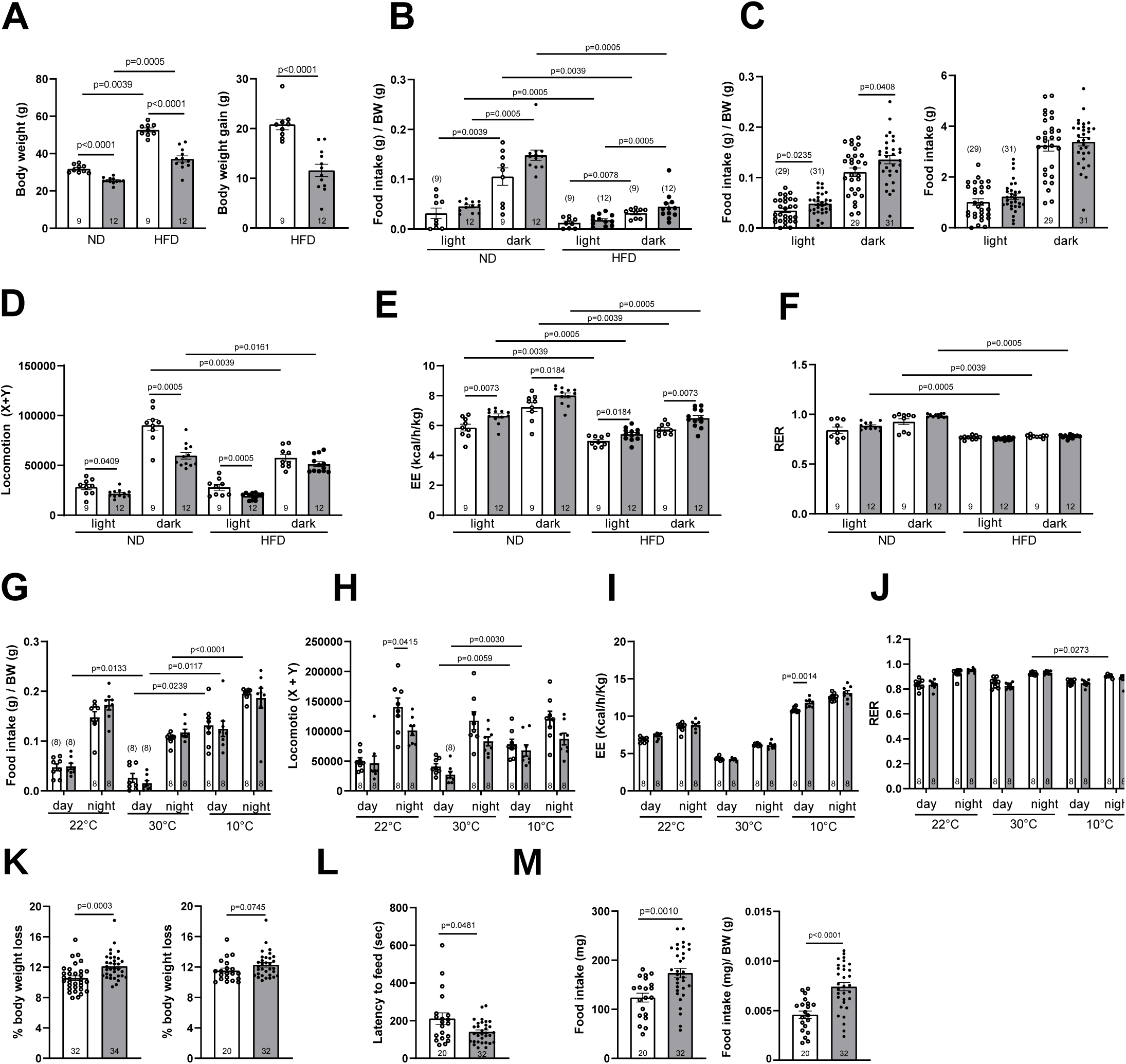
Adult Snord116/115-KO mice display altered energy balance without overt alterations in feeding behavior. **(A)** Snord116/115-KO mice display reduced body weight (left) under both normal diet (ND) and 14-week high-fat diet (HFD) conditions, together with decreased body weight gain upon HFD exposure (right) **(B-F)** Indirect calorimetry was performed in cohort 1 (males, 3-4 months age) maintained on ND and HFD (12 week) to assess food intake **(B-C)**, locomotor activity **(D)**, energy expenditure **(E)** and respiratory exchange ratio **(F)**. **(C)** Food intake measurements with (left) and without (right) normalization to body weight, using pooled data from cohorts 1, 2, and 3. (G-J) Indirect calorimetry was performed in cohort 2 (males, 2 months age), initially housed at 22°C, then shifted to thermoneutrality (30°C) for 24h, followed by cold exposure for 24h (10°C), to measure food intake **(G)**, locomotor activity **(H)**, energy expenditure **(I)** and respiratory exchange ratio **(J)**. **(K-M)** Novelty-Suppressed Feeding (NSF) test. **(K)** Snord-deficient mice exhibit increased body weight loss during the fasting period (left), a difference that is largely attenuated when the analysis is restricted to mice that lost at least 10% of their body weight (right). **(L)** Snord-deficient mice display a reduced latency to feed in an anxiogenic environment. **(M)** Snord-deficient mice show increased food intake, both in absolute terms (left) and when normalized to body weight (right) immediately upon return in their home-cages. The number of analyzed individuals or biological samples is indicated within each histogram or, where space is limited, in brackets above the histogram. White and grey bars represent WT and Snord116/115-KO mice, respectively.

Despite the absence of clear effects on food intake, Snord-deficient mice displayed reduced spontaneous locomotor activity during the active dark phase under ND, but not HFD in cohort 1 (Figure 6D) and, to a lesser extent, in cohort 3 (Supplementary figure 8). A similar defect was observed in cohort 2 during the dark phase at regular (22°C) temperature (Figure 6H). Energy expenditure (EE) was also increased in Snord116/115-KO mice from cohorts 1 and 3 under both dietary conditions (Figure 6E; Supplementary Figure 8), whereas no consistent increase was observed in cohort 2, which may reflect age-dependent differences (Figure 6I). The respiratory exchange ratio (RER) did not differ substantially between genotypes in any of the three cohorts, except during the light phase in cohort 3 (Supplementary figure 8). However, and as expected, RER values tended to decrease under HFD compared to ND, indicating a shift toward lipid oxidation (Figure 6F). Finally, response to rapid changes in ambient temperature remained largely unaltered, as evidenced by decreased food intake, locomotor activity and EE when the temperature was shifted from 22°C to 30°C, while conversely, these parameters increased during a shift from 30°C to 10°C (Figure 6G-J). Only a slight genotype effect was observed for EE during the light phase at 10°C.

### Snord116/115-KO mice display enhanced refeeding following the Novelty-Suppressed Feeding (NSF) test

Open field and Elevated Plus Maze (EPM) assays did not reveal increased anxiety-like behavior in Snord116/115-KO mice (Supplementary Figure 10). We therefore assessed refeeding behavior in an anxiogenic context using the Novelty-Suppressed Feeding (NSF) test, in which mice are placed in a brightly lit arena following a 24-hour fasting period. Three independent cohorts were studied several years apart (Supplementary Figure 11). In this paradigm, latency to initiate feeding is used as an index of anxiety-like behavior [59]. Remarkably, Snord116/115-KO mice exhibited greater body weight loss during the fasting period (Figure 6K-left). However, when the analysis was restricted to animals that had lost at least 10% of their body weight - a standard threshold for ensuring consistent NSF measurements - the difference between genotypes was less apparent (Figure 6K-right). Subsequent analyses were therefore conducted on this subset of mice, resulting in the exclusion of 2 mutant and 12 WT animals. Under these conditions, Snord116/115-KO mice displayed a reduced latency to feed (Figure 6L). As a control, food intake was measured immediately upon return to the home cage. Remarkably, Snord116/115-KO mice showed a marked increase in food consumption, regardless of whether intake was normalized to body weight (Figure 6M-left and 6M-right), indicating an enhanced refeeding response. This phenotype does not allow a clear distinction between reduced anxiety and increased hunger drive. It nonetheless points to an increased, context-dependent motivation to eat. Whether this reflects altered feeding behavior (*e.g*., true hyperphagia) or increased metabolic sensitivity to fasting remains unclear, as body weight loss tended to be greater in mutant mice.

### rRNA modification profile is not affected in the whole brain of Snord116/115-deficient mice

To gain deeper insight into the consequences of SNORD loss, the phylogenetic and phenotypic analyses described above were complemented by molecular studies. We previously showed that ribose-methylated nucleotides on rRNAs in hypothalamus remained unchanged in Snord115-deficient mice [46]. Building on more recent findings that SNORDs can also guide rRNA N4-acetylation [1,2], we then assessed whether Snord116 and Snord115 loss might - directly or indirectly - impact other rRNA modification types. This was achieved using Nanopore direct RNA sequencing (DRS), a recently developed technology that enables the detection of multiple RNA modification types within a single experiment at single-molecule resolution [60]. We then prepared native DRS libraries of WT and Snord116/115-KO whole brain in biological duplicates and sequenced them in a multiplexed FLO-MIN106 MinION flowcell. As assessed by TapeStation analysis, the steady-state levels of 18S and 28S rRNA in KO compared to WT were comparable, arguing against a major impairment of ribosome synthesis (supplementary Figure 12A). We then used two different RNA modification detection methods - EpiNano [61] and NanoConsensus [62] - to identify differentially modified sites. Our analysis revealed that neither of the two algorithms were able to identify reproducible differentially modified sites in KO *vs*. WT samples (Figures 7A-B). We should note that prior works benchmarking these tools have demonstrated that individual differentially modified sites identified only in a single replicate cannot be trusted as true positives. Nevertheless, we visually examined in IGV all the individual sites identified in a single replicate in 18S and 28S rRNAs (supplementary Figure 12B-C). This visual inspection did not suggest that the sites identified by an individual replicate would in fact be true differentially modified sites. Finally, as a positive control, we examined whether our pipeline would be able to identify differentially modified sites when comparing a different set of samples. To this end, we applied the Epinano based approach on WT and Snord90-KO knockout ([63]; supplementary Figure 12D). Our analysis revealed that position 18S:355 was identified as differentially modified upon the KO, which is the reported target site of Snord90, demonstrating that Epinano is robust for detecting differentially modified rRNA modifications. We therefore conclude that the lack of Snord116 and Snord115 does not lead to significant loss of any rRNA modification types.

**Figure 7-.**
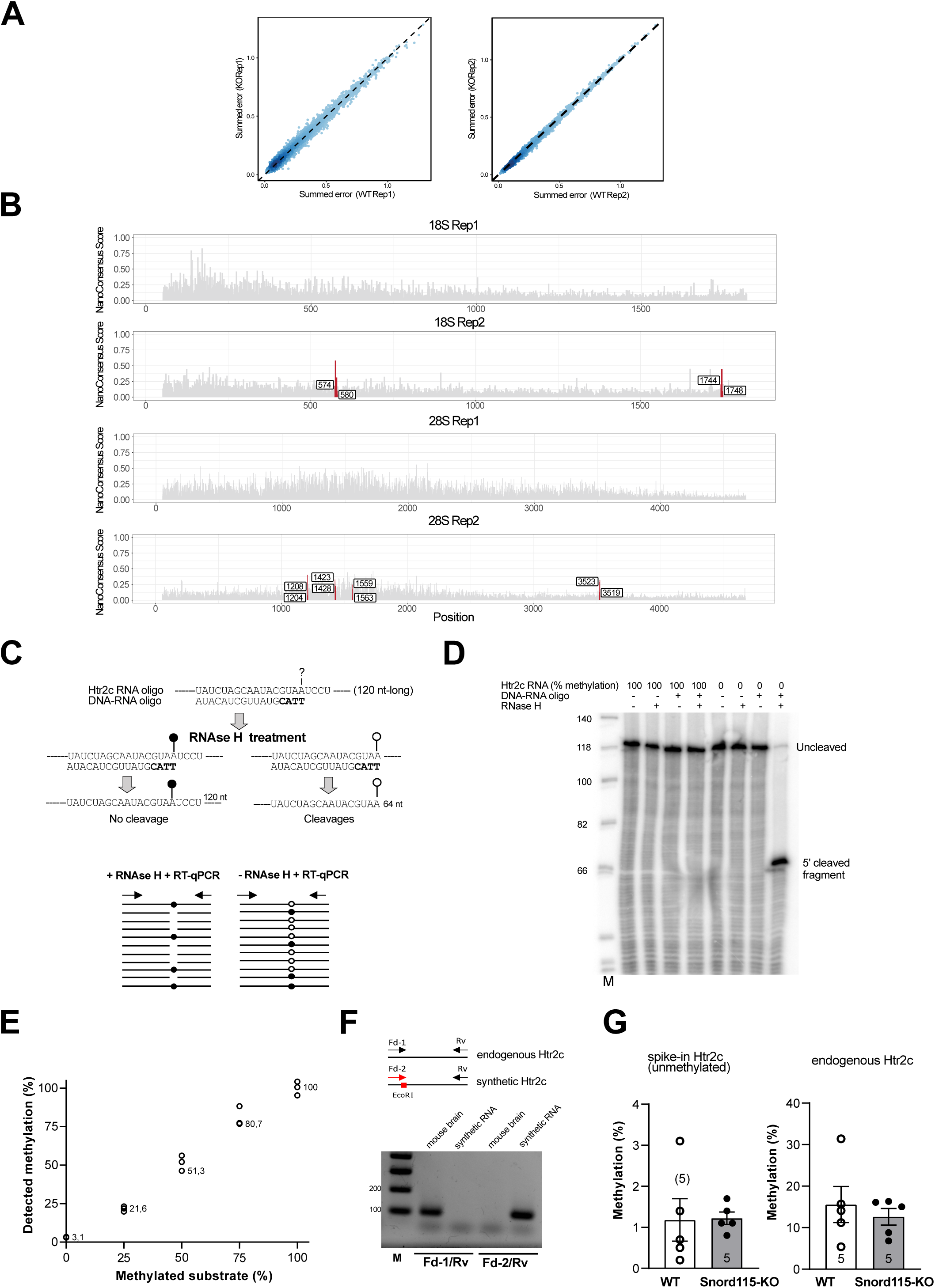
Snord loss does not alter the overall rRNA modification profile, nor the predicted 2′-O-methylation (Am) at Htr2c. **(A-B)** Nanopore direct-RNA sequencing shows no differences in rRNA modifications between WT and Snord116/115-KO mice. (A) Epinano summed error scatterplots for WT-SNORD116/115-KO comparisons (replicate 1, left; replicate 2, right) across all cytosolic rRNA molecules (5S, 5.8S, 18S, 28S). **(B)** NanoConsensus tracks for WT vs. Snord116/115-KO. NanoConsensus integrates signals from multiple nanopore RNA modification detection tools (EpiNano [61], Nanopolish [96], Tombo (bioRxiv. https://doi.org/10.1101/094672) and Nanocompore [97] by converting their outputs to Z-scores and identifying candidate sites within expanded 5-mer windows around predicted modification positions. Sites supported by at least two tools and exceeding the NanoConsensus score threshold are called as differentially modified and are highlighted in red. The rest of the sites are shown in gray. **(C-G)** Attempts to detect Snord115-mediated 2’-O methylation on endogenous Htr2c mRNA through Nm-VAQ. **(C)** Top: Schematic representation of RNA-DNA hybrids formed between a chimeric RNA-DNA oligonucleotide (DNA bases are in bold) and Htr2c region suspected to harbor a 2′-O-methylated adenosine targeted by Snord115 (indicated by “?”, see also Figure 1A). In this design, the target nucleotide is precisely positioned at the RNase H cleavage site, which is cleaved only when the substrate is unmethylated. Bottom: Following RNase H treatment, the uncleaved RNA is quantified by RT-qPCR using primers flanking the target residue (filled and open circles represent methylated and unmethylated adenosines, respectively). Untreated samples serve as input controls and the difference in Ct values (ΔCt) between treated and untreated conditions reflects the fraction of RNA protected from cleavage and thus 2′-O-methylation level. **(D)** The specificity and efficiency of RNase H cleavage were assessed by Northern blotting using an antisense oligonucleotide probe capable of detecting both the uncleaved precursor (120 nt, with or without Am) and the 5’ cleaved product (64 nt). **(E)** Correlation between expected and observed 2′-O-methylation ratios of an in vitro Htr2c substrate spiked into L929 total RNA. **(F)** Ethidium bromide-stained agarose gel following RT-PCR using two primer pairs designed to discriminate between endogenous Htr2c mRNA (mouse brain) and a synthetic Htr2c RNA carrying a 6-nt tag (EcoRI site) matching the 3′ end of the forward primer. **(G)** % of detected Am in whole brain from WT (white) and Snord115-KO mice (grey) on an exogenously added, unmethylated synthetic Htr2c RNA (left) and endogenously expressed Htr2c mRNA (right).

### Lack of detectable 2′-O-methylation at the Htr2c mRNA in vivo

Htr2c mRNA has been proposed as a target of SNORD115 for over two decades [12]. Yet, no attempts have been made to detect the predicted SNORD115-dependent ribose-methylated residue *in vivo,* the detection of which would provide definitive evidence for the formation of transient Snord115:Htr2c mRNA duplexes, which are otherwise difficult to capture biochemically in brain. We then employed Nm-VAQ, a recently-developed approach enable to quantify 2’-O-methylation at the single nucleotide level ([64], Figure 7C-top). We first assessed RNase H-mediated cleavages using two *in vitro* synthesized 120-nt-long Htr2c RNAs, one containing and one lacking a ribose-methylated adenosine at the position predicted to be targeted by Snord115. As shown in Figure 7D, Nm-VAQ proved efficient and specific, as the expected 64-nt-long 5’ cleaved product was detected only when the chimeric DNA-RNA oligonucleotide was annealed to the unmethylated Htr2c substrate in the presence of RNase H. We then assessed our ability to detect lower levels of Am by spiking 12.5 fmol of *in vitro* synthesized Htr2c RNA, containing defined proportions of Am (0, 25, 50, 75 and 100%) into total RNA prepared from L929 fibroblast cells which do not express htr2c mRNA. As shown in Figure 7E, the uncleaved-to-cleaved ratio, as inferred from ΔCt (cycle threshold) values obtained without and with RNase H treatment (Figure 7C-bottom), demonstrated correlation between observed and expected 2′-O-methylation levels. Having validated this approach, we applied Nm-VAQ to total RNA extracted from whole brains of WT and Snord115-KO mice (n = 5 per genotype). Each RNA preparation was also spiked with 5 fmol of the unmethylated synthetic Htr2c used as an internal negative control (*i.e*., positive control for RNAse H cleavages). To distinguish endogenous from spike-in RNA, we designed a forward PCR primer whose 3′ end annealed to an EcoRI (GAATTC) tag uniquely present in the synthetic RNA (Figure 7F, top). This strategy enabled the specific amplification of the exogenously added htrr2C RNA (Figure 7F, bottom). As shown in Figure 7G-left, and as expected, the uncleaved-to-cleaved ratio for spike-in htr2c RNA was minimal in both WT and KO samples (∼1%). By contrast, the methylation ratio for endogenous Htr2c mRNA reached ∼15% in WT samples but was, unexpectedly, comparable in KO samples (Figure 7G-right). To account for this unexpected observation, we hypothesize that the Snord115-independent methylation-like signal arises from A-to-I RNA editing. In particular, inosine incorporation within the RNA duplex, especially at the target site, may reduce the efficiency of RNase H-mediated cleavage, resulting in an apparent methylation signal. We therefore favor the more parsimonious explanation that, if Htr2c mRNA undergoes Snord115-dependent modification, the extent of 2′-O-methylation is very low.

### Snord116/115-KO mice exhibited modest alterations in gene expression profiles across the hypothalamus, cortex and cerebellum

The above-mentioned epitranscriptomic analyses were then complemented by gene expression profiling (mRNA-seq) of the hypothalamus at P1 (4 WT *vs.* 4 KO), P11 (6 WT *vs*. 6 KO) and adulthood (4 WT *vs.* 4 KO), as well as adult prefrontal cortex (6 KO *vs.* 6 WT) and cerebellum (5 KO *vs*. 5 WT). As shown in Figure 8A and Supplementary Table 2, only a relatively modest impact on the transcriptome was observed across the conditions, as reflected by the limited number of differentially expressed genes (DEGs), most with fold changes of less than two. In agreement with RT-qPCR analysis (Figure 3C), Ndn was found upregulated in both P1 and adult hypothalamic RNA samples. Remarkably, Dgkk and Htr2c, were also among the list of DEGs in the hypothalamus at P11 and their altered mRNA levels were further confirmed by RT-qPCR using different RNA preparations (Figure 8B). A similar effect on the steady-state level of Htr2c mRNA was not, however, observed in the adult hypothalamus of Snord116/115-KO mice (Figure 8B). We also examined alternative splicing of Htr2c pre-mRNA using RT-qPCR, as well as A-to-I RNA editing by quantifying A-to-G transitions in sequencing reads mapping to exon V of Htr2c. We did not detect major alterations in A-to-I RNA editing (Figure 8C) while a slight change in alternative mRNA splicing - a trend toward an increased RNA2/RNA1 ratio - was observed in adult hypothalamus of Snord116/115-deficient-mice (Figure 8B).

**Figure 8-.**
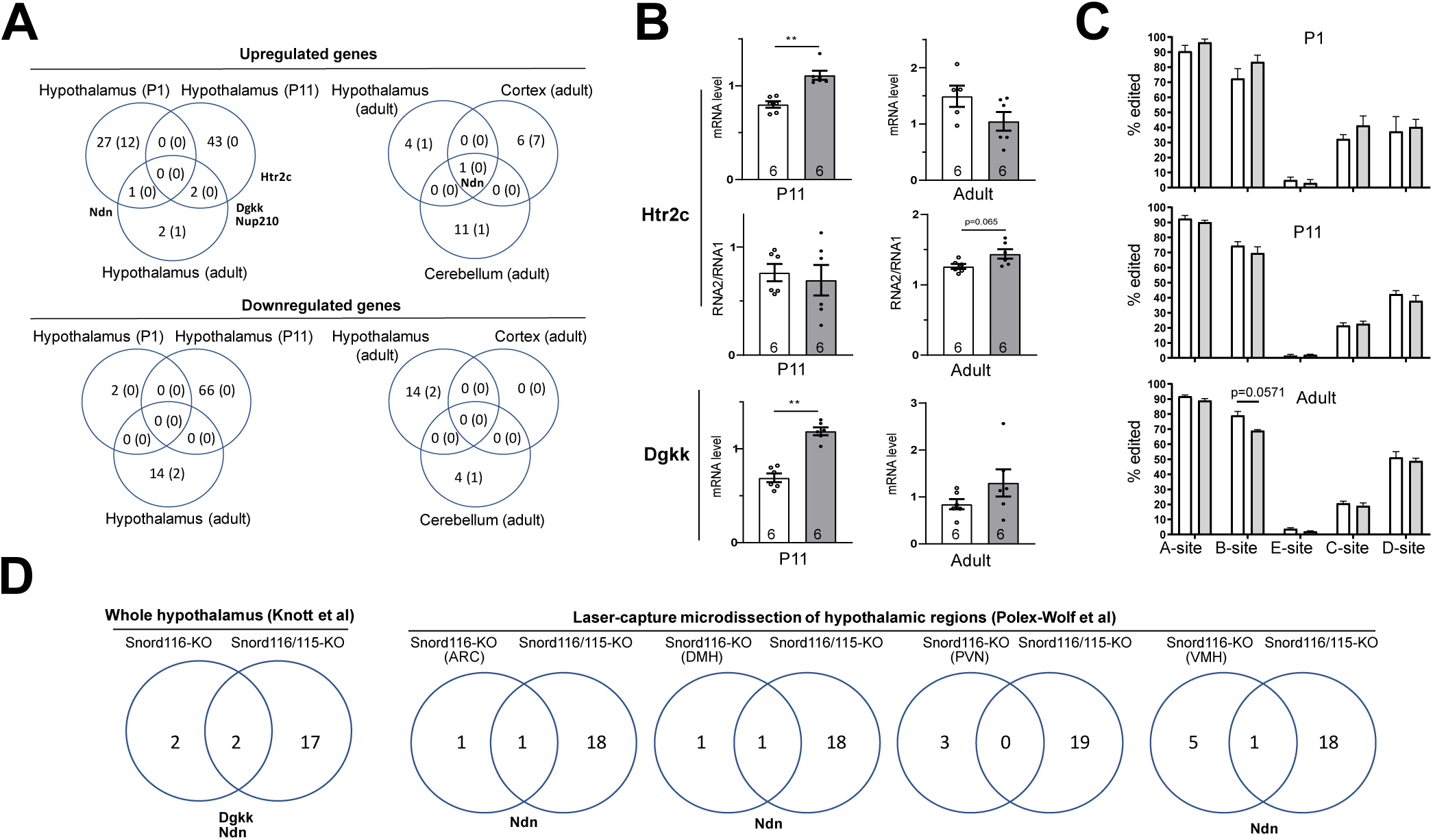
Modest impact of Snord116 and Snord115 loss on transcriptome output. **(A)** Venn diagram comparing DEGs (adj. p value < 0.05) identified in manually dissected brain regions of Snord116/115-KO vs. WT mice. The number of DEGs with a fold change greater than twofold is indicated in brackets. **(B)** Expression levels of Htr2c and Dgkk, as well as alternative splicing of Htr2c pre-mRNA (RNA1-to-RNA2 isoform ratio), were measured by RT-qPCR in the whole hypothalamus of Snord116/115-KO (grey) and WT (white) mice at P11 and in adulthood. RNA expression levels were normalized to those of Gapdh mRNA. **(C)** Percentage of A-to-I RNA editing events at individual sites (A-E) of Htr2c mRNA in the whole hypothalamus of Snord116/115-KO (grey) and WT (white) mice at P1, P11 and adulthood, as inferred from A-to-G mutations in sequencing reads. **(D)** Venn diagram comparing DEGs (adjusted p value < 0.05) identified in the hypothalamus of Snord116/115-KO mice with DEGs identified in two published transcriptomic analyses of Snord116-KO hypothalami. Note that sequencing reads mapping to the deleted region, and consequently downregulated in the KO samples, were not included among the DEGs (Supplementary Table 2).

We next compared DEGs identified in the adult hypothalamus of Snord116/115-KO mice with previously published datasets from adult SNORD116-KO mice, encompassing whole hypothalamus [38] or laser-captured hypothalamic nuclei [33]. To enable direct comparison, we re-analyzed all datasets using a consistent pipeline (Supplementary Table 2). As shown in Figure 8D, the hypothalamic transcriptomes of adult Snord116-KO mice remain also largely unaffected. This was also the case in other studies following SNORD116 loss in the perinatal hypothalamus [65] or in the whole brain [66]. Although the limited impact of SNORD deficiencies provided little insight, several gene-specific effects are nevertheless noteworthy. Snord116 deletion is associated with upregulation of Dgkk and Ndn in both Snord116-KO and Snord116/115-KO mice while Nup210 expression shows slight upregulation, yet reproducible in both Snord116/115-KO and Snord115-KO [46]. Note that Htr2c levels remain normal in cerebellum and prefrontal cortex while expression of Dgkk was too low to be detected in these two brain areas. Finally, expression of Nhlh2 mRNA, another putative Snord116 target involving non-canonical base-pairing interactions [16], remain unchanged across all Snord-deficient mice and conditions. Thus, the combined deletion of Snord116 and Snord115 exerts only modest molecular effects on transcriptome output and, consistent with perinatal and adult phenotypes, it does not result in greater alterations beyond those observed in Snord116-KO mice.

## Discussion

### “A rat is not a big mouse” - Imprinted domains as SNORD nurseries ?

One of the key insights from our comparative analyses is that PWS-associated SNORDs exhibit substantial birth-and-death dynamics, involving not only changes in gene copy number but also, perhaps more unexpectedly, sequence diversification. Indeed, such evolutionary changes are more extensive than previously anticipated, even within Euarchontoglires where these SNORDs have been proposed to contribute to neural development through adaptive processes [47]. This evolutionary divergence is exemplified by the relatively poor conservation of the single-copy SNORD107, SNORD64 and SNORD108 genes, in marked contrast to the strong conservation of canonical rRNA methylation-guiding SNORDs [47], thereby raising questions about the functional relevance and evolutionary selective pressures acting on some family members in certain mammalian species. For the tandemly repeated SNORD116 and SNORD115 genes, functional redundancy among related gene copies is expected to relax selective constraints, promoting sequence diversification and ultimately resulting in the pseudogenization, genetic drift, or neofunctionalization of individual copies [67,68]. In line with this evolutionary framework, SNORD115 genes have diversified along distinct evolutionary trajectories. While many SNORD115 copies are relatively well conserved both within and across species (e.g., *H. sapiens* and *M. musculus*), consistent with purifying selection preserving an essential regulatory function such as base pairing with Htr2c mRNA [12], other species display substantial sequence divergence. This is particularly evident in Lorisiform primates and the hedgehog *S. araneus*, where nucleotide differences in the AE preclude interaction with Htr2c mRNA. Whether this rapid evolution reflects pseudogenization, whereby SNORD115 is expressed but no longer functional, or neofunctionalization through the acquisition of novel RNA targets, remains an intriguing question. In some more extreme cases, such as in the genome of guinea pig (*C. porcellus*), SNORD115 gene cluster appears lacking, raising the possibility that its functions are either dispensable or compensated for by alternative mechanisms in some species. Regardless, our findings reveal that SNORD115 has followed a complex evolutionary trajectory. This is in marked contrast to SNORD116, which appears to be the most evolutionarily constrained of all PWS-derived SNORDs. At the same time, our study also revealed that SNORD109 appears more widespread than previously believed, including species-specific expansion giving rise to previously undescribed large arrays of repeated SNORD109 genes. This recalls observations at the imprinted Dlk1-Dio3 domain where members of the SNORD113/14 families expanded only in the *Rattus* genus [69]. Collectively, our findings suggest that the imprinted PWS region may be prone to gene duplication and sequence diversification, potentially enabling the rapid evolution of novel SNORD genes in specific mammalian lineages

SNORD gene loss or amplification raises questions about the stability of SNORD copy both within and between individuals and, by extension, whether copy number influences functionality through gene dosage effects. Accurately determining the copy number of repeated sequences is inherently challenging, as estimates are sensitive to sequencing technologies and genome assembly algorithms, as well as potentially to the biological origin of the analyzed samples. A similar issue was recently addressed in the Pig genome [70]. Taking these important technical considerations into account, our analyses of domesticated species suggest variability in SNORD116 - but not SNORD115 - copy number among individuals. Whether such “asymmetry” reflects domestication, genetic drift or merely genome assembly artifacts remain an open question that warrants further detailed investigation. Although an expression of concern has been issued, it is noticeable that one study in C57BL/6J mice also reported inter-individual variation in Snord116 copy number associated with craniofacial morphology features [71]. In human, small deletions affecting the number of SNORD115 copies were also detected in African (Yoruba) population [72], the relevance of which, if any, is totally unknown. Because the clinical penetrance of rare SNORD116-centered deletions in PWS is unknown, it would therefore be informative to investigate whether changes in SNORD116 copy number - or even complete loss - exists in unaffected individuals. Indeed, gene-phenotype relationships in PWS remain complex, as evidenced by studies showing that rare deletions simultaneously affecting the paternal alleles of MAGEL2, NDN and MKRN3 are associated with either absent or mild clinical manifestations [73–75] or, conversely, with a full PWS phenotype [76].

### Snord116 in the pre-weaning period - A role of postnatal leptin surge?

The fact that Snord116/115-deficient pups exhibit partially penetrant, genetic background-dependent neonatal survival rates is consistent with the “fine-tuner hypothesis” according to which SNORDs may promote robustness, especially in challenging biological contexts. Indeed, the transition to extrauterine life represents one of the most physiologically demanding events experienced by placental mammals. [77,78] and perinatal phenotypes are frequently documented in mouse models disrupted for imprinted genes [79], including those at the PWS locus [40]. Given the absence of perinatal phenotypes in Snord115-KO [46] and Ipw-KO mice (this study), the perinatal alterations in Snord116/115-KO mice are most likely attributable to Snord116 deficiency ([31,36,80], Wolff et al, bioRxiv, our study), although the possibility that certain specific defects arising from the combined deletion of Snord116, Ipw and Snord115 cannot be formally excluded.

Although rates of neonatal death upon Snord116 loss vary widely, from apparently not reported [37], 15% [31], 40% (this study) and up to 75% (Wolff et al, bioRxiv), perinatal growth impairment and metabolic disturbances represent the most consistently described abnormalities in Snord116-KO across independent studies (reviewed in [40]). Importantly, these abnormalities can be rescued by re-expressing only 15% of Snord116 from the normally maternally silenced allele [81]. As such, these Snord116-dependent postnatal developmental and metabolic alterations recapitulate the failure-to-thrive phenotype observed in infants with PWS. Although the precise contribution of Snord116 loss to these outcomes remains unclear, our observations in the FVB genetic background demonstrate that impaired metabolic status and growth retardation alone are insufficient to account for neonatal death. One possibility is that loss of Snord116, either directly or indirectly, disrupts *in utero* development and/or activation of the hypothalamic-autonomic nervous axis, with such defects becoming mostly evident shortly after birth. This may explain how the absence of Snord116 in the brain can, in a non-cell-autonomous manner, impact simultaneously multiple peripheral tissues, ultimately leading to fatal outcomes whose penetrance is likely influenced by intertwined stochastic factors (*e.g.,* timing of first suckling, maternal care, intra-litter competition…) and environmental variables (*e.g.,* ambient temperature, microbiota composition…).

In such a complex picture, the absence of the postnatal leptin surge at P11 in Snord116/115-KO or Snord116-KO pups which, to our knowledge, has never been reported in other PWS mouse models, is noteworthy. Indeed, the postnatal rise of leptin plays a critical role in the connectivity of hypothalamic feeding circuits [82]. Moreover, adverse *in utero* and/or perinatal events, including changes in leptin levels, can have long-term effects on metabolic and behavioral trajectories [83]. Thus, the postnatal metabolic and hormonal alterations experienced by Snord116-deficient pups imply that any adult phenotypes may originate, at least in part, from “perinatal programming.” Ongoing work is underway to test this hypothesis and to precisely define how Snord116 may shape hypothalamic circuits, metabolic homeostasis and the endocrine transitions during the pre-weaning period.

### Snord116-KO vs. Snord116/115-KO mice - What role for Snord115?

Contrary to expectations inferred from our initial redundancy hypothesis, the overall phenotype of Snord116/115-KO mice are not substantially more severe than those seen in Snord116-KO mice, leaving the role of Snord115 unresolved. This does not, of course, rule out functional interactions between Snord116 and Snord115, nor does it imply that Snord115 lacks a biological function. Indeed, our most recent findings indicate that, whereas Snord116-KO mice exhibit defects in sleep architecture, as previously reported [35], neither Snord115-KO nor, surprisingly, Snord116/115-KO mice display such sleep defects (Rivière et al., bioRxiv). These unanticipated findings suggest that deleting Snord115 genes normalizes somehow the impact of Snord116 deletion, perhaps revealing the context-dependent roles of Snord115. Our failure to observe robust overeating phenotype or anxiety in Snord116/115-KO mice, contrary to some report in Snord116-KO mice [30,32,37], may also reflect such counterintuitive interplay where “1+1=0”. In any case, our observations emphasizes the importance of carefully examining combined paternal allele deletions, as has recently been done for the paternally expressed Ndn and Magel2 protein-coding genes [84].

### Lost in (epi-)transcriptomics - What molecular defects drive phenotypes in Snord-deficient models ?

In this study, we adopted a SNORD-centered perspective, in which molecular and phenotypic defects are attributed to disruption of SNORD-mediated post-transcriptional processes. However, we acknowledge that SNORD loss through large constitutive deletions, as observed in mouse models and PWS patients, inevitably affects SNORD host gene transcripts including noncoding RNAs with snoRNA-derived ends [85] and the repetitive architecture of the locus [86], all of which may potentially contribute to the observed phenotypes. Therefore, elucidating the precise molecular mechanisms underlying the functions of murine Snord115 and Snord116 remains both a conceptual and technical challenge *in vivo*.

Our observations that rRNA modification profiles are unaffected in Snord116/115-KO mice provide direct experimental support to the notion that Snord115 and Snord116 are *bona fide* orphan SNORDs, prompting the obvious question of their RNA targets. Although the idea that SNORDs may interact with and potentially guide 2′-O-methylation of mRNAs has attracted increasing attention, robust and consistent evidence supporting this mechanism in tissues remains scarce [12,15,17,18,64,87–92]. This is largely due to the technical challenges associated with quantifying partial 2′-O-methylation at single-nucleotide resolution in low-abundance RNA substrates. Because “the absence of evidence does not constitute evidence of absence”, our inability to detect Am on Htr2c mRNA should therefore be interpreted with extreme caution, especially given that, under the “fine-tuner hypothesis”, this putative Snord115-guided methylation may occur at dynamic, low levels in a cell type-dependent manner. Alternatively, Snord115 regulatory role may involve antisense-based mechanisms independent of ribose methylation. It is, however, worth noting that SNORD116-dependent methylation of human DGKK mRNA has recently been mapped, although the stoichiometry was not reported [90].

A further major challenge is the small number of DEGs detected by mRNA-seq, including at ZT0 (Zeitgeber Time 0, corresponding to the onset of the light phase), as we recently documented (Rivière et al., bioRxiv). Although this could merely reflect cellular heterogeneity where the diversity of neuronal populations dilutes cell type-specific effects, Snord115 and Snord116 may also regulate protein output while inducing only modest changes in mRNA abundance that are difficult to detect consistently. In this context, where the transcriptome appears only minimally affected, the increase in mRNA levels of the two most convincing predicted Snord116/115-regulated targets, Dgkk and Htr2c, in the hypothalamus of Snord116/115-deficient pups is noteworthy and highlights the postnatal period as a key window for future investigations. Strikingly, and in contrast to *in vivo* mouse studies, numerous DEGs were recently detected across four different *in vitro* cultivated human neuronal cell models lacking either SNORD116 or SNORD115 [18,93,94]. However, these studies, each involving only a limited number of clones, and in some cases only a single clone, show little to no overlap in the top DEGs identified. Even more surprisingly, while FGF13 mRNA predicted to be targeted by SNORD116 displayed decreased steady state levels in human SNORD116-KO embryonic stem (H9 and CT2) cells differentiated into cortical neurons [18], its expression was upregulated in SNORD116-KO Lund Human Mesencephalic (LUHMES) cells differentiated into dopaminergic neurons [93]. In addition, SNORD116 loss in CT2 - but not H9 - neurons was associated with decreased expression of the maternally expressed ncRNA genes hosting the SNORD113/SNORD114 family [95] whereas, conversely, SNORD113 and SNORD114 were upregulated in SNORD116-KO iPSC-derived dopaminergic neurons ([94]; our observations), indicating very likely stochastic (epigenetic?) deregulation. In contrast, the expression of maternally expressed ncRNA genes at the imprinted Dlk1-Dio3 locus remain unchanged in Snord116/115-KO mice, at least in the brain regions we analyzed. Finally, neither DGKK, NHLH2 nor NDN showed consistent evidence of misexpression in these *in vitro* cultured neuronal SNORD-deficient human cell models.

In conclusion, by integrating evolutionary, biological, and molecular perspectives, our study underscores the limitations of inferring the functional significance of PWS-associated SNORDs from a single experimental system. Regardless of their putative RNA targets or mechanism of action (whether dependent on ribose methylation or not), our findings suggest that SNORD-mediated gene regulation is context-dependent and that the consequences of SNORD loss - whether arising naturally through evolution or following targeted deletion in mouse models - may be buffered by alternative compensatory mechanisms. Future studies should adopt comparative, multi-system approaches, including single-cell analyses, to elucidate how these recently and rapidly evolved, epigenetically regulated PWS-associated SNORD genes fine-tune brain function, phenotypic diversity, and disease susceptibility in the context of PWS. Such approaches may be particularly informative during the perinatal period, when SNORD deficiency appears to produce the most robust and reproducible phenotypic effects, offering a valuable opportunity to investigate the earliest stages of PWS pathogenesis.

---

Supplementary Table 1 - List of the mammalian genomes analyzed, including the genome assembly version and the sequencing technology used.

**Supplementary Figure 1 - (A)** Estimates of SNORD116 and SNORD115 copy numbers in the assembly of the genome of one Charolais heifer obtained with Oxford Nanopore, Pacific Biosciences (HiFi, CLR) and 10X Genomics linked-reads. White and grey bars represent SNORD116 and SNORD115, respectively. **(B)** Estimates of SNORD116 and SNORD115 copy numbers in 64 high-quality de novo genome assemblies of a French cattle breed generated using a standardized methodology and further refined through three iterative polishing steps using both PacBio CLR long-read and Illumina short-read data. **(C)** Estimates of SNORD116 and SNORD115 copy numbers in 16 high-quality, haplotype-resolved genomes generated using PacBio HiFi sequencing. Black and white bars represent SNORD116 and SNORD115, respectively.

**Supplementary Figure 2** - Sequence alignment of the RNA segment of Htr2c thought to be targeted by SNORD115 shows its conservation in placental (Eutherian) mammals but not in Metatherian or Prototherian. The positions of the five A-to-I editing sites are indicated, with the C-site corresponding to the site believed to be targeted for ribose methylation by SNORD115.

**Supplementary Figure 3** - Sequences of some newly identified SNORD115, SNORD115-like and SNORD116-like RNA species identified in this study.

**Supplementary Figure 4 – Canonical SNORD115 gene clusters are not detected in Eulipotyphla and Caviidae species (A)** Top: Identification of SNORD115-like gene cluster in Eulipotyphla. Left: simplified phylogenetic tree illustrating that SNORD115 loss is conserved in Eulipotyphla. Bottom: Dot plot analysis identified a large SNORD115-like gene cluster located between the SNORD116 cluster and the Ub3A gene in the European shrew (S. araneus). **(B)** Top: Northern blot detection of Snord116, Snord115 and Snord64 in total brain RNA from C. porcellus. Bottom: Dot plot analysis failed to detect a repeated array between the SNORD116 and the Ube3a gene in C. porcellus.

Supplementary Figure 5 - Sequences of detected SNORD64, SNORD107, SNORD108 and SNORD109 across mammalian species. Sequence logo alignments and multiple sequence alignments are also shown.

Supplementary Figure 6 - Snord116-deficient pups display altered circulating hormone levels compared to WT, as assessed at P1 and P11. The number of individuals analyzed is indicated within the bars. White and grey bars represent WT and Snord116/115-KO mice, respectively.

**Supplementary Figure 7 - (A)** Sequence alignment of exon-C of Ipw gene across selected Eutherian species. **(B)** Targeted genomic regions are indicated, with predicted Cas9 cleavage sites marked by arrowheads and PAM sequences underlined. Expected and observed breakpoint junction sequences (after cloning and Sanger sequencing of deletion PCR products) are shown.

**Supplementary Figure 8** - Indirect calorimetry of cohort 3 (Snord116/115-KO vs. WT) on a normal diet. **(A)** Body weight. **(B)** Food intake shown as absolute values (left) and normalized to body weight (right). **(C)** Locomotor activity. **(D)** Energy expenditure. **(E)** Respiratory exchange ratio (RER). The number of individuals analyzed is indicated within the bars or, where space is limited, in brackets above the histogram. White and grey bars represent WT and Snord116/115-KO mice, respectively.

**Supplementary Figure 9** - Indirect calorimetry of cohorts 1, 2 and 3 (pooled measurements) maintained on a normal diet. **(A)** Snord116/115-KO mice exhibit reduced body weight compared to WT. **(B)** Scatter plots showing the relationship between food intake and body weight during the light (left) and dark (right) phases. **(C)** Locomotor activity. **(D)** Energy expenditure. **(E)** Respiratory exchange ratio (RER). The number of individuals analyzed is indicated within the bars or, where space is limited, in brackets above the histogram. White and grey bars represent WT and Snord116/115-KO mice, respectively.

**Supplementary Figure 10** - Snord116/115-KO mice display a normal emotional response. **(A)** Elevated Plus Maze (EPM). Bars show the time spent in the open (anxiogenic) and closed arms of the apparatus during a 5 min session. **(B)** Open Field (OF). Bars show the distance travelled (left) and the time spent in the center (anxiogenic) of the arena (right) during a 10 min session. The number of individuals analyzed and sexes is indicated within and under the bars. White and grey bars represent WT and Snord116/115-KO mice, respectively. Under this design, a decrease in travelled distance was observed in mutant males but not in females.

**Supplementary Figure 11** - Snord116/115-KO mice display increased food intake in the Novelty-Suppressed Feeding (NSF) test. The detailed measurements of the three independent cohorts are shown. The pooled data are given in Figure 6. **Top:** Snord116/115-KO mice show a reduced latency to eat food placed in the bright (anxiogenic) center of the apparatus during a 10 min session. **Middle-Bottom:** Snord-deficient mice exhibit increased food intake upon returning to their home cages, both in absolute terms (middle) and normalized to body weight (bottom). The number of individuals analyzed is indicated within the bars. White and grey bars represent WT and Snord116/115-KO mice, respectively. Note: these are raw data, including mice that did not lose at least 10% of their initial body weight (see main text).

**Supplementary Figure 12** - Nanopore direct-RNA sequencing shows no differences in rRNA modifications between WT and Snord116/115-KO mice **(A-C)**. **(A)** Tapestation results for total RNA extracted from whole brain of WT and Snord116/115-KO. **(B)** IGV tracks for 18S sites shown as significantly different between WT and Snord116/115-KO replicate 2 samples. **(C)** IGV tracks for 28S sites shown as significantly different between WT and Snord116/115-KO replicate 2 samples. The allele frequency threshold used in all IGV tracks is 0.2. **(D)** Nanopore direct-RNA sequencing shows decreased 18S:Um355 methylation in Snord90-KO neurons. Epinano summed error scatterplots for the WT-Snord90-KO comparison across all cytosolic rRNA molecules (5S, 5.8S, 18S, 28S). Sites with Δ summed errors > 0.3 are labeled in red. Note that ribose methylations can influence the nanopore signal not only at the modified nucleotide itself but also at neighboring nucleotides (e.g., positions −2, −1, +1 and +2). This is illustrated here by the signal at U353, which lies two nucleotides upstream of the cognate SNORD90-mediated Um355 modification.

**Supplementary Table 2** - Lists of differentially expressed genes (DEGs) identified by mRNA-seq in Snord116/115-KO mice (this study) and Snord116-KO mice (publicly available datasets).

**Supplementary Table 3** - List of oligonucleotides used.

## Material and Methods

### Mice housing, breeding and genotyping

All animal procedures were conducted in accordance with the European Union Directive 2010/63/EU on the protection of animals used for scientific purposes and were approved by the local ethics committee for animal experimentation in Toulouse. They were registered with the French Ministry of Research of Education and Research (protocols APAFIS #13372-2018011214542827, #30068-202012181642973, #35424-2022021410167542, #36977-202204151214375 and #60796-2026042816437774). The animal housing facility met CNRS standards.

Mice were housed in standard plastic cages with access to water and food *ad libitum*, either regular chow diet or HFD diet (ssniff® DIO-60 kJ% fat (Lard)) in a temperature-controlled room, with a 12-h light-dark cycle at 22 ± 1 °C and 50% humidity. Given that Snord115 and Snord116 are expressed only from the paternal chromosome, Snord116/115-KO mice and their WT littermate controls were thus generated by crossing a heterozygous male with a WT female. Mice on a pure C57BL/6J or FVB/NRj genetic background were purchased from Charles River and Janvier Laboratories, respectively. The SnoRD116-Ipw-115 deletion was backcrossed to the C57BL/6J background for at least eight generations prior to behavioral and metabolic analyses. To generate C57BL/6J-FVB hybrid pups, FVB females were crossed with C57BL/6J males. Upon euthanasia, tissues were harvested, weighed, and immediately snap-frozen in liquid nitrogen before storage at −80°C. Blood was collected in Microvette CB 300 K2E tubes containing EDTA, centrifuged for 10 min at 10,000 rpm and 4°C, and the plasma was subsequently stored at −80°C. For genotyping routinely performed between P14 and P28 (weaning), genomic DNA was extracted using a Wizard SV Genomic DNA Purification System (Promega) and PCR was performed with FirePol Master Mix (Solis Biodyne). The sequences of PCR primers are given in the supplementary Table 3.

Breeding was performed overnight and females exhibiting a copulation plug the following morning were designated as embryonic day 0.5 (E0.5). Neonates were assessed immediately after birth (postnatal day 0, P0). To monitor perinatal death, which predominantly occurs within the first 48 hours, pup counts were recorded at P0, followed by twice-daily observations (9 a.m. and 6 p.m.) till P2, and once daily thereafter. All deceased pups that could be recovered (*i.e.,* those not subject to maternal cannibalism) were collected for genotyping. To minimize maternal stress, cages were only opened when necessary for observations. In the reduced litter size assay, designed to decrease competition among pups for maternal milk, litters were artificially reduced to three pups at P0 (n=10), while control litters containing 6-9 pups (n=6) were left unmanipulated. Pups were dissected at P10 at 5:00 p.m. for further analyses. To assess spontaneous activity in neonates, pups were placed 15 cm away from the nest, and the time taken to return was recorded with a maximum tolerated time of 120 seconds.

### Generation of Snord116/115-KO and Ipw-KO mouse models

The double KO mouse model deficient for both Snord116 and Snord115 genes was generated using the same CRISPR-Cas9 strategy used to generate the single Snord115-KO line, as previously described in [46]. Briefly, *in vitro* transcribed Cas9 encoding mRNA (GeneArt CRISPR nuclease mRNA, Invitrogen) and two guide RNAs (MEGAshortscript T7 Transcription Kit; Ambion) designed to base-pair upstream and downstream of Snord116-Ipw-Snord115 gene array (crispr.mit.edu tool) were coo-injected into zygotes isolated from (C57BL/6JxCBA) F1 mice. Briefly, prepubertal (C57BL/6J×CBA) F1 female mice (4-6 weeks old) were superovulated by intraperitoneal injection of 5 IU pregnant mare serum (PMS), followed 46 hours later by 5 IU human chorionic gonadotropin (hCG), and then mated overnight with adult (C57BL/6J×CBA) F1 males. Zygotes were collected 22 hours after hCG injection. Microinjections were performed using an Eppendorf microinjector. A few picoliters of Cas9 mRNA (100 ng/ml) and *in vitro-*transcribed sgRNAs (25 ng/ml each) were co-injected into the cytoplasm and pronuclei of the zygotes. Injected embryos were cultured in M16 medium until the blastocyst stage to assess cleavage efficiency or were transferred directly into the oviducts of 0.5-day post-coitum (p.c.) pseudopregnant recipient female mice (approximately 20 embryos per recipient). The Ipw-KO mouse model was generated by electroporation of frozen C57BL/6JRj zygotes obtained from Janvier Laboratory (SE-ZYG-CJP) using CRISPR/Cas9 ribonucleoprotein (RNP) complexes. The RNP complexes consisted of Cas9 protein (IDT, 1081060) and two sgRNAs designed to target sequences upstream and downstream of the IPW locus (designed using the CRISPOR tool and synthesized by IDT). Briefly, zygotes were thawed and incubated in M2 medium for a couple of hours. They were then electroporated with the RNP complexes (Cas9: 500 ng/µl; sgRNA: 100 ng/µl) using a NEPA21 electroporator (NEPAGENE). Following electroporation, zygotes were cultured in M16 medium until the blastocyst stage to assess cleavage efficiency, or directly transferred into the oviductal ampullae of 0.5-day p.c. pseudopregnant recipient female mice (approximately 20 embryos per female).

### Search and sequence analyses of PWS-associated SNORD genes

The identification and enumeration of PWS-associated SNORD genes were performed using BLAST searches with default parameters (word size = 7) against genome assemblies available in the NCBI database, with priority given to high-quality reference assemblies generated using long-read sequencing technologies (PacBio, Oxford Nanopore). In most cases, results were further verified by manual curation. The full list of analyzed genomes is provided in Supplementary Table 1. Multiple sequence alignments were performed using MultAlin (http://multalin.toulouse.inra.fr/multalin/) and sequence logos were created using WebLogo (https://weblogo.threeplusone.com/create.cgi). Percent identity between SNORD copies was estimated using the Alignment Quality Scorer (https://punnettsquare.org/alignment-quality-scorer/). Species phylogenetic relationships were obtained from https://timetree.org/ and mammalian species classification was based on the NCBI Taxonomy database (https://www.ncbi.nlm.nih.gov/taxonomy).

### Metabolic Parameter Analysis

Blood glucose levels were measured using a hand-held glucometer (Accu-Chek Guide, Roche Diagnostics). Samples below the detection limit (10 mg/dL) were arbitrarily assigned a value of 9 mg/dL. Ketone bodies were measured using a hand-held ketone meter (FreeStyle Optium Neo, Abbott). Hepatic glycogen content was determined using a direct enzymatic assay according to the manufacturer’s instructions (Glycogen Assay Kit II, Colorimetric; Abcam, ab169558). Serum leptin and IGF-1 levels were measured in pups at different developmental stages using the Mouse Leptin ELISA Kit (Millipore) and the Mouse IGF-1 Quantikine ELISA Kit (R&D Systems), respectively. Hormone quantification was also performed using the MILLIPLEX MAP Mouse Metabolic Disease Panel (Merck Millipore) and analyzed on a Luminex 100 IS system at the IBISA-labeled Transgenesis, Zootechny and Functional Exploration Core Facility of Toulouse (Anexplo/GenoToul).

### RNA Extraction, RT-qPCR and Northern Blot

Total RNA was extracted using TRI Reagent (Euromedex) and treated with RNase-free RQ1 DNase (Promega) and Proteinase K (Sigma) to remove DNA and proteins. For mRNA analysis, whole-cell RNA was reverse transcribed with random hexamer primers using the GoScript Reverse Transcriptase Kit (Promega) at 42°C for 60 min. Gene expression was quantified by RT-qPCR using IQ Custom SYBR Green Supermix (Bio-Rad). Relative expression levels were determined using the standard curve method, with technical triplicates for each data point. For Northern blot analysis, 1-10 µg of total RNA was fractionated on a 6% polyacrylamide/7 M urea denaturing gel (PAGE) and electrotransferred onto a nylon membrane (Amersham Hybond-N, GE Healthcare), followed by UV crosslinking (Stratalinker). Membranes were hybridized overnight at 50°C with 5′-end ^32^P-labeled DNA oligonucleotide probes in hybridization buffer (5X SSPE, 5X Denhardt’s, 1% SDS, 150 µg/ml yeast tRNA) and washed twice in 0.1× SSPE, 0.1% SDS for 15 min at room temperature. Radioactive signals were detected using a Typhoon Biomolecular Imager (Amersham) and quantified with Multi Gauge V3.0 software. Primer sequences are listed in Supplementary Table 3.

### Indirect calorimetry and body composition

Three cohorts of adult mice were subjected to indirect calorimetry. Cohort 1 was used to assess the impact of a high-fat diet (HFD). Indirect calorimetry was first performed in 3-4-month-old male mice maintained on a chow diet, followed by a second assessment after 14 weeks of HFD. Cohort 2 (males aged 2 months) was used to evaluate temperature homeostasis. These mice were initially monitored at 22°C, then transferred to 30°C for 24 hours, followed by 10°C for 24 hours. Cohort 3 consisted of slightly older males aged 7-9 months maintained on chow diet. Indirect calorimetry was performed after 24 hours of acclimatization in individual cages. Mice that consumed less than 0.5 g during a 24-hour period were excluded from the analyses. Oxygen consumption (VO), carbon dioxide production (VCO), food and water intake, and spontaneous locomotor activity were measured in individual mice at 15-minute intervals over a 24-hour period at a constant temperature of 22°C using the Phenomaster system (TSE Systems, Bad Homburg v.d.H., Germany). The Respiratory Exchange Ratio (RER) and Energy Expenditure (EE) were calculated. [RER] = Vco2/Vo2 and [EE] in kcal/day/kg0^.75^ = (CVo2 x Vo2 + CVco2 x Vco2)/1000, with CVo2=3.941 et CVco2=1.106. Ambulatory activities of the mice were monitored by infrared photocell beam interruption. For assessment of body composition, total body fat and lean mass were measured in mice maintained on a chow diet. Mice were placed in a clear plastic holder without anesthesia or sedation and inserted into a Minispec whole-body composition analyzer (Bruker, LF50) following the manufacturer’s instructions.

### Behavioral Procedures

Behavioral analyses were performed blind during the light phase (8:30 a.m. to 1:00 p.m.). Anxiety-like phenotypes were assessed using the Elevated Plus Maze (EPM) and the Open Field (OF) tests. For the EPM, WT and Snord116/115-KO male mice (cohort 7; 3.5-4 months old) were placed in a plus-shaped maze consisting of two closed and two open arms (30×5×35 cm) extending from a central platform (5×5 cm). The apparatus was elevated 50 cm above the floor. Each trial began with the mouse placed in the central zone. During the first 5 min, the number of entries into the open and closed arms and the time spent in each arm were recorded manually using Ethoc software (CRCA, France). For the OF test, WT and Snord116/115-KO male and female mice (cohort 8; 4-6 months old) were assessed in a circular arena (diameter 40 cm; height 30 cm) located in a featureless room illuminated by white light (20 lx). Locomotor activity and exploratory behavior were monitored using Ethovision XT (Noldus), and the time spent in the center versus the periphery of the arena was scored during a 10-minute session. Feeding behavior under an anxiogenic environment was evaluated using the Novelty-Suppressed Feeding (NSF) test. Three independent cohorts of male mice (cohorts, 4, 5 and 6; 4-4,5 months) were tested (see also supplementary Figure 11). Mice were food-deprived for 24 hours prior to the test. Each mouse was placed individually in a 50×50×20 cm box filled with bedding and illuminated by a 70 W lamp. A single food pellet was positioned on white paper in the center of the arena. Latency to approach and consume the pellet within a 10-minute session was recorded as a measure of motivation (i.e., a longer latency is interpreted as a measure of increased anxiety-like behavioral). Mice that did not eat during the testing period were excluded from the analysis. Immediately after testing, each mouse was returned to its home cage, and the amount of food consumed over 5 minutes was measured to control for potential changes in appetite.

### mRNA-seq analyses

Library preparation (poly(A)+ selection) and Illumina sequencing (2×150 bp paired-end) were performed by Genewiz Azenta. Quality control of raw sequencing files (FASTQ) was assessed using FastQC (https://www.bioinformatics.babraham.ac.uk/projects/fastqc/). Reads were aligned to the mouse reference genome (GRCm39) and quantified against the GENCODE annotation (GRCm39 GTF) using the STAR aligner (STAR_2.5.4a). The raw count matrix was subsequently filtered using the Bioconductor HTSFilter package (HTSFilter_1.42.0) with parameters s.min = 1 and s.max = 50. Differential gene expression analysis, including normalization as well as estimation of log2 fold changes and adjusted p-values, was performed using the Bioconductor DESeq2 package (v1.42.0), with each mutant condition (KO) analyzed independently against the WT group. Raw sequencing data have been deposited in the Sequence Read Archive (SRA) under accession number GSE329283.

### Direct RNA nanopore library preparation

All nanopore sequencing runs were performed using the MinION sequencer (flow cell type: FLO-MIN106, sequencing kit: SQK-RNA002). DNase treatment was performed by mixing 2 µg of total RNA per sample with 5 µl of DNase buffer (Life Technologies, #AM2239), 1 µl of RNaseOUT (Invitrogen, #10777019) and 2 µl of Turbo DNase (Life Technologies, #AM2239) in a total volume of 50 µL. The digestion was done by incubating the samples for 10 minutes at 37 °C. The RNA was cleaned using RNA XP beads (Beckman Coulter, #A63987), resuspended in 20 µl of RNase-free water and quantified using NanoDrop spectrophotometer and Qubit™. Poly(A)-tailing was performed by mixing 1 µg of DNase-treated total RNA with the following reagents: 2 µl of poly(A)-tailing buffer (NEB, #B0276SVIAL), 2 µl of 10 mM ATP (NEB, #B0756AVIAL), 1 µl of E. coli Poly(A) polymerase (NEB, #M0276SVIAL), 0.5 µl of SUPERase•In™ (Life Technologies, #AM2694) in a total volume of 20 µl. The reaction mixtures were incubated for 15 minutes at 37 °C. Poly(A)-tailed RNA was then cleaned using Zymo RNA Clean and Concentrator (Zymo, #R1015) and quantified using NanoDrop spectrophotometer and Qubit™. RNA profiles of DNase-treated and poly(A)-tailed RNA were checked using the Tapestation™. Four samples per flow cell were barcoded by ligating custom-made oligonucleotides by mixing the following reagents: 250 ng of poly(A)-tailed total RNA per sample, 1.5 µl NEBNext Quick Ligation Reaction Buffer, 0.75 µl T4 DNA Ligase, 0.5 µl RNaseOUT (Invitrogen, 10777019) and 0.5 µl of pre-annealed custom-made oligonucleotides. The reaction mixtures were mixed by pipetting and incubated for 10 minutes at room temperature. After the ligation, reverse transcription was performed directly by adding 6.5 µl of RNase-free water, 1 µl of 10 mM dNTPs (Thermo Fisher, #18427013), 4 µL of Maxima RT buffer and 1 µl of Maxima reverse transcriptase (Life Technologies, #EP0751). The samples were mixed by pipetting and incubated for 30 min at 60 °C. The RNA-cDNA hybrids were then cleaned using RNA XP beads (Beckman Coulter, #A63987), resuspended in 5 µl of RNase-free water and placed into a clean 1.5 mL Eppendorf DNA LoBind tube (Eppendorf, 30108051). RNA and cDNA were quantified using Qubit™. 50 ng of each sample was pooled together and mixed with 8 µl of NEBNext Quick Ligation Reaction Buffer (NEB, #B6058S), 6 µl of RNA adaptor (RMX, ONT), 3 µl of RNase-free water, and 3 µl of T4 DNA ligase (NEB, #M2200L) in a total volume of 20 µl. The reaction mixture was mixed by pipetting and incubated for 10 minutes at room temperature. The library was cleaned using RNA XP beads (Beckman Coulter, #A63987), with Wash Buffer (WSB, ONT) in the washing steps, for a total of two washes. The beads were then eluted in 21 µl of Elution Buffer (EB, ONT). RNA and cDNA were quantified using Qubit™. The library was mixed with 17.5 µl of RNase-free water and 37.5 µl RRB buffer (ONT) and loaded onto the flow cell.

### Nanopore DRS data analysis

Raw fast5 files were basecalled, demultiplexed and mapped using the mop_preprocess module of MoP2 [98]. The demultiplexing was performed using DeePlexiCon [99]. Fasta files used as a reference for rRNA mapping were retrieved from GenBank, are correspond to the following annotations: NC_000074.6 (5S rRNA), NR_003280.2 (5.8S rRNA), NR_003278.3 (18S rRNA) and NR_003279.1 (28S rRNA). The mop_preprocess output was used as input for mop_mod analysis, and the mop_mod output was used for the mop_consensus analysis [62]. Epinano was used to obtain summed errors calculated at per-site resolution, for each sample. Briefly, the summed error (SE) values were calculated by summing the mismatch, deletion and insertion frequencies at the per-site level for each position in all rRNA molecules, as the frequency of basecalling errors was reported to correlate with the modification status of a given site [61,100]. Raw sequencing data have been deposited in the European Nucleotide Archive (ENA) under accession number PRJEB114655.

### RNase H-mediated cleavages and Nm-VAQ assay

Two 120-nt-long synthetic Htr2c RNA oligonucleotides (with or without 2′-O-methylation, Am) and a chimeric (2′-O-methyl/DNA) probe were synthesized by Integrated DNA Technologies (IDT). A 6-nt GAATTC (EcoRI) tag was inserted into both the methylated and unmethylated Htr2c RNAs, enabling their discrimination from endogenous Htr2c mRNA. Sequences are provided in Supplementary Table 3. Briefly, 2.5 pmol of Htr2c RNA oligonucleotides (with or without Am) were incubated in the presence or absence of 25 pmol of chimeric probe, heated at 95 °C for 2 min and gradually cooled to 22 °C at a rate of 0.1 °C/s, followed by a 5 min incubation at this temperature to allow duplex formation. Samples were then divided into two equal aliquots. One aliquot (5 µl) was incubated with RNase H (5 U; New England Biolabs, M0297) in 1× RNase H Reaction Buffer while the second aliquot was processed in parallel under identical conditions without RNase H. Reactions were carried out at 37 °C for 30 min and terminated by heating at 90 °C for 10 min. Following phenol/chloroform extraction and ethanol precipitation, cleavage products were analyzed by Northern blot using a 5′-end ³²P-labeled DNA oligonucleotide probe (sequence provided in Supplementary Table 3). Methylated and unmethylated Htr2c RNA oligonucleotides were mixed to obtain RNA substrates with defined methylation levels (0, 25, 50, 75 and 100%). For each condition, 0.0125 pmol of substrate was incubated with 2.5 µg of total RNA from L929 cells and 12.5 pmol of chimeric probe and the RNase H cleavage assay was carried out as described above. After phenol/chloroform extraction and ethanol precipitation, one-tenth of the products was reverse-transcribed using random hexamer primers and GoScript Reverse Transcriptase Kit (Promega) at 42 °C for 60 min. qPCR was performed using 1 µl of 10-fold diluted cDNA with iQ Custom SYBR Green Supermix (Bio-Rad). Each sample was analyzed in triplicate. ΔCt (cycle threshold) was calculated as follows: ΔCt = Ct (RNase H-treated sample) - Ct (RNase H minus treated samples and the percentage of methylation was calculated as follows: % methylation = 100 × 2^(−ΔCt). To determine the methylation status of endogenously expressed Htr2c transcripts, 6 µg of total RNA prepared from whole mouse brain (WT and KO, n = 5 per genotype) was incubated with 50 pmol of chimeric probe and spiked with 5 fmol of an unmethylated synthetic Htr2c RNA oligonucleotide and processed as described above.

### Statistical Analysis

Statistical analyses were performed using GraphPad Prism (version 8.4.3) and Jamovi (version 2.6.44). Data are presented as mean ± SEM. Normality and homogeneity of variances were assessed using the Shapiro-Wilk and Levene tests, respectively. For comparisons between two groups, a Student’s t-test was used when assumptions of normality and equal variances were met; otherwise, the non-parametric Mann-Whitney test was applied. To assess the effect of high-fat diet (HFD), within-genotype comparisons between normal diet (ND) and HFD were performed using the Wilcoxon signed-rank test. Analysis of covariance (ANCOVA) was used where indicated to adjust for potential covariates. For the open-field test, data were analyzed using two-way ANOVA followed by Sidak’s multiple comparisons test.

## Author Declaration

All authors have approved the final version of the manuscript and confirm that it is not under consideration for publication elsewhere. The authors declare that they have no conflict of interest.

## Declaration of generative AI-assisted technologies

The authors used OpenAI’s ChatGPT solely to correct grammatical errors and improve the clarity of the English language. The authors reviewed and edited all AI-assisted text and take full responsibility for the final content. No AI tools were used to conduct bibliography searches, generate scientific ideas or hypotheses, interpret data or draw conclusions.

## Acknowledgments

We are grateful to S. Labialle (IMoPA-CNRS, Nancy) for careful and critical reading of the manuscript; to R. Feil (IGMM-CNRS, Montpellier) for providing the JF1 mice; to B. Guiard (CRCA-CNRS, Toulouse) for helpful discussions regarding the use and interpretation of NSF; to V. Sorin and M. Boussaha (INRAE, AgroParisTech) for discussions on the technical aspects of Bos taurus genome assembly; and to Q. Chen (University of Macau) for discussions about Nm-VAQ. We also thank the ABC Facility of ANEXPLO (Toulouse), as well as the CBI facilities, notably the Big-A platform for NGS analyses and the Mouse Behavioral Core (MBC) for access to behavioral equipment. Finally, JC would like to acknowledge a colleague - whose name, unfortunately, he can no longer recall - for reminding him that “a rat is not a big mouse.”

## Fundings

This work was supported by the Centre National de la Recherche Scientifique (CNRS), the University of Toulouse, the Agence Nationale de la Recherche (ANR-22- CE12-0020 to J.C.) and the Foundation for Prader-Willi Research (to J.C.). EMN received funding from the project PID2024-157315NB-I00 and the European Union’s Horizon Europe through the European Research Council (ERC) under the grant agreement number 101042103 and 101187456. The funders had no role in study design, data collection or interpretation.

## Notes

### Competing Interest Statement

The authors have declared no competing interest.

