## Supplementary figures and images for "Evolutionary and biological insights into the paternally expressed Snord116-Ipw-Snord115 gene array at the imprinted Prader-Willi syndrome domain"

### Supplementary Figure 1

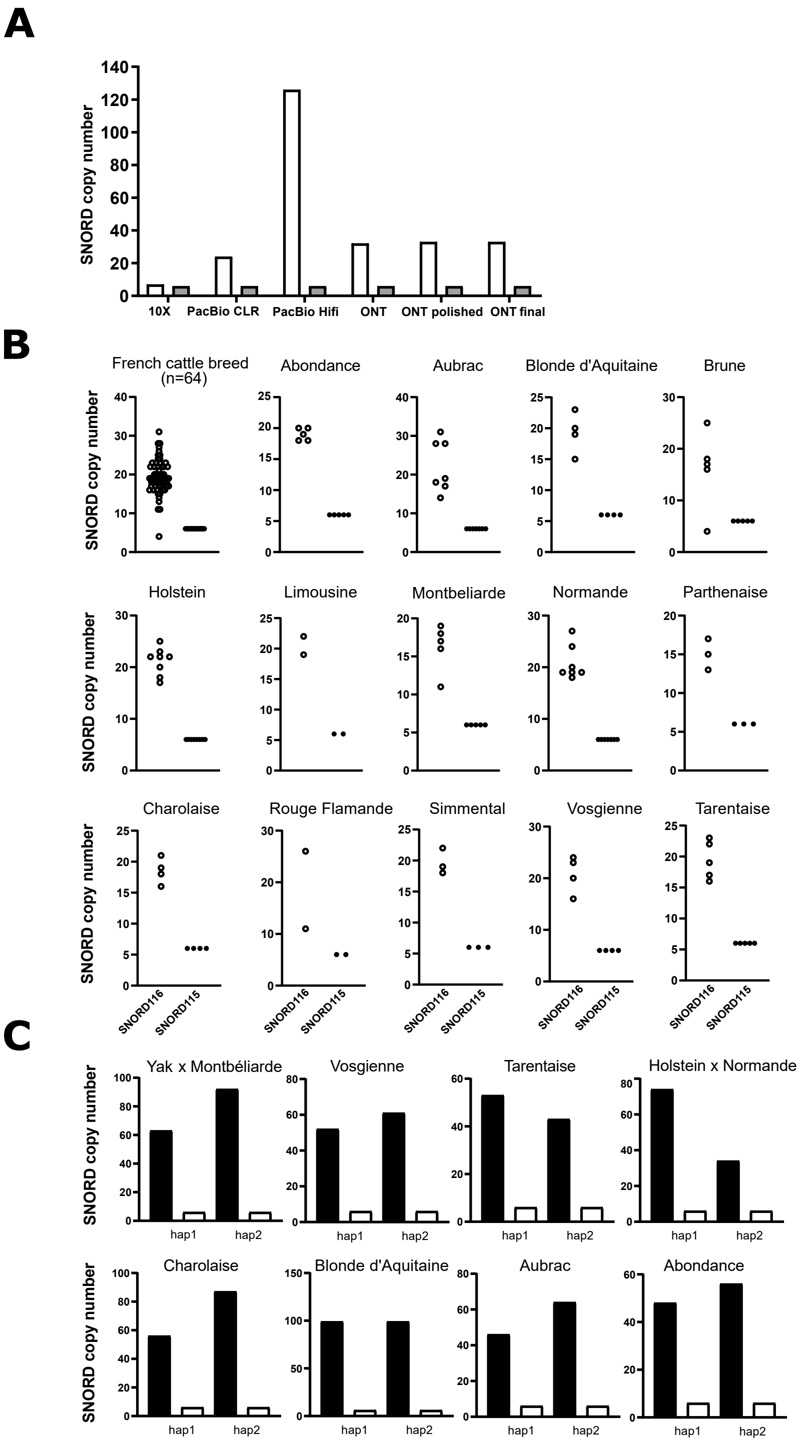


Supplementary Figure 1 - Marty *et al*

### Supplementary Figure 2

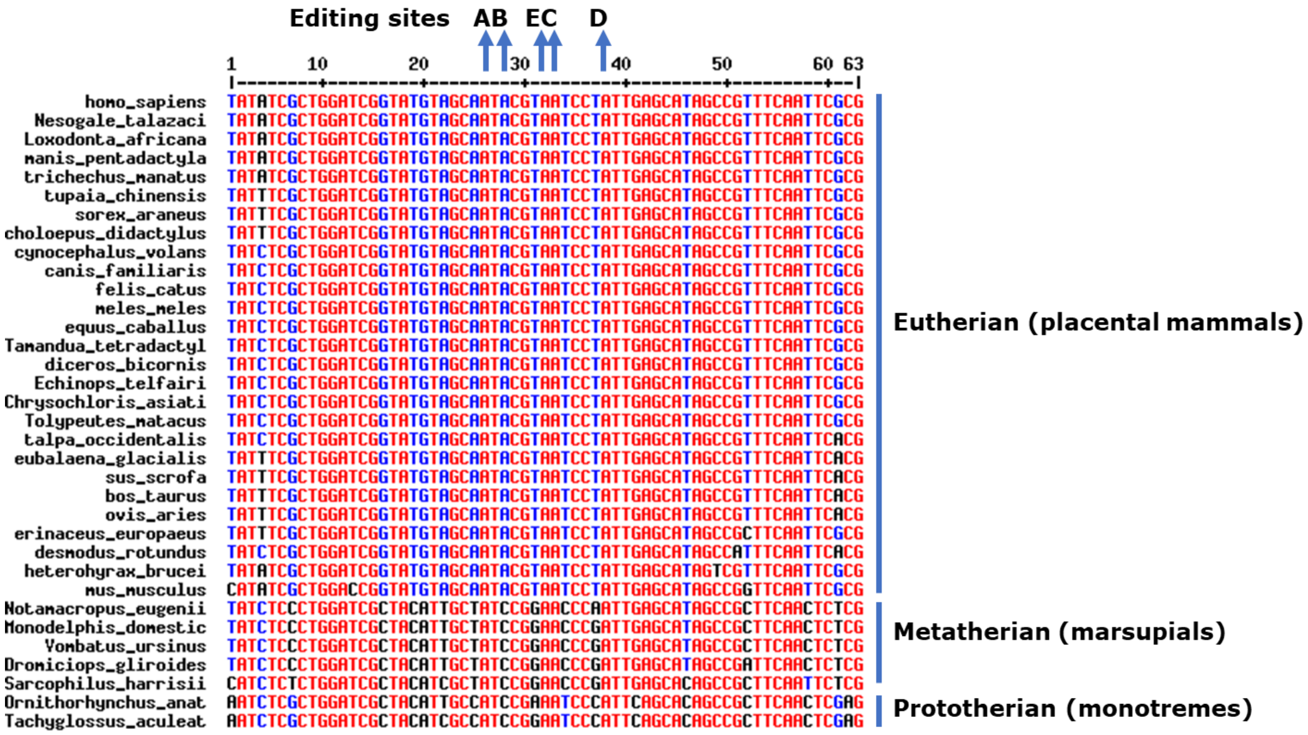


Supplementary Figure 2, Marty *et al*

### Supplementary Figure 4

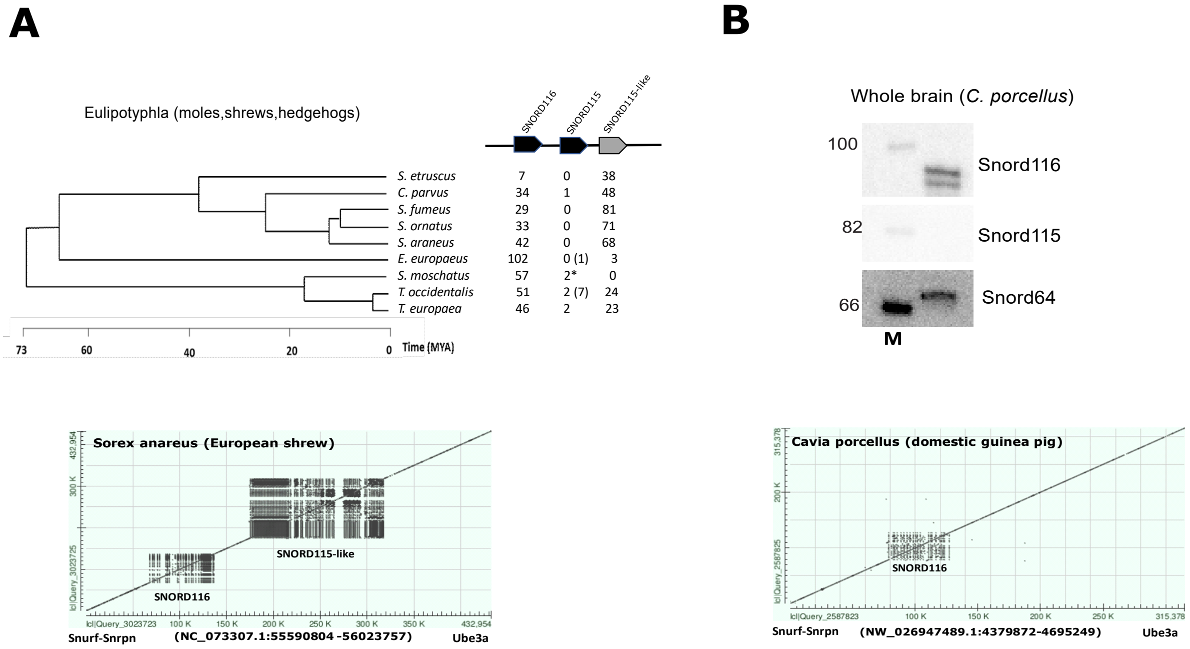


Supplementary Figure 4 - Marty *et al*

### Supplementary Figure 6

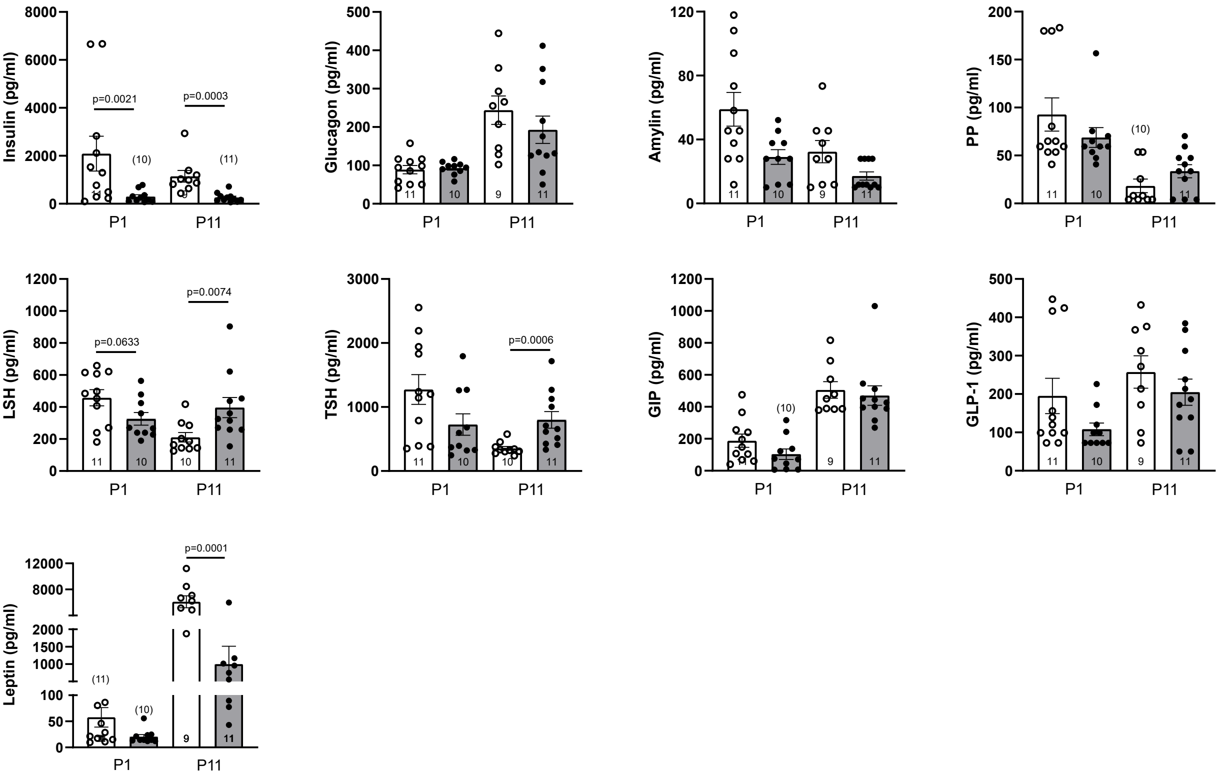


Supplementary Figure 6 - Marty *et al*

### Supplementary Figure 7

**A**

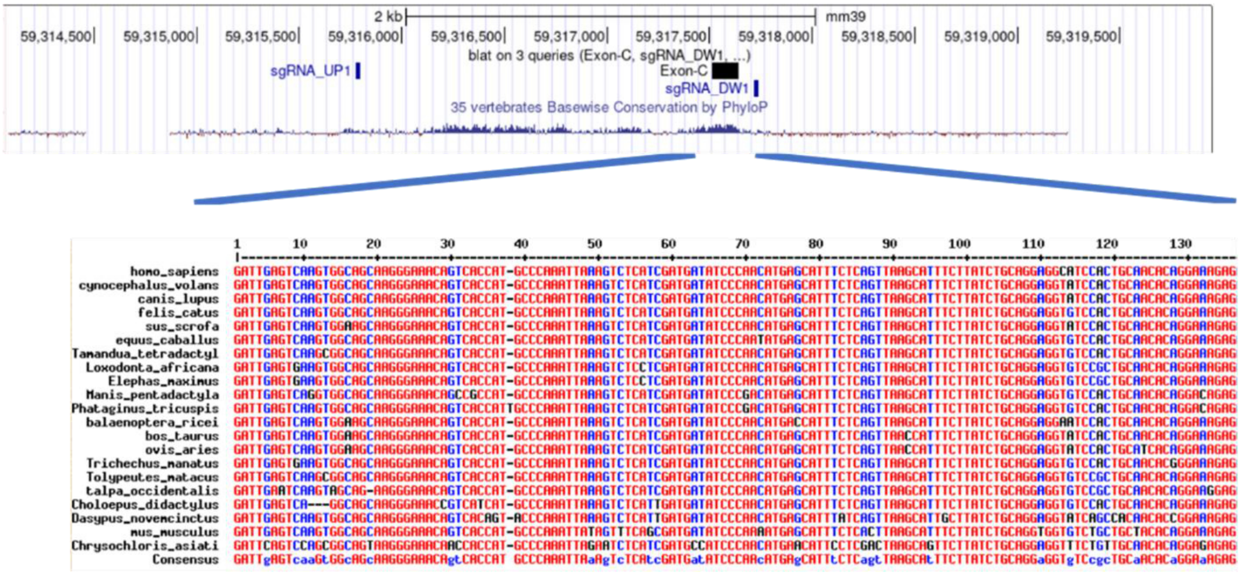


**B**

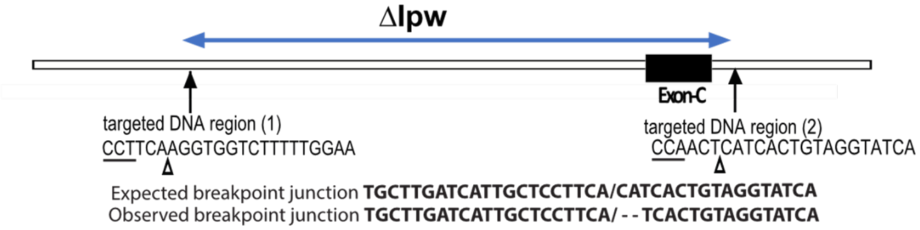


Supplementary Figure 7 - Marty *et al*

### Supplementary Figure 8

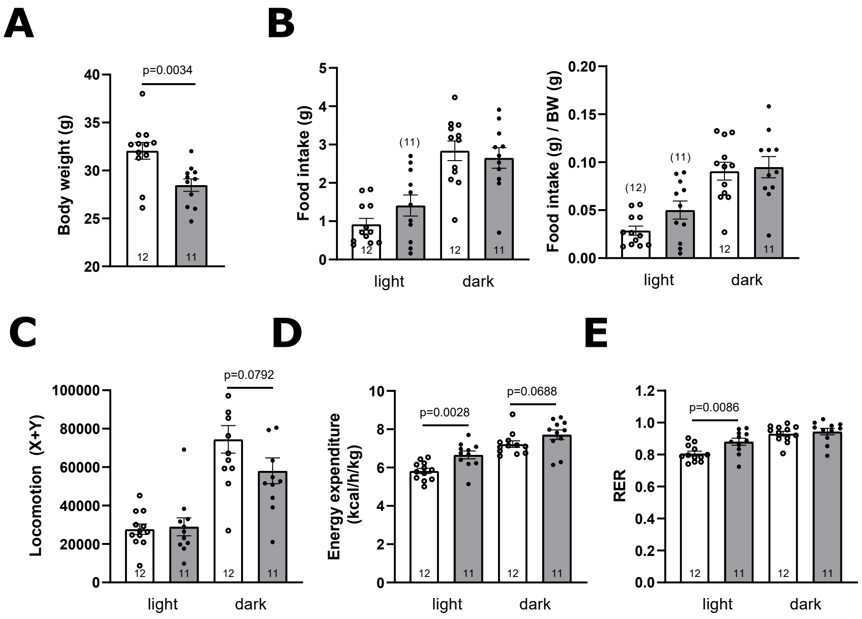


Supplementary Figure 8 - Marty *et al*

### Supplementary Figure 9

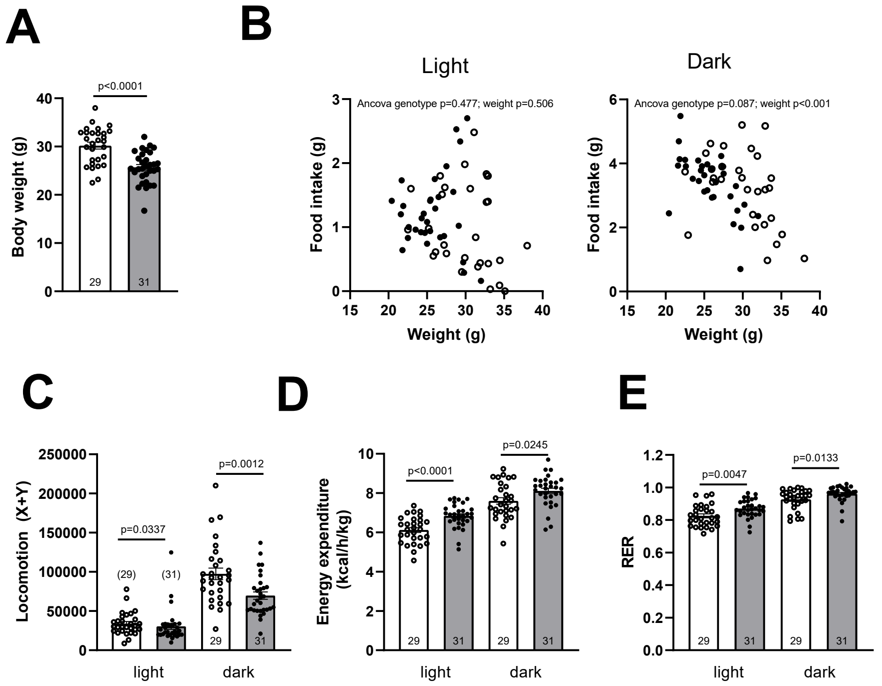


Supplementary Figure 9 - Marty *et al*

### Supplementary Figure 10

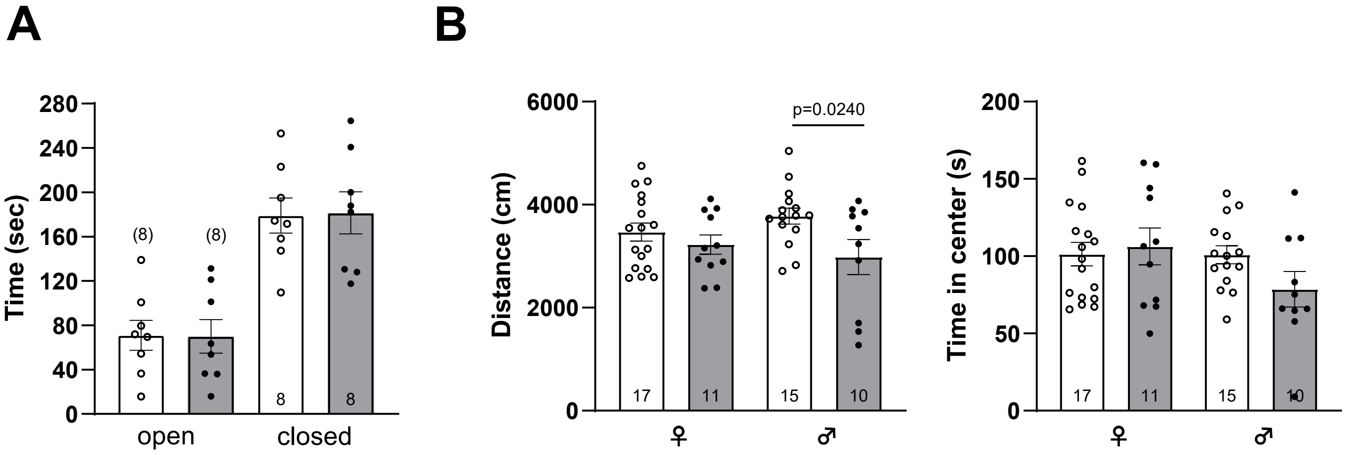


Supplementary Figure 10 - Marty *et al*

### Supplementary Figure 11

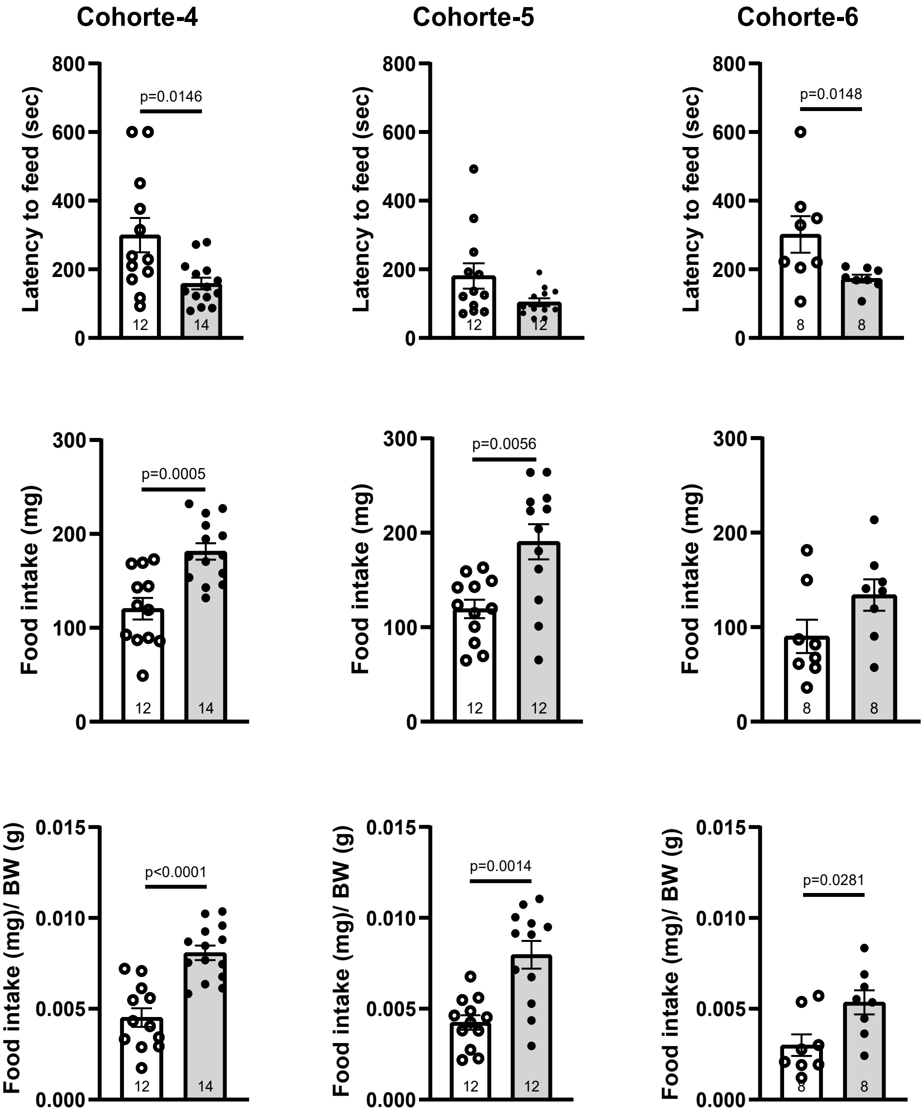


Supplementary Figure 11 - Marty *et al*

### Supplementary Figure 12

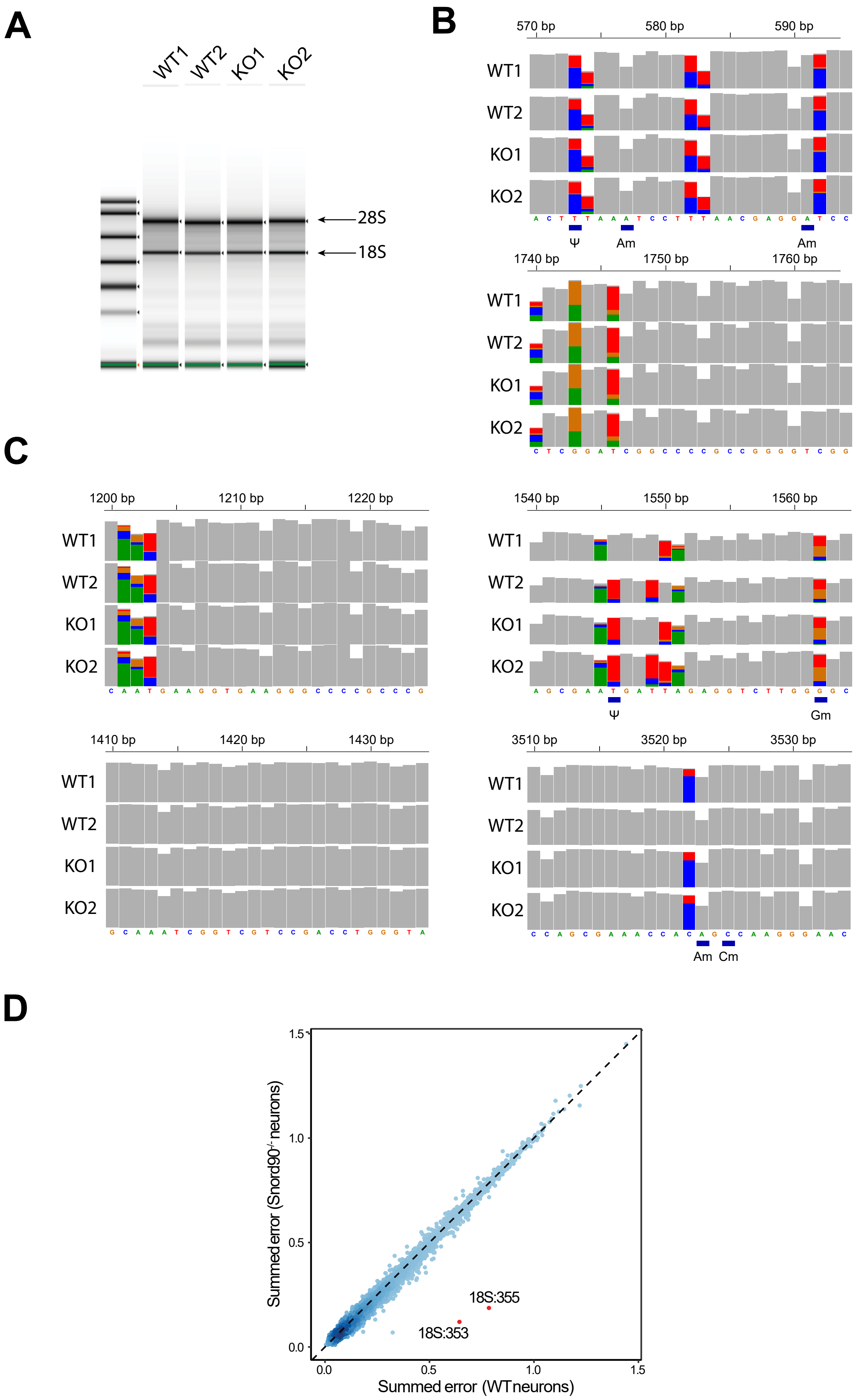


Supplementary Figure 12 - Marty *et al*
