## Supplementary Figure 3 for "Evolutionary and biological insights into the paternally expressed Snord116-Ipw-Snord115 gene array at the imprinted Prader-Willi syndrome domain"

**Cluster of SNORD115 in *L. tardigradus* (slender loris)**

>SNORD115-1
GGGTTGATGATGAGGACCTAAATTGTCTGAAGAGAGGTGATAAATAAAAAAATCATACACAAGAGGATTGAGCTG
AGGCCC
>SNORD115-2
GGGTGGATGATGATGACCTAAATTGTCTGAAGAGAGGTGATGAATAAAAAAATCATACACCAGAGGATTGAGCTG
AGGCCC
>SNORD115-3
GGGTTGATGATGACAACCTAAATTGTCTGAAGAGAAGTGATGAATAAAAAAATCATACACCAGAGGATTGAGCTG
AGGCCC
>SNORD115-4
GGGTCAATGATGACGACCTAAATTGTCTGAAGAGAGGTGATGAATAAAAAAATCATACACCAGAGGATTGAGCTG
AGGCCA
>SNORD115-5
GGGTTGATGATGACGACCTAAATTGCCTGAAGAGAGGTGATGAATACAAAAGTCATACACCAGAGGATTGAGCTG
AGGCCA
>SNORD115-6
GGGTAGATGATGATGACCTAAATTGCCTGAAGAGAGGTGATAAATAAAAAAAATCATACACCAGAGGATTGAACT
GAGGCCC
>SNORD115-7
GGGTTGATGATGACAACCTAATGTCTGAAGAGAGGTGATGAATAAAAAAATCATACACCAGAGGATTGAGCTGAG
GCCC
>SNORD115-8
GGGTCGATGATGACAACCTAAATTGTCTGAATAGAGGTGATAAATAAAAAAATCATAACCAGAGGATTGAGCTGA
GGCCC
>SNORD115-9
GGGTTGATGATGACAACCTAAATTGTCTGAAGAGAGGTGATGAATAAAAAAATCATAACCACAGGATTGAGCTGA
GGCCC
>SNORD115-10
GGGTTGATGATGATGACCTAAATTGTGTGAAGAGAAGTGATGAATAAAAAAATCATAAACCAGAGGATTGAGCTG
ACGCCC
>SNORD115-11
GGGTTGATGATGATAACCTAAATTGCCTGAAAAGAGGTGATGAATAAAGAAATCATACACCAGAGGATTGAGCTG
AGGCCC
>SNORD115-12
ggataGATGATGATGACCTAAATTGCCTGAAGAGAGGTGATGAATAAAAATATCATACACCAGAGGATTGAGCTG
AGGCCC
>SNORD115-13
GGGTCGATGATGACAACCTAAATTGTCTGAAGAGAGGTGATGAATAAAAAAATCATAACCACAGGATTGAGCTGA
GGCCC
>SNORD115-14
GGGTAGATGATGACGACCTAATTTGCCTGAAGAGAGGTGATGAATTAAAAAAATCATACACCAGAGGATTGAGCT
GAGGCCA
>SNORD115-15
GGGTTGATGATGACAACCTAATGTCTGAAGAGAGGTGATGAATAAAAAAAATCATACACCAGAGGATTGAGCTGA
GGCCC
>SNORD115-16
GGGTTGATGATGATGACCTAAATTGTGTGAAGAGAAGTGAAGAATAATAAAATCATACTCCAGAGGATTGAGCTG
AGGCCC
>SNORD115-17

GGGCTGATGATGACGACCTAAATTGTGTGAAGAGAAGTGATGAATAAAGAAATCATACACCAGAGGATTTAGCTG
AGGCCC
>SNORD115-18
GGGTTGATGATGATGACCTAAATTTTGTGAAGAGAAGTGATGACTAAAAAAAATCATACACCAGGGGATTGAGCTG
AGGCCC
>SNORD115-19
GGGTTGATGATGACAACCTAATGTCTGAAGAGAGGTGATGAATAAAAAAAAATCATACACCAGAGGATTGAGCTGA
GGCCC
>SNORD115-20
GGGTCGATGATGACAATCTAATGTCTGAAGAGAGGTGATGAATAAAAAAAATCATACACCAGAGGATTGAGCTGAG
GCCC
>SNORD115-21
GGGTCGATGATGACAATCTAATGTCTGAAGAGAGGTGATGAATAAAAAAAATCATACACCAGAGGATTGAGCTGAG
GCCC
>SNORD115-22
GGGTCAATGATGACAACCTAATGTCTGAAGAGAGGTGATGAATAAAAAAAATCATACACCAGAGGATTGAGCTGAG
GCCC
>SNORD115-23
GGGTCGATGATGACAACCTAATGTCTGAAGAGAGGTGATGAATAAAAAAAATCATACGCCAGAGGATTGAGCTGAG
GCCC
>SNORD115-24
GGATTGATGATGACCACCTAAATTGTCTGAAGAGAAGTGATAAATAAAATAATCATACACCAGAATTGAGCTGAG
GCCC
>SNORD115-25
GGGTCAATGATGACAACCTAATGTCTGAAGAGAGGTGATGAATAAAAAAAATCATACACCAGAGGATTGAGCTGAG
GCCC
>SNORD115-26
GGGTCGATGATGACAACCTAATGTCTGAAGAGAGGTGATGAATAAAAAAAAATCATACACCAGAGGATTGAGCTGA
GGCCC
>SNORD115-27
GGGTCGATGATGACAACCTAATGTCTGAAGAGAGGTGATGAATAAAAAAAAATCATACACCAGAGGATTGAGCTGA
GGCCC
>SNORD115-28
GGGTCGATGATGACAACCTAATGTCTGAAGAGAGGTGATGAATAAAAAAAAATCATACACCAGAGGATTGAGCTGA
GGCCC
>SNORD115-29
GGGTCGATGATGACAACCTAATGTCTGAAGAGAGGTGATGAATAAAAAAAAATCATACACCAGAGGATTGAGCTGA
GGCCC
>SNORD115-30
GGGTCGATGATGACAACCTAATGTCTGAAGAGAGGTGATGAATAAAAAAAAATCATACACCAGAGGATTGAGCTGA
GGCCC
>SNORD115-31
GGGTCGATGATGACAACCTAATGTCTGAAGAGAGGTGATGAATAAAAAAAAATCATACACCAGAGGATTGAGCTGA
GGCCC
>SNORD115-32
GGGTCGATGATGACAACCTAATGTCTGAAGAGAGGTGATGAATAAAAAAAAATCATACACCAGAGGATTGAGCTGA
GGCCC
>SNORD115-33
GGGTCGATGATGACAACCTAATGTCTGAAGAGAGGTGATGAATAAAAAAAAATCATACACCAGAGGATTGAGCTGA
GGCCC
>SNORD115-34
GGGTCGATGATGACAACCTAATGTCTGAAGAGAGGTGATGAATAAAAAAAAGTCATACACCAGAGGATTGAGCTGA
GGCCC

>SNORD115-35
GGGTCGATGATGACAACCTAATGTCTGAAGAGAGGTGATGAATAAAAAAAAGTCATACACCAGAGGATTGAGCTGA
GGCCC
>SNORD115-36
GGGTCGATGATGACAACCTAATGTCTGAAGAGAGGTGATGAATAAAAAAAAATCATACACCAGAGGATTGAGCTGA
GGCCC
>SNORD115-37
GGGTCGATGATGACAACCTAATGTCTGAAGAGAGGTGATGAATACAAAAAAATCATACACCAGAGGATTGAGCTGA
GGCCC
>SNORD115-38
GGGTCGATGATGACAACCTAATGTCTGAAGAGAGGTGATGAATACAAAAAAATCATACACCAGAGGATTGAGCTGA
GGCCC
>SNORD115-39
GGGTCGATGATGACTACCTAATGTCTGAAGAGAGGTGATGAATAAAAAAAAAATCATACACCAGAGGATTGAGCTG
AGGCCC
>SNORD115-40
GGGTCGATGATGACAACCTAATGTCTGAAGAGAGGTGATGAATAAAAAAAAAATCATACACCAGAGGATTGAGCTG
AGGCCC
>SNORD115-41
GGGTCAATGATGACAACCTAATGTCTGAAAAGAGGTGATGAATAAAAAAAATCATACACCAGAGGATTGAGCTGAG
GCCC
>SNORD115-42
GGGTCGATGATGACAAGCTAATGTCTGAAAAGAGGTGATGAATAAAAAAAATCATACACCAGAGGATTGAGCTGAG
GCCC
>SNORD115-43
GGGTCGATGATGACAACCTAATAGCTGAAGAGAGGTGATGAATAAAAAAAATCATACACCAGAGGATTGAGCTGAG
GCCC
>SNORD115-44
GGGTCGATGATGACAACCTAATGTCTGAAGAGAGGTGATGAATAAAAAAAATCATGCACCAGAGGATTGAGCTGAA
GCCC
>SNORD115-45
ggatcGATGATGACAACCTAATGTCTGAAGAGAGGTGATGAATAAAAAAAATCATACACCAGAGGATTGAGCTGAG
GGCC
>SNORD115-46
GGGTCGATGATGACAACCTAATGACTGAAGAGAGGTGATGAATAAAAAAAATCATGCACCAGAGGATTGAGCTGAG
GCCC
>SNORD115-47
GGGTCGATGATGACAACCTAATGTCTGAAGAGAGGTGATGAATAAAAAAAAACATGCACCAGAGGATTGAGCTGAG
GCCC
>SNORD115-48
GGGTCAATGATGACAACCTAATGTCTGAAGAGAGGTGATGAATAAAAAAAATCATAAACCAGAGGATTGAGCTGAG
GCCC
>SNORD115-49
GGGTCGATGATGACAACCTAATGTCTGAAGAGAGGTGATGAATAAAAAAAAAAATCATACACCAGAGGATTGAGCT
GAGGCCC
>SNORD115-50
gggtcagTGATGACAACCTAATGTCTGAAGAGAGGTGATAAATAAAAAAAAATCATACACTAGAGGATTGAGCTGA
GGCCC
>SNORD115-51
AGGTTTATGATGATGACCTAAATTGTCTGAAGAGAAGTGATGAGTATAAAAATCATACACCAGAGGACTGAGCTG
AAGCCG
>SNORD115-52

GGGTCGATGATGACAACCTAATGTCTGAAGAGACGTGATGAATAAAAAAAATCATACACCAGAGGATTGAGCTGA
GGCCC
>SNORD115-53
GGGTCGATGATGACAACCAAATGTCTGAAGAGAGGTGATGAATAAAAAAAATCATACACCAGAGGATTGAGCTGA
GGCCC
>SNORD115-54
GGGTCGATGATGACAACCTAATATCTGAAGAGAGGTGATGAATAAAAAAAATCATACACCAGAGGATTGAGCTGA
GGCCC
>SNORD115-55
GGGTCGATGATGACAATCTAATGTCTGAAGAGAGGTGATGAATAAAAAAAATCATACACCAGAGGATTGAGCTGA
GGCCC
>SNORD115-56
GGGTCAATGATGACAACCTAATGTCTGAAGAGAGGTGATGAATAATAAAAATCATACACCAGAGGATTGAGCTGA
GGCCC
>SNORD115-57
GGGTCAATGATGACAACCTAATGTCTGAAGAGAGGTGATGAATAATAAAAATCATACACCAGAGGATTGAGCTGA
GGCCC
>SNORD115-58
GGGTCGATGATGACAACCTAATGCCTGAAGAGAGGTGATGAATAAAAAAAATCATACACCAGAGGATTGAGCTGA
GGCCC
>SNORD115-59
GGGTCGATGATGACAACCTAATGTCTGAAGAGAGGTGATGAATAAAAAAAATCATGCACCAGAGGATTGAGCTGA
GGCCC
>SNORD115-60
GGGTCGATGATGACAACCTAATGTCTGAAGAGAGGTGATGAATAAGAAAAATCATACACCAGAGGATTGATCTGA
GGCCC
>SNORD115-61
GGGTCGATGATGACAACCTAATGTCTGAAGAGAGGTGATGAATAATAAAAATCATACACCAGAGGATTCAGCTGA
GGCCT
>SNORD115-62
GGGTCGATGATGACAACCTAATATCTGAAGAGAGGTGATGAATAAAAAAAAATCATACACCAGAGGATTGAGCTG
AGGCCC
>SNORD115-63
GGGTCGATGATGACAACCTAATGTCTGAAGAGAGGTGATGAATAAAAAAAAATCATACACCAGAGGATTGAGCTG
AAGCCC
>SNORD115-64
CGGTTGGTGATGACGACCTAAATTTTGTGAAGAGAAGTGATGAATAAAGTAATCATACACCATAGGATTGAGCTG
AGGCCC
>SNORD115-65
GGTTCGATGATGACAATCTAATGTCTGAAGAGAGGTGATGAATAAAAAAAATCATACACCAGAGGATTGAGCTGA
GGCCC
>SNORD115-66
GGGTCGATGATGACAACCTAATGTCTGAGAGAGGTGATGAATAAAAAAAATCATACACCAGAGGATTGAGCTGAG
GCCC
>SNORD115-67
GGGTCGATGATGATGACCTAAATTGTCTGAAGACTGGTGATGAATAAAAAAATCATAACCACAAGATTGAGCTGA
GACCC
>SNORD115-68
GGGTCGATGATGACAACCTAATAGCTGAAGAGAGGTGATGAATAAAAAAAATCATACACCAGAGGATTGAGCTGA
GGCCC
>SNORD115-69
GGGTCGATGATGACAACCTAATAGCTGAAGAGAGGTGATGAATAAAAAAAATCATACACCAGAGGATTGAGCTGA
GGCCC

>SNORD115-70
GGGTCGATGATGACAACCTAATAGCTGAAGAGAGGTGATGAATAAAAAAAATCATACACCAGAGGATTGAGCTGA
GGCCC
>SNORD115-71
GGGTCGATGATGACAAGCTAATGTCTGAAAAGAGGTGATGAATAAAAAAAATCATACACCAGAGGATTGAGCTGA
GGCCC
>SNORD115-72
GGGTCGATGATGACAACCTAATACCTGAAGAGAGGTGATGAATAAAAAAAATCATACACCAGAGGATTGAGCTGA
GGCCC
>SNORD115-73
GGGTCGATGATGACAACCTAATGTTTGAAGAGAGGTGATGAATAAGAAATATCATACACCAGAGGATTGAGCTGA
GGCCC
>SNORD115-74
GGGTCGATGATGACAACTTATTGTCTGAAGAGAGGTGATGAATAAAAAAAATTATACACCAGAGGATTGAGCTGA
GGCCC
>SNORD115-75
GGGTCAATGATGACAACCTAATGACTGAAGAGAGGTGATGAATAAAAAAATCATACACCAGAGGATTGAGCTGAG
ccta
>SNORD115-76
GGGTCGATGATGACAAGCTAATGTCTGAAAAGAGGTGATGAATAAAAAAAAATCATACACCAGAGGATTGAGCTG
AGGCCC
>SNORD115-77
GGGTCGATGATGACAGCCTAATGTCTGAAGAGAGGTGATGAATAAAAAAAAATCACACACCAGAGGATTGAGCTG
AGGCCC
>SNORD115-78
GGGTCGATGATGACAACCTAATGTCGGAAAAGAGGTGATGAATAAAAAAAACATACACCAGAGGATTGATCTGAG
GCCC
>SNORD115-79
GGGTCGATGATGACAACCTAATGTCTGAAGAGGTGATGAATAAAAAAAATCATACACCAGAGGATTGAGCTGAGG
CCC
>SNORD115-80
GGGTCGATGATGACAACCTAATGTCTGAGAGAGGTGATGAATAAAAAAAATCATACACCAGAGGATTGAGCTGAG
CCCC
>SNORD115-81
GGGTCGATGATGACAACCTACTGTCTGAGAGAGGTGATGAATAAAAAAAATCATACACCAGAGGATTGAGCTGAG
GCCC
>SNORD115-82
GGGTCGATGATGACAACCTAATGTCTGAGGGAGGTGATGAATAAAAAAAATCATACACCAGAGGATTGAGCTGAG
GCCC
>SNORD115-83
GGGTCGATGATGACAACCTGATGTCTGAAGAAAGGTAATGAATAAAAAAAATCATACACCAGAGGATTGAGCTGA
GGCCC
>SNORD115-84
GGGTCAATGATGACAACCTAATGTCTGAAGAGAGGTGATGAATAAAAAAAAAAAAATCATACACCAGAGGATTGA
GCTGAGGCCC
>SNORD115-85
GGGTCGATGATGACAACCTAATGTCTGAGAGAGGTGATGAATAAAAAAAATCATATACCAGAGGATTGAGCTGAG
GCCA
>SNORD115-86
GGGTCGATGATGACAACCTAAATTGAAGAGAAGTGATGAATAAAAAAATCATAACCACAGGATTGAGCTGAGGCC
C
>SNORD115-87

GGGTCAATGATGACAACCTAATGTCTGACGAGTGGTGATGAATAAAAAAAATCATACACCAGAGGATTGAGCTGA
Gtcac
>SNORD115-88
GGGTCAATGATGACAACCTAATGTCTGAAGAAGGTGATGAACAAAAAAAATCATAAACCAGAGGATTGAGCTGAG
GCCC
>SNORD115-89
tgataGATGATGACCACCTAATTTGTCTAAAGAAAAAGTGATGAATAAAAAATTCATACATCAGATGATTGAGCT
GAGGCTC

>SNORD115-90
GGGTCGATGATGACAACCTAATGTCTGAAGAGAGGTGATGAATAAAAAAAATCATACACCAGTGGTTGAGCTCAG
GCCC
>SNORD115-91
GGGTCGATGATGATGACCTAAATTGTCTGAAGAGAGGTGATGAATAGAATAATCATACACCAGAGGCTTGAGCTA
AGGCCC
>SNORD115-92
GGGTAGATGATGACGACCTAAATTGCCTGAAGAGTGGTGATGAATAAAAAATCATACACCAGAGGATTGAGCTCA
GGCCC
>SNORD115-93
tgggcGATGATGACGACCTAAATTGTCTAAAGAGAGGTGATGAATAGAATAATCATACACCAGAGGCTTGAGCTA
AGGCCC
>SNORD115-94
GGATTGATGATGACAACCTAAATTGTCTGAAGAGAAGTGATGAATAAAATAATCATACACCAGAATTGAGCTAAG
GCCC
>SNORD115-95
GGGTCGATGATGACAACCTAATGTCATGAAGAGAGGTGATGAATAAAAAAAACATACACCAGAGGATTGAGCAGA
GGCCC
>SNORD115-96
GGGTCGATGATGACAACCTAATAGCTGAAGAGAGGTGATGAATAAAAAAAATGATACACAAGAGGATTGAGCAGA
ttccc
>SNORD115-97
gagtcaattttgaccacctaacGTCTGAACGGAGGTGATGAATAAAAAAAATCTACACCAGAGGATTGAGCTGAG
GCCC
>SNORD115-98
GGGTCTAGGATGACAACCTAATGTCTGAAGAGAGGTGATGAATAAAAAAATCATACACCAGAGGATTGAGCTGAG
GCCC
>SNORD115-99
GGGTCAATAATGACAACCTAATGTCTGAAAAGAGGTGATGAATTAAAAAAATCATACACTAGACGATTGAGCTGA
GCCCA
>SNORD115-100
TGGTCGATAATGACCACCTAAATTGTCTGAAGAAGTGATCAATAAAAAATTCGTACATCAGAGGATTGAGCTGAG
GCCC
>SNORD115-101
GGGTCAATGATAACCACCAAAATTGTCTGAAGAAAAGTGATGAATAAAAAATTCATACATCAGAGGATTGAGCTG
AGGCCC
>SNORD115-102
TGGTCGATAATGACCACCTAAATTGTCTGAAGGAGTGATCAATAAAAAATTCATACATCAGAGGATTGAGCTGAG
GCCC
>SNORD115-103
GGGTCGATGATGGCAAACTAATGTCTGAAGAGAGGTGATGAATAAAAAAAATCATACACCAGAGGATTGAGCTGA
GGTCC
>SNORD115-104

GGGTCAATGATAACCACCTAAATTGTCTGAAGAAAAGTGATAAATAAAAAGCTCATACATCAGAGGATTGAGCAG
AGGCCC
>SNORD115-105
GGGTCAAGGATGACAACCTAATGTCTGAAGAGAGGTGATGAATAATAAAAATCATACACCAGAGGATTGAGCTGA
GGCCC
>SNORD115-106
GGGTCGATGACGACAACCTAATGTCTGAAGAGAGGTGATGAATAAAAAAAATCATACACCAGAGGATTGAGCTGA
GGCCC
>SNORD115-107
cctgcgtctATGACGACCTAAATTGCTTGAAGAGACGTGATGAATAAAAAAATCATACACCAGAGGATTGAGCTG
AGGCCT
>SNORD115-108
GGGTCGATGATGGCAACCTAATGTCTGAAGAGAGGTGATGAATAAAAAAAATCATACACCAGAGGATTGAGCTGA
GGCCC
>SNORD115-109
GGGTTGATTATGACAACCTAATGTCTGAAGAGAGGTGATGAATAAAAAAAATCATACACCAGAGGATTGAGCTGA
GGTCC

### Cluster of SNORD115 in *N. coucang* (slow loris)

>SNORD115-1
GGGTCAATGATGACAACCTAATATCTGAAGAGAGGTGATGAATAAAAAAATCATACACCA
GAGGATTGAGCTGAGGCCC
>SNORD115-2
GGGTCGATGATGACAACCTAATGTCTGAAGAGAGGTGATGAATAAAAAAAATCATACACC
AGAGGATTGAGCTGAGGCCC
>SNORD115-3
GGGTCGATGATGACAACCTAATGTCTGAAGAGAGGTGATGAATAAAAAAAATCATACACC
AGAGGATTGAGCTGAGGCCC
>SNORD115-4
GGGTCGATGATGACAACCTAATGTCTGAAGAGAGGTGATGAATAAAAAAAATCATACACC
AGAGGATTGAGCTGAGGCCC
>SNORD115-5
GGGTCGATGATGACAACCTAATGTCTGAAGAGAGGTGATGAATAAAAAAAATCATACACC
AGAGGATTGAGCTGAGGCCC
>SNORD115-6
GGGTCGATGATGACAACCTAATGTCTGAAGAGAGGTGATGAATAAAAAAAATCATACACC
AGAGGATTGAGCTGAGGCCC
>SNORD115-7
GGGTCGATGATGACAACCTAATGTCTGAAGAGAGGTGATGAATAAAAAAAATCATACACC
AGAGGATTGAGCTGAGGCCC
>SNORD115-8
GGGTCGATGATGACAACCTAATGTCTGAAGAGAGGTGATGAATAAAAAAAATCATACACC
AGAGGATTGAGCTGAGGCCC
>SNORD115-9
GGGTCGATGATGACAACCTAATGTCTGAAGAGAGGTGATGAATAAAAAAAATCATACACC
AGAGGATTGAGCTGAGGCCC
>SNORD115-10
GGGTCGATGATGACAACCTAATGTCTGAAGAGAGGTGATGAATAAAAAAAATCATACACC
AGAGGATTGAGCTGAGGCCC
>SNORD115-11
GGGTCGATGATGACAACCTAATGTCTGAAGAGAGGTGATGAATAAAAAAAATCATACACC
AGAGGATTGAGCTGAGGCCC
>SNORD115-12
GGGTCGATGATGACAACCTAATGTCTGAAGAGAGGTGATGAATAAAAAAAATCATACACC
AGAGGATTGAGCTGAGGCCC
>SNORD115-13
GGGTCGATGATGACAACCTAATGTCTGAAGAGAGGTGATGAATAAAAAAAATCATACACC

```
AGAGGATTGAGCTGAGGCCC
>SNORD115-14
GGGTCGATGATGACAACCTAATGTCTGAAGAGAGGTGATGAATAAAAAAAATCATACACC
AGAGGATTGAGCTGAGGCCC
>SNORD115-15
GGGTCGATGATGACAACCTAATGTCTGAAGAGAGGTGATGAATAAAAAAAATCATACACC
AGAGGATTGAGCTGAGGCCC
>SNORD115-16
GGGTCGATGATGACAACCTAATGTCTGAAGAGAGGTGATGAATAAAAAAAATCATACACC
AGAGGATTGAGCTGAGGCCC
>SNORD115-17
GGGTCGATGATGACAACCTAATGTCTGAAGAGAGGTGATGAATAAAAAAAATCATACACC
AGAGGATTGAGCTGAGGCCC
>SNORD115-18
GGGTCGATGATGACAACCTAATGTCTGAAGAGAGGTGATGAATAAAAAAAATCATACACC
AGAGGATTGAGCTGAGGCCC
>SNORD115-19
GGGTCGATGATGACAACCTAATGTCTGAAGAGAGGTGATGAATAAAAAAAATCATACACC
AGAGGATTGAGCTGAGGCCC
>SNORD115-20
GGGTCGATGATGACAACCTAATGTCTGAAGAGAGGTGATGAATAAAAAAAATCATACACC
AGAGGATTGAGCTGAGGCCC
>SNORD115-21
GGGTCGATGATGACAACCTAATGTCTGAAGAGAGGTGATGAATAAAAAAAAATCATACAC
CAGAGGATTGAGCTGAGGCCC
>SNORD115-22
GGGTCGATGATGACAACCTAATGTCTGAAGAGAGGTGATGAATTAAAAAAATCATACACC
AGAGGATTGAGCTGAGGCCT
>SNORD115-23
GGGTCGATGATAACAACCTAATGTCTGAAGAGAGGTGATGAATAAAAAAAATCATACACC
AGAGGATTGAGCTGAGGCCC
>SNORD115-24
GGGTCTATGATGACGACCTAAATTGTCTGAAGAGAGGTGATGAATAAAAAAATCATACAC
CAGAGGATTGAGCTGAGGCCC
>SNORD115-25
GGGTCGATGATAACAACCTAATGTCTGAAGAGAGGTGATGAATAAAAAAAATCATACACC
AGAGGATTGAGCTGAGGACC
>SNORD115-26
GGGTCAATGATGACAACCTAATGTCTGAAGAGAGGTGATGAATAAAAAAAATCATACACA
AGGGGATTGAGCTTAGGCCC
>SNORD115-27
GGGTCTATGATGACGACCTAAATTGTCTGAAGAGAGGTGATAAATAAAAAAATCATACAC
CAGAGGATTGAGCTGAGGCCC
>SNORD115-28
GGCTCGATGATGACAACCGAATGTCTGAACAGAGGTGATGAATAAGAAAAATCATACACC
AGAGGATTGAGCTGAGGCCC
>SNORD115-29
GGGTTGATGATGACAACCTAATCTCTGAAGAGACGTGATGAATAAAAAAAAATCATACAC
AAGAGGATTGAGCTGAGGCCC
>SNORD115-30
TGGTCGATGATGACAACTTAATGTCTGAGGAGAGATGATGAATAAAAAAAATCATACACC
AGAGGATTGAGCTGAGGCCC
>SNORD115-31
GGGTCGATGATGACAACCTAATGTCTAAAGAGAGGTGATGAATAAAAAAAAAAAATCATAC
ACCAGAGGATTGAGCTAAGGTCC
>SNORD115-32
GGGTCTATGATGACGACCTAAATTGTCTGAAGAGAGGTGATGAGTAAAAAATCATACACC
AGAGGATTGAGCTGAGGCCC
>SNORD115-33
GGGTCGATGATGACGACAACCTGATGTCTGAAGAGAGGTGATGAATAAAAAAAAAAACATA
CACCAGAGGATTGAGCTGAGGCCC
```

>SNORD115-34
GGGTCTATGATGACGACCTAAATTGTCTGAAGAGAGGTTATGAATAAAAAAAAATCAAAC
ACCAGAGGATTGAGCTCAGGCCC
>SNORD115-35
GGGTTGATGATGATGACCTAAATTGTCTGAAGAGAGGTGATGAATAGAATAATCCTACAC
CAGAGGATTGAGCTGAGGCCC
>SNORD115-36
GGGTTGATGATGACAACCTAATGTCTGAAAAGTAGTGATGAATAAAAAAAAAATAATACA
CCAGGGGATTGAGCTGAGGCCC
>SNORD115-37
GGGTTGATGATGACCACCTAAATTGTCTGAAGAAAAGTGATGAATAAAAAATTCATACAT
CAGAGGATTGAGCTGAGACCC
>SNORD115-38
GGGTTGATGATGATGACCTAAATTGTCTGAAGACAAATGATGAATAAAAAAATCATACAC
CAGAGGATTGAGCTGAGGTCC
>SNORD115-39
TGGTTGATGATGATGACCTAAATTGTCTGAAGAGAGGTGATGAATAAAAAGTCATACACC
AGAGGATTGAGCTGAGGTCC
>SNORD115-40
GGGTCAATGATGACGATCTAAATTGTCTAAAGGAAGGTGATGAATAAAAAAATCATAAAC
AAGAGGACTGAGCTGAGGTCA
>SNORD115-41
GAGGCCATGATGACGACCTAAATTGTCTGAAGAAAGGTGACGAATAAAAAAATCATGAAC
AGAGGATTGAGCTGAGGCCC
>SNORD115-42
GAGGCCATGATGACGACCTAAATTGTCTGAAGAAAGGTGACGAATAAAAAAATCATGAAC
AGAGGATTGAGCTGAGGCCC
>SNORD115-43
GAGGCCATGATGACGACCTAAATTGTCTGAAGAAAGGTGACGAATAAAAAAATCATGAAC
AGAGGATTGAGCTGAGGCCC
>SNORD115-44
GAGGCCATGATGACGACCTAAATTGTCTGAAGAAAGGTGACGAATAAAAAAATCATGAAC
AGAGGATTGAGCTGAGGCCC
>SNORD115-45
GAGGCCATGATGACGACCTAAATTGTCTGAAGAAAGGTGACGAATAAAAAAATCATGAAC
AGAGGATTGAGCTGAGGCCC
>SNORD115-46
GAGGCCATGATGACGACCTAAATTGTCTGAAGAAAGGTGACGAATAAAAAAATCATGAAC
AGAGGATTGAGCTGAGGCCC
>SNORD115-47
GGGTTGATGATGACTACCTCAATTGTGGAAGAGAAGTGATGAATAAAAAAATCATAACCT
GAGGATTGAGCTGAGGCCC
>SNORD115-48
GGGACGATGATGATGACCTAAATTGTCTGAAGAAAGGTGATGAATAAAAAATCTTACCCA
GAGGATTGAGCTGAGGCCC
>SNORD115-49
GAGGCCATGATGACGACCTAAATTGTCTGAAGAAAGGTGACGAATAAAAAAAATCATAAA
CAGAGGATTGAGCTGAGGCCC
>SNORD115-50
GAGGCCATGATGACGACCTAAATTGTCTGAAGAAAGGTGACGAATAAAAAAAATCATAAA
CAGAGGATTGAGCTGAGGCCC
>SNORD115-51
CGGTTGATGATGACGACCTCAATTGTGGAAGAGAAGTGATGAATAAAAAAATTATAACCT
GAGGATTGAGCTGAGGCCC
>SNORD115-52
CGGTTGATGATGACGACCTCAATTGTGGAAGAGAAGTGATGAATAAAAAAATTATAACCT
GAGGATTGAGCTGAGGCCC
>SNORD115-53
CGGTTGATGATGACGACCTCAATTGTGGAAGAGAAGTGATGAATAAAAAAATTATAACCT
GAGGATTGAGCTGAGGCCC
>SNORD115-54

CGGTTGATGATGACGACCTCAATTGTGGAAGAGAAGTGATGAATAAAAAAATTATAACCT
GAGGATTGAGCTGAGGCCC
>SNORD115-55
CGGTTGATGATGACGACCTCAATTGTGGAAGAGAAGTGATGAATAAAAAAATTATAACCT
GAGGATTGAGCTGAGGCCC
>SNORD115-56
CGGTTGATGATGACGACCTCAATTGTGGAAGAGAAGTGATGAATAAAAAAATTATAACCT
GAGGATTGAGCTGAGGCCC
>SNORD115-57
GAGGCCATGATGACGACCTAAATTGTCTGAAGAAAGTGACGAATAAAAAAAATCATAAAC
AGAGGATTGAGCTGAGGCCC
>SNORD115-58
CGGTTGATGATGACGACCTCAATTGTGGAAGAGAAGTGATGAATAAAAAAATTATAACCT
GAGGATTGAACTGAGGCCC
>SNORD115-59
GGGTTGATGTTGACGACCTCAATTGTGGAAGAGAAGTGATAAATAAAAAAATTATAACCT
GAGGATTGAGCTGAGGCCC
>SNORD115-60
GGGTTGATGGTGACGACCTCAATTGTGGAAGAGAAGTGATGAATAAAAAAATCATAACCT
GAGGACTGAGCTGAGGCCC
>SNORD115-61
GGGTTGATGTTGACGACCTCAATTGTGGAAGAGAAGTGATGAATAAAAAAATTATAACCT
GAGGATTGAGCTGAGGCCC
>SNORD115-62
GGGTCGATGGTGACGACCTAAATTGTCTGAAGAGAGGTGATGAATTAAAAAGTCATATAC
CAGGGCGGCGCCTGTGGCTC

### Cluster of SNORD115 in *C. bancanus* (Horsfield’s tarsier)

>SNORD115-1
GGTCAATGATGAGAACTTTATATTGTCTGAAGAGAGGTGATGACTTAAAAATCATGCTCAATAGGATTATGCTGA
GGCCC
>SNORD115-2
GGTCAATGATGAGAACTTTATATTGTCTGAAGAGTGATGATGACTTAAAAATCATGCTCAATAGGATTATGCTGA
GGCCA
>SNORD115-3
GGTAAATGATGAGAACTTTATATTGTCTGAAGAGAGGTGATGACGTAAAAATCATGCTCAATAGAATTATGCTGA
GGCCA
>SNORD115-4
GGTCAATGATGAGAACTTTATATTGCCTGAAGAGAGGTGAGGACTTAAAAATCATGCTCAATAGGATTACTCTGA
GGCCC
>SNORD115-5
AGTCAATGATGAGAACTTTATATTGTCTAAAGAGAAGTGATGGCTTAAAAATCATGCTCAATAGTATTATGCTGA
GGCCC
>SNORD115-6
AGTCTATGATGAGAACTTTATATTGTCTGAAGAGAGGTGATGGCTTAAAAACCATGCTCAATAGGATTATGCTGA
GATTC
>SNORD115-7
GGTCAATGATGAGAACTTTATATTGTCTTAAAGAGAGGTGATGACATAAAAATTATGCTCAATAGGATTATGCTG
AGCCCA
>SNORD115-8
GGTCAATGATGACATCTTTATATTGTCTGAAGTGAGGTGGTGACTTAAAAATCATGCTCAGTAGGATTATGCTGA
GGCTC
>SNORD115-9
GGTCAATGATGAGAACTTTATACTGTCTGAAGAGACGTGATGACTTAAAAATCATGCTTAATAAGATTACGCTGA
GGCAC
>SNORD115-10

GGTCAATGATGAGAACTTTATATTGACTGAAGAGAGATGATGACCTAAAAATCATGCTCAACAGAATTACGCTGA
GGCCC
>SNORD115-11
GGTCAATGATGAGAAATTTATATTGTCTGAAGAGAGGTGATGACCTAAAAATCATGCTCAATAAGATTACTCTGA
GACCC
>SNORD115-12
GCTCTATGATGAGAGCTGTATATTGTCTTGAAGAGAGGTGATGACTTAAAAATCATGCTCAATAGGATTACACTG
AGGCCC
>SNORD115-13
GGTCAATGATGAGCACGTTATATTGTCTTGAAGAGAGGTGATCACTTAAAAATGCTCAATAGGATTATGCTGAGG
CCC
>SNORD115-14
GGTCAATGATGAGACATTTACATTGTCTGTAGATAGATGATGACTTCAAAATCATGCTCAATAGGATTAGGCTGA
GGCCC
>SNORD115-15
GGTCAATGATGAGACCTTTATATTGTCTGAGAAGAGGCGATGACTTAAATATTATGCTCAACAGGATTACACTGA
GGCCC
>SNORD115-16
AGTCAATGATGAGAAGTTTATGTTATCTGAAGACAGGTGATGACTTAAAATCATGCTTGATAGGATTATGCTGAG
ACCC
>SNORD115-17
GGTCAATGATGAAGACTTTATATTGTCTTGAAGAGAGGTGATGACTTAAAATTCATGTACAATGGCATGATGCTG
AGGCCC
>SNORD115-18
GGTCAATGATGAGACCATTGTATTGTCTGTAGAGAGATGATGACTTAAAAGTCATGCCCAATAGGATTTCCCTGA
GTCCC
>SNORD115-19
TGTCAATGATGAGATCTTTTCACTGCTGTAGAGAGATGATAACTTAAAAATCATGCTCAATAGGATTACACTGAG
GCCC
>SNORD115-20
GGTCAATGATGAGGACTTTATATTGCCTTGAAGAGCGGTGATGACTTAAATATCATGTACAATGGCATGATGCTG
AGACCC
>SNORD115-21
GATCCCTGATGAGAACTTTATGTTGTCCTGAAGTAAACTGATGACTTAAAAATCATGCTCAATAGGATTACACTG
AGTCCC
>SNORD115-22
GGTCAATGATGAGTACTTTATATTGCCTTGAGGAGCGGTGATGACTTAAATATCAAGTACAATGGCATGATGCTG
AGGCCC
>SNORD115-23
GATCAATGATGAGGACTTTATATTGCCTTGAAGAGCAGTGATGACTTAAATATCATGCACAATGGCATGATGCTG
AGATGC
>SNORD115-24
GGTCAATGATGAGGACTTTATATTGCCTTGAGGAGCGGTGATGACTTAAATATCACGTACAATGGCATGATGCTG
AGGCCT
>SNORD115-25
GGTCGATGATGAAAACTTTGCCTTGTCTCTAGAGAGGTGTTGTCTTAAAAGTCATGCTCAATAGAGTTAGGCTGA
GGCCC
>SNORD115-26
GGTCAATGATGAGTACTTTATATTGCCTTGAGGAGCGGTGATGACTTAAATATCAAGTACAATGGCATGACGCTG
AGGCCC
>SNORD115-27
GGTCAATGATGAGTACTTCATATTGCCTTGAGGAGAGGTGATGACTTAAATATCAAGTACAATGGCATCACGCTG
AGGCCC
>SNORD115-28
GGTCAATGATGAGGACTTTATATTGCCTTGAGGAGTGGTGATGACTTAAATATCATGTACAATGGCATGACACTG
AGGCCC
>SNORD115-29

GGTCAATGATGAGGACTTTATATTGTCTTGAGAAGTGGTGATGACTTAAATATCGTGTACAATGGCATGACGCTG
AGGCCC
>SNORD115-30
GGTCAATGATGAGTACTTTATATTGTCTTGAGGAGTGGTGATGACTTAAATATCATGCACAATATGGAAATGTCA
CTGAGCCC
>SNORD115-31
GTTCAGTGATGAAACCTTTGTATTGTGTTTAGAGAGGTGATGACTTAAATGTCACTCTCAATAGAATTATGCTGA
GGTCC
>SNORD115-32
GGTCAATGATGAGTACTTTATATTGCCTTGAGGAGCGGTGATGACTTAAGTATCACGTACAATGGCATGATGCTG
AGGCCT
>SNORD115-33
GGTCAATGATGAGTACTTTATATTGCCTTGAGGAGCAGGTGATGATGTAAATATCAAGTTCAATGGCATGATGCT
GAGGCCT
>SNORD115-34
AGTCAATGATGAGAACTTTATATTGCCTTGAGGAAAGGTGATGACTTAAATATCATGTACAGTGGCATGACACTG
AGGCCA
>SNORD115-35
GGTCAATGATGAAACTTTTGTATGGTCTCTAGAGAGATGATGACTTAAATGTCATACTCAGTAGGATTACGCTGA
GACCA
>SNORD115-36
GGTAAATGATGAGTACTTTATATTGCCTGGAAGAGCGGTGATGACTTCAATATCACGTCCAATGGCATGAGGCTG
AGGCCC
>SNORD115-37
GGTCAATGATGAGTACTTTATATTGCCTTGAGGAGTGGTGATGACTCAAATATCAAGTACAATGGCATGACGCTG
AGGCCC
>SNORD115-38
GGTCAATGATGAGTACTTTATATTGCCTTGAGGAGCGGTGATGACTCAAATATCAAGTACAATGGCATGACGCTG
AGGCCC
>SNORD115-39
GGTCAATGATGAGTACTTTATATTGCCTTGAGGAGCGGTGATGACTCAAATATCAAGTACAATGGCATGACGCTG
AGGCCC
>SNORD115-40
GGTCAATGATGAGTACTTTATATTGCCTTGAGGAGCGGTGATGACTCAAATATCAAGTACAATGGCATGACGCTG
AGGCCC
>SNORD115-41
GGTCAATGATGAGTACTTTATATTGCCTTGAGGAGCGGTGATGACTCAAATATCAAGTACAATGGCATGACGCTG
AGGCCC
>SNORD115-42
GGTCAATGATGAGTACTTTATATTGCCTTGAGGAGCGGTGATGACTTAAATATCAAGTACAATGGCATGACACTG
AGGCCC
>SNORD115-43
GGTCACTGATGAAGACTTTATATTGCCTTGAGGAGTGGTGATGACTTAAAGATCATGTACAATGGCATGACGCTG
AGGCCC
>SNORD115-44
GGTCAATGATGAAACCTTTGCATTGTATATAGAGAGCTGATGAATTAAACGTCATGCTCAACAGAATTATGCTGA
GTTCC
>SNORD115-45
GGTCAATGATGAAATCTTAACCTTGTCTCTAGAGAGGTGTTGTCTTAAAATTCATGCACAATAGAGTTACGCTGA
GGACC
>SNORD115-46
GGTCAATGATGAGTACTTTATATTGCTTTGAGGAGCGGTAATGACTTAAATATCACACACAATGGATGACGCTGA
GGCAC
>SNORD115-47
GGTCAATGATGAGAACTTTATATTGCCTTGAGGAGTGGTGATGACATAATATCACATACAATGGCATGATGCTGA
GGCCC
>SNORD115-48

GGTCAATGATGAGTACTTTATATTGCCTAGAGGAGAGATGATGACTTAAACATCAAGTACAATGGTGTGACGCTG
AGGCTC
>SNORD115-49
GGTTGATGATGAGGACTTTATATTGCCTTGAGGAGCGGTGATGACTTAAATATCATGTACAATGGCATGACACTG
AGGCCC
>SNORD115-50
GGTCAATGATGAGTACTTTATGTTGCTTTGAGGAGCGGTGATGACTTAAATGTCATGTACAATGGCATGACACTG
AGGCCC
>SNORD115-51
GGTCAATGATGAGTACTTTATATTGCCTTGAGGAGTGGTGATGACTTAAATGTCACGTACAATGGCATGAAGCTG
AGTCCC
>SNORD115-52
GGCCAATGATGAGTGCTTTATATTGCCCTGAGGAGCGGTGATGACTTAAATATCAAGTACAATGGCATGACGCTG
AGGCCC
>SNORD115-53
GGCCAATGATGAGTGCTTTATATTGCCCTGAGGAGCGGTGATGACTTAAATATCAAGTACAATGGCATGACGCTG
AGGCCC
>SNORD115-54
GGTCAATGATGAGTACTTTATATTGCCTTGAGGAACGGTGATGACTTCAATATCAAGTACAATGGCATCACGCTG
AGGCCC
>SNORD115-55
GGTCAATGATGAGTACTTTATATTGCTTTGAGGAGCAGTGATGACTTAAATATCAGGTACAATGGCATGACGCTG
AGTCCC
>SNORD115-56
GGTCAATGATGAGGACTTTATATTGCCTTGAGGAGCGGTGATGACTTAAATATCACGTACAATGGCCTGACGCTG
AGGACC
>SNORD115-57
GGTCAATGATGAGTACTTTATATTGCCTTGACAGCGGTGATAACTTAAATATCACGTACAATGGCATGACGCTGA
GTCCC
>SNORD115-58
GGTCAATGATGAGTACTTTATATTGCCTGAGGAGCTGTGATGACTTAAATATCAAGTACAATGGCATGTCACTGA
AGCCC
>SNORD115-59
GGTCAATGATGAAGACTTTATATTGCTTTGAGGAGAAGTGATGATTTAAATACCAGCACAATGGCATGATGCTGA
GGTCC
>SNORD115-60
GATCAATGATGAGTACTTTATATTGCCTTGAGGAGCAGTGATGACTTAAATATCACATACAATGGCATGACGCTG
AGGCCC
>SNORD115-61
GGTCAATGATGAAATCTTTCCCTTGTCTCTAAAGAGGTGCTGTCTTAAAAGTCATGCTCAATAGAGTTTCGCTGA
GGATC
>SNORD115-62
GGTCAATGATGATGACTTTATATTGCCTTGAGACGAAGCGATGTCTTAAATACCATGCATAATTGAATGATGCTG
AGGCCC
>SNORD115-63
GGTCAGTGATGAGGACTTTGTATTTCCTTGAGGAGCAGTGGTGACTTAAATATCATGTACAATGGCATGATGCTG
AGGCCC
>SNORD115-64
GGTCAATGATGAGTACCTTATATTGCCTTGAGCAGCGGTGATGACTTAAATATCAAGTACAATGGCATGACTCTG
AGGCCC
>SNORD115-65
GGTTAATGATGAGTACTTTATATTGCCTTGAGGAGCGGTGATGATGTAAATATCAAGTACAATGGCATGACGCTG
AGGCCC
>SNORD115-66
GGTCAATGATGAGGACTTTATATTGCCTTGAGAAGCAGTGATGACTTAAATATCACATACAATGGCATGACGCTG
AGGCCC
>SNORD115-67

AGCCAATGATGAGTACTTTATATTGCCCTGAGGAACGGAGATGACTTAAATATCAAGTACAATGGCATGACGCTG
AGGCCC
>SNORD115-68
TGTCAATGATGAGTATTTTATATTGCCTTGAGGAGTGGTGATGATTTAAATATCAAGTACAATGGCATGACACTG
AGGCCC
>SNORD115-69
GGTCAATGATGAAAACTTTGCCTTGTCTCTAGAGAGGTGTGGTCTTAAAAGTCTTGTGCAATAGAGTTACGCTGA
GGGCC
>SNORD115-70
GGTCAATGATGAGTACCTTATATTGCTTTGAGGAGCATTGATGAATTCAATATCATGTACAATGGCATGACGCTG
AGGCCC
>SNORD115-71
GGTCAATGATGAAATGTTTGACTTGTCTCTAAAGAGGTGTTGTCTTAAAAGTCTTGCACAATAGAGTTATGCTGA
GGAAA
>SNORD115-72
AGTCAGTGATGAAAACTTTGTCCTGTCTCAAGAGAGGTGTTGTTTTAAAATCTTGTGCAATTGAGTTATGCTGAG
GCCC
>SNORD115-73
GGTCAATGATGAGTACTTTATATTGCCTTGAGGAACGGTAATGACATCAATATCAAGTACAATGGCATGAAGCTG
AGGCCC
>SNORD115-74
GGTCAATGATGAGTACTTAATATTGCTTTGAAGAGCAGTGATGACTTCAATATCACATACAATGGCATGACACTG
AGGCCC
>SNORD115-75
AGTCAATGATGAAATCTTTGCCTTGTCTCTAGAGAGGTGTTGTGTTAAAAGTCATGCACAATAGATTTACACTGA
GGACC
>SNORD115-76
GGTCAATGATGAAGACTTTATGTTGCCTTGAGGAGCAGTGAGGACTTAAATATCACGTACAATGGCATGACGCTG
AGGCCC
>SNORD115-77
GGTCAATGATGAATACTTTATATTGCCTTGAGGAGCGGTGATGACTTAAGTATCACATACAATGGCATGACGCTG
AGGTCG
>SNORD115-78
GGTCAATGATGAGTACTTTATATTACCTTGAAGAGGGTTGATGGCTTACATGTCACGTACAATGGCATGATGCTG
AGGAAC
>SNORD115-79
GGTCAATGATGAGTACTTTATATTGCCTTGAGGAGCAGTGATGACTTAAATATCACATACGATGGTAATACGCTG
AGTCTC
>SNORD115-80
GGTCAATGATGAATACTTTATAATACCTTGAGGAGCGGTGATGACTTAAATATCAAGTACAATGGCATGACACTG
AGGCCG
>SNORD115-81
GGTCAGTGATGAAATCTTTGCCTTGTCTCTAGAGAGGTGTTGTCTTATAAGTCTTGCTCAATAGAGCTACGCTGA
GGACC
>SNORD115-82
CATCAATGATGAAATCTATGCCTTGTCTATAGTGAGGTGTTGTCTTAAATATCGGGCTCAATAGAGTTATGCTGA
GGAAC
>SNORD115-83
GGTCAATGATGAAATCTTTGTCTTGACTCTAGTGAGGTGTTGTCTTAAAAGTCTTGTGCAAAAGAGTTATGCTGA
GGAAC
>SNORD115-84
GGTCAGTGATGAAATCTTTGTCTTGTCTCCAGAGAGGTGTTGTCTTAAAAGTTTTGCACAATATAGTTGTGCTGA
GGACA
>SNORD115-85
GGTCAATGATGAGTACTTTATATTGCCTTGAGTAACAGTGATGACTTCAATATCAAGTACAATGGCATGACACTG
AGGCCC
>SNORD115-86

GGTCAATGATGAATACTTTATACTGCCTTGAGAATCGGTGATGACTTGAATAACATGTACAATGGCATGACGCTG
AGGCCC
>SNORD115-87
GGTCAATGATGAGTACTTTATATTTCCTTGAGGAACGGTGATGACTTAAATATCACATACAATGGCATGACACTG
AGTCCC
>SNORD115-88
GATCAATGATGAAATCTTTGCTTTGTCTCTAGAGAGGTGTTGTCTTAAATGTCTTGCACAAAGAGTTACGCTGAG
GCCC
>SNORD115-89
GGTCAATGATGAAATCTTTGCATTGTCTCTAGACAGGTGTTGTCTTAAATCTTGCACAATAGAGTTACACTGAGG
TCC
>SNORD115-90
GGTCAATGATGAAATCTTGCCTTGTCTCTAGAGAGGTGTTATTTTAAAAGTCATGCGCAATATAGTTATGCTGAG
GACC
>SNORD115-91
GGTCAATGATGAAATCAATGCCTTGTCTCTAGAGAGGTGTGGTCTTAAAACTCTTGCCCAATCGTGTTATGCTGA
GGACC
>SNORD115-92
GGTCAATGATGAGTACTTTATATTGCCTTATGGATCAGTGATGACTTAAATATCACATACAAGGCATGACGCTGA
GGCCC
>SNORD115-93
TGTCAATGATGAAATATTTCCCTTGTCTCTAGAGAGGTGCTGTCTTAAAAGTCTTGTGCAATAGAGTTACACTGA
GGCTC
>SNORD115-94
GGTCAATGATGAATACTTCATGTTGCCTTGAGGAGCAGTGATGACTTGAATATCACTTACAATGGCATGATGCTG
AGGCAA
>SNORD115-95
GGTCAGTGATGAAATCTTTGTCTTGTCTCTACAGAGGTGTTGTCTTAAATGTCTTGTGCAATAGAGTTACGCTGA
GTAAC
>SNORD115-96
GGTCACTGATGAGCATTTTATATTGCCTGGAGGAGTGATGATGACTTAAAGTTCACACATTATGCTGAGGCCCAG
CCTGAATA
>SNORD115-97
GGTCAATGATGAGCACTTCATACTGCCTTGAGGACCAGTGATGACTTAAATACCGTGAACAATGGCATGACGCTG
AGACCG
>SNORD115-98
GGCCAATGATGAGTTTATATTGTTTTGTGGAGCAGTGATGACTTAAGTATCACGTACAATGGCATGACGCTGAGG
CCC
>SNORD115-99
GGTCAATGATGAAATATTTGCCTTGCATCTAGAGATCTGTTGTCTTAAAAGTCATGCTCAGTAAATTTATGCTGA
GGACC
>SNORD115-100
GGTCAATGATGAAATTCTTGCCTTGTCTGTAGAAAGGTGTTGTGTTAAAATTCATGTGCAATAGAGTTACACTGA
GGACC
>SNORD115-101
GGTCATGATGATTACTTTTTTTTTTGGCTTGAGGAGTGGTGATGACTTAAATATCACGTACAATAGCATGACACT
GAGGCCC
>SNORD115-102
GGTCAATGATGAAATCTTTTCCTTGTCTCTAGAGAGGTGTTGTTTCAAAAGTCTTGCATAATAGAGTTACTCTGA
GGACC
>SNORD115-103
GGTCAATGATGAGTACTTCATGTTTCCTTGAGGAGCAGTGATGACTAAATATCACATACAATGGCATGACGCTGA
GGCCC
>SNORD115-104
GATCAATGATGAACCTTTATATTGTCATAAAGAAAGTTGATGACATGAAAATTATGACAAAGAAGACTGTTCTGA
GAACC
>SNORD115-105

GGTCAATGATGAATACTTTATGCTGCCTTGAGGAGCAGTGATCACTTGAATATCACATACAATGGCATGACGCTG
AGGCCC
>SNORD115-106
GGTCAATGATGAAATCTTGGCCTGGTCATTAGAGAGGTGTTGTCTTAAAAGTCATGCACAATAGAGCTACACTGA
GCACC
>SNORD115-107
GGTCAATGATGAGTACTTTATATTGCCTTGAGGAACGGTAATGACTTCAATATTAATACAATTGCATGACACTGA
GACCC
>SNORD115-108
TGTCAATGATGAAATATTTGCCTTTTCTCTAGAGAGGTGTTGTCTTAAGAGTCTTGCACAATAGAGTTACACTGA
GGACA
>SNORD115-109
TGTCAATGATGAAATCTTTGCCTTGTCTCTAGAAAGGTGTTGTCTTAAGTGTCCTGCACAATAGAGTTACACTGA
GGACT
>SNORD115-110
GATCAGTGATGAAATCTTTGCCTCGTGTCTAGAGAGGGGTTGTCTTAAAAATCTTGCACAATAGAGTTACACTGA
GGACC
>SNORD115-111
GGTCAATGATGAGGACTTAAATTGCCTTGAGGTGCAGTGATGACTTAAATATCGCATACAATGGCCTATCGCTGA
GGCCC
>SNORD115-112
GGCCAATGATGAGGAGTATATATTGCCTTGAGGAGCAGTGATAACTTAAATACCACATACAATGGCATGACACTG
AGGCCC
>SNORD115-113
GATCAATGATGAAATCTTTGCCTTCTCTCTAGAGAGGTGTGGTTTTAAGAGTCTTGCTCAATGGAGTTAGGCTGA
GTACC
>SNORD115-114
GGTCAATGATGAGTAGTTTATATTGACTTTAGGGGCAGTGATGACTCAAATATCAGGTACATGGCATGACACTGA
GGCTC
>SNORD115-115
GGTCAATGATGAAATCTTTGCCCAATCTCTAGAGAGTTGCTGTCTTAAAAGTCATGCAAAATAGAGTTGCACTGA
GGACC
>SNORD115-116
GGTCAGTGATGAAATCTATGCCTTGTCTCTAGGGAGGTGTTGTCTTAAGAGTCTTGTGCAATTGACTTACGCTGA
GGACC
>SNORD115-117
GTTCCATGATGATATTTTTGCCTTCTCTCTGGAGAGGAGTTGTCTTAAAAGTCATTCTCTATAGAGTTACACTGA
GGACC
>SNORD115-118
GGTCAATGATGAGTACTTTATATTGCCTTGAGGAATGGTGATGACTTCAATATCAAACACAATGACCTGATGCTG
TTGCCC
>SNORD115-119
GGTCAATGATGAGCATTTTATATTGCCTGGTGGAGTCATGATGACTGAAAGTTCACGCATGACACTGAGGCCCAG
GCTAGGTA
>SNORD115-120
GGTCAATGATGAGCATTTTATATTGCCTGGAGGATTGGTGATGACTGAAAGTTCACGCATGACGCTGAGGCCCAG
CCTAGGTA
>SNORD115-121
GGTCAATGATGAGCGTTTAATATTGCCTGGAGGAGCGGTGATGACTGAAATTTCATGCATGACACTGAGACCCAG
CCTAGGTA
>SNORD115-122
GGTCAATGATGAGCATTTTATATTGTCTGGAAGAGCAGTGGTGACGGAAAGTTCATGCATGATGCTGAGGAGCAG
CCTGGTGA
>SNORD115-123
GGTCAATGATGAGCATTTTATATTGCCTGGAGGAGCAGTGATGACTGAAAGCTCACGCATGATGCTGAGGCCCAG
CCTAGGAA
>SNORD115-124

GGTCAATGATGAGCATTTTATATTGCCTGGAGGAGCGGTGATGACTGAAAGCTCATGCATGACGCTGAGGCCCAG
CCTAGGTA
>SNORD115-125
GGTCAATGATGAGCATTTTAAATTGCCTGGAGGAGTGGTGATGACTGAAAGATCACGCATGATGCTGAGGCCCAG
GCTAGGTA
>SNORD115-126
GGTCAATGATGAGCATTTTATATTGTCTGGAGGAGCGGTGATGACTGAAAGTTTGACATGATGCTGAGGCCCAGC
CTAGGTGA
>SNORD115-127
GGTCAATGATGAGCATTTTATATTGTCTGGAAGATCGGTGATGACTGAAAGTTCACGGATGACACTGAGGGCCAG
CCTACGTA
>SNORD115-128
GGTCAATGATGAGCATTTTATATTGCCTGGAGGAGCGGTGATGACTGAAAGCTCATGCATGACGCTGAGGCCCAG
CCTAGGTA
>SNORD115-129
GGTCAGTGATGAAATCTCTGTGTTGTCTCTACAGGTGTTGTCTTAAAAGTCTTGCACAATAGAGTTATTCTAAGG
ACC
>SNORD115-130
GGGCAATGATGAACCCTTTTCCTTAATTCTAGAGATGCGTTGTCTTAAAAGTCATGCACAATATAATTATGCTTA
GGACC
>SNORD115-131
GGTCAGTGATGAAACCTTTACATTGTGTGTAGATATGTGCTGACTTAAAGTTCATGCTCAATGGAATTACGATGA
GACCC
>SNORD115-132
GGTCAATGATGAGTACTTTATACTGCCTTGAGGAGTGATGGTGACTTAAATATCATGTGCAATGGCATGACAATG
AGGCCC
>SNORD115-133
GGTCAATGATGAGTACTTTATATTGCCTTGAGGAGAGATGATGATTAAATATCATGTACAAGCCGGGCTCGGTGT
CTC
>SNORD115-134
GGTCAATGATGAAATTTTTGCTTTGTCTCTAGAAAAGTGTTGTCTTAAAAGTCATGCTCAATAGAGTTACGCTGC
AGACC
>SNORD115-135
GGTCAATGATGAGTATTTTATATTACCTTGAGAAGCGGTGATGACTTAAATATCACATACAATGGCATGAAGTTG
AGGCCC
>SNORD115-136
GGTCAATGATGAGAACTTTATATTGCCTTGAGGAGCATGATGACTTAAATATCACACACACGCTGGAGCCCAGCA
GGTCG
>SNORD115-137
TGTCAATGTTGAGTACTTTATATTGCCTTGAGGAGCGGTGATGACCTAAATATCAAGTACACTGGCATGACACCG
TGGCCC
>SNORD115-138
GGTCAATGATGAGTACTTTATGTTTCCTTGAGGAGCAGTGGTTACTTAAATATCACATACAATGGCATGATGCTA
AGGCCA
>SNORD115-139
GGTCAATGATGAGCATTTTATATTGCCTGGAGAAGCGGTGATAACTTAAATTTCACGCGTGACACAGAGGCCCAG
CCTAGGTT
>SNORD115-140
GGTCAATGATGAAATCTTTGCATTGTATCTAGAGAGATGATGACTTCAAAGGTATGCTCAGTGGTGTTACATGAG
GACC
>SNORD115-141
AGTCAATGATGAGACTTTATTTGGTCTTAAAGACAGAAGATCACATAAAAATGATGGTCAATAAAAACTCTGGGG
ACC
>SNORD115-142
TGTCAATGATGAAATATTTGCCTTGTCTCTAGAGAGGTGTTGTCTTAAGAATCTTGCAAAATAGAGTTACGTTTA
GGACC
>SNORD115-143

GGTCAGTGATGAAATATTTTCTTTGTCTCTAGAGAGGTGCTGTCTTAAAAGTCTTGTGCAATAGAGTTACGATGA
GAACC
>SNORD115-144
GGTCCATGATGAAATCTTTGACTTATTCTAGAGATGTGTTGTCTTAAAAGTCATGTTCAATATAGTTATGCTTAG
GACA
>SNORD115-145
GGTCAATGATGAAATCTTTGCCTTTTCTCTAGAGAGGTGTTGTCTTAAAAGTTTTGTGCAATGGAGTTATGTTGA
AGACC
>SNORD115-146
GGTCAATGATGAAATCTTTGTCTTGACTCAAGTGAGGTGTTGTCTTGAAAGCCTTGTGCAATAGAGTGATAGTGA
GGACC
>SNORD115-147
GGTCAATGATGAGTACTTTATATTGCCTTGAGGAGAGGTAATGACTTAAATATCATGTACAATGGCATGACGCTG
GGGCCC
>SNORD115-148
TGTCGATGATGATAATGCTATATTATCTTGAAGAGAAGTGATGACTTAAAAAATCATGCTCAATAAGAATTTGCT
AAGATCC
>SNORD115-149
GATCAATGATGAGAACTTTATATTTTCTTGAAGAGAGGTGATGACATAAAAATCATGCCCAAGTTGAGGCCCAGC
CCCAGTGA
>SNORD115-150
GGTCAGTGATGAGAACTTTATATTGTCTGAAGAGAGATGATGGCTTAAACATCATGCTCAATATTGTCATCCAGC
CTAGATGA
>SNORD115-151
GGTCAATGATGAGAACTTTATACTGTCTTGAAGAGATGTGATGACCTAGAAATCATGCTCAACAGGATTACTTTG
AGGCCC
>SNORD115-152
GGTCAATGATGAGAACTATGTATTGTCTTGAAGAGAAGTAATGACTAAGAAATCATGCTCAATAAGATTATGTTG
AGGCCA
>SNORD115-153
GGTCAATGATGAGATGTTTACATTGTCTTAGGAGAGGGGATGACTGAAAAATCATGCTCAATAGGATTACACAGA
GGCCC
>SNORD115-154
GAACAATGATGAGGACTTTATATTGTCTTAAAGAGAGGTGATGACTTAAAAATCATGCTCAATAGCATTGCACTA
TGCCCA
>SNORD115-155
GGTCAATGATGAGATATTTGTATAGTCTGAAGAGAGTTGATGACTTAAAAAATCATGCTCAATAGAATTATACTT
AGGTTC
>SNORD115-156
GGGCAATGATGAAACCTTTGCTCTAATTCTAGAGATGTGTTGTCTTAAAAGTCATGCAAAATATAATTACGCTTA
GGACC
>SNORD115-157
GGTCAATGATGAAAACTTCATATTGTCTGGAGAGCAGTGATGACTTAAAAATCATGCTCAATAGGATTATGATGA
GGCCT
>SNORD115-158
GGTCAATGATGAGAACTTTATATTGTCTAGAGGAGAGATGATGACTTAAAAATCATGCTCAATATGATTATACTG
TGGCCC
>SNORD115-159
GGTCAATGATGAGAACTTTATATTGTCTAGAAGAGAGATGATGACTTAAAAAGCATGCTCAATAGAATTATGCTG
TGTCCC
>SNORD115-160
GGTCAATTACAAGAACTTTACATTGTCTGAAACCTGGAGATGATTTATAAATCATACCCAATAAGATTAT
GCTGAGTCCC
>SNORD115-161
GGTCAATCATGAGGAATTTATATTGCCTTGAGGAGAGGTGATGACTTAAATATCATGTACAATGGCATGA
CACTGAAGCCC
>SNORD115-162

GGTCAAAGATGGGAATTTTATATTGCCTTGAGGAGAGGTGATGACTTAATTATCATGTACAATGACATGA
CGCTGAGGCCC
>SNORD115-163
CATCAATGGTGAGGACTTCATATTGCCTTGAAAAGTGGTGATGACTTAAATATCAAGTACAATAGCATGTCG
CTGAGGCCC
>SNORD115-164
GGTCAATGGTGAAATCTTTGCATTCTCTAGAGAGGTGATGGCTTAAAAGTCATGCTCAATGGAGTTATGT
TGAGGACC
>SNORD115-165
GGTCAAAGATGAGGAGTCTATATCACACTGAGGAGCGGTGATGACTTAAATATCATGTGCAATGGCATGA
TGCTGAGGCCC
>SNORD115-166
GGTCAATGATAGTACTTTATATTGCCTTAAGGAAGAGTGATGACTTAAATATAAAGTACAATGGCATGAT
GCTGAGGCCC
>SNORD115-167
GGTCAATGATTAATCCTTTGCATTGTCTACATAGAGGCCATGACTTAAAAGATATACTCAATAGAATTAT
GCTGAGGACC
>SNORD115-168
GGTCAATTATGAAATCTTTGCCTTGTCTCTAGAGATGTGTTATCTTAAAAGTCATGCACAATAGAGTTAT
GCTGAGGATC
>SNORD115-169
GGTCAATAATGAGTACTTTATATTGCCTTGAGAAGTGGTGATGACTTAAACATCAAGTACAATGGCATGA
CACTGAGGCGC
>SNORD115-170
GGTCAGCGATGAAATCTTTGTCTTTTCTCAAGAGTGGTGTTGACTTAAACATCTTGTGCAATAGGGTTAC
GCTGAGGACA
>SNORD115-171
GGTCAGTGGTGAGGACTTTATAATGTCTTGAGGAGAGGTGATGACTTAAATATCATGTACAATGGCATGA
CACTGAGGCCC
>SNORD115-172
GGTCAATTATGAGATTTTTATATCGGCTGAAGAGAGGTGATGACTTAAAAATTATGCTCGCCCGTGGGTTCGAGA
CTGGCCTG
>SNORD115-173
GGTAAATGATTATCTTTATATTGTTTGAAGAGAGGTGATGACTTAAAATGATGCTCAGTAGGAATACACT
GAGGTCC
>SNORD115-174
GGTCAATAATGAGGTCTTTATATTGTCTTGAAGAAAGTTGATCACTTAAAATTCATGTACAATGACATGA
TGCTGAGGCCC
>SNORD115-175
GGTTAATTATGAGAACTTTGCATTGTCTTTAGAGACGTGATGACAAAAGTCATGCTCAATAGGATTACAC
TGAGACCC
>SNORD115-176
GGTCAATGATAAGTACTTTATATTGCCTTGAGGAGAAGTGATGACTTAAATATCAAGTACAATGGCATGA
TGCTGAGGCCC
>SNORD115-177
GGTCAATAATGAGGACTTTATATTGCCTTGAGGAGTGGTGATGACTTAAATATCATGTACAATGGCATGA
TGCTGAGACCC
>SNORD115-178
GGTCAATGATGGGAACTTTGTATTGTCTTTAATAGAGGTGATGACTTAAAAATCATGTTCAATAACATTACACTG
CCCC
>SNORD115-179
GGTCAATGATGGGAACTTGATATTGTCTGAAGAGTGATGATAATGTAAAAATCATGCTCAATAGGACTAC
AATAAGGCCC
>SNORD115-180
GGTTTATGACGAGAACTTTATATTGTCTTGAAGAGAGTTGATGACATAAAAATCATGCTCAGTGAGGTTATGCTG
AAGCCC
>SNORD115-181

GGTCAATGCTGAGACTTACATTGTCTGTACATAGGTGATGACTTAAAAATCATGCTCAATAGGATTACAC
TGAGGTCC
>SNORD115-182
GGTCAATGACGAATACTTTATATTGCCTTGAAGAGAGGTGATGACTTAAATATCAAGTATAATGGCACAA
CGCAGAGGCCC
>SNORD115-183
GGTCCACGATAAGTACTTTATATTGCCTTGAGGAGTGGTGATGACTTAAATATCAAGTACAATGGCATGA
CGCTGAGGCCC
>SNORD115-184
GGTCAATGAAGAGAACTTAATATTGCCTTGAGGAACGGTGATGATTTCAATATCAAGTACAATGGCATGA
CGCTGAGGCCC
>SNORD115-185
GGTCCATGATGTGTACTTTATGTTGCCTTGAGAAGCGGTGATGACTTAAATATCACATACAATGGCATGA
TGCTGAGGCCC
>SNORD115-186
AGTCAATAATGAAATCTTTGTCTTGTCTCTAAAGAGAAGTTGTCTTAAAAGTCATGCATAATAGAGTTATG
CTGAGGACC
>SNORD115-187
GGTCAATGTTGAGTACTTTATATTGCCTTGAGAAGAGGTGATTTTTAAATATCAAGTACAATGGCATGA
CGCTGAGGACC
>SNORD115-188
GGTCAATTATGCCTACATTATATTGCCTTGAGGAGAGGTGATGACTTAAATATCAAGTACAATGACATGA
AACTGAGGCCC
>SNORD115-189
CATCAGCGATGAAATCTTTGTTTTGTGTCTACAGAGGTGTTGTCTTAAAAGTCTTGCACAATAGAATTATGCTAA
GG
TCC
>SNORD115-190
GGTCAATGTTGAGATGTTATATTGTCTTGAACAGGGATGATGACATAAAAATTATATTCAATGGAATTAT
GGGAGAGCCA
>SNORD115-191
GGTCAATGGGAAAATCTTTGACTTGTCTCTAGAGAGGTGTTGTCTTAAAAGTTATGCTCAATAGAGTTAC
ACTGAGGACC
>SNORD115-192
GGACAGTGATGTAACCTTTGCCTTGTCTCTAGAGAGGTGTTGTCTTAAAAGTCTTGCACAATAGAGTTAC
GCTGAGGACC
>SNORD115-193
GGTCAATGAAGAAATCTTTGTCTTGACTCTAGTGAGGTCTTGTCTTAAAAGTCTTGTGCAATAGAGTTAT
GCTGAGGACC
>SNORD115-194
GGTCAATAATGAAATCTTGGCCTTGTCTCTAGAGAGCTGTTGTCTGAAAAGTCATGCACAATAGAGTTAC
GCTGAGGACC
>SNORD115-195
GGTCAAAGATAAAATCTTTGCCTTGTCTCTAGAGAGGTGTTGTCTTTAAAGTCATGAGCAATAGAGTTAC
GCTGAGGACC
>SNORD115-196
GGTCAATCATGAGGACTTTATCTTGTCTTGAGGAACGGTGATGACTTTAATATCACATACAATGGCATGA
CGCTGAGATCC
>SNORD115-197
GGTCAATGATCAGACCTTTGCCTTGATTCTAGAGATGTGTTTTCTTGAAAGTCATGCTCAATATAATTAT
GCTTAGGACT
>SNORD115-198
GTTCAAATATGCAATCTTTGCATTGTCTATAGAGAGGTTATGACTTAAATGTCATGTTCAATGGAATTA
CACTGAGATCT
>SNORD115-199
GGCCAATGGTAAGTACTTTATATTGCCTTGAGGAGAGATGATAACTTAAATATCATGTACAAGGCATGAT
GTTCAGGCCC

>SNORD115-200
GGTCAATATTGAGGACTTTATATTACCTTGAGAAGCGGTGATGACTTAAATATCACGTACAGTATCATGA
CGCTGAGACCC
>SNORD115-201
GGTCAACAATGAGTTCTTAATATTGCCTTGAGGAGCGATGTTGACTTAAATATCAAGTACAATGGTATGA
CGCTGAGGCCC
>SNORD115-202
GGTCAATGGTTAGTACTTTATATTGACTTGAGGAACGATGATGACTTCAACATCGAGTACAATGGCATGA
CGCTGAGGCCC
>SNORD115-203
GGTCAATCATGTGTACTTTATATTGCCTTGAGGAGCAGTAATAACTTAAGTATCATGTACAATGGCATGA
CGCTGAGTCCC
>SNORD115-204
GGATCAGGATGAAATCATTGTCTTGTCTCTATAGAGGTGTTGTCTTAAAAATCTCGTGCAATAGAGTTATGCTGA
GG
ACC
>SNORD115-205
GGTCAGGCGTGAAATCTTTGTCTTGTCTCTAGAGAGGTGTTGCCTTAAAAGTCTTGCGCAATAGAGCTACGCTGA
GGACA
>SNORD115-206
TGTCAATGATCAAAACTTTGACTTGATTCTAGAGATGTGTTGTCTTAAAAGTCATGCTCAATATAGTTACACTTA
GGACC
>SNORD115-207
GGTCAATAATGGGCATTTTATGTTGCCTGGAGGAGCAGTGATGACTTAAATATCATGCACGACTCTGAGGACCAG
TCTAGGTGA
>SNORD115-208
GGTCAATGATAAGCATTTTATATTGCCTGGAGAAGTGGTGATGACTGAAAGCTCACGCATGACGCTGAGGCCCAG
CCTAGGTGA
>SNORD115-209
GGTCAAATATTAAGTCTTTACCTTGTCTCTAGAGAGGTGTTGTCTTAATGGTCATGCACAATAGCGCTA
CGCTGAGGACC
>SNORD115-210
GGTCACCGATGAAATCTTTGTATCTTGTCTAGAGAGGTGTTATCTTAAAATTCTTGCACAATACAATTAC
ACTGAGGACA
>SNORD115-211
GGTCAATGATCAAACCTTTGACTTGATTCTAGAGATGTGCTGTCTTAAAAGTCATGCTCAATACAGTTAACGCTT
AGGACC
>SNORD115-212
GGTCAATGATCAAATCTTTCCTTGATTCTAGAGATGGGCTGTCTTAAAAGTCATGGTCAATATAATTATG
CTTAGGACC
>SNORD115-213
GGTCACTAATAAATTCTTTATATTGCCTTGAGGAGCGGTGATGACTAAAATATCACGTACAATGGCATGACACTG
AGGCCT
>SNORD115-214
GGTCAATAATGAAATCTTTGTCTTGTCTCTAGAAGGTGTTGTCTTAAAAGTCTTGCTCAGCCAGGCTCAGTGGCT
C
>SNORD115-215
GCTCAATTCTGAAATATTTGCCTTGTCTCTAGAGAGGTGTTGTCTTAAAAGTCATGCACAATAGAGCTTCGCTGA
GGACC
>SNORD115-216
AATCAATGATAAGACCTTATATTCTCTCTATGAGAAAAAAATGACTTAAAATTATTCTCAGTAGGATTATGC
TGAGTCTC
>SNORD115-217
GGTAAATGATCAAACCTTTGTCTTGATTCTTGAGATGTGTTGTCTTAAAAGTCATGCTCAATATAATTAC
ACTTAGGACC
>SNORD115-218
GGTCAACATGTGTACATTATATTGCCTCGAGGAGTGGTAATGTGTTAAATATCAAGTACAATGGCATGAA

GCTGAGGCCC
>SNORD115-219
GGACGGGCATAAAATCTTTGTCTTGTCTCTAGAGAGGTGTTGTCTTAAATGTCTTGCACAATAGAGTTACTCTGA
GGACA
>SNORD115-220
GGTCAGTGATTAAATCTATGTCTTGTCTATAGACAGGTGTTGTCTTAAAAGTCTTGCACAATAAAGTTACACTGA
GGACA
>SNORD115-221
GGTCAGCAATGAAATCTTTGTCTTTTCTCTAGAGAGGTGTTGTCTTAAAAGTCTTGTGCAATAGAGTTAC
TCTGAGGACA
>SNORD115-222
GGTCAATAATCAAACTTTGCCTTGATTCTAGAGATGTGTTGTCTTAAATGTCATGCTCAATATAGTTAC
GCTTAGGACC
>SNORD115-223
GGCCAGCGATGAAATCTTTGTCTTTTCTCTAGAGAGGTGTTGTCTTAAGAGTCTCGCACAATAGCATTACACTGA
GGACA
>SNORD115-224
GGTGAGTAATGAAATCTTTGCCTTGTCTCTACAGAGGTGTTGTCTTCAAAGTCTTGCACAATAGAGTTAT
GCTGCAAACT
>SNORD115-225
GGTCAATGATCAAACATTTGCCTTGATTCTAGAGATGTGTTGTCTTAAAAGTCATGCTCAATGTAATTAA
GCTTAGTACC
>SNORD115-226
GGTCAATGGTGAGCATTTTATATTGCATGGAGGAGCGGTAATGACTGAAAGTTCATGCATGACAATGAGGCCCAG
CCTAGGTGA
>SNORD115-227
GTTCAATGATCAAATATTTGCCTTATCTCTAGAAAGGAGTTGTCTTAAAAGTCATGCTAATAGACTTATGCT
GGGGACC
>SNORD115-228
GGTCAATAATCAAACCTTTGACTTGATTCTAGAGATGTGTTGTCTTAAAAGTCATGCTCAATATAGTTACACTTA
GTGCT
>SNORD115-229
GGTCAATAATAAAACCTTTGCCTTGTTTCTAGAGATGTGTTGTCTTAAAAGTCATGTTCAATATACTTACACTTG
GGACC
>SNORD115-230
GGTCAATAATGAAATCTTTGCCTTGTCTCTAGAGAACAGTTGTCTTAAATGTCATGCTCAATATAGTCACCCTGA
GGACC
>SNORD115-231
GGTCAAAGATGAACATTTTATATTGCCTGGAGGAGCGGTGATGACTGAAGGCGCTGAGGCCCAGTCTAGGTGA
>SNORD115-232
TGTCAATGATCAAACCTTTGACTTGATTCTAGAGATGTTTTCTTAAAAGTCATGCTCAATATAGTTATGCT
TAGGAAC
>SNORD115-233
GGTCAATGGTAGATCCTTTTCCTTGTTTTTAGAGAGGTGTTGTCTTAAAATTCATCCTCATTATAGTTAT
GGTAAGGACA
>SNORD115-234
GGTCAATGATCATATTTTATATTGCCTGGAGGAATGGTGATGACTGAAAGTTCATGCATGATGCTAAGGCCCAGT
CTAGGTAT
>SNORD115-235
GGTCAGCGATGAAATCTTTGTATTTTCTCTAGAGAGGTGTTGTGTTAAACAAATTGTACAATAGAGTTAC
GCTAAAGATG
>SNORD115-236
CATCAGTAATGAAATCTTTGTCTTGTCTCTAGAGAGGTGTTGTCTTAAAAGTCTCGTGCCATTGAGTTACGCTGT
GGACC
>SNORD115-237
GGTCAGCGATTAAATCTTTGTCTTTTCTCTAAAGAGGTGTTGTCTTAAATGTCTTGCGCAATGGAGTTAC
GCTGAGAACT

>SNORD115-238
GGTCAATGATCAAACCTTTGACTTGAGTCTAGAGATGTGTTGTCTTAAGAGTCATGCTAAATATAGGTACGCTTA
GGACC
>SNORD115-239
GGTCAACGATTAAACCTTTGACTTGATTCTATAGATGTGTTGTCTTAAAAGTCATGCTCAATATAGTTACGCCTA
GGACC
>SNORD115-240
GATCAAAGATGTGTACTTTATATTGCCTTCAGGAGTGGGAATGACTTAAATATTATGTACAATGGCATGATG
CTGAGGCCC
>SNORD115-241
GGTCAGCGATGAGTAATTTTATTGCCTTGAGGAGCAGTGATGACTTAAATATCATGTACAATGGCATGAC
GCTGAGGCCC
>SNORD115-242
GGTCAATAATGAGTACTTTATATTGCCTTGAGGAACGCTGATGACTTAAATGTCACGTACAACAGCATGA
CGCTGAGGCCG
>SNORD115-243
GGATGATGGTGGGAACTTTATATTGTCTTAAAGAGTGGTGATGACTTAAAAAGCATGTCTAGGTGACAATTGTGG
AAA
>SNORD115-244
ATGGAAAGGTGAGGACTTTGTATTGTCTTGAAAAGAGGTGATGATTTAATATTCAAGTACAATGGCATGATGCTG
AGGT
CC
>SNORD115-245
GGTAAATGCCAAAATCTTTGCATCGTCTGTTGAGAGGTGATGATTTAAAGACATGCTCAATAGAATTTTGCTGAG
GACT
>SNORD115-246
GGTCAATGATCAAACCTTTGACTTGATTCTAGAGATGTGTTGTCTTAAAAGTCATGCTCAATATAATTAT
GCTTAGGACC
>SNORD115-247
GGTAAACAATGAGAACTTTATATTGTCTGAAGAGAGGTGATGACTTAAAAATCATGCTCAATAGGATTAC
ACTGAGGCCC
>SNORD115-248
GGTCAATTATGAGAACTTTATATTGTCCTGAAGAGAGGTGATGACTTAAAAATCATGCTTGATAGGATTA
TGCTGAGGCCC
>SNORD115-249
GGTCAATAATGAGAACTTTATATTGTTTTGAAAGGAGGTGATGACTTCAAAATCATGCTCAATAGGATTA
TGCTGAGGCCC
>SNORD115-250
GGTCAATAATGAGAACTTTATATTGTCTAGAACAGAGGTGATGACTTAAAAATCATGCTCAATAGGATTA
CGATGAGGGCC
>SNORD115-251
GGTCAATGATAAGATCTTTATATTGTCTGAAGAGAGATGATGACTTAAAAACTATGCTCAATAGGATTGC
GCTGAAGTCC
>SNORD115-252
GATCAGTGCTGAGAACTTTATATTGTCTGAAGAGAGGTGATGACATAATAATCATGCTCAATAGGAATACGC
TGTGACCT
>SNORD115-253
GGTGAACGATGAGAACTTTATATTGTCTGAAGAGAGGTGATGACTTAAAAATTGTGCTCAGTAAGATTAC
ACTGAGGCCC
>SNORD115-254
TAACAGTGATAAGAACTTTATATTGTCTAGAAGAGAGGTGATGACTTAAAAATCATGCTTAATAGGATTTCAC
TGAGGTCC
>SNORD115-255
GGTCAATGATAAGGACTTCATATTGTCTTGAAGAGAAGTGATGACTTAAAAATCATGTTCGGTGGGATTA
TGCTGAGGCCT
>SNORD115-256
GGTAAATGATAAGACCTTTGTATTGTCTGTAGAGAGGTTATGACTTAAAAGTCATGCTCAATAGGCTTAT

GCTGAAACCC
>SNORD115-257
GGTCAATGATTAGAATTTTATATTGTCTTAAAGAGAAGTGATGGCTTAAAAATCATGCTCAGTAGTTTTA
CACTGAGGCCC
>SNORD115-258
GGTCAGTGTTGAGAACTTTATATTGTCTTGAAGAGAGGTAATGACTTAAAAGTCACGATCAATAGAATTA
CGCTGATGTCC
>SNORD115-259
GGGCAAAGATCCAACCTTTGCCTTAATTCTAGAGATGTGTTGTCTTAAAAGTCATGCTCAATATAATTACGCTTA
GGACC
>SNORD115-260
GGGCAAAGATCCAACCTTTGCCTTAATTCTAGAGATGTGTTGTCTTAAAAGTCATGCTCAATATAATTACGCTTA
GGACC
>SNORD115-261
GGTCAATGATTAAATCTTTGCCTTGTCTCTATAGAGTTGTTGTCTTAAAAGTAATGCGCGATAGAGTTAC
ACTGAGGACC
>SNORD115-262
GGTCAAAGATGTAGTCTTTGCCTTGTCACTAGAGAGGTGCTGTCTTAAAAGTCATGTGCAATAGAGTCAC
GCTGAGGACC
>SNORD115-263
GGTCAATTATGAGTTTTTATATTGCCTTGAGGAGGGGTGATGACTTAAATATCAAGTACAATGGCCTGAT
GCTGAAGCCC

### Cluster of SNORD115 in *M. auratus* (Golden hamster)

>SNORD115_1
GGGTCTATGATGAGAAACCTACGTCCTGAAGAGAGGTGATGACATAAAAATCATGCTCAGTCTCGGTCTGCTGAG
GCCC
>SNORD115_2
GGGTCTATGATGACAAACTAATGTCATGAAGAGAGGTGATGACAACAATATCATGTTCAGTATGGTTCTGCTGAG
GCCC
>SNORD115_3
GGGTCTATGATGAGAAACCTACGTCCTGAAGAGAGGTGATGACATAAAAATCATGCTCAGGTTCCGGCTGCTGAG
GCCC
>SNORD115_4
GGGTCTATGATGAGAAACCTACGTCCTGAAGAGAGGTGATGACATAAAAATCATGCTCAGGTTCCGGTTGCTGAG
GCCC
>SNORD115_5
GGCTATATGATGAAAAACCAATGTCATGAAGAGAGGTGATGACATAAAAATGATGCTCAATAGCATTACGCTGAA
GACC
>SNORD115_6
GGGTCTATGATGACAATCCAATGTCAGGAAGAGAGGCGGTGACATAAATATCATGCTTAGTATGGTTCCGCTGAG
GCCC
>SNORD115_7
GGGACTATGATGACAAACCATTGTCATGAACTCAGTTGATGATATAAAAACCATGCTCAATAGGATTACACTGAG
GCAC
>SNORD115_8
GGGTCTATGATGACAAACCAATGTCATGAAGAGAGGTGATGACAACAATATCATGTTCAGTATGGTTCTGCTGAG
GCCC
>SNORD115_9
GGGTCTATGATGACAAACCAATGTCATGAAGAGAGGTGATGACAACAATATCATGTTCAGTATGGTTCTGCTGAG
GCCC
>SNORD115_10
GGGTCTATGATGACAAACCAATGTCATGAAGAGAGGTGATGACATAAAAATCATGCTCACTTGGATTACGCTGAG
GCCC

>SNORD115_11
GGGTCTATGATGACAAACCAATGTCATGAAGAGAGGTGATGACAACAATATCATGTTCAGTATGGTTCTGCTGAG
GCCC
>SNORD115_12
GGGTCTATGATGAGAAACCTACGTCCTGAAGAGAGGTGATGACATAAAAATCATGCTCTGTCTCGGTCTGCTGAG
GCCC
>SNORD115_13
GGGTCTATGATGACAAAAAAATGTCATGAAGAGAGGTGATGACAACAATATCATGTTCAGTATGGTTCTGCTGAG
GCCC
>SNORD115_14
GGGTCTATGATGACAAACCAATGTCATGAAGAAAGGTGATGACAATAATATCATGTTCAGTATGGTTCTGCTGAG
GCCC
>SNORD115_15
GGGTATATGATGACAAACCAATGCCATGAAGAGAGGTGATGACGTCAAATTCATGCTCAGAAGGGATACGCTGAG
GCCC
>SNORD115_16
GGGTCTATGATGATAAACCAAAGTCATGAAGAGAGGTGATGACATAAAAATCATGCTCACGGGGTTCCGCTGAGG
CCG
>SNORD115_17
GGGTCTATGATGACAAACCAATGTCATGAAGAGAGGTGATGACATAAAAATCATGCTCACTTGGATTACGCTGAG
GCCC
>SNORD115_18
AGGTTTATGATGACAAACAAATGTCATGAAGAGTGCTGATGACATAAAAATGATGCTCAATGAGATTTTGCTGAG
GCCC
>SNORD115_19
GGGTCTATGATGAGAAACCTATGTCCTGAAGAGAGGTGATGTCATAAAAATCATTCTCAGATTCCGGTTGCTGAG
GCCC
>SNORD115_20
GGGTCTATGATGACAAACCAATGTCATGAAGAAAGGTGATGACAATAATATGATGTTCAGTATGGTTCTGCTGAG
GCCC
>SNORD115_21
GGGTCTATGATGACAAACCTACGTCATGAAGAGAGGTGATGACATAAAAATGATGCTCAGGTTTCAGGCTGCTGA
GGCCC
>SNORD115_22
GGGTCTATGATGACAAACCAAAGTCATGAAGAGAGGTGATGACATAAATATCATGCTCAGGTTCTGGCTGCTGAG
GCCC
>SNORD115_23
GGGTCTATGATGACAAACCAAAGTCATGAAGAGAGGTGATGACATAAATATCATGCTCAGGTTCTGGCTGCTGAG
GCCC
>SNORD115_24
GGGTCTATGATGACAAACCAAAGTCATGAAGAGAGGTGATGACATAAATATCATGCTCAGGTTCCGGCTGCTGAG
GCCC
>SNORD115_25
GGGTCTATGATGACAAACCAATGTCATGAAGAGAGGTGATGACATAAAAATCATGCTCACTTGGATTACGCTGAG
GCCC
>SNORD115_26
GGGTCTATGATGACATACTAATGTCCTGAAAAGTGCTGATAACATATAAATCAGCCTAAATAGGATAACACTGAA
GCTC
>SNORD115_27
GGGTGTATGATGAGAAACCTACGTCCTGAAGAGAGGTGATGACATAAAAATCATGCTCAGGTTCCGGCTGCTGAG
GCCC
>SNORD115_28
CGGTCTATGATGACAAACCAATGACATGAAGATAGGTGATGACATAAAATGATGCTCAAAGGATTATGCTGAGGC
CC
>SNORD115_29
GGGTCTATGATGACAAACTAACGTCATGAAGAGAGGTGATGACATAAATATCATGCTGTGGTTGGGGCTGCTGAG
GCCC

>SNORD115_30
GGGTCTATGATGACAAACCAATGTCATGAAGAGAGGTGATGACATAAAAATCATGCTCACTTGGATTACGCTGAG
GCCC
>SNORD115_31
GGGTCTATGATGACAAACCAACGTCATGAAGAGAGGTGATGACATAAATATCATGCTGAGGTTGGGGCTGCTGAG
GCCC
>SNORD115_32
GCGTCTATGATGACAAACCAATGTCATCAAGAGAGGTGCTGACATAAATATCATGCTCAGGGGAGTACGCTGAGG
CCC
>SNORD115_33
GGGTGTATGATGAGAAACCTACGTCCTGAAGAGAGGTGATGACATAAAAATAATGCTCAGGTTCCGGCTGCTGAG
GCCC
>SNORD115_34
AGGTTTATGATGACAAACCAATGTCATGAATAGAGCTGATGACACAAAAATGATGCACAATCAGTTTATGCTGAG
GCCC
>SNORD115_35
AGGTTTATGATGACAAACCAATGTCATGAATAGAGCTGATGACACAAAAATGATGCACAATCAGTTTATGCTGAG
GCCC
>SNORD115_36
AGGTTTATGATGACAAGCCAATGTCATGAATAGAGCTGATGACACAAAAATGATGCAAAATCAGATTATGCTGAG
GCCC
>SNORD115_37
GGGTCTATGATGACAAACCTACTTCCTGAAGAGAGGTGATGACATAAAAATCAATCTCGGGTTGTGGTTGCTGAG
GCCC
>SNORD115_38
GGGTCTATGATGACAAACCAATGTCATGAAGAGAGGTGATGACATAAAAATCATGCTCACTTGGATTACGCTGAG
GCCC
>SNORD115_39
GGGTCTCTGATGACAAACCAATGTCATGAAGAGAAGTGATAACATAATAATCATGCTCAATAGGACTTTGCTGAT
GTGC
>SNORD115_40
AGGAATCTGATGACAATGAATGTCACTAAGAGAGGTGATGACATAAAAATCATACTCAATAGCATGACACTGAGT
CCC
>SNORD115_41
GGGTCTATGATGACAAACCTACGTCCTGAAGAGAGGTGATGATATAAAAATCAATGCTCGGGTTGCGGTTGCTGA
GGCCC
>SNORD115_42
CCGGTTTTGATGACAAACCAATGTCATGAAGACAGGTGATGACATAAAAATCATATTCAATGTGATTATGCTGAA
GCCC
>SNORD115_43
AGGTCTATGATGACAAACCTACGTCATGAAGACAGGTGATGACATAAAAATCATGCTCAGGTTCCAGCTGCTGAG
GCCC
>SNORD115_44
GGGACTATGATGACAAACCAATGTCATGAAGAGAGGTAATGACATAAAAATCATGTTCAGTATGGTAGAGCTGAG
GCAC
>SNORD115_45
GGGTCTATGATGACAAACCAATGTCATGAAGAGAGGTGATGACATAAAAATCATACTCAATAGGACTACGCTGAG
GCCC
>SNORD115_46
GGGTCTATGATGAACAACCAACGTCATGAAGAGAGGTGATGACATAAATATCATGCTGAGGTTGGGGCTGCTGAG
ACCC
>SNORD115_47
GGGTCTATGATGACAAACCAACGTCATTAAGAGAGGTGATGACATAAATATCATGCTGTGGTTGGGGCTGCTGAG
ACCC
>SNORD115_48
GGGTCTATGATGACAAACCAACGTCATTAAGAGAGGTGATGACATAAATATCATGCTGTGGTTGGGGCTGCTGAG
ACCC

```
>SNORD115_49
GGGTCTATGATGACAAACCTATGTCATGAAGAGAGGTGGTGACATAAATATCATGCTCTGGTTCTGGCAGCTGAG
GCCC
>SNORD115_50
GGGTCTATGATGACACACCTACGTCATGAAGAGAGGTGATGACAAAAATATCATGTTCAGGTTCCTGCTGCTGAG
GCCC
>SNORD115_51
GGGTCTATGATGACACACCTACGTCATGAAGAGAGGTGATGACAAAAATATCATGTTCAGGTTCCTGCTGCTGAG
GCCC
>SNORD115_52
GGGTCTATGATGAGAAACTAAAGATATGAAGAGAGGTGATGACAAAATATCATGCTGAGGTTGAGGCTGCTGAGG
CCC
>SNORD115_53
GGGTCTATGATGACAAACTAAAGGCATGAAGAGAGGTGATGACAAAATATCATGCTGAGTTTGAGGCTGCTGAGG
CCC
>SNORD115_54
AGGTCTATGATGACAAACCTACCTCATGAAGAGAGGTGATGACATAAAAATCTTGCTCAGGTTATGGCTGCTGAG
GCCC
>SNORD115_55
GGGTCTATGATGACAAACCAACATCATGAAGAGAGGTGATGACAAAAATATCATGCTGTGGTTGGGGCTGCTGAG
ACCC
>SNORD115_56
GGGTCTATGATGACAAACCAACGTCATGAAGAGAGGTGATGACATACATATCATGCTGGGGTTGTGGCTGCTGAG
ACCC
>SNORD115_57
TGGTCTATGATGACAAACATATGTCATGAAGACAGGTGATGGCATAAAAATCATGCTCAGGTTCCGGATGCTGAG
GCCC
>SNORD115_58
AGGTCTATGATGACAAACCTACGTCCTGAAGAGAGGTGATGACATAAAAATCATTCTCAGGTTCCGGCTGCTGAG
TCCC
>SNORD115_59
GGGTCTATGATGACAAATAAATGTCATGAAGAGAGGTGGTGACATAAAAATCATGCTCAATATGATTACACTGAG
GCCC
>SNORD115_60
GGGTCTATGATGACACCAATGTCATGAAGAGAGGTGATGAAAACAATATCATGTTCAGTATGGTTCTGCTGAGGG
CC
>SNORD115_61
GGATCTATGATGACAAACCTATATCATGAAGAGAGGTGATGACATAAATATCATGCTCTGGTTCCGGCAGCTGAG
GCCC
>SNORD115_62
AGGTCTCTGATGACAAATGAATGTCACTAAGAGAGGTGATGACATAAAAATCATACTATGTAACATGACACTGAG
TCCC
>SNORD115_63
GGGTCTATGATGACAAACCAACGTCATGAAGATAGGTGATGACATAAATATAATTCTGTGGTTTGTGCTGCTGAG
ACCC
>SNORD115_64
AGGTCTATGATGAAAAACCTACGTCATGAAGAGAGGTGATGACATAAATATCATGCTCTGGTTCTGGCTACTGAG
GCCA
>SNORD115_65
GGGTCTATGATGACAAACCTACGTCATCAATAGAGGTGATGACATAAATATTATGCTCTGGTTCCAGCTGCTGAG
GCCT
>SNORD115_66
CGGTCTATGATGACAAACCAACGTCATGAAGAGAGGTGATGACATAAATATCATGCTGAGGTAGGGGCTGCTGAG
ACCC
>SNORD115_67
GGGTCTATGATGACAAACCAATGTCATAAAAAGAGGTGATGACATAAAAATCATGCTCACTTGGATTACGCTGAG
GCCC
```

>SNORD115_68
GGATCTATGATGAAAAACCAACGTCATGAAGAGAGGTAATGACATAAATATCATGCTGAGGTTGGGGCTGCTGAG
GCCC
>SNORD115_69
TGGTCTATGATGACAAACCTACGTCCTGAATAGAGGTGATGACATAGAAATCATGGTCATGTTCCAGCTGCTGAG
GCCC
>SNORD115_70
GGATCTATGATGACAAACCAACGTCATGAAGAGAGGTAATGACATAAATATCATGCTGAGGTTGGGGCTGCTGAG
GCCC
>SNORD115_71
GGGTCTATGATGACAAACCAAAGGCATGAAGAGAGGTGATGACAAAATATCATGCAGAGTTTGAGGCTGCTGAGG
CCC
>SNORD115_72
GGGTCTATGATGACAAACCAAAGGCATGAAGAGAGGTGATGACAAAATATCATGCAGAGTTTGAGGCTGCTGAGG
CCC
>SNORD115_73
GGGTCTATGATGACACACCTACCTCATGAAGAGAGGTGATGACAAAAATATCATGTTCATGTTCCTGCTGCTGAG
GCCT
>SNORD115_74
TGGTCTATGATGACAAACCAACGACATGAAGAGAGGTGATGACATAAATATCATGCTGTGGTTGGGGCTGCTGAG
ACCC
>SNORD115_75
TGGTCTATGATGACAAACCAACGACATGAAGAGAGGTGATGACATAAATATCATGCTGTGGTTGGGGCTGCTGAG
ACCC
>SNORD115_76
CGGTCTATGATGACAAACCAACGTCATGAAGAGAGGTGATGACATAAATATCATGCTGAGGTTGGGGCTGCTGAG
ACTC
>SNORD115_77
GGGTCTATGATGACAAACCAATGTCACGAAGAGAGGTGATGACATAAAAATCATGCTCAATAGGACTACTCTGAG
GCCC
>SNORD115_78
GGGTCTATGATGACAAACCAAAGGCATGAAGAGAGGTGATGACAAAATATCATGCAGAGTTTGCGGCTGCTGAGG
CCC
>SNORD115_79
GTACCTATGATGACATACTAATTTCCCAAAAAGGTTTGATGACATAGAAATCATGCTGAATAGGATTACACTAAA
GGTC
>SNORD115_80
GGTTCTATGATGAGAAACCTACGTCCTGAAGAGAGCAGATGTCATAAAAGTCATTCTCAGGTTCCGGTTGCTGAG
GCCC
>SNORD115_81
TGGACTATGATGACAAACCAACATCATGAAGAGAGATGATGACATAAAAATCATGCTGTGGTTGGGGCTGCTGAG
ACTT
>SNORD115_82
GGGTCTATGATGACAAACCAAAGGCATGAAGAGAGGTGATGACAAAATATCATGCAGAGGTTGTGGCTGCTGAGG
CCC
>SNORD115_83
GGGTCTATGATGACAAACCAAAGGCATGAAGAGAGGTGATGACAAAATATCATGCAGAGGTTGTGGCTGCTGAGG
CCC
>SNORD115_84
GGGTCTATGATGACAAACCAAAGGCATGAAGAGAGGTGATGACAAAATATCATGCAGAGGTTGTGGCTGCTGAGG
CCC
>SNORD115_85
GGGTCTATGATGACACACCAATGTCATGAAGAGAGGTGATGACATAAAAATCAAGCTCAATAGGATTACTCTGAG
GCCC
>SNORD115_86
GGGTCTATGATGACAAACCAAAGGCATGAAGAGAGGTGATGACAAAATATCATGCAGAGGTTGTGGCTGCTGAGG
CCC

>SNORD115_87
GGGTCTATGATGACAAACCAAAGGCATGAAGAGAGGTGATGACAAAATATCATGCAGAGGTTGCGGCTGCTGAGG
CCC
>SNORD115_88
GGGTCCATGATGACATGACTACGTCATGAAGAGAGGTGATGACAAAAATATCATGCTCTGGTTCCAGCTGCTGAG
GCCA
>SNORD115_89
GGGTCCATGATGACATACCTACATCATGAAGAGAGGTGATGACAAAAATATCATGCTCTGGTTCCAGCTGCTGAG
GCCA
>SNORD115_90
TGGTCTATGATGACAAACCAAAGGCATGAAGAGAGGTGATGACAAAATATCATGCAGAGTGTGAGGCTGCTGAGG
CCA
>SNORD115_91
GGGTCTATGATGACAAACCAAGGCATGAAGAGAGGTGATGACAAAATATCATGCAGAGGTTGCGGCTGCTGAGGC
CC
>SNORD115_92
GGGTCTATGATGACAAAACTATGTCATGAAGAGAGGAAATGACATAAAAATCATGCTCAGTAGGATTACACTGAG
GCCC
>SNORD115_93
GGGTCTATGATGAAAAACCAAAGGCATGAAGTGAGGTGATGACAAAATATCATGCAGAGGTTGTGGCTGCTGAGG
CCC
>SNORD115_94
GGGTCTATGATGACAAACCAATGTCACAAAAGAGGTGATGACATAAAAATCATGCTCAATAGGATTACACTGAGG
CCC
>SNORD115_95
TGGTCTATGATGACAAACCAAAGGCACGAAGAGAGGTGATGACAAAATATCATGCAGTGTGTGAGGCTGCTGAGG
CCA
>SNORD115_96
GGGTCTATGATGACAAAACAATGTCAAGAAGAGAGGTGATGACATAATAATCATGCTCAATAGGATTACTCTGAG
GCCC
>SNORD115_97
GGATCTATGATGACAAACCAATGTTATGAAGAGAGGTGATGACATAAAAATCATGCTCAATGGGATTACGCTGAT
GCCC
>SNORD115_98
GGGTCTATGATGAAAAACCAAAGTCGTGAAGAGAGGTGATGACATAAAAATCATGCTCACTGGGGTTCCGCTGAG
GCCC
>SNORD115_99
AGGTCTATGATGACAAACCAAAGTCATGAAGAGTGGTGATGACAAAAAAATCATGCTCAATAGGATTATGCTGAG
ACCC
>SNORD115_100
GGGTCTATGATGACTAACCAATGTCATGAAGAGAGATGGTGACAGAAAATTCATGATCAATAGGATTATGCTGAG
GCCC
>SNORD115_101
GGGTATGTGATGACAAACCAAAGTTACAAAGAGAGTTGATGACATAAAATTCATGCTCAATAGGATTATGCTGAG
GCCC
>SNORD115_102
GGGTCTATGATGAGAAACCTACGTCCTGAAGAGAGGTGATGACATAAAAATCATGCTCAGGTTCCGGCCGCTGAG
GCCC
>SNORD115_103
GGGTATATGATGACAAACTAGTGACATGAATATAGGTGATGACATAAAAATCATGTTCAATAGGACTATGCTGAG
TCCC
>SNORD115_104
GGGTCTATGATGAGAAACCTACGTCCTGAAGAGAGGTGATGACATAAAAATCATGCTCAGTCTCGGTCTGCTGAG
GCCC
>SNORD115_105
GGGTCTATGATGAGAAACCTACGTCCTGAAGAGAGGTGATGACATAAAAATCATGCTCAGTCTCGGTCTGCTGAG
GCCC

>SNORD115_106
GGGTCTATGATGACAAACCAATCTCATGAAGAGAGGTGATGACATAAAAATCATGCTCACTTGGATTACGCTGAG
GCCC
>SNORD115_107
AGGTTTATGATGACAAACCAATGTCATGAATAGAGCTGATGACACAAAAATGATGCACAATCAGTTTATGCTGAG
GCCC
>SNORD115_108
AGGAATCTGATGAAAAATGAATGTCTCTTAAAGAGGTGATGACATAAAAATCATATTCAATAACATGACACTGAG
TCCC
>SNORD115_109
GGGTCTATGATGACAAACCAAAGTCATGAAGAGAGGTGATGACATAAATATCATGCTGAGGTTCTGGCTGCTGAG
GCCC
>SNORD115_110
GGGTCTATGATGACAAACTAATGTCATGAAGAGAGGTGATGACAACAATATCATGTTCAGTATGGTTCTGCTGAG
GCCC
>SNORD115_111
GGGTCTATGATGACAAACCAATGTCATGAAGAGAGGTGATGACAACAATATCATGTTCAGTATGGTTCTGCTGAG
GCCC
>SNORD115_112
GGGTCTATGATGACAAACCAATGTCATGAAGAGAGGTGATGACAACAATATCATGTTCAGTATGGTTCTGCTGAG
GCCC
>SNORD115_113
GGGTCTATGATGACAAACCAATGTCATGAAGAGAGGTGATGACAACAATATCATGTTCAGTATGGTTCTGCTGAG
GCCC
>SNORD115_114
GGGTCTATGATGACAAACCAATGTCATGAAGAGAGGTGATGACATAAAAATCATGCTCACTTGGATTACGCTGAG
GCCC
>SNORD115_115
AGGAATCTGATGAAAAATGAATGTCACTTAAAGAGGTGATGACATAAAAATCATATTCAATAACATGACACTGAG
TCCC
>SNORD115_116
GGGTCTATGATGACAAACCAACGTCATGAAGAGAGGTGATGACATAAATATCATGCTGAGGTTAGTGATGCTGGG
GCCC
>SNORD115_117
AGAACTATGATGACAAACCAATGTCATGAAGAGAGTTGATGACATAAAAATCATGCACAATAGGTCTATGTTGAG
GCAC
>SNORD115_118
TGGTCTATGATGACAAACCAATGTCATGAAGAGAGGTGATGATATACAACCACATGCTCACTTGGATTACGCTTG
AGGCCC
>SNORD115_119
AGGAATCTGATGACAATGAATGTCACTAAGAGAGGTGATGACATAAAAATCATACTCAATAACATGACACAGAGT
CAC
>SNORD115_120
GGGTCTATGATGACAAACCAATGTCATGAACAGAGGTTATGACATAAAAATCATGCTCAATAGGATTATGCTGTG
GCCC
>SNORD115_121
AGAACTATGATGACAAACCAATGTCATGAAGAGAGTTGATGACATAAAAACCATGCACACGAGGCCTATGTTGAG
GCAC
>SNORD115_122
AAGTATATGATGACATACCAATGTCCCAAAAAGAGCTGATGACATAGTAATTATGGTCAATTAGATAACGTTGAA
GCTC
>SNORD115_123
GGGTCGATTTTGACAAACCAATGTCAGGAAGAGTGGTGATGACATAAAAATCATGCTCAATAGGATTACGATAAG
GCCC
>SNORD115_124
AGGTCTATGATGACAAACCAATGTCATGCAGAGAGATGATGACATAAAAATCATTCTCAATAGGATTGCGATGAG
GTCC

>SNORD115_125
GGGTCTATGATGACAAACCAATGTCATGAAGAGAGGTGATTACATAAAAATCATGCTCAATGGGATTACGATTAT
TCCC
>SNORD115_126
AGGTTTATGATGACAAACCAATGTCATGAAGAGAGATGATTACATAAAAACGATGCTCAATAAGATTATGCGGAG
GCCC
>SNORD115_127
GGGTCTATGATGACATACCAATGTCATGCAGGTCGATGAAGACGTAAAAATCATTTTCAATAGGATTACGATGAG
GCCC
>SNORD115_128
GGGTGTGTGATGACAAACCAATGTTATGAAGAAAGGTGATGACAAAAAAATCATGCTCAATAGGATTATGCTAAG
GCCC
>SNORD115_129
AGAACTATGATGAAAAACCAATGTCATGAAGTGAGTTGATGACATAAAAATCATGCACACAAGGCCTATGTTGAG
GCAC
>SNORD115_130
CGGTCTATGATGACAAACCAACGTCATGAAGAGTGGTGATGACATAAATATCATGCTGAGGTTGGGGCTGCTGGA
GACC
>SNORD115_131
AGAACTATGATGAAAAACCAATGTCATGAAGTGAGTTGATGACATAAAAATCATGCACACAAGGCCTATGTTGAG
GCAC
>SNORD115_132
GGGTCTATGATGACAAACCAATGTCATGAAGAGAGGTTATGACATAACAATCATGCTCAATAGGATTATACTCTG
GCCC
>SNORD115_133
GGGTCTATAATGACAAATCAATGTCATGAAGAGAGGTGATGACATACAAATCATGCTCAATAGGATTATGCTCAG
GCCC
>SNORD115_134
AGGAATATGGTGATAAATGAATGTCTCTAAGAGAGTTGAAGACCCAAAAATCATACTCAATAACATAACACTGAA
TCCC
>SNORD115_135
AGGTCTGTAGTGACAAACGAATGATCTGAAGACAAGTGATGATATAAAAACCATGATTAAAAGGTTATGCTGAAG
CCC
>SNORD115_136
GGGTCTATGGGGACAAACCAACGTCATGAAGAGAGGTGATGACAAAGATATCATGCTCAGGATGGGGCTGCTGAG
TCCC
>SNORD115_137
GGATATATGAGGACAAACCAATGTTATGAACAGAGGTGATGACATAAAAATCATGCTCAATGGGATTACGCTGAT
GCCC
>SNORD115_138
GGGTCTATAATGACAAATCAATGTCATGAAGAGAGGTGAAGACATACAAATCATGCTCAATAGGATTATGCTCAG
GCCC
>SNORD115_139
AGGTTGATTTTGACAAACCAATATCATGAAGAGAGGTGATGACATAAAAATCATGCTCAATAGGATTATGCTGAG
GCCA
>SNORD115_140
GGGTCTATAATGACAAACCAATGTCATGAAGAGAGGTGATGACATCAATATCATGTTCAGTATGGTTCTGCTGAG
GCCC
>SNORD115_141
GGGTCTATAATGACAAATCAATGTCATGAAGAGAGGTGATGACATACAAATCATGCTCAATAGGATTATGCTCAG
GCCC
>SNORD115_142
GGATCTATGAGGACAAACCAATGAAAGGAAGAGAGGTGATGACATAAAAATCATGCTCAATGGGATTACGCTGAT
GCCC
>SNORD115_143
GGGTCTAGGATGGCAAACCAATGTCATGAAGAGAGATGATAACATCAAATCATGCTCAATAGGATTATGCTGAGG
CCC

```
>SNORD115_144
GGGTCAATTTTGACAAACCAATGTCAGGAAGAGAGGTGATGACATAAAAATCATGCTCAATAGGATTACGCAGAG
GTCC
>SNORD115_145
GGGTTGATTTTGACAAACCAAAGTCACGAAGAGAGGTGATGACATAAAAATCATGCTCAATATGATTACGCTGAG
GCCC
>SNORD115_146
GGGTCTATGATGGCAAACCAATGTCATGAAGAGAGATGATAACATCAAATCATGCTCAATAGGATTATGCTGAGG
CCC
>SNORD115_147
GGGTCTATGATGGCAAACCAATGTCATGAAGAGAGATGATAACATCAAATCATGCTCAATAGGATTATGCTGAGG
CCC
>SNORD115_148
GGGTATATAATGACAAATCAATGTCATGAAGAGAGGTGATGACATACAAATCATGCTCAATAATATTATGCTCAG
GCCC
>SNORD115_149
GGGTCTATGATAAAAAACCAATGTCATGAGAAGAGCTGATGACATAAAAATCATGCTCAATAGGATTATGCTGAG
ACAC
>SNORD115_150
AGGAATATGGTGATAAATGAATGTCTCTAAAAGAGTTGATGACACAAAAATCATACTCAATAACATAACAATTAA
TCCC
>SNORD115_151
AGGTCTGTAGTGACAAACGAATGATCTGAAGAGAAGTGATGATATAAAAACCATGATTAAAAGGTTATGCTGAAG
CCC
>SNORD115_152
GGGTCTATGATGACATAACAATGTCATGCAGATCGATAAACACATAAAAATCATTTTCAATAGGATTATGATGAG
ACCC
>SNORD115_153
GGGTCTAGGATGGCAAACCAATGTCATGAAGAGAGGTGATGACATCAAATCATGCTCAATAGGATTATGCTGAGG
CCC
>SNORD115_154
GGGTCTATAATGACAAATCAATGTCATGAAAAGAAATGATGATATACAAATCATGCTCAATAGGATTACGCTCAG
GCCC
>SNORD115_155
GGATGTATAATGACAAATCAATGTCATGAAGAGAGGTGATGACATACAAATCATGCTCAATTGTATTATGTTCAG
GCCC
>SNORD115_156
GGGTCTATGATCACAAACCAATGTCATGAAGAGAGGTGATAACATAAAAATAATGCTTAATCGGATTATGCTGAG
GGCC
>SNORD115_157
AGGTCTATGATGGCAAACCAATGTCATGAAGAGAGGTAATGGCATATAAATCATGCTCAGTAGGGTTACGCTGAG
GCCC
>SNORD115_158
GGGTCTGTTATGACAAATCAATGTCATGAAGAAAGGTGAAGACAAAATAATCATGCTCAATAGGATTACACTAAG
GCCC
>SNORD115_159
GGGTCTGTGATGGCAAACCAATGTCATGAAGAGAGGTGATGACATCAAAATCATGATCAATAGGATTACACTGAG
GCCC
>SNORD115_160
AGGTCTATGATGGCAAGCCAATGTCACAAAAAGAGGTGATGACATGAAATTCATGCTCAAGAGGATTATGCTGAG
GCCC
>SNORD115_161
GGGTATATTATGACAAACCAATGTAATGAAGAGAGGGGATGACATAAAAATCATGTTCACTAGGGTTCTGCTGAG
ACCC
>SNORD115_162
GGGTCTATGACGACAAACCAATGTCATGAAGAGAGGTGATGACAACAATATCATGTTCAGTATGGTTCTGCTGAG
GCCC
```

>SNORD115_163
GGGTATGTGACGACAAACCAAAGTCATGAAGAGAGTTGACGATATAAAATCCATGCTCAATAGGGTTATGCTGAG
GCCC
>SNORD115_164
GAGTCTATGATGACAAACCAATGCCATGCAGAGTGATGATGACATAAAAATCATTCTCAATAGGATTATGATGAG
GCCG
>SNORD115_165
GAGTCTATGATGACAAACCAATATCATGAAGAGAGGTGATGTCATAAATATCATGTTCAGTATGGTTCTGCTGTG
GCCC
>SNORD115_166
GGGTCTATGATGACAAACCATTGTCATGAAAGAGGTGATGACGGAAAAATCATGGTCAATAGGATTACAATGAGC
CA
>SNORD115_167
GGGTCTGTAGTGACAAACGAATGATATGAAGAGAAGAGATGATATAAAAACCATGATTAAAAGGGTTATGCTGAA
GCCC
>SNORD115_168
GGGTCTATGAGGTGAAACCTACGTCCTGAAGACAGGTGATGACATAAAAATCATGCTCAGGTTCCGGCTGCTGAG
GCCC
>SNORD115_169
GAGCCTATCATGACAAACCAATATGTCAGGAATAGAGGTGATGACATAACTATTATGGTCAATAGGATTACTCTG
AGGCCC
>SNORD115_170
GGGTCTAGGAAGCCAAAACAATGTCAGGAAGAGAGGTGATGACATATAAATCATGCTCAATATAATTATTCTGAT
GTCC
>SNORD115_171
GGGACAATGACGACAAACCTACATCATCAATAGAGGTGATGACATAAATATTATGCTCTGGTTCCGGCTGCTGAG
GCCC
>SNORD115_172
GGGTCTATGAGGACAAACCAACGTCATGAAGAGAGGTGATGACAAAAAATATCATGCTCAGGATGGGGCTGCTGA
GTCCC
>SNORD115_173
GGGTCTATAATGACAAACCAACGTCATTAAGAGAGTTGATGACATAAATATCATGCTGTGGTTGGGGCTGCAGAG
ACCC
>SNORD115_174
GGGTCTTTAGTGGCAAAACAATGTTAAAAAGAGAAGTGATGATATAAAAACCATGATTTAAAGGATTATACTGAA
GCCC
>SNORD115_175
GGGTCTATAATGACAAACCAACATCATCAAGAGAGGTGATGACATAAATATCATGCTGTGGTTGGGGCTGCTGAG
ACCC
>SNORD115_176
GGGTCTATGGGGACAAACCAACGTCATGAAGAGAGGTGATGACATAAATATCATGCTCAGGATGGGGCTGCTGAG
TCCC
>SNORD115_177
GGGTCTGTAGTGAAAAACGAATGATATGAAGAGAAGTGATGATATAAAAAAGCATTATTAAAAGGGTTATGCTGA
AGCCC
>SNORD115_178
GGGTCTATGATTACAAAGCAATGCCATGAAGAGAGGTGATGACATAAAAATCATGCTCAATAGGATTACGCTGAG
GCCC
>SNORD115_179
GGGTCTATGTTGACAAACCAATATCATGAAGAGAGGTGACGACATAAATATCATGCTGTGGTTGGGGCTGCTGAG
ACCC
>SNORD115_180
GGGTCTATGAAGACAAACCTATGTCATGAAGAGAGGTGGTGACATAAATATCATGCTCTGGTTCCGGCAGCTGAG
GCCC
>SNORD115_181
GGGTCTATGTTGACAAACCAACATCATGAAGAGAGGTGACGACATAAATATCATGCTGTGGTTGGCGCTGCTGAG
ACCC

>SNORD115_182
GGGTCTATGTTGACAAACCAACATCATGAAGAGAGGTGACGACATAAATATCATGCTGTGGTTGGCGCTGCTGAG
ACCC
>SNORD115_183
GGGTCAATAATGACATATCTACGTCATGAAGAGAGGTGATGACAAAAATATCATGCTCTGGTTCCAGCTGCTGAG
GCCA
>SNORD115_184
GGGTCTATGGGGACAAACCAACGTCATGAAGAGAGGTGATGACATAAATATCATGCTCAGGATGGGGCTTCTGAG
TCCC
>SNORD115_185
TTGTCTAAGATGACGTACTAATTTTCAAAAAGAGCTGATGACATAGAAATCATGATCAATAGGATTACGCTGAAG
CTC
>SNORD115_186
GGGTCTATGATGGCAAGCCAATGTCATGAAGAGAGGTGATGACATAAAAATCATGCTCAATAGGATTACTCTGAG
GCCC
>SNORD115_187
AGGAATATGGTTATAAACGAATGTCTCTAAGAGAGTTGATGACACAAAAATCATACTCAATAACATTACACTGAA
TCCC
>SNORD115_188
GGGTCTATGTTGACAAACCAACATAATGAAGAGAGGTGACAACATAAATATCATGCTGTGGTTGGGGCTGCTGAG
ACCC
>SNORD115_189
GGGAATATGGTGACAAATGAATGTCTCTAATATAGGTGATGACACAAAAATCATACTCAATAACATTAAACTGAT
TCCC
>SNORD115_190
AGGTCTATTATGACAAACCTACCTCATGAAGAGAGGTGATGACATAAAACTCGTGCTCAGGTTCTGGCTGCTGAG
GCCC
>SNORD115_191
GGGTCTATAGTGACAAACAAATGCTACAAATGGAAATGATGACATAAAAATCATGATTAAAAGGTTTATGCTGAA
GCAA
>SNORD115_192
GGGTCTATGATGTCAGACCAATGTCACGGATAAAAGGTGATGACATAAAAATCATGTTCAATATGATTACGGTAA
ATGCCA
>SNORD115_193
GGGAATATGGTAACAATTGAATGACTCTAAGTGAGGTGATGACAGAAAAAAATCATAGTCAATAACATTACACCG
AGTCCT
>SNORD115_194
GAGTCTCTGATGACATACTGATTTCCCAAAAAGATCTGATGACATAGAAATCATGCTCAATATGATTACGCTTAA
GCTC
>SNORD115_195
GGGTCTATGTTGACAAACCAACATCATGAAGAGAGGTGACGACATAAATATCATGCTGTGGTTGGGGCTGCTCAG
ACCC
>SNORD115_196
AGGAATCTGATTACGAATGAATGTCACTAAGAGAGGTGATGACATAAAAATCATACTCAATAACATGACACTGAA
TCCC
>SNORD115_197
GGTTCTATGATTATACACTAGTGTCCTGAAAAGGGCTGATGACATAGAAATCATGTTCAATATGGTTACCCTAAA
GCTC
>SNORD115_198
AGGAATATGGTTATAAATGAATGTCTCTAAGAGAGTTGATGACACAAAAATTCATACTCAATAACATTACACTGA
ATCCC
>SNORD115_199
GGGTCCATGATGGCATACCTACGTCATGAAGAGAGGTGATGACAAAAATATCATGCTCTGGTTCCAGCTGCTGAG
GCCA
>SNORD115_200
GGGTCTACGATGACAAACCAAAGGCATGAAGAGAGGTGATGACAAAATATCATGCAGAGGTTGTGGCTGCTGAGG
CCC

### Cluster of SNORD115-like sequences in *S. araneus* (European shrew)

>SNORD115-like_1
GGGTCAATGATGAGAAAGCCAGGCATTGCTTCTGAAGGAGCTATGCTGATCATAAACGGTTAACTCAAGGGTGTC
ACACTGAGGCCC
>SNORD115-like_2
GGGTCAATGATGAGAAAGCCAGGCATTGCTTCTGAAGGAGCTATGCTGATCATAAACGGTTAACTCAAGGGTGTC
ACACTGACGCCC
>SNORD115-like_3
GGGTCAATGATGAGAAAGCCAGGCATTGCTTCTGAAGGAGCTATGCTGATTATAAAGGGTTAACTCAAGGGTGTC
ACACTGAGGCCC
>SNORD115-like_4
GGGTCAATGATGAGAAACCCAGGCATTGCTTCTGAAGGAGCTATGCTGATCATAAAGGGTTAGCTCAAGGGTGTC
ACACTGAGGCCC
>SNORD115-like_5
GGGTCAATGATGAGAAACCCAGGCATTGCTTCTGAAGGAGCTATGCTGATCATAAAGGGTTAGCTCAAGGGTGTC
ACACTGAGGCCC
>SNORD115-like_6
GGGTCAATGATGAGAAACCCAGGCATTGCTTCTGAAGGAGCTATGCTGATCATAAAGGGTTAGCTCAAGGGTGTC
ACACTGAGGCCC
>SNORD115-like_7
GGGTCAATGATGAGAAACCCAGGCATTGCTTCTGAAGGAGCTATGCTGATCATAAAGGGTTAGCTCAAGGGTGTC
ACACTGAGGCCC
>SNORD115-like_8
GGGTCAATGATGAGAAACCCAGGCATTGCTTCTGAAGGAGCTATGCTGATCATAAAGGGTTAGCTCAAGGGTGTC
ACACTGAGGCCC
>SNORD115-like_9
GGGTCAATGATGAGAAACCCAGGCATTGCTTCTGAAGGAGCTATGCTGATCATAAAGGGTTAGCTCAAGGGTGTC
ACACTGAGGCCC
>SNORD115-like_10
GGGTCAATGATGAGAAACCCAGGCATTGCTTCTGAAGGAGCTATGCTGATCATAAAGGGTTAGCTCAAGGGTGTC
ACACTGAGGCCC
>SNORD115-like_11
GGGTCAATGATGAGAAACCCAGGCATTGCTTCTGAAGGAGCTATGCTGATCATAAAGGGTTAGCTCAAGGGTGTC
ACACTGAGGCCC
>SNORD115-like_12
GGGTCAATGATGAGAAACCCAGGCATTGCTTCTGAAGGAGCTATGCTGATCATAAAGGGTTAGCTCAAGGGTGTC
ACACTGAGGCCC
>SNORD115-like_13
GGGTCAATGATGAGAAACCCAGGCATTGCTTCTGAAGGAGCTATGCTGATCATAAAGGGTTAGCTCAAGGGTGTC
ACACTGAGGCCC
>SNORD115-like_14
GGGTCAATGATGAGAAACCCAGGCATTGCTTCTGAAGGAGCTATGCTGATCATAAAGGGTTAGCTCAAGGGTGTC
ACACTGAGGCCC
>SNORD115-like_15
GGGTCAATGATGAGAAACCCAGGCATTGCTTCTGAAGGAGCTATGCTGATCATAAAGGGTTAGCTCAAGGGTGTC
ACACTGAGGCCC
>SNORD115-like_16
GGGTCAATGATGAGAAACCCAGGCATTGCTTCTGAAGGAGCTATGCTGATCATAAAGGGTTAGCTCAAGGGTGTC
ACACTGAGGCCC
>SNORD115-like_17

GGGTCAATGATGAGAAACCCAGGCATTGCTTCTGAAGGAGCTATGCTGATCATAAAGGGTTAGCTCAAGGGTGTC
ACACTGAGGCCC
>SNORD115-like_18
GGGTCAATGATGAGAAACCCAGGCATTGCTTCTGAAGGAGCTATGCTGATCATAAAGGGTTAGCTCAAGGGTGTC
ACACTGAGGCCC
>SNORD115-like_19
GGGTCAATGATGAGAAACCCAGGCATTGCTTCTGAAGGAGCTATGCTGATCATAAAGGGTTAGCTCAAGGGTGTC
ACACTGAGGCCC
>SNORD115-like_20
GGGTCAATGATGAGAAACCCAGGCATTGCTTCTGAAGGAGCTATGCTGATCATAAAGGGTTAGCTCAAGGGTGTC
ACACTGAGGCCC
>SNORD115-like_21
GGGTCAATGATGAGAAACCCAGGCATTGCTTCTGAAGGAGCTATGCTGATCATAAAGGGTTAGCTCAAGGGTGTC
ACACTGAGGCCC
>SNORD115-like_22
GGGTCAATGATGAGAAACCCAGGCATTGCTTCTGAAGGAGCTATGCTGATCATAAAGGGTTAGCTCAAGGGTGTC
ACACTGAGGCCC
>SNORD115-like_23
GGGTCAATGATGAGAAACCCAGGCATTGCTTCTGAAGGAGCTATGCTGATCATAAAGGGTTAGCTCAAGGGTGTC
ACACTGAGGCCC
>SNORD115-like_24
GGGTCAATGATGAGAAACCCAGGCATTGCTTCTGAAGGAGCTATGCTGATCATAAAGGGTTAGCTCAAGG
GTGTCACACTGAGGCCC
>SNORD115-like_25
GGGTCAATGATGAGAAACCCAGGCATTGCTTCTGAAGGAGCTATGCTGATCATAAAGGGTTAGCTCAAGGGTGTC
ACACTGAGGCCC
>SNORD115-like_26
GGATCAATGATGAGAAACCCAGGCATTGCTTCTGAAGGAGCTATGCTGATCATAAAGGGTTAACTCAAGGGTGTC
ACACTGAGGCCC
>SNORD115-like_27
GGGTCAATGATGAGAAACCCAGGCATTGCTTCTGAAGGAGCTATGCTGATCATAAAGGGTTAGCTCAAGGGTGTC
ACACTGAGGCCC
>SNORD115-like_28
GGGTCAATGATGAGAAACCCAGGCATTGCTTCTGAAGGAGCTATGCTGATCATAAAGGGTTAGCTCAAGGGTGTC
ACACTGAGGCCC
>SNORD115-like_29
GGGTCAATGATGAGAAACCCAGGCATTGCTTCTGAAGGAGCTATGCTGATCATAAAGGGTTAGCTCAAGGGTGTC
ACACTGAGGCCC
>SNORD115-like_30
GGGTCAATGATGAGAAACCCAGGCATTGCTTCTGAAGGAGCTATGCTGATCATAAAGGGTTAGCTCAAGGGTGTC
ACACTGAGGCCC
>SNORD115-like_31
GGGTCAATGATGAGAAACCCAGGCATTGCTTCTGAAGGAGCTATGCTGATCATAAAGGGTTAGCTCAAGGGTGTC
ACACTGAGGCCC
>SNORD115-like_32
GGGTCAATGATGAGAAACCCAGGCATTGCTTCTGAAGGAGCTATGCTGATCATAAAGGGTTAGCTCAAGGGTGTC
ACACTGAGGCCC
>SNORD115-like_33
GGGTCAATGATGAGAAACCCAGGCATTGCTTCTGAAGGAGCTATGCTGATCATAAAGGGTTAGCTCAAGGGTGTC
ACACTGAGGCCC
>SNORD115-like_34
GGGTCAATGATGAGAAACCCAGGCATTGCTTCTGAAGGAGCTATGCTGATCATAAAGGGTTAGCTCAAGGGTGTC
ACACTGAGGCCC
>SNORD115-like_35
GGGTCAATGATGAGAAACCCAGGCATTGCTTCTGAAGGAGCTATGCTGATCATAAAGGGTTAGCTCAAGGGTGTC
ACACTGAGGCCC
>SNORD115-like_36
GGGTCAATGATGAGAAACCCAGGCATTGCTTCTGAAGGAGCTATGCTGATCATAAAGGGTTAGCTCAAGGGTGTC
ACACTGAGGCCC
>SNORD115-like_37

GGGTCAATGATGAGAAACCCAGGCATTGCTTCTGAAGGAGCTATGCTGATCATAAAGGGTTAGCTCAAGGGTGTC
ACACTGAGGCCC
>SNORD115-like_38
GGGTCAATGATGAGAAACCCAGGCATTGCTTCTGAAGGAGCTATGCTGATCATAAAGGGTTAGCTCAAGGGTGTC
ACACTGAGGCCC
>SNORD115-like_39
GGGTCAATGATGAGAAACCCAGGCATTGCTTCTGAAGGAGCTATGCTGATCATAAAGGGTTAGCTCAAGGGTGTC
ACACTGAGGCCC
>SNORD115-like_40
GGGTCAATGATGAGAAACCCAGGCATTGCTTCTGAAGGAGCTATGCTGATCATAAAGGGTTAGCTCAAGGGTGTC
ACACTGAGGCCC
>SNORD115-like_41
GGGTCAATGATGAGAAACCCAGGCATTGCTTCTGAAGGAGCTATGCTGATCATAAAGGGTTAGCTCAAGGGTGTC
ACACTGAGGCCC
>SNORD115-like_42
GGGTCAATGATGAGAAACCCAGGCATTGCTTCTGAAGGAGCTATGCTGATCATAAAGGGTTAGCTCAAGGGTGTC
ACACTGAGGCCC
>SNORD115-like_43
GGGTCAATGATGAGAAACCCAGGCATTGCTTCTGAAGGAGCTATGCTGATCAAAATGGTTAACTCAAGGGTGTCA
CACTGAGGCCC
>SNORD115-like_44
GGGTCAATGATGAGAAACCCAGGCATTGCTTCTGAAGGAGCTATGCTGATCAAAATGGTTAACTCAAGGGTGTCA
CACTGAGGCCC
>SNORD115-like_45
GGGTCAATGATGAGAAACCCAGGCATTGCTTCTGAAGGAGCTATGCTGATCATAAAGGGTTAGCTCAAGGGTGTC
ACCCTGAGGCCC
>SNORD115-like_46
GGGTCTATGATGAGAAACCCAGGCATTGCTTCTGAAGGAGCTATGCTGATCATAAAGGGTTAGCTCAAGGGTGTC
ACACTGAGGCCC
>SNORD115-like_47
GGGTCAATGATGAGAAACCCAGGCATTGCTTCTGAAGGAGCTATGCTGATCACAAAGGGTTAGCTCAAGGGTGTC
ACACTGAGGCCC
>SNORD115-like_48
GGGTCAATGATGAAAAACCCAGGCATTGCTTCTGAAGGAGCTATGCTGATCATAAAGGGTTAGCTCAAGGGTGTC
ACACTGAGGCCC
>SNORD115-like_49
GGGTCAATGATGAGAAACCCAGGCATTGCTTCTGAAGGAGCTATGCTGATCAACATGGTAAACTCAAGGGTGTCA
CACTGAGGCCC
>SNORD115-like_50
GGGTCAATGATAAGAAAGCCAGACATTGCTTCTGAAGGAGCTATGCTGATCAAAATGGTTAACTCAAGGGTGTCA
CACTGGGGCCC
>SNORD115-like_51
GGGTCAATGATGAGAATCCCAGGCATTGCTTCTGAAGGAGCTATGCTGATCATAACGGGTTAGCTCATGGGTGTC
ACACTGAGGCCC
>SNORD115-like_52
GGGTCAATGATGAGAACCCCAGGCATCGCTTCTGAAGGAGCTATGCTGATCATAAGGGTCAACTCAAGGGTGTCA
CACTGAGGCCC
>SNORD115-like_53
GGGTCAATGATGAGAACCCCAGGCATCGCTTCTGAAGGAGCTATGCTGATCATAAGGGTCAACTCAAGGGTGTCA
CACTGAGGCCC
>SNORD115-like_54
GGATCACTGATGAGAAACCCAGGCATGGCTTCTGAAGGAGCTATGCTGATCATAATGGTTAACTCAAGGGTGTCA
CACTGAGGCCC
>SNORD115-like_55
GGGTCAATGATGAGAACCCCAGGCATCGCTTCTGAAGGAGCTGTGCTGATCATAAGGGTCAACTCAAGGGTGTCA
CACTGAGGCCC
>SNORD115-like_56
GGGTCAATGATGAGAAACCCAGGCGGTGCTTCTGAAGGAGCTATGCTGATTCAAAGGGTTAACTCAAGGGTGTCA
CACTGAGGCCC
>SNORD115-like_57

GGGTCAATGATGAGAAACCATGCATTGCTTCTGAAGGAGCTATGCTGTTCTTATAGGTTAACTCAAGGGTGTCAC
ATTGATGCCC
>SNORD115-like_58
GGGTCAATGATGAGAACCCCAGGCATCACTTCTGAAGGAGCTATGCTGATTGTATGGGTTAACTCAAGGGTGTCA
CACTGAGGCCG
>SNORD115-like_59
GTGTCAATGATGAGAAACCCAGGCATCTCTTCTGATGGAGCTATGCTGATCATAAGGGTTAACTCAAGGTTATCA
CACTGAGCCCC
>SNORD115-like_60
GGGTCAATGATGAGAAACCCAGGCATCATTTCTGAAGGAGCTATGCTGATCTTATGGGTTAATTCAAGGGTGTTA
CACTGAGGCCC
>SNORD115-like_61
GAGTCAATGATGAGAAACTCAGACATTGCTTCTGCAGGAGCTCTGCTGATCAAAAGGGTTAACTCAAAGATGTCA
CACTGAGACCT
>SNORD115-like_62
TAGTCAATGATGATAAACCCAGGCATTACTTCTGAAGGAGCTCTGCTCATCTTATGGGTTAATTCAAAGGTGTCA
CACTGAGGTCC
>SNORD115-like_63
GGACCTGTTGATATCCATGATAGTCTTTAGTCTGACTCTTCTGAAGGAGCTATGCGGATCAAAATGGTTAATTCA
AGGGTGTCACACTGAGGCCC
>SNORD115-like_64
GGGTCAGTGATGAGAAACCCAGGCATCGCTTCTGAAGAAACTATACTGATCTTATGGGTTAATTGAAGGTTTTCA
CACTGAGACTA
>SNORD115-like_65
TGTCCCATAATGAACACCCCAGGCATTGCTACTGAAGAAGCTATGCTGATCATATGGATTAATTCAAGAGTGTCA
CACTGAGGTCC
>SNORD115-like_66
TGGGCATTGGTGAGAAACCAGGCATCACTTCTACAGGCAGCTATGCTGATCATATGTGTTATCTCAAGGTTGTCA
CACTGAGGCCC
>SNORD115-like_67
GGGTCAATGATGAGAACACCAGCCATCATTTCTGAAAGAGCTATGCTGATCTTTGGATCAATTCAAGGGTGTTACACTGAGGT
CC
>SNORD115-like_68
GGGTCAATGTTGAGAAACCCAGGCATCACTTCTAAAGGAGCTATGCTGATTTTATTTAATCATTAGATAATTCTAAGGTGCCA
CACTGAGGCCC

**Cluster of SNORD116-like sequences in *L. catta* (the ring-tailed lemur)**

>SNORD116-LIKE_1
TGCCTGGATCGATGATGTCTAAAAAAAATGGAAACTTTTGAAATCTGAACAAAATGAGTGAGGACTCGAGCTGAG
GTCCA
>SNORD116-LIKE_2
CGTACGTGTGCCTGGATCGATGATGTCTAAAAAAAATGGAAACTTTTGAAATCTGAACAAAATGAGTGAGGACTC
GAGCTGAGGTCC
>SNORD116-LIKE_3
TGTGCCTGGATCGATGATGTCTAAAAAAAATGGAAACTTTTGAAATCTGAACAAAATGAGTGAGGACTCGAGCTG
AGGTCC
>SNORD116_LIKE_4
CGTGTGCCTGGATCGATGATGTCTAAAAAAAATGGAAACTTTTGAAATCTGAACAAAATGAGTGAGGACTCGAGC
TGAGGTCC
>SNORD116_LIKE_5
CTGGATCGATGATGTCTAAAAAAAATGGAAACTTTTGAAATCTGAACAAAATGAGTGAGGACTCGAGCTGAGGTC
C
>SNORD116_LIKE_6
TGCCTGGATCGATGATGTCTAAAAAAAATGGAAACTTTTGAAATCTGAACAAAATGAGTGAGGACTCGAGCTGAG
GTCCA
>SNORD116_LIKE_7
ACGTGTGCCTGGATCGATGATGTCTAAAAAAAATGGAAACTTTTGAAATCTGAACAAAATGAGTGAGGACTCGAG
CTGAGGTCCAGCACG
>SNORD116_LIKE_8

TGCCTGGATCGATGATGTCTAAAAAAAATGGAAACTTTTGAAATCTGAACAAAATGAGTGAGGACTCGAGCTGAG
GTCCA
>SNORD116_LIKE_9
ACGTGTGCCTGGATCGATGATGTCTAAAAAAAATGGAAACTTTTGAAATCTGAACAAAATGAGTGAGGACTCGAG
CTGAGGTCCAGCACG
>SNORD116_LIKE_10
CTGGATCGATGATGTCTAAAAAAAATGGAAACTTTTGAAATCTGAACAAAATGAGTGAGGACTCGAGCTGAGGTC
CAGCA
>SNORD116_LIKE_11
TGCCTGGATCGATGATGTCTAAAAAAAATGGAAACTTTTGAAATCTGAACAAAATGAGTGAGGACTCGAGCTGAG
GTCCA
>SNORD116_LIKE_12
GTGTGCCTGGATCGATGATGTCTAAAAAAAATGGAAACTTTTGAAATCTGAACAAAATGAGTGAGGACTCGAGCT
GAGGTCC
>SNORD116_LIKE_13
TACGTGTGCCTGGATCGATGATGTCTAAAAAAAATGGAAACTTTTGAAATCTGAACAAAATGAGTGAGGACTCGA
GCTGAGGTCCAGCAC
>SNORD116_LIKE_14
GCCTGGATCGATGATGTCTAAAAAAAATGGAAACTTTTGAAATCTGAACAAAATGAGTGAGGACTCGAGCTGAGG
TCCAG
>SNORD116_LIKE_15
TGTGCCTGGATCGATGATGTCTAAAAAAAATGGAAACTTTTGAAATCTGAACAAAATGAGTGAGGACTCGAGCTG
AGGTCCAGCACGGGG
>SNORD116_LIKE_16
GTACGTGTGCCTGGATCGATGATGTCTAAAAAAAATGGAAACTTTTGAAATCTGAACAAAATGAGTGAGGACTCG
AGCTGAGGTCCA
>SNORD116_LIKE_17
GTGCCTGGATCGATGATGTCTAAAAAAAATGGAAACTTTTGAAATCTGAACAAAATGAGTGAGGACTCGAGCTGA
GGTCC
>SNORD116_LIKE_18
GTGCCTGGATCGATGATGTCTAAAAAAAATGGAAACTTTTGAAATCTGAACAAAATGAGTGAGGACTCGAGCTGA
GGTCC
>SNORD116_LIKE_19
ACGTGTGCCTGGATCGATGATGTCTAAAAAAAATGGAAACTTTTGAAATCTGAACAAAATGAGTGAGGACTCGAG
CTGAGGTCCAG
>SNORD116_LIKE_20
GCCTGGATCGATGATGTCTAAAAAAAATGGAAACTTTTGAAATCTGAACAAAATGAGTGAGGACTCGAGCTGAGG
TCCAG
>SNORD116_LIKE_21
ACGTGTGCCTGGATCGATGATGTCTAAAAAAAATGGAAACTTTTGAAATCTGAACAAAATGAGTGAGGACTCGAG
CTGAGGTCCA
>SNORD116_LIKE_22
CTGGATCGATGATGTCTAAAAAAAATGGAAACTTTTGAAATCTGAACAAAATGAGTGAGGACTCGAGCTGAGGTC
CAGCA
>SNORD116_LIKE_23
TTTCGTACGTGTGCCTGGATCGATGATGTCTAAAAAAAATGGAAACTTTTGAAATCTGAACAAAATGAGTGAGGA
CTCGAGCTGAGGTCCAGC
>SNORD116_LIKE_24
TGTGCCTGGATCGATGATGTCTAAAAAAAATGGAAACTTTTGAAATCTGAACAAAATGAGTGAGGACTCGAGCTG
AGGTCCA
>SNORD116_LIKE_25
CTGGATCGATGATGTCTAAAAAAAATGGAAACTTTTGAAATCTGAACAAAATGAGTGAGGACTCGAGCTGAGGTC
CAGCA
>SNORD116_LIKE_26
TGTGCCTGGATCGATGATGTCTAAAAAAAATGGAAACTTTTGAAATCTGAACAAAATGAGTGAGGACTCGAGCTG
AGGTC
>SNORD116_LIKE_27
CTGGATCGATGATGTCTAAAAAAAATGGAAACTTTTGAAATCTGAACAAAATGAGTGAGGACTCGAGCTGAGGTC
CAGCA
>SNORD116_LIKE_28

GTGTGCCTGGATCGATGATGTCTAAAAAAAAATGGAAACTTTTGAAATCTGAACAAAATGAGTGAGGACTCGAGCT
GAGGTCCA
>SNORD116_LIKE_29
ACGTGTGCCTGGATCGATGATGTCTAAAAAAAAATGGAAACTTTTGAAATCTGAACAAAATGAGTGAGGACTCGAG
CTGAGGTCCAGCACG
>SNORD116_LIKE_30
CGTACGTGTGCCTGGATCGATGATGTCTAAAAAAAAATGGAAACTTTTGAAATCTGAACAAAATGAGTGAGGACTC
GAGCTGAGGTCCAGC
>SNORD116_LIKE_31
TGTGCCTGGATCGATGATGTCTAAAAAAAAATGGAAACTTTTGAAATCTGAACAAAATGAGTGAGGACTCGAGCTG
AGGTCC
>SNORD116_LIKE_32
TACGTGTGCCTGGATCGATGATGTCTAAAAAAAAATGGAAACTTTTGAAATCTGAACAAAATGAGTGAGGACTCGA
GCTGAGGTCC
>SNORD116_LIKE_33
TGCCTGGATCGATGATGTCTAAAAAAAAATGGAAACTTTTGAAATCTGAACAAAATGAGTGAGGACTCGAGCTGAG
GTCCA
>SNORD116_LIKE_34
TGGATCGATGATGTCTAAAAAAAAATGGAAACTTTTGAAATCTGAACAAAATGAGTGAGGACTCGAGCTGAGGTCC
AGCAC
>SNORD116_LIKE_35
GTGCCTGGATCGATGATGTCTAAAAAAAAATGGAAACTTTTGAAATCTGAACAAAATGAGTGAGGACTCGAGCTGA
GGTCC
>SNORD116_LIKE_36
ACGTGTGCCTGGATCGATGATGTCTAAAAAAAAATGGAAACTTTTGAAATCTGAACAAAATGAGTGAGGACTCGAG
CTGAGGTCCAGCACG
>SNORD116_LIKE_37
CCTGGATCGATGATGTCTAAAAAAAAATGGAAACTTTTGAAATCTGAACAAAATGAGTGAGGACTCGAGCTGAGGT
CCAGC
>SNORD116_LIKE_38
GTGCCTGGATCGATGATGTCTAAAAAAAAATGGAAACTTTTGAAATCTGAACAAAATGAGTGAGGACTCGAGCTGA
GGTCC
>SNORD116_LIKE_39
GTGTGCCTGGATCGATGATGTCTAAAAAAAAATGGAAACTTTTGAAATCTGAACAAAATGAGTGAGGACTCGAGCT
GAGGTCCA
>SNORD116_LIKE_40
CCTGGATCGATGATGTCTAAAAAAAAATGGAAACTTTTGAAATCTGAACAAAATGAGTGAGGACTCGAGCTGAGGT
CCAGC
>SNORD116_LIKE_41
CGTGTGCCTGGATCGATGATGTCTAAAAAAAAATGGAAACTTTTGAAATCTGAACAAAATGAGTGAGGACTCGAGC
TGAGGTCCAGCACGG
>SNORD116_LIKE_42
GCACGTGTGCCTGGATCGATGATGTCTAAAAAAAAATGGAAACTTTTGAAATCTGAACAAAATGAGTGAGGACTCG
AGCTGAGGTCC
>SNORD116_LIKE_43
CTGGATCGATGATGTCTAAAAAAAAATGGAAACTTTTGAAATCTGAACAAAATGAGTGAGGACTCGAGCTGAGGTC
CAGCA
>SNORD116_LIKE_44
CTGGATCGATGATGTCTAAAAAAAAATGGAAACTTTTGAAATCTGAACAAAATGAGTGAGGACTCGAGCTGAGGTC
C
>SNORD116_LIKE_45
GTGTGCCTGGATCGATGATGTCTAAAAAAAAATGGAAACTTTTGAAATCTGAACAAAATGAGTGAGGACTCGAGCT
GAGGTCC
>SNORD116_LIKE_46
ACGTGTGCCTGGATCGATGATGTCTAAAAAAAAATGGAAACTTTTGAAATCTGAACAAAATGAGTGAGGACTCGAG
CTGAGGTCCAGCACG
>SNORD116_LIKE_47
GTACGTGTGCCTGGATCGATGATGTCTAAAAAAAAATGGAAACTTTTGAAATCTGAACAAAATGAGTGAGGACTCG
AGCTGAGGTCCAGCA
>SNORD116_LIKE_48

TACGTGTGCCTGGATCGATGATGTCTAAAAAAAATGGAAACTTTTGAAATCTGAACAAAATGAGTGAGGACTCGA
GCTGAGGTCCAGCAC
>SNORD116_LIKE_49
TACGTGTGCCTGGATCGATGATGTCTAAAAAAAATGGAAACTTTTGAAATCTGAACAAAATGAGTGAGGACTCGA
GCTGAGGTCCA
>SNORD116_LIKE_50
GTGCCTGGATCGATGATGTCTAAAAAAAATGGATACTTTTGAAATCTGAACAAAGTGAGTGAGGACTCGAGCTGA
GGTCC
>SNORD116_LIKE_51
GTGTGCCTGGATCGATGATGTCTAAAAAAAATGGAAACTTTTGAAATCTGAACACAGTGAGTGAGGACTCGAGCT
GAGGTCC
>SNORD116_LIKE_52
TACGTGTGCCTGGATCGATGATGTCTAAAAAAAATGGAAACTTTTGAAATCTGAACACAGTGAGTGAGGACTCGA
GCTGAGGTCCAGCAC
>SNORD116_LIKE_53
ATGTGCCTGGATCGATGATGTCTGAAAAAAATGGAAACTCTTGAAATCTGAACAAAATGAGTGAGGACTCGAGCT
GAGGTCCGGCACAGG
>SNORD116_LIKE_54
TACGTGTGCCTGGATCGATGATGTCTAAAAATATGGAAACTTTTGAAATCTGAACAAAATGAGTGAGGACTCGAG
CTGAGGTCCAGCACG
>SNORD116_LIKE_55
GCCTGGATCGATGATGTCTAAAAATATGGAAACTTTTGAAATCTGAACAAAATGAGTGAGGACTCGAGCTGAGGT
CCAGC

**Cluster of SNORD109 in *L. catta* (the ring-tailed lemur)**

>SNORD109_1
GGATCGATGATGAGCATAATTATCTGAGGATGCTGAGGGACTCATTGCAGATGTCAATCTAAAGTCT
>SNORD109_2
GGATCGATGATGAGCATAAGTGTCCGAGGACAGTGAGCGACACACTGCAGATGTCAATCTGAGGTCA
>SNORD109_3
GGATCGATGATGAGCCTAAGTGTCTGAGGACGGTGATGACACATGGCAGATGTCAATCTGAGGTCC
>SNORD109_4
GGATCGATGATGAGCCTAAGTGTCCGAGGACGGTGATGACACATGGCAGATGTCAATCTGAGGTCC
>SNORD109_5
GGATCGATGATGAGCCTAAGTGTCCGAGGACGGTGACGACACATGGCAGATGTCAATCTGAGGTCC
>SNORD109_6
GGATCGATGATGAGCCTAAGTGTCCGAGGACGGTGATGACACATTGCAGATGTCAATCTGAAGGTCC
>SNORD109_7
GGATCGATGATGAGCCTAAGTGTCCGAGGACGGTGATGACACATTGCAGATGTCAATCTGAAGGTCC
>SNORD109_8
GGATCGATGATGAGCCTAAGTGTCCGAGGACGGTGATGACACATTGCAGATGTCAATCTGAAGGTCC
>SNORD109_9
GGATCGATGATGAGCCTAAGTGTCCGAGGACGGTGATGACACATTGCAGATGTCAATCTGAAGGTCC
>SNORD109_10
GGATCGATGATGAGCCTAAGTGTCCGAGGACGGTGATGACACATGGCAGATGTACATCTGAGGTCC
>SNORD109_11
GGATCGATGATGAGCCTAAGTGTCCGAGGACGGTGATGACACATGGCAGATGTACATCTGAGGTCC
>SNORD109_12
GGATCGATGATGAGCCTAAGTGTCCGAGGACGGTGATGACACATGGCAGATGTACATCTGAGGTCC
>SNORD109_13
GGATCGATGATGAGCCTAAGTGTCCGAGGACGGTGACGACACATGGCAGATGTACATCTGAGGTCC
>SNORD109_14
GGATCGATGATGAGCCTAAGTGTCCGAGGACGGTGACGACACATGGCAGATGTACATCTGAGGTCC
>SNORD109_15
GGATCGATGATGAGCCTAAGTGTCCGAGGACGGTGACGACACATGGCAGATGTACATCTGAGGTCC
>SNORD109_16
GGATCGATGATGAGCCTAAGTGTCCGAGGACGGTGACGACACATGGCAGATGTACATCTGAGGTCC
>SNORD109_17

GGATCGATGATGAGCCTAAGTGTCCGAGGACGGTGACGACACATGGCAGATGTACATCTGAGGTCC
>SNORD109_18
GGATCGATGATGAGCCTAAGTGTCCGAGGACGGTGACGACACATGGCAGATGTACATCTGAGGTCC
>SNORD109_19
GGATCGATGATGAGCCTAAGTGTCCGAGGACGGTGACGACACATGGCAGATGTACATCTGAGGTCC
>SNORD109_20
GGATCGATGATGAGCCTAAGTGTCCGAGGACGGTGACGACACATGGCAGATGTACATCTGAGGTCC
>SNORD109_21
GGATCGATGATGAGCCTAAGTGTCCGAGGACGGTGACGACACATGGCAGATGTACATCTGAGGTCC
>SNORD109_22
GGATCGATGATGAGCCTAAGTGTCCGAGGACGGTGACGACACATGGCAGATGTACATCTGAGGTCC
>SNORD109_23
GGATCGATGATGAGCCTAAGTGTCCGAGGACGGTGACGACACATGGCAGATGTACATCTGAGGTCC
>SNORD109_24
GGATCGATGATGAGCCTAAGTGTCCGAGGACGGTGACGACACATGGCAGATGTACATCTGAGGTCC
>SNORD109_25
GGATCGATGATGAGCCTAAGTGTCCGAGGACGGTGACGACACATGGCAGATGTACATCTGAGGTCC
>SNORD109_26
GGATCGATGATGAGCCTAAGTGTCCGAGGACGGTGACGACACATGGCAGATGTACATCTGAGGTCC
>SNORD109_27
GGATCGATGATGAGCCTAAGTGTCCGAGGACGGTGACGACACATGGCAGATGTACATCTGAGGTCC
>SNORD109_28
GGATCGATGATGAGCCTAAGTGTCCGAGGACGGTGACGACACATGGCAGATGTACATCTGAGGTCC
>SNORD109_29
GGATCGATGATGAGCCTAAGTGTCCGAGGACGGTGACGACACATGGCAGATGTACATCTGAGGTCC
>SNORD109_30
GGATCGATGATGAGCCTAAGTGTCCGAGGACGGTGACGACACATGGCAGATGTACATCTGAGGTCC
>SNORD109_31
GGATCGATGATGAGCCTAAGTGTCCGAGGACGGTGACGACACATGGCAGATGTACATCTGAGGTCC
>SNORD109_32
GGATCGATGATGAGCCTAAGTGTCCGAGGACGGTGACGACACATGGCAGATGTACATCTGAGGTCC
>SNORD109_33
GGATCGATGATGAGCCTAAGTGTCCGAGGACGGTGACGACACATGGCAGATGTACATCTGAGGTCC
>SNORD109_34
GGATCGATGATGAGCCTAAGTGTCCGAGGACGGTGACGACACATGGCAGATGTACATCTGAGGTCC
>SNORD109_35
GGATCGATGATGAGCCTAAGTGTCCGAGGACGGTGACGACACATGGCAGATGTACATCTGAGGTCC
>SNORD109_36
GGATCGATGATGAGCCTAAGTGTCCGAGGACGGTGACGACACATGGCAGATGTACATCTGAGGTCC
>SNORD109_37
GGATCGATGATGAGCCTAAGTGTCCGAGGACGGTGACGACACATGGCAGATGTACATCTGAGGTCC
>SNORD109_38
GGATCGATGATGAGCCTAAGTGTCCGAGGACGGTGACGACACATGGCAGATGTACATCTGAGGTCC
>SNORD109_39
GGATCGATGATGAGCCTAAGTGTCCGAGGACGGTGACGACACATGGCAGATGTACATCTGAGGTCC
>SNORD109_40
GGATCGATGATGAGCCTAAGTGTCCGAGGACGGTGACGACACATGGCAGATGTACATCTGAGGTCC
>SNORD109_41
GGATCGATGATGAGCCTAAGTGTCCGAGGACGGTGACGACACATGGCAGATGTACATCTGAGGTCC
>SNORD109_42
GGATCGATGATGAGCCTAAGTGTCCGAGGACGGTGACGACACATGGCAGATGTACATCTGAGGTCC
>SNORD109_43
GGATCGATGATGAGCCTAAGTGTCCGAGGACGGTGACGACACATGGCAGATGTACATCTGAGGTCC
>SNORD109_44
GGATCGATGATGAGCCTAAGTGTCCGAGGACGGTGACGACACATGGCAGATGTACATCTGAGGTCC
>SNORD109_45
GGATCGATGATGAGCCTAAGTGTCCGAGGACGGTGACGACACATGGCAGATGTACATCTGAGGTCC
>SNORD109_46
GGATCGATGATGAGCCTAAGTGTCCGAGGACGGTGACGACACATGGCAGATGTACATCTGAGGTCC
>SNORD109_47
GGATCGATGATGAGCCTAAGTGTCCGAGGACGGTGACGACACATGGCAGATGTACATCTGAGGTCC

>SNORD109_48
GGATCGATGATGAGCCTAAGTGTCCGAGGACGGTGACGACACATGGCAGATGTACATCTGAGGTCC
>SNORD109_49
GGATCGATGATGAGCCTAAGTGTCCGAGGACGGTGACGACACATGGCAGATGTACATCTGAGGTCC
>SNORD109_50
GGATCGATGATGAGCCTAAGTGTCCGAGGACGGTGACGACACATGGCAGATGTACATCTGAGGTCC
>SNORD109_51
GGATCGATGATGAGCCTAAGTGTCCGAGGACGGTGACGACACATGGCAGATGTACATCTGAGGTCC
>SNORD109_52
GGATCGATGATGAGCCTAAGTGTCCGAGGACGGTGACGACACATGGCAGATGTACATCTGAGGTCC
>XSNORD109_53
GGATCGATGATGAGCCTAAGTGTCCGAGGACGGTGACGACACATGGCAGATGTACATCTGAGGTCC
>SNORD109_54
GGATCGATGATGAGCCTAAGTGTCCGAGGACGGCGACGACGCATGGCAGATGTACATCTGAGGTCC
>SNORD109_55
GGATCGATGATGAGCCTAAATGTTTGAGGACGGTTAAGGACACATGGCAGATGTACATCTGACGTCC

**Cluster of SNORD109 in *M. oregoni* (creeping vole)**

>SNORD109_1
GGATCGATGATGAGAAACCCTGTCTGAGGATGCTGAGAGACTCATGTCAATGGGAATCTGAGGTCC
>SNORD109_2
GGATTGATGATGATAAGAAGTCTCTGAGGATGCTGACTGACTCATACAAATGGCAATCTGAGGTCC
>SNORD109_3
GGATCGATGATGACAAGAAGTCTCTGAGGACAGTGACTGATTCAAACAAATGGCAATCTGAGGTCC
>SNORD109_4
GGATTGATGATGAGAACAAGTTTCTGAGGACGCTGATGGATTCAAAGAGAAAGCCTCTGAAGTCC
>SNORD109_5
GGATTGATGATGAGAACAAGTTTCTGAGGACGCTGATGGATTCAAAGAGAAAGCCTCTGAAGTCC
>SNORD109_6
GGATTGATGATGAAAACAGTTCTCTGAATATGGTGAATTACTCATAGAAATGGCAATCAGAGATCC
>SNORD109_7
GGATCGATGATGACAACAGGTTTCTGAGGACGCTGATTGACTCAAACATAACCCCTCTGAGGTCC
>SNORD109_8
GGATCGATGATGACAACAGGTTTCTGAGGACGCTGATTGACTCAAACATAACCCCTCTGAGGTCC
>SNORD109_9
GGATCGATGATGACAACAGGTTTCTGAGGACGCTGATTGACTCAAACATAACCCCTCTGAGGTCC
>SNORD109_10
GGATCGATGATGACAACAGGTTTCTGAGGACGCTGATTGACTCAAACATAACCCCTCTGAGGTCC
>SNORD109_11
GGATCGATGATGACAACAGGTTTCTGAGGACGCTGATTGACTCAAACATAACCCCTCTGAGGTCC
>SNORD109_12
GGATCGATGATGACAACAGGTTTCTGAGGACGCTGATTGACTCAAACATAACCCCTCTGAGGTCC
>SNORD109_13
GGATCGATGATGACAACAGGTTTCTGAGGACGCTGATTGACTCAAACATAACCCCTCTGAGGTCC
> SNORD109_14
GGATCGATGATGACAACAGGTTTCTGAGGACGCTGATTGACTCAAACATAACCCCTCTGAGGTCC
>SNORD109_15
GGATCGATGATGACAACAGGTTTCTGAGGACGCTGATTGACTCAAACATAACCCCTCTGAGGTCC
>SNORD109_16
GGATCGATGATGACAACAGGTTTCTGAGGACGCTGATTGACTCAAACATAACCCCTCTGAGGTCC
>SNORD109_17
GGATCGATGATGACAACAGGTTTCTGAGGACGCTGATTGACTCAAACATAACCCCTCTGAGGTCC
>SNORD109_18
GGATCGATGATGACAACAGGTTTCTGAGGACGCTGATTGACTCAAACATAACCCCTCTGAGGTCC
>SNORD109_19
GGATCGATGATGACAACAGGTTTCTGAGGACGCTGATTGACTCAAACATAACCCCTCTGAGGTCC
>SNORD109_20

GGATCGATGATGACAACAGGTTTCTGAGGACGCTGATTGACTCAAACATAACCCCTCTGAGGTCC
>SNORD109_21
GGATCGATGATGACAACAGGTTTCTGAGGACGCTGATTGACTCAAACATAACCCCTCTGAGGTCC
>SNORD109_22
GGATCGATGATGACAACAGGTTTCTGAGGACGCTGATTGACTCAAACATAACCCCTCTGAGGTCC
>SNORD109_23
GGATCGATGATGACAACAGGTTTCTGAGGACGCTGATTGACTCAAACATAACCCCTCTGAGGTCC

>SNORD109_24
GGATCGATGATGACAACAGGTTTCTGAGGACGCTGATTGACTCAAACATAACCCCTCTGAGGTCC
>SNORD109_25
GGATCGATGATGACAACAGGTTTCTGAGGACGCTGATTGACTCAAACATAACCCCTCTGAGGTCC
>SNORD109_26
GGATCGATGATGACAACAGGTTTCTGAGGACGCTGATTGACTCAAACATAACCCCTCTGAGGTCC
>SNORD109_27
GGATCGATGATGACAACAGGTTTCTGAGGACGCTGATTGACTCAAACATAACCCCTCTGAGGTCC
>SNORD109_28
GGATCGATGATGACAACAGGTTTCTGAGGACGCTGATTGACTCAAACATAACCCCTCTGAGGTCC
>SNORD109_29
GGATCGATGATGACAACAGGTTTCTGAGGACGCTGATTGACTCAAACATAACCCCTCTGAGGTCC
>SNORD109_30
GGATCGATGATGACAACAGGTTTCTGAGGACGCTGATTGACTCAAACATAACCCCTCTGAGGTCC
>SNORD109_31
GGATCGATGATGACAACAGGTTTCTGAGGACGCTGATTGACTCAAACATAACCCCTCTGAGGTCC
>SNORD109_32
GGATTGATGATGAGAACATGTTTCTGAGGACGCTGATGGAGTCAAAGAGAAAGCCTCTGAGGTCC
>SNORD109_33
GGATTGATGATGAGAACATGTTTCTGAGGACGCTGATGGAGTCAAAGAGAAAGCCTCTGAGGTCC
>SNORD109_34
GGATTGATGATGAGAACATGTTTCTGAGGACGCTGATGGAGTCAAAGAGAAAGCCTCTGAGGTCC
>SNORD109_35
GGATTGATGATGAGAACATGTTTCTGAGGACGCTGATGGAGTCAAAGAGAAAGCCTCTGAGGTCC
>SNORD109_36
GGATTGATGATGAGAACATGTTTCTGAGGACGCTGATGGAGTCAAAGAGAAAGCCTCTGAGGTCC
>SNORD109_37
GGATCGATGATGACAACAGGTTTCTGAGGACGCTGATTGACTCAAACATAACCCCTCTCAGGTCC
>SNORD109_38
GGATCAGTGATAAGAACAACAATGTGAGGATGCTGAACAACTCTTAGGAATTGCAAACTCAGGTCC

**Cluster of SNORD109 in Meriones unguiculatus (Mongolian gerbil)**

>SNORD109_1
GATCTATGATGACAACAGTTATCTGAGGACGTTGACTGACTTGAGCAGATGTAATCTGAGGTCC
>SNORD109_2
GATCTATGATGACAACAGTTATCTGAGGACGTTGACTGACTTGAGCAGATGTAATCTGAGGTCC
>SNORD109_3
GATCTATGATGACAACAGGTATCTGAGGACGTTGACTGACTTGAGCAGATGTAATCTGAGGTCC
>SNORD109_4
GATCTATGATGACAACAGGTATCTGAGGACGTTGACTGACTTGAGCAGATGTAATCTGAGGTCC
>SNORD109_5
GATCTATGATGACAACAGGTATCTGAGGACGTTGACTGACTTGAGCAGATGTAATCTGAGGTCC
>SNORD109_6
GATCTATGATGACAACAGGTATCTGAGGACGTTGACTGACTTGAGCAGATGTAATCTGAGGTCC
>SNORD109_7
GATCTATGATGACAACAGGTATCTGAGGACGTTGACTGACTTGAGCAGATGTAATCTGAGGTCC
>SNORD109_8
GATCTATGATGACAACAGGTATCTGAGGACGTTGACTGACTTGAGCAGATGTAATCTGAGGTCC

>SNORD109_9
GATCTATGATGACAACAGGTATCTGAGGACGTTGACTGACTTGAGCAGATGTAATCTGAGGTCC
>SNORD109_10
GATCTATGATGACAACAGGTATCTGAGGACGTTGACTGACTTGAGCAGATGTAATCTGAGGTCC
>SNORD109_11
GATCTATGATGACAACAGGTATCTGAGGACGTTGACTGACTTGAGCAGATGTAATCTGAGGTCC
>SNORD109_12
GATCTATGATGACAACAGGTATCTGAGGACGTTGACTGACTTGAGCAGATGTAATCTGAGGTCC
>SNORD109_13
GATCTATGATGACAACAGGTATCTGAGGACGTTGACTGACTTGAGCAGATGTAATCTGAGGTCC
>SNORD109_14
GATCTATGATGACAACAGGTATCTGAGGACGTTGACTGACTTGAGCAGATGTAATCTGAGGTCC
>SNORD109_15
GATCTATGATGACAACAGGTATCTGAGGACGTTGACTGACTTGAGCAGATGTAATCTGAGGTCC
>SNORD109_16
GATCTATGATGACAACAGGTATCTGAGGACGTTGACTGACTTGAGCAGATGTAATCTGAGGTCC
>SNORD109_17
GATCTATGATGACAACAGGTATCTGAGGACGTTGACTGACTTGAGCAGATGTAATCTGAGGTCC
>SNORD109_18
GATCTATGATGACAACAGGTATCTGAGGACGTTGACTGACTTGAGCAGATGTAATCTGAGGTCC
>SNORD109_19
GATCTATGATGACAACAGGTATCTGAGGACGTTGACTGACTTGAGCAGATGTAATCTGAGGTCC
>SNORD109_20
GATCTATGATGACAACAGGTATCTGAGGACGTTGACTGACTTGAGCAGATGTAATCTGAGGTCC
>SNORD109_21
GATCTATGATGACAACAGGTATCTGAGGACGTTGACTGACTTGAGCAGATGTAATCTGAGGTCC
>SNORD109_22
GATCTATGATGACAACAGGTATCTGAGGACGTTGACTGACTTGAGCAGATGTAATCTGAGGTCC
>SNORD109_23
GATCTATGATGACAACAGGTATCTGAGGACGTTGACTGACTTGAGCAGATGTAATCTGAGGTCC
>SNORD109_24
GATCTATGATGACAACAGGTATCTGAGGACGTTGACTGACTTGAGCAGATGTAATCTGAGGTCC
>SNORD109_25
GATCTATGATGACAACAGGTATCTGAGGACGTTGACTGACTTGAGCAGATGTAATCTGAGGTCC
>SNORD109_26
GATCTATGATGACAACAGGTATCTGAGGACGTTGACTGACTTGAGCAGATGTAATCTGAGGTCC
>SNORD109_27
GATCTATGATGACAACAGGTATCTGAGGACGTTGACTGACTTGAGCAGATGTAATCTGAGGTCC
>SNORD109_28
GATCTATGATGACAACAGGTATCTGAGGACGTTGACTGACTTGAGCAGATGTAATCTGAGGTCC
>SNORD109_29
GATCTATGATGACAACAGGTATCTGAGGACGTTGACTGACTTGAGCAGATGTAATCTGAGGTCC
>SNORD109_30
GATCTATGATGACAACAGGTATCTGAGGACGTTGACTGACTTGAGCAGATGTAATCTGAGGTCC
>SNORD109_31
GATCTATGATGACAACAGGTATCTGAGGACGTTGACTGACTTGAGCAGATGTAATCTGAGGTCC
>SNORD109_32
GATCTATGATGACAACAGGTATCTGAGGACGTTGACTGACTTGAGCAGATGTAATCTGAGGTCC
>SNORD109_33
GATCTATGATGACAACAGGTATCTGAGGACGTTGACTGACTTGAGCAGATGTAATCTGAGGTCC
>SNORD109_34
GATCTATGATGACAACAGGTATCTGAGGACGTTGACTGACTTGAGCAGATGTAATCTGAGGTCC
>SNORD109_35
GATCTATGATGACAACAGGTATCTGAGGACGTTGACTGACTTGAGCAGATGTAATCTGAGGTCC
>SNORD109_36
GATCTATGATGACAACAGGTATCTGAGGACGTTGACTGACTTGAGCAGATGTAATCTGAGGTCC
>SNORD109_37

GATCTATGATGACAACAGGTATCTGAGGACGTTGACTGACTTGAGCAGATGTAATCTGAGGTCC
>SNORD109_38
GATCTATGATGACAACAGGTATCTGAGGACGTTGACTGACTTGAGCAGATGTAATCTGAGGTCC
>SNORD109_39
GATCTATGATGACAACAGGTATCTGAGGACGTTGACTGACTTGAGCAGATGTAATCTGAGGTCC
>SNORD109_40
GATCTATGATGACAACAGGTATCTGAGGACGTTGACTGACTTGAGCAGATGTAATCTGAGGTCC
>SNORD109_41
GATCTATGATGACAACAGGTATCTGAGGACGTTGACTGACTTGAGCAGATGTAATCTGAGGTCC
>SNORD109_42
GATCTATGATGACAACAGGTATCTGAGGACGTTGACTGACTTGAGCAGATGTAATCTGAGGTCC
>SNORD109_43
GATCTATGATGACAACAGGTATCTGAGGACGTTGACTGACTTGAGCAGATGTAATCTGAGGTCC
>SNORD109_44
GATCTATGATGACAACAGGTATCTGAGGACGTTGACTGACTTGAGCAGATGTAATCTGAGGTCC
>SNORD109_45
GATCTATGATGACAACAGGTATCTGAGGACGTTGACTGACTTGAGCAGATGTAATCTGAGGTCC
>SNORD109_46
GATCTATGATGACAACAGGTATCTGAGGACGTTGACTGACTTGAGCAGATGTAATCTGAGGTCC
>SNORD109_47
GATCTATGATGACAACAGGTATCTGAGGACGTTGACTGACTTGAGCAGATGTAATCTGAGGTCC
>SNORD109_48
GATCTATGATGACAACAGGTATCTGAGGACGTTGACTGACTTGAGCAGATGTAATCTGAGGTCC
>SNORD109_49
GATCTATGATGACAACAGGTATCTGAGGACGTTGACTGACTTGAGCAGATGTAATCTGAGGTCC
>SNORD109_50
GATCTATGATGACAACAGGTATCTGAGGACGTTGACTGACTTGAGCAGATGTAATCTGAGGTCC
>SNORD109_51
GATCTATGATGACAACAGGTATCTGAGGACGTTGACTGACTTGAGCAGATGTAATCTGAGGTCC
>SNORD109_52
GATCTATGATGACAACAGGTATCTGAGGACGTTGACTGACTTGAGCAGATGTAATCTGAGGTCC
>SNORD109_53
GATCTATGATGACAACAGGTATCTGAGGACGTTGACTGACTTGAGCAGATGTAATCTGAGGTCC
>SNORD109_54
GATCTATGATGACAACAGGTATCTGAGGACGTTGACTGACTTGAGCAGATGTAATCTGAGGTCC
>SNORD109_55
GATCTATGATGACAACAGGTATCTGAGGACGTTGACTGACTTGAGCAGATGTAATCTGAGGTCC
>SNORD109_56
GATCTATGATGACAACAGGTATCTGAGGACGTTGACTGACTTGAGCAGATGTAATCTGAGGTCC
>SNORD109_57
GATCTATGATGACAACAGGTATCTGAGGACGTTGACTGACTTGAGCAGATGTAATCTGAGGTCC
>SNORD109_58
GATCTATGATGACAACAGGTATCTGAGGACGTTGACTGACTTGAGCAGATGTAATCTGAGGTCC
>SNORD109_59
GATCTATGATGACAACAGGTATCTGAGGACGTTGACTGACTTGAGCAGATGTAATCTGAGGTCC
>SNORD109_60
GATCTATGATGACAACAGGTATCTGAGGACGTTGACTGACTTGAGCAGATGTAATCTGAGGTCC
>SNORD109_61
GATCTATGATGACAACAGGTATCTGAGGACGTTGACTGACTTGAGCAGATGTAATCTGAGGTCC
>SNORD109_62
GATCTATGATGACAACAGGTATCTGAGGACGTTGACTGACTTGAGCAGATGTAATCTGAGGTCC
>SNORD109_63
GATCTATGATGACAACAGGTATCTGAGGACGTTGACTGACTTGAGCAGATGTAATCTGAGGTCC
>SNORD109_64
GATCTATGATGACAACAGGTATCTGAGGACGTTGACTGACTTGAGCAGATGTAATCTGAGGTCC
>SNORD109_65
GATCTATGATGACAACAGGTATCTGAGGACGTTGACTGACTTGAGCAGATGTAATCTGAGGTCC

>SNORD109_66
GATCTATGATGACAACAGGTATCTGAGGACGTTGACTGACTTGAGCAGATGTAATCTGAGGTCC
>SNORD109_67
GATCTATGATGACAACAGGTATCTGAGGACGTTGACTGACTTGAGCAGATGTAATCTGAGGTCC
>SNORD109_68
GATCTATGATGACAACAGGTATCTGAGGACGTTGACTGACTTGAGCAGATGTAATCTGAGGTCC
>SNORD109_69
GATCTATGATGACAACAGGTATCTGAGGACGTTGACTGATTTGAGCAGATGTAATCTGAGGTCC
>SNORD109_70
GATCTATGATGACAACAGGTATCTGAGGACGTTGACTGACTTGAGCAGATGTGATCTGAGGTCC
>SNORD109_71
GATCTATGATGACAACAGGTATCTGAGGACGTTGACTGACTTGAGCAGATGTGATCTGAGGTCC
>SNORD109_72
GATCTATGATGACAACAGGTATCTGAGGACGTTGACTGACTTGAGCAGATGTGATCTGAGGTCC
>SNORD109_73
GATCTATGATGACAACAGGTATCTGAGGACGTTGACTGACTTGAGCAGATGTGATCTGAGGTCC
>SNORD109_74
GATCTGTGATGACAACAGGTATCTGAGGACGTTGACTGACTTGAGCAGATGTAATCTGAGGTCC
>SNORD109_75
GATCTATGATGACAACAGGTATCTGAGGACGTTGACTGACTTGAGCAGATGTAATCTGATGTCC
>SNORD109_76
GATCTATGATGACAACAGGTATCTGAGGACGTTGACTGACTTGAGCAGATGTGATCTGAGGTCC
>SNORD109_77
GATCTATGATGACAACAGGTATCTGAGGACGTTGACTGACTTGAGCAGATGTAATCTGATGTCC
>SNORD109_78
GATCTATGATGACAACAGGTATCTGAGGACGTTGACTGACTTGAGCAGATGTGATCTGAGGTCC
>SNORD109_79
GATCTATGATGACAACAGGTATCTGAGGACGTTGACTGACTTGAGCAGATGTAATCTGAGATCC
>SNORD109_80
GATCTATGATGACAACAGGTATCTGAGGACGTTGACTGACTTGAGCAGATGTAATCTGAGATCC
>SNORD109_81
GATCTATGATGACAACAGGTATCTGAGGACGTTGACTGACTTGAGCAGATGTAATCTGAGATCC
>SNORD109_82
GATCTATGATGACAACAGGTATCTGAGGACGTTGACTGACTTGAGCAGATGTAATCTGAGATCC
>SNORD109_83
GATCTATGATGACAACAGGTATCTGAGGACGTTGACTGACTTGAGCAGATGTAATCTGAGATCC
>SNORD109_84
GATCTATGATGACAACAGGTATCTGAGGACGTTGACTGACTTGAGCAGATGTAATCTGAGATCC
>SNORD109_85
GATCTATGATGACAACAGGTATCTGAGGACGTTGACTGACTTGAGCAGATGTAATCTGAGATCC
>SNORD109_86
GATCTATGATGACAACAGGTATCTGAGGACGTTGACTGACTTGAGCAGATGTAATCTGAGATCC
>SNORD109_87
GATCTATGATGACAACAGGTATCTGAGGACGTTGACTGACTTGAGCAGATGTAATCTGAGATCC
>SNORD109_88
GATCTATGATGACAACAGGTATCTGAGGACGTTGACTGACTTGAGCAGATGTAATCTGAGATCC
>SNORD109_89
GATCTATGATGACAACAGGTATCTGAGGACGTTGACTGACTTGAGCAGATGTAATCTGAGATCC
>SNORD109_90
GATCTATGATGACAACAGGTATCTGAGGACGTTGACTGACTTGAGCAGATGTAATCTGAGATCC
>SNORD109_91
GATCTATGATGACAACAGGTATCTGAGGACGTTGACTGACTTGAGCAGATGTAATCTGAGATCC
>SNORD109_92
GATCTATGATGACAACAGGTATCTGAGGACGTTGACTGACTTGAGCAGATGTAATCTGAGATCC
>SNORD109_93
GATCTATGATGACAACAGGTATCTGAGGACGTTGACTGACTTGAGCAGATGTGATCTGAGGTCC
>SNORD109_94

GATCTATGATGACAACAGGTATCTGAGGACGTTGACTGACTTGAGCAGATGTGATCTGAGGTCC
>SNORD109_95
GATCTATGATGACAACAGGTATCTGAGGACGTTGACTGACTTGAGCAGATGTGATCTGAGGTCC
>SNORD109_96
GATCTATGATGACAACAGGTATGTGAGGACGTTGACTGACTTGAGCAGATGTAATCTGAGGTCC
>SNORD109_97
GATCTATGATGACAACAGGTATGTGAGGACGTTGACTGACTTGAGCAGATGTAATCTGAGGTCC
>SNORD109_98
GATCTATGATGACAACAGGTATCTGAGGACGTTGACTGACTTGAGCAGATGTGATCTGAGGTCC
>SNORD109_99
GATCTATGATGACAACAGGTATCTGAGGACGTTGACTGACTTGAGCAGATGTGATCTGAGGTCC
>SNORD109_100
GATCTATGATGACAACAGGTATCTGAGGACGTTGACTGACTTGAGCAGATGTGATCTGAGGTCC
>SNORD109_101
GATCTATGATGACAACAGGTATCTGAGGACGTTGACTGACTTGAGCAGATGTGATCTGAGGTCC
>SNORD109_102
GATCTATGATGACAACAGGTATCTGAGGACGTTGACTGACTTGAGCAGATGTGATCTGAGGTCC
>SNORD109_103
GATCTATGATGACAACAGGTATCTGAGGACGTTGACTGACTTGAGCAGATGTGATCTGAGGTCC
>SNORD109_104
GATCTATGATGACAACAGGTATCTGAGGACGTTGACTGACTTGAGCAGATGTGATCTGAGGTCC
>SNORD109_105
GATCTATGATGACAACAGGTATCTGAGGACGTTGACTGACTTGAGCAGATGTGATCTGAGGTCC
>SNORD109_106
GATCTATGATGACAACAGGTATCTGAGGACGTTGACTGACTTGAGCAGATGTGATCTGAGGTCC
>SNORD109_107
GATCTATGATGACAACAGGTATCTGAGGACGTTGACTGACTTGAGCAGATGTAATCTAAGGTCC
>SNORD109_108
GATCTATGATGACAACAGGTATCTGAGGACGTTGACTGAATTGAGCAGATGTAATCTGAGGTCC
>SNORD109_109
GATCTATGATGACAACAGGTATCTGAGGACGTTGACTGACTTGAGCAGATGTGATCTGAGGTCC
>SNORD109_110
GATCTATGATGACAACAGGTATCTGAGGATGTTGACTGACTTGAGCAGATGTAATCTGAGGTCC
>SNORD109_111
GATCTATGATGACAATAGGTATCTGAGGACGTTGACTGACTTGAGCAGATGTAATCTGAGGTCC
>SNORD109_112
GATCTATGATGACAATAGGTATCTGAGGACGTTGACTGACTTGAGCAGATGTAATCTGAGGTCC
>SNORD109_113
GATCTATGATGACAACAGGTATCTGAGGACGTTGACTGACTTGAGCAGATGTGATCTGAGGTCC
>SNORD109_114
GATCTATGATGACAACAGGTATCTGAGGACGTTGACTGACTTGAGCAGATGTGATCTGAGGTCC
>SNORD109_115
GATCTATGATGACAACAGGTATCTGAGGACGTTGACTGACTTGAGCAGATGTGATCTGAGGTCC
>SNORD109_116
GATCTATGATGACAACAGGTATCTGAGGACGTTGACTGAATTGAGCAGATGTAATCTGAGGTCC
>SNORD109_117
GATCTATGATGACAACAGGTATCTGAGGACGTTGACTGACTTGAGCAGATGTGATCTGAGGTCC
>SNORD109_118
GATCTATGATGACAACAGGTATCTGAGGACGTTGACTGACTTGAGCAGATGTGATCTGAGGTCC
>SNORD109_119
GATCTATGATGACAATAGGTATCTGAGGACGTTGACTGACTTGAGCAGATGTAATCTGAGGTCC
>SNORD109_120
GATCTATGATGACAACAGGTATCTGAGGACGTTGACTGACTTGAGCAGATGTGATCTGAGGTCC
>SNORD109_121
GATCTATGATGACAACAGGTATCTGAGGACGTTGACTGACTTGAGCAGATGTAATCTAAGGTCC
>SNORD109_122
GATCTGTGATGACAACAGGTATCTGAGGACGTTGACTGACTTGAGCAGATGTAATCTGAGGTCC

>SNORD109_123
GATCTGTGATGACAACAGGTATCTGAGGACGTTGACTGACTTGAGCAGATGTAATCTGAGGTCC
>SNORD109_124
GATATATGATGACAACAGGTATCTGAGGACGTTGACTGACTTGAGCAGATGTAATCTGAGGTCC
>SNORD109_125
GATATATGATGACAACAGGTATCTGAGGACGTTGACTGACTTGAGCAGATGTAATCTGAGGTCC
>SNORD109_126
GATCTATGATGACAACAGGTATCTGAGGACGTTGACTGACTTGAGCAGATGTGATCTGAGGTCC
>SNORD109_127
GATCTATGATGACAACAGGTATCTGAGGACGTTGACTGACTTGAGCAGATGTGATCTGAGGTCC
>SNORD109_128
GATCTGTGATGACAACAGGTATCTGAGGACGTTGACTGACTTGAGCAGATGTAATCTGAGGTCC
>SNORD109_129
GATCTATGATGACAACAGGTATCTGAGGACGTTGACTGACTTGAGCAGATATAATCTGAGGTCC
>SNORD109_130
GATCTGTGATGACAACAGGTATCTGAGGACGTTGACTGACTTGAGCAGATGTAATCTGAGGTCC
>SNORD109_131
GATCTATGATGACAACAGGTATCTGAGGACGTTGACTGACTTGAGCAGATGTGATCTGACGTCC
>SNORD109_132
GATCTATGATGACAACAGGTATCTGAGGATGTTGACTGATTTGAGCAGATGTAATCTGAGGTCC
>SNORD109_133
GATCTATGATGACAACAGGTATCTGAGGACGTTGACTGACTTGAACAGATGCAATCTGAGGTCC
>SNORD109_134
GATCTGTGATGACAACAGGTATCTGAGGACGTTGACTGACTTGAGCAGATGTGATCTGAGGTCC
>SNORD109_135
GATCTATGATGACAACAGGTATCTGAGGACGTTGACTGACTTGAGCAGATGTGATCTGACGTCC
>SNORD109_136
GATCTATGATGACAACAAGTATCTGAGGACGTTGACTGACTTGAGAAGATGTAATCTGAGGTCC
>SNORD109_137
GATCTCTGATGACAACAGGTATCTGAGGACGTTGACTGACTTGAGCAGATGTGATCTGAGGTCC
>SNORD109_138
GATCTATGATGACAACAGGTATCTGAGGACATTGACTGACTTGAGCAGATGTGATCTGAGGTCC
>SNORD109_139
GATCTATGATGACAACAGGTATCTGAGGACATTGACTGACTTGAGCAGATGTGATCTGAGGTCC
>SNORD109_140
GATCTATGATGACAACAGGTATCTGAGGACATTGACTGACTTGAGCAGATGTGATCTGAGGTCC
>SNORD109_141
GATCTATGATGACAACAGTTATCTGAGGACATTGACTGACTTGAGAAGATGTGATCTGAGGTCC
>SNORD109_142
GATCTATGATGACAACAGGTATCTGAGGACATTGACTGACTTGAGCAGATGTGATCTGAGGTCC
>SNORD109_143
GATCTGTGATGACAACAGATATCTGAGGACGTTGACTGACTTGAGCAGATGTGATCTGAGGTCC
>SNORD109_144
GATCTGTGATGACAACAGATATCTGAGGACGTTGACTGACTTGAGCAGATGTGATCTGAGGTCC
>SNORD109_145
GATCTGTGATGACAACAGGTATCTGAGGACGTTGACTGACTTGAGCAGATGTGATCTGAGGTCC
>SNORD109_146
GATCTGTGATGACAACAGGTATCTGAGGACATTGACTGACTTGAGCAGATGTAATCTGAGGTCC
>SNORD109_147
GATCTGTGATGACAACAGGTATCTGAGGACATTGACTGACTTGAGCAGATGTAATCTGAGGTCC
>SNORD109_148
GATCTATGATGACAACAGGTATCTGAGGACGTTGACTGACTTGAACAGATGCGATCTGAGGTCC
>SNORD109_149
GATCTATGATGACAACAGGTATCTGAGGACGTTGACTGACTTGAACAGATGCGATCTGAGGTCC
>SNORD109_150
GATCTATGATGACAACAGGTATCTGAGGACGTTGACTGACTTGAACAGATGCGATCTGAGGTCC
>SNORD109_151

GATCTATGATGACAACAGGTATCTGAGGACGTTGACTGACTTGAACAGATGCGATCTGAGGTCC
>SNORD109_152
GATCTATGATGACAACAGGTATCTGAGGACGTTGACTGACTTGAACAGATGCGATCTGAGGTCC
>SNORD109_153
GATCTATGATGACAACAGGTATCTGAGGACGTTGACTGACTTGAACAGATGCGATCTGAGGTCC
>SNORD109_154
GATCTATGATGACAACAGGTATCTGAGGACGTTGACTGACTTGAACAGATGCGATCTGAGGTCC
>SNORD109_155
GATCTATGATGACAACAGGTATCTGAGGACGTTGACTGACTTGAACAGATGCGATCTGAGGTCC
>SNORD109_156
GATCTATGATGACAACAGGTATCTGAGGACGTTGACTGACTTGAACAGATGCGATCTGAGGTCC
>SNORD109_157
GATCTATGATGACAACAGGTATCTGAGGACGTTGACTGACTTGAACAGATGCGATCTGAGGTCC
>SNORD109_158
GATCTATGATGACAACAGGTATCTGAGGACGTTGACTGACTTGAACAGATGCGATCTGAGGTCC
>SNORD109_159
GATCTATGATGACAACATGTATCTGAGGACGTTGACTGACTTGAACAGATGTGATCTGAGGTCC
>SNORD109_160
GATCTATGATGACAACAGGTATCTGAGGACGTTGACTGACTTGAACAGATGCGATCTGAGGTCC
>SNORD109_161
GATCTATGATGACAACAGGTATCTGAGGACGTTGACTGACTTGAACAGATGCGATCTGAGGTCC
>SNORD109_162
GATCTATGATGACAACAGGTATCTGAGGACGTTGACTGACTTGAACAGATGCGATCTGAGGTCC
>SNORD109_163
GATCTATGATGACAACAGGTATCTGAGGACGTTGACTGACTTGAACAGATGCGATCTGAGGTCC
>SNORD109_164
GATCTATGATGACAACAGGTATCTGAGGACGTTGACTGACTTGAACAGATGCGATCTGAGGTCC
>SNORD109_165
GATCTATGATGACAACAGGTATCTGAGGACGTTGACTGACTTGAACAGATGCGATCTGAGGTCC
>SNORD109_166
GATCTATGATGACAACAGGTATCTGAGGACGTTGACTGACTTGAACAGATGCGATCTGAGGTCC
>SNORD109_167
GATCTATGATGACAACAGGTATCTGAGGACGTTGACTGACTTGAACAGATGCGATCTGAGGTCC
>SNORD109_168
GATCTATGATGACAACAGGTATCTGAGGACGTTGACTGACTTGAACAGATGCGATCTGAGGTCC
>SNORD109_169
GATCTATGATGACAACAGGTATCTGAGGACGTTGACTGACTTGAACAGATGCGATCTGAGGTCC
>SNORD109_170
GATCTATGATGACAACAGGTATCTGAGGACGTTGACTGACTTGAACAGATGCGATCTGAGGTCC
>SNORD109_171
GATCTATGATGACAACAGGTATCTGAGGACGTTGACTGACTTGAACAGATGCGATCTGAGGTCC
>SNORD109_172
GATCTATGATGACAACAGGTATCTGAGGACGTTGACTGACTTGAACAGATGCGATCTGAGGTCC
>SNORD109_173
GATCTATGATGACAACAGGTATCTGAGGACGTTGACTGACTTGAACAGATGCGATCTGAGGTCC
>SNORD109_174
GATCTATGATGACAACAGGTATCTGAGGACATTGACTGACTTGAGAAGATGTGATCTGAGGTCC
>SNORD109_175
GATCTATGATGACAACAGGTATCTGAGGACGTTGACTGACTTGAACAGATGCGATCTGAGGTCC
>SNORD109_176
GATCTATGATGACAACAGGTATCTGAGGACGTTGACTGACTTGAACAGATGCGATCTGAGGTCC
>SNORD109_177
GATCTATGATGACAACAGGTATCTGAGGACGTTGACTGACTTGAACAGATGCGATCTGAGGTCC
>SNORD109_178
GATCTATGATGACAACAGGTATCTGAGGACGTTGACTGACTTGAACAGATGCGATCTGAGGTCC
>SNORD109_179
GATCTATGATGACAACAGGTATCTGAGGACGTTGACTGACTTGAACAGATGCGATCTGAGGTCC

>SNORD109_180
GATCTATGATGACAACAGGTATCTGAGGACGTTGACTGACTTGAACAGATGCGATCTGAGGT
>SNORD109_181
GATCTATGATGACAACAGGTCTCTGAGGACGTTGACTGACTTGAACAGATGCGATCTGAGGTCC
>SNORD109_182
GATCTCTGATGACAACAGGTATCTGAGGACGTTGACTGACTTGAACAGATGCGATCTGAGGTCC
>SNORD109_183
GATCTATGATGACAACAGGTATCTGAGGACGTTGACTGACTTGAACAGATGCGATATGAGGTCC
>SNORD109_184
GATCTATGATGACAACAGGTCTCTGAGGACGTTGACTGACTTGAACAGATGCGATCTGAGGTCC
>SNORD109_185
GATCTATGATGACAACAGGTCTCTGAGGACGTTGACTGACTTGAACAGATGCGATCTGAGGTCC
>SNORD109_186
GATCTATGATGACAACAGGTCTCTGAGGACGTTGACTGACTTGAACAGATGCGATCTGAGGTCC
>SNORD109_187
GATCTATGATGACAACAGGTCTCTGAGGACGTTGACTGACTTGAACAGATGCGATCTGAGGTCC
>SNORD109_188
GATCTATGATGACAACAGGTCTCTGAGGACGTTGACTGACTTGAACAGATGCGATCTGAGGTCC
>SNORD109_189
GATCTATGATGACAACAGGTCTCTGAGGACGTTGACTGACTTGAACAGATGCGATCTGAGGTCC
>SNORD109_190
GATCTATGATGACAACAGGTCTCTGAGGACGTTGACTGACTTGAACAGATGCGATCTGAGGTCC
>SNORD109_191
GATCTATGATGACAACAGGTCTCTGAGGACGTTGACTGACTTGAACAGATGCGATCTGAGGTCC
>SNORD109_192
GATCTATGATGACAACAGGTCTCTGAGGACGTTGACTGACTTGAACAGATGCGATCTGAGGTCC
>SNORD109_193
GATCTATGATGACAACAGGTCTCTGAGGACGTTGACTGACTTGAACAGATGCGATCTGAGGTCC
>SNORD109_194
GATCTATGATGACAACAGGTCTCTGAGGACGTTGACTGACTTGAACAGATGCGATCTGAGGTCC
>SNORD109_195
GATCTATGATGACAACAGGTCTCTGAGGACGTTGACTGACTTGAACAGATGCGATCTGAGGTCC
>SNORD109_196
GATCTATGATGACAACAGGTCTCTGAGGACGTTGACTGACTTGAACAGATGCGATCTGAGGTCC
>SNORD109_197
GATCTATGATGACAACAGGTCTCTGAGGACGTTGACTGACTTGAACAGATGCGATCTGAGGTCC
>SNORD109_198
GATCTATGATGACAACAGGTCTCTGAGGACGTTGACTGACTTGAACAGATGCGATCTGAGGTCC
>SNORD109_199
GATCTATGATGACAACAGGTCTCTGAGGACGTTGACTGACTTGAACAGATGCGATCTGAGGTCC
>SNORD109_200
GATCTATGATGACAACAGGTCTCTGAGGACGTTGACTGACTTGAACAGATGCGATCTGAGGTCC
>SNORD109_201
GATCTATGATGAAAACAGGTATCTGAGGACGTTGACTGACTTGAACAGATGCGATCTGAGGTCC
>SNORD109_202
GATCTATGATGACAACAGGTCTCTGAGGACGTTGACTGACTTGAACAGATGCGATCTGAGGTCC
>SNORD109_203
GATCTATGATGACAACAGGTATCTGAGGACGTTGACTGACTTGAACAGATGCGATATGAGGTCC
>SNORD109_204
GATCTATGATGACAACAGGTATCTGAGGACGTTGACTGACTTGAACAGATGCGATATGAGGTCC
>SNORD109_205
GATCTATGATGACAACAGGTATCTGAGGACGTTGACTGACTTGAACAGATGCGATATGAGGTCC
>SNORD109_206
GATCTATGATGACAACAGGTATCTGAGGACGTTGACTGACTTGAACAGATGCGATATGAGGTCC
>SNORD109_207
GATCTATGATGACAACAGGTCTCTGAGGACGTTGACTGACTTGAACAGATGCGATCTGAGGTCC
>SNORD109_208

GAACTATGATGACAACAGGTATCTGAGGACGTTGACTGACTTGAACAGATGCGATCTGAGGTCC
>SNORD109_209
GATCTATGATGACAACAGGTCTCTGAGGACGTTGACTGACTTGAACAGATGCGATCTGAGGTCC
>SNORD109_210
GATCTCTGATGACAACAGGTATCTGAGGACGTTGACTGACTTGAACAGATGCGATCTGAGGTCC
>SNORD109_211
GATCTATGATGACAACAGGTATCTGAGGACGTTGACTGACTTGAACAGATGCGATATGAGGTCC
>SNORD109_212
GATCTATGATGACAACAGGTCTCTGAGGACGTTGACTGACTTGAACAGATGCGATCTGAGGTCC
>SNORD109_213
GATCTATGATGACAACAGGTCTCTGAGGACGTTGACTGACTTGAACAGATGCGATCTGAGGTCC
>SNORD109_214
GATCTATGATGACAACAGGTCTCTGAGGACGTTGACTGACTTGAACAGATGCGATCTGAGGTCC
>SNORD109_215
GATCTATGATGACAACAGGTCTCTGAGGACGTTGACTGACTTGAACAGATGCGATCTGAGGTCC
>SNORD109_216
GATCTATGATGACAACAGGTCTCTGAGGACGTTGACTGACTTGAACAGATGCGATCTGAGGTCC
>SNORD109_217
GATCTATGATGACAACAGGTCTCTGAGGACGTTGACTGACTTGAACAGATGCGATCTGAGGTCC
>SNORD109_218
GATCTATGATGACAACAGGTATCTGAGGACGTTGACTGACCTGAACAGATGCGATCTGAGGTCC
>SNORD109_219
GATCTATGATGACAACAGGTCTCTGAGGACGTTGACTGACTTGAACAGATGCGATCTGAGGTCC
>SNORD109_220
GATCTATGATGACAACAGGTATCTGAGGACGTTGACTGACTTGAACAGATGCGATCTGAGTTCC
>SNORD109_221
GATCTATGATGACAACAGGTATCTGAGGACGTTGACTGACTTGAACAGATGCGATATGAGGTCC
>SNORD109_222
GATCTATGATGACAACAGGTCTCTGAGGACGTTGACTGACTTGAACAGATGCGATCTGAGGTCC
>SNORD109_223
GATCTATGATGACAACAGGTCTCTGAGGACGTTGACTGACTTGAACAGATGCGATCTGAGGTCC
>SNORD109_224
GATCTATGATGACAACAGGTTTCTGAGGACGTTGACTGACTTGAACAGATGCGATCTGAGGTCC
>SNORD109_225
GATCTATGATGACAACAGGTCTCTGAGGACGTTGACTGACTTGAACAGATGCGATCTGAGGTCC
>SNORD109_226
GATCTATGATGACAACAGGTCTCTGAGGACGTTGACTGACTTGAACAGATGCGATCTGAGGTCC
>SNORD109_227
GATCTATGATGACAACAGGTCTCTGAGGACGTTGACTGACTTGAACAGATGCGATCTGAGGTCC
>SNORD109_228
GATCTATGATGACAACAGGTATCTGAGGAGGTTGACTGACTTGAACAGATGCGATCTGAGGTCC
>SNORD109_229
GATATATGATGACAACAGGTATCTGAGGACGTTGACTGACTTGAACAGATGCGATCTGAGGTCC
>SNORD109_230
GATCTATGATGACAACAGGTCTCTGAGGACGTTGACTGACTTGAACAGATGCGATCTGAGGTCC
>SNORD109_231
GATCTATGATGACAACAGGTCTCTGAGGACGTTGACTGACTTGAACAGATGCGATCTGAGGTCC
>SNORD109_232
GATCTATGATGACAACAGGTCTCTGAGGACGTTGACTGACTTGAACAGATGCGATCTGAGGTCC
>SNORD109_233
GATCTATGATGACAACAGGTCTCTGAGGACGTTGACTGACTTGAACAGATGCGATCTGAGGTCC
>SNORD109_234
GATCTATGATGACAACAGGTCTCTGAGGACGTTGACTGACTTGAACAGATGCGATCTGAGGTCC
>SNORD109_235
GATCTATGATGACAACAGGTCTCTGAGGACGTTGACTGACTTGAACAGATGCGATCTGAGGTCC
>SNORD109_236
GATCTATGATGACAACAGGTCTCTGAGGACGTTGACTGACTTGAACAGATGCGATCTGAGGTCC

>SNORD109_237
GATCTATGATGACAACAGGTCTCTGAGGACGTTGACTGACTTGAACAGATGCGATCTGAGGTCC
>SNORD109_238
GATCTATGATGACAACAGGTCTCTGAGGACGTTGACTGACTTGAACAGATGCGATCTGAGGTCC
>SNORD109_239
GATCTATGATGACAACAGGTCTCTGAGGACGTTGACTGACTTGAACAGATGCGATCTGAGGTCC
>SNORD109_240
GATCTATGATGACAACAGGTCTCTGAGGACGTTGACTGACTTGAACAGATGCGATCTGAGGTCC
>SNORD109_241
GATCTATGATGACAACAGGTCTCTGAGGACGTTGACTGACTTGAACAGATGCGATCTGAGGTCC
>SNORD109_242
GATCTATGATGACAACAGGTCTCTGAGGACGTTGACTGACTTGAACAGATGCGATCTGAGGTCC
>SNORD109_243
GATCTATGATGACAACAGGTCTCTGAGGACGTTGACTGACTTGAACAGATGCGATCTGAGGTCC
>SNORD109_244
GATCTATGATGACAACCGGTATCTGAGGACGTTGACTGACTTGAACAGATGTGATATGAGGTCC
>SNORD109_245
GATCTATGATGACAACCGGTATCTGAGGACGTTGACTGACTTGAACAGATGTGATATGAGGTCC
>SNORD109_246
GATCTATGATGACAACAGGTCTCTGAGGACGTTGACTGACTTGAACAGATGCGATCTGAGGTCC
>SNORD109_247
GATCTATGATGACAACAGGTATCTGAGGACGTTGACTGACTTGAACAGATGCGATATGAGGTCC
>SNORD109_248
GATCTATGATGACAACAGGTCTCTGAGGACGTTGACTGACTTGAACAGATGCGATCTGAGGTCC
>SNORD109_249
GATCTATGATGACAACAGGTCTCTGAGGACGTTGACTGACTTGAACAGATGCGATCTGAGGTCC
>SNORD109_250
GATCTATGATGACAACAGGTCTCTGAGGACGTTGACTGACTTGAACAGATGCGATCTGAGGTCC
>SNORD109_251
GATCTATGATGACAACAGGTCTCTGAGGACGTTGACTGACTTGAACAGATGCGATCTGAGGTCC
>SNORD109_252
GATCTATGATGACAACAGGTCTCTGAGGACGTTGACTGACTTGAACAGATGCGATCTGAGGTCC
>SNORD109_253
GATCTATGATGACAACAGGTCTCTGAGGACGTTGACTGACTTGAACAGATGCGATCTGAGGTCC
>SNORD109_254
GATCTATGATGACAACAGGTCTCTGAGGACGTTGACTGACTTGAACAGATGCGATCTGAGGTCC
>SNORD109_255
GATCTATGATGACAACAGGTATCTGAGGATGTTGACTGACTTGAACAGATGCGATCTGAGGTCC
>SNORD109_256
GATCTATGATGACAACAGGTATCTGAGGACGTTGACTGACTTGAACAGATGCGATATGAGGTCC
>SNORD109_257
GATATATGATGACAACAGGTATCTGAGGACGTTGACTGACTTGAACAGATGCGATCTGAGGTCC
>SNORD109_258
GATCTATGATGACAACAGGTCTCTGAGGACGTTGACTGACTTGAACAGATGCGATCTGAGGTCC
>SNORD109_259
GATCTATGATGACAACAGGTCTCTGAGGACGTTGACTGACTTGAACAGATGCGATCTGAGGTCC
>SNORD109_260
GATCTATGATGACAACAGGTCTCTGAGGACGTTGACTGACTTGAACAGATGCGATCTGAGGTCC
>SNORD109_261
GATCTATGATGACAACAGGTCTCTGAGGACGTTGACTGACTTGAACAGATGCGATCTGAGGTCC
>SNORD109_262
GATCTATGATGACAACAGGTCTCTGAGGACGTTGACTGACTTGAACAGATGCGATCTGAGGTCC
>SNORD109_263
GATCTATGATGACAACAGGTATCTGAGGACGTTGACTGACTTGAACAGATGCGATATGAGGTCC
>SNORD109_264
GATCTATGATGACAACAGGTCTCTGAGGACGTTGACTGACTTGAACAGATGCGATCTGAGGTCC
>SNORD109_265

GATCTATGATGACAACAGGTCTCTGAGGACGTTGACTGACTTGAACAGATGCGATCTGAGGTCC
>SNORD109_266
GATCTATGATGACAACAGGTCTCTGAGGACGTTGACTGACTTGAACAGATGCGATCTGAGGTCC
>SNORD109_267
GATCTATGATGACAACAGGTCTCTGAGGACGTTGACTGACTTGAACAGATGCGATCTGAGGTCC
>SNORD109_268
GATCTATGATGACAACAGGTCTCTGAGGACGTTGACTGACTTGAACAGATGCGATCTGAGGTCC
>SNORD109_269
GATCTATGATGACAACAGGTATCTGAGGACGTTGACTGACTTGAACAGATGCGATATGAGGTCC
>SNORD109_270
GATCTATGATGACAACAGGTCTCTGAGGACGTTGACTGACTTGAACAGATGCGATCTGAGGTCC
>SNORD109_271
GATCTATGATGACAACAGGTCTCTGAGGACGTTGACTGACTTGAACAGATGCGATCTGAGGTCC
>SNORD109_272
GATCTATGATGACAACAGGTCTCTGAGGACGTTGACTGACTTGAACAGATGCGATCTGAGGTCC
>SNORD109_273
GATCTATGATGACAACAGGTCTCTGAGGACGTTGACTGACTTGAACAGATGCGATCTGAGGTCC
>SNORD109_274
GATCTATGATGACAACAGGTCTCTGAGGACGTTGACTGACTTGAACAGATGCGATCTGAGGTCC
>SNORD109_275
GATCTATGATGACAACAGGTCTCTGAGGACGTTGACTGACTTGAACAGATGCGATCTGAGGTCC
>SNORD109_276
GATCTATGATGACAACAGGTCTCTGAGGACGTTGACTGACTTGAACAGATGCGATCTGAGGTCC
>SNORD109_277
GATCTATGATGACAACAGGTCTCTGAGGACGTTGACTGACTTGAACAGATGCGATCTGAGGTCC
>SNORD109_278
GATCTATGATGACAACAGGTCTCTGAGGACGTTGACTGACTTGAACAGATGCGATCTGAGGTCC
>SNORD109_279
GATCTATGATGACAACAGGTCTCTGAGGACGTTGACTGACTTGAACAGATGCGATCTGAGGTCC
>SNORD109_280
GATCTATGATGACAACAGGTCTCTGAGGACGTTGACTGACTTGAACAGATGCGATCTGAGGTCC
>SNORD109_281
GATCTATGATGACAACAGGTATCTGAGGACATTGACTGACTTGAACAGATGCGATCTGAGGTCC
>SNORD109_282
GATATATGATGACAACAGGTATCTGAGGACGTTGACTGACTTGAACAGATGCGATCTGAGGTCC
>SNORD109_283
GATCTATGATGACAACAGGTCTCTGAGGACGTTGACTGACTTGAACAGATGCGATCTGAGGTCC
>SNORD109_284
GATCTATGATGACAACAGGTCTCTGAGGACGTTGACTGACTTGAACAGATGCGATCTGAGGTCC
>SNORD109_285
GATCTATGATGACAACAGGTCTCTGAGGACGTTGACTGACTTGAACAGATGCGATCTGAGGTCC
>SNORD109_286
GATCTATGATGACAACAGGTCTCTGAGGACGTTGACTGACTTGAACAGATGCGATCTGAGGTCC
>SNORD109_287
GATCTATGATGACAACAGGTCTCTGAGGACGTTGACTGACTTGAACAGATGCGATCTGAGGTCC
>SNORD109_288
GATCTATGATGACAACAGGTCTCTGAGGACGTTGACTGACTTGAACAGATGCGATCTGAGGTCC
>SNORD109_289
GATCTATGATGACAACAGGTCTCTGAGGACGTTGACTGACTTGAACAGATGCGATCTGAGGTCC
>SNORD109_290
GATCTATGATGACAACAGGTCTCTGAGGACGTTGACTGACTTGAACAGATGCGATCTGAGGTCC
>SNORD109_291
GATCTATGATGACAACAGGTCTCTGAGGACGTTGACTGACTTGAACAGATGCGATCTGAGGTCC
>SNORD109_292
GATCTATGATGACAACAGGTCTCTGAGGACGTTGACTGACTTGAACAGATGCGATCTGAGGTCC
>SNORD109_293
GATCTATGATGACAACAGGTCTCTGAGGACGTTGACTGACTTGAACAGATGCGATCTGAGGTCC

>SNORD109_294
GATCTATGATGACAACAGGTCTCTGAGGACGTTGACTGACTTGAACAGATGCGATCTGAGGTCC
>SNORD109_295
GATCTATGATGACAACAGGTCTCTGAGGACGTTGACTGACTTGAACAGATGCGATCTGAGGTCC
>SNORD109_296
GATCTATGATGACAACAGGTCTCTGAGGACGTTGACTGACTTGAACAGATGCGATCTGAGGTCC
>SNORD109_297
GATCTATGATGACAACAGGTCTCTGAGGACGTTGACTGACTTGAACAGATGCGATCTGAGGTCC
>SNORD109_298
GATCTATGATGACAACAGGTCTCTGAGGACGTTGACTGACTTGAACAGATGCGATCTGAGGTCC
>SNORD109_299
GATCTATGATGACAACAGGTCTCTGAGGACGTTGACTGACTTGAACAGATGCGATCTGAGGTCC
>SNORD109_300
GATCTATGATGACAACAGGTTTCTGAGGACGTTGACTGACTTGAACAGATGCGATCTGAGGTCC
>SNORD109_301
GATCTATGATGACAACAGGTCTCTGAGGACGTTGACTGACTTGAACAGATGCGATCTGAGGTCC
>SNORD109_302
GATCTATGATGACAACAGGTCTCTGAGGAAGTTGACTGACTTGAACAGATGCAATCTGAGGTCC
>SNORD109_303
GATCTATGATGACAACAGGTCTCTGAGGACGTTGACTGACTTGAACAGATGCGATCTGAGGTCC
>SNORD109_304
GATCTATGATGACAACAGGTCTCTGAGGACGTTGACTGACTTGAACAGATGCGATCTGAGGTCC
>SNORD109_305
GATCTATGATGACAACAGGTCTCTGAGGACGTTGACTGACTTGAACAGATGCGATCTGAGGTCC
>SNORD109_306
GATCTATGATGACAACAGGTCTCTGAGGACGTTGACTGACTTGAACAGATGCGATCTGAGGTCC
>SNORD109_307
GATCTATGATGACAACAGGTCTCTGAGGACGTTGACTGACTTGAACAGATGCGATCTGAGGTCC
>SNORD109_308
GATCTATGATGACAACAGGTCTCTGAGGACGTTGACTGACTTGAACAGATGCGATCTGAGGTCC
>SNORD109_309
GATCTATGATGACAACAGGTCTCTGAGGACGTTGACTGACTTGAACAGATGCGATCTGAGGTCC
>SNORD109_310
GATCTATGATGACAACAGGTCTCTGAGGACGTTGACTGACTTGAACAGATGCGATCTGAGGTCC
>SNORD109_311
GATCTATGATGACAACAGGTCTCTGAGGACGTTGACTGACTTGAACAGATGCGATCTGAGGTCC
>SNORD109_312
GATCTATGATGACAACAGGTCTCTGAGGACGTTGACTGACTTGAACAGATGCGATCTGAGGTCC
>SNORD109_313
GATCTATGATGACAACAGGTCTCTGAGGACGTTGACTGACTTGAACAGATGCGATCTGAGGTCC
>SNORD109_314
GATCTATGATGACAACAGGTCTCTGAGGACGTTGACTGACTTGAACAGATGCGATCTGAGGTCC
>SNORD109_315
GATCTATGATGACAACAGGTCTCTGAGGACGTTGACTGACTTGAACAGATGCGATCTGAGGTCC
>SNORD109_316
GATCTATGATGACAACAGGTCTCTGAGGACGTTGACTGACTTGAACAGATGCGATCTGAGGTCC
>SNORD109_317
GATCTATGATGACAACAGGTCTCTGAGGACGTTGACTGACTTGAACAGATGCGATCTGAGGTCC
>SNORD109_318
GATCTATGATGACAACAGGTCTCTGAGGACGTTGACTGACTTGAACAGATGCGATCTGAGGTCC
>SNORD109_319
GATCTATGATGACAACAGGTCTCTGAGGACGTTGACTGACTTGAACAGATGCGATCTGAGGTCC
>SNORD109_320
GATCTATGATGACAACAGGTCTCTGAGGACGTTGACTGACTTGAACAGATGCGATCTGAGGTCC
>SNORD109_321
GATCTATGATGACAACAGGTCTCTGAGGACGTTGACTGACTTGAACAGATGCGATCTGAGGTCC
>SNORD109_322

GATCTATGATGACAACAGGTCTCTGAGGACGTTGACTGACTTGAACAGATGCGATCTGAGGTCC
>SNORD109_323
GATCTATGATGACAACAGGTCTCTGAGGACGTTGACTGACTTGAACAGATGCGATCTGAGGTCC
>SNORD109_324
GATCTATGATGACAACAGGTCTCTGAGGACGTTGACTGACTTGAACAGATGCGATCTGAGGTCC
>SNORD109_325
GATCTATGATGACAACAGGTCTCTGAGGACGTTGACTGACTTGAACAGATGCGATCTGAGGTCC
>SNORD109_326
GATCTATGATGACAACAGGTCTCTGAGGACGTTGACTGACTTGAACAGATGCGATCTGAGGTCC
>SNORD109_327
GATCTATGATGACAACAGGTCTCTGAGGACGTTGACTGACTTGAACAGATGCGATCTGAGGTCC
>SNORD109_328
GATCTATGATGACAACAGGTCTCTGAGGACGTTGACTGACTTGAACAGATGCGATCTGAGGTCC
>SNORD109_329
GATCTATGATGACAACAGGTCTCTGAGGACGTTGACTGACTTGAACAGATGCGATCTGAGGTCC
>SNORD109_330
GATCTATGATGACAACAGGTCTCTGAGGACGTTGACTGACTTGAACAGATGCGATCTGAGGTCC
>SNORD109_331
GATCTATGATGACAACAGGTCTCTGAGGACGTTGACTGACTTGAACAGATGCGATCTGAGGTCC
>SNORD109_332
GATCTATGATGACAACAGGTCTCTGAGGACGTTGACTGACTTGAACAGATGCGATCTGAGGTCC
>SNORD109_333
GATCTATGATGACAACAGGTCTCTGAGGACGTTGACTGACTTGAACAGATGCGATCTGAGGTCC
>SNORD109_334
GATCTATGATGACAACAGGTCTCTGAGGACGTTGACTGACTTGAACAGATGCGATCTGAGGTCC
>SNORD109_335
GATCTATGATGACAACAGGTCTCTGAGGACGTTGACTGACTTGAACAGATGCGATCTGAGGTCC
>SNORD109_336
GATCTATGATGACAACAGGTCTCTGAGGACGTTGACTGACTTGAACAGATGCGATCTGAGGTCC
>SNORD109_337
GATCTATGATGACAACAGGTCTCTGAGGACGTTGACTGACTTGAACAGATGCGATCTGAGGTCC
>SNORD109_338
GATCTATGATGACAACAGGTCTCTGAGGACGTTGACTGACTTGAACAGATGCGATCTGAGGTCC
>SNORD109_339
GATCTATGATGACAACAGGTTTCTGAGGACGTTGACTGACTTGAACAGATGCGATCTGAGGTCC
>SNORD109_340
GATCTATGATGACAACAGGTCTCTGAGGAAGTTGACTGACTTGAACAGATGCAATCTGAGGTCC
>SNORD109_341
GATCTATGATGACAACAGGTCTCTGAGGACGTTGACTGACTTGAACAGATGCGATCTGAGGTCC
>SNORD109_342
GATCTATGATGACAACAGGTCTCTGAGGACGTTGACTGACTTGAACAGATGCGATCTGAGGTCC
>SNORD109_343
GATCTATGATGACAACAGGTCTCTGAGGACGTTGACTGACTTGAACAGATGCGATCTGAGGTCC
>SNORD109_344
GATCTATGATGACAACAGGTCTCTGAGGACGTTGACTGACTTGAACAGATGCGATCTGAGGTCC
>SNORD109_345
GATCTATGATGACAACAGGTCTCTGAGGACGTTGACTGACTTGAACAGATGCGATCTGAGGTCC
>SNORD109_346
GATCTATGATGACAACAGGTCTCTGAGGACGTTGACTGACTTGAACAGATGCGATCTGAGGTCC
>SNORD109_347
GATCTATGATGACAACAGGTCTCTGAGGACGTTGACTGACTTGAACAGATGCGATCTGAGGTCC
>SNORD109_348
GATCTATGATGACAACAGGTCTCTGAGGACGTTGACTGACTTGAACAGATGCGATCTGAGGTCC
>SNORD109_349
GATCTATGATGACAACAGGTCTCTGAGGACGTTGACTGACTTGAACAGATGCGATCTGAGGTCC
>SNORD109_350
GATCTATGATGACAACAGGTCTCTGAGGACGTTGACTGACTTGAACAGATGCGATCTGAGGTCC

>SNORD109_351
GATCTATGATGACAACAGGTCTCTGAGGACGTTGACTGACTTGAACAGATGCGATCTGAGGTCC
>SNORD109_352
GATCTATGATGACAACAGGTCTCTGAGGACGTTGACTGACTTGAACAGATGCGATCTGAGGTCC
>SNORD109_353
GATCTATGATGACAACAGGTCTCTGAGGACGTTGACTGACTTGAACAGATGCGATCTGAGGTCC
>SNORD109_354
GATCTATGATGACAACAGGTCTCTGAGGACGTTGACTGACTTGAACAGATGCGATCTGAGGTCC
>SNORD109_355
GATCTATGATGACAACAGGTCTCTGAGGACGTTGACTGACTTGAACAGATGCGATCTGAGGTCC
>SNORD109_356
GATCTATGATGACAACAGGTCTCTGAGGACGTTGACTGACTTGAACAGATGCGATCTGAGGTCC
>SNORD109_357
GATCTATGATGACAACAGGTCTCTGAGGACGTTGACTGACTTGAACAGATGCGATCTGAGGTCC
>SNORD109_358
GATCTATGATGACAACAGGTCTCTGAGGACGTTGACTGACTTGAACAGATGCGATCTGAGGTCC
>SNORD109_359
GATCTATGATGACAACAGGTCTCTGAGGACGTTGACTGACTTGAACAGATGCGATCTGAGGTCC
>SNORD109_360
GATCTATGATGACAACAGGTCTCTGAGGACGTTGACTGACTTGAACAGATGCGATCTGAGGTCC
>SNORD109_361
GATCTATGATGACAACAGGTCTCTGAGGACGTTGACTGACTTGAACAGATGCGATCTGAGGTCC
>SNORD109_362
GATCTATGATGACAACAGGTCTCTGAGGACGTTGACTGACTTGAACAGATGCGATCTGAGGTCC
>SNORD109_363
GATCTATGATGACAACAGGTCTCTGAGGACGTTGACTGACTTGAACAGATGCGATCTGAGGTCC
>SNORD109_364
GATCTATGATGACAACAGGTCTCTGAGGACGTTGACTGACTTGAACAGATGCGATCTGAGGTCC
>SNORD109_365
GATCTATGATGACAACAGGTCTCTGAGGACGTTGACTGACTTGAACAGATGCGATCTGAGGTCC
>SNORD109_366
GATCTATGATGACAACAGGTCTCTGAGGACGTTGACTGACTTGAACAGATGCGATCTGAGGTCC
>SNORD109_367
GATCTATGATGACAACAGGTCTCTGAGGACGTTGACTGACTTGAACAGATGCGATCTGAGGTCC
>SNORD109_368
GATCTATGATGACAACAGGTCTCTGAGGACGTTGACTGACTTGAACAGATGCGATCTGAGGTCC
>SNORD109_369
GATCTATGATGACAACAGGTCTCTGAGGACGTTGACTGACTTGAACAGATGCGATCTGAGGTCC
>SNORD109_370
GATCTATGATGACAACAGGTCTCTGAGGACGTTGACTGACTTGAACAGATGCGATCTGAGGTCC
>SNORD109_371
GATCTATGATGACAACAGGTCTCTGAGGACGTTGACTGACTTGAACAGATGCGATCTGAGGTCC
>SNORD109_372
GATCTATGATGACAACAGGTCTCTGAGGACGTTGACTGACTTGAACAGATGCGATCTGAGGTCC
>SNORD109_373
GATCTATGATGACAACAGGTCTCTGAGGACGTTGACTGACTTGAACAGATGCGATCTGAGGTCC
>SNORD109_374
GATCTATGATGACAACAGGTCTCTGAGGACGTTGACTGACTTGAACAGATGCGATCTGAGGTCC
>SNORD109_375
GATCTATGATGACAACAGGTCTCTGAGGAAGTTGACTGACTTGAACAGATGCAATCTGAGGTCC
>SNORD109_376
GATCTATGATGACAACAGGTCTCTGAGGACGTTGACTGACTTGAACAGATGCGATCTGAGGTCC
>SNORD109_377
GATCTATGATGACAACAGGTCTCTGAGGACGTTGACTGACTTGAACAGATGCGATCTGAGGTCC
>SNORD109_378
GATCTATGATGACAACAGGTCTCTGAGGACGTTGACTGACTTGAACAGATGCGATCTGAGGTCC
>SNORD109_379

GATCTATGATGACAACAGGTATCTGAGGACGTTGACTGACTTGAACAGATGCGATATGAGGTCC
>SNORD109_380
GATCTATGATGACAACAGGTATCTGAGGGCGTTGACTGACTTGAACAGATGCGATCTGAGGTCC
>SNORD109_381
GATCTATGATGACAACAGGTATCTGAGGGCGTTGACTGACTTGAACAGATGCGATCTGAGGTCC
>SNORD109_382
GATCTATGATGACAACAGGTCTCTGAGGACGTTGACTGACTTGAACAGATGCGATCTGAGGTCC
>SNORD109_383
GATCTATGATGACAACAGGTCTCTGAGGACGTTGACTGACTTGAACAGATGCGATCTGAGGTCC
>SNORD109_384
GATCTATGATGACAACAGGTCTCTGAGGACGTTGACTGACTTGAACAGATGCGATCTGAGGTCC
>SNORD109_385
GATCTATGATGACAACAGGTATCTGAGGACGTTGACTGACTTGAACAGATGCGATATGAGGTCC
>SNORD109_386
GATCTATGATGACAACAGGTATCTGAGGACATTGACTGACTTGAACAGATGAGATCTGAGGTCC
>SNORD109_387
GATCTATGATGACAACAGGTATCTGAGGACGTTGACTGACTTGAACAGATGCGATCTGAAGTCC
>SNORD109_388
GATCTATGATGACAACAGGTATCTGAGGACGTTGACTGACTTGAACAGATGGGATCTAAGGTCC
>SNORD109_389
ATCTATGATGACAACAGGTCTCTGAGGACGTTGACTGACTTGAACAGATGCGATCTGAGGTCC
>SNORD109_390
GATCTATGATGACAACAGGTCTCTGAGGACGTTGACTGACTTGAACAGATGCGATCTGAGGT
>SNORD109_391
GATCTATGATGACAACAGGTCTCTGAGGACGTTGACTGACTTGAACAGATGCGATCTGAGGT
>SNORD109_392
GATCTATGATGACAACAGGTATCTGAGGACGTTGACTGACATGAACAGCTGGGATCTGAGGTCC
>SNORD109_393
GATCTATGATGACAACAGGTATCTGAGGATGTTGACTGACATGAACAGATGCGATCTGAGGTCC
>SNORD109_394
GATCTATGATGACAACAGGTCTCTGAGGACGTTGACTGATTTGAACAGATGCGATCTGAGGTCC
>SNORD109_395
GATCTATTATGACAACAGGTATCTGAGAACGTTGACTGACTTGAACAGATGCGATCTGAGGTCC
>SNORD109_396
GATCTATTATGACAACAGGTATCTGAGAACGTTGACTGACTTGAACAGATGCGATCTGAGGTCC
>SNORD109_397
GATCTAGGATGACAACAGGTCTCTGAGGACGTTGACTGACTTGAACAGATGCGATCTGAGGTCC
>SNORD109_398
GATCTATGATGAAAACAGGTATCTGAGGACGTTGACTGACTTGAATAGATGCGATCTGAGGTCC
>SNORD109_399
GATCTATGATGACAACAGGTCTCTGAGGACGTTGAATGACTTGAACAGATGCGATCTGAGGTCC
>SNORD109_400
GATCTATGAAGACAACAGGTCTCTGAGGACGTTGACTGACTTGAACAGATGCGATCTGAGGTCC
>SNORD109_401
GATCTATGATGACAACAGGTCTCTGAGGACGTTGACTGACCTGAACAGATGCGATCTGAGGTCC
>SNORD109_402
GATCTATGATGACAACAGGTATCTGAGGACGTTGACCTACTTGAACAGATGCGATCTGAGGTCC
>SNORD109_403
GATCTATTATGACAACAGGTCTCTGAGGACGTTGACTGACTTGAACAGATGCGATCTGAGGTCC
>SNORD109_404
GATCTATGATGACAAAAGGTCTCTGAGGACGTTGACTGACTTGAACAGATGCGATCTGAGGTCC
>SNORD109_405
GATCTATGATGACAACAGGTATCTGAGGACGTTGACTGACTAGAACAGATGCGATATGAGGTCC
>SNORD109_406
GATCTATTATGACAACAGGTCTCTGAGGACGTTGACTGACTTGAACAGATGCGATCTGAGGTCC
>SNORD109_407
GATATATGATGACAACAGGTATCTGAGGACATTGACTGACTTGAACAGATGCGATCTGAGGTCC

>SNORD109_408
GATCTATGATGACAACAGGTCTCTGAGGATGTTGACTGACTTGAACAGATGCGATCTGAGGTCC
>SNORD109_409
GATCTATGATGACAACAGGTCTCGGAGGACGTTGACTGACTTGAACAGATGCGATCTGAGGTCC
>SNORD109_410
GATCTATGATGACAACAGGTATCTGAGGACGTTGACTGACTAGAACAGATGCGATATGAGGTCC
>SNORD109_411
GATCTATGATGACAACAGGTCTCTGAGGATGTTGACTGACTTGAACAGATGCGATCTGAGGTCC
>SNORD109_412
GATCTATGATGACAACAGGTCTCTGAGGACGTTGACTGACTTGAACAGATGCGATCTAAGGTCC
>SNORD109_413
GATCTATGATGACAACCGGTATCTGAGGACGTTGACTGACTTGAACAGATGCGATATGAGGTCC
>SNORD109_414
GATCTATGATGACAACAGGTCTCTGAGGACGTTGACTGACTTGAACAGATGCGATCTGAAGTCC
>SNORD109_415
GATCTATGATGACAACAGGTCTCTGAGGACGTTGACTGACTTGAACAGATGCGATCTGAAGTCC
>SNORD109_416
GATCTATGATGACAACAGGTATATGAGGACGTTGACTGACTCGAACAGATGCGATCTGAGGTCC
>SNORD109_417
GATCTATGATGACAACAGGTCTCTGAGGACGTTGACTGACTTGAACAGATGCGATGTGAGGTCC
>SNORD109_418
GATCTATGATGACAACAGGTATCTGAGGACGTTGACTGACTAGAACAGATGCGATATGAGGTCC
>SNORD109_419
GATCTATGATGACAACAGGTCTCTGAGGAAGTTGCCTGACTTGAACAGATGCAATCTGAGGTCC
>SNORD109_420
GATCTATGATGACAACAGGTCTCTGAGGACGTTGACTGACTTGAACAGATGCGATCAGAGGTCC
>SNORD109_421
GATCTATGATGACAACAGGTCTCTGAGGACGTTGACTGACTTGAACAGATGCGATCAGAGGTCC
>SNORD109_422
GATCTATGATGACAACAGGTATCTGAGGACGTTGACTGACTAGAACAGATGCGATATGAGGTCC
>SNORD109_423
GATCTATGATGACAACAGGTCTCTGAGGACGTTGACTGACTTGAACAGATGCGATCTGAAGTCC
>SNORD109_424
GATCTATGATGACAACAGGTATCTGAGGACGTTGACTGACTAGAACAGATGCGATATGAGGTCC
>SNORD109_425
GATCTATGATGACAACAGGTCTCTGAGGACGTTGACTGACTTTAACAGATGCGATCTGAGGTCC
>SNORD109_426
GATCTATGATGACAACAGGTATCTGAGGACGTTGACCTACTTGAACAGATGCGATCTGAGGTCC
>SNORD109_427
GATCTATGATGACAACAGGTATCTGAGGACGTTGACTGACTAGAACAGATGCGATATGAGGTCC
>SNORD109_428
GATCTATGATGACAACAGGTCCCTGAGGACGTTGACTGACTTGAACAGATGCGATCTGAGGTCC
>SNORD109_429
GATCTATGATGACAACAGGTATCTGAGGACGTTGACTGACTAGAACAGATGCGATATGAGGTCC
>SNORD109_430
GATCTATGATGACAACAGGTCTCTGAGGACGTTGACTGACTTGAACAGATGCTGTCTGAGGTCC
>SNORD109_431
GATCTATGATGACAACAGGTATCTGAGGACGTTGACTGACTAGAACAGATGCGATATGAGGTCC
>SNORD109_432
GATCTATGATGACAACAGGTCTCTGAGGACGTTGACTTACTTGAACAGATGCGATCTGAGGTCC
>SNORD109_433
GATCTATGATGACAACAGGTATCTGAGGACGTTGACTGACTAGAACAGATGCGATATGAGGTCC
>SNORD109_434
GATCTATGATGACAACAGGTATCTGAGGACGTTGACTGACTAGAACAGATGCGATATGAGGTCC
>SNORD109_435
GATCTATGATGACAACAGGTATCTGAGGACGTTGACTGACTAGAACAGATGCGATATGAGGTCC
>SNORD109_436

GATCTATGATGACAACAGGTCTCTGAGGACGTTGACTGACTTTAACAGATGCGATCTGAGGTCC
>SNORD109_437
GATCTATGATGACAACAGGTATCTGAGGACGTTGACCTACTTGAACAGATGCGATCTGAGGTCC
>SNORD109_438
GATCTATGATGACAACAGGTATCTGAGGACGTTGACTGACTAGAACAGATGCGATATGAGGTCC
>SNORD109_439
GATCTATGATGACAACAGGTCTCTGAGGAAGTTGCCTGACTTGAACAGATGCAATCTGAGGTCC
>SNORD109_440
GATCTATGATGACAGCAGGTCTCTGAGGACGTTGACTGACTTGAACAGATGCGATCTGAGGTCC
>SNORD109_441
GATCTCTGATGACAACAGGTCTCTGAGGACGTTGACTGACTTGAACAGATGCGATCTGAGGTCC
>SNORD109_442
GATCTATGATGACAACAGGTATCTGAGGACGTTGACTGACTAGAACAGATGCGATATGAGGTCC
>SNORD109_443
GATCTATGATGACAGCAGGTATCTGAGGACGTTGACTTACTTGAACAGATGCGATCTGAGGTC
>SNORD109_444
GATCTCTGATGAAAACAGTATCTGAGGACGTTGACTGACTTGAACAGATGCGATCTGAGGTCC
>SNORD109_445
GATCTCTGATGAAAACAGGTATCTTAGGACGTTGACTGACTTGAACAGATGCGATCTGAGGTCC
>SNORD109_446
GATCTCTAATGAAAACAGGTATCTGAGGACGTTGACTGACTTGAACAGATGCGATCTGAGGTCC
>SNORD109_447
GATCTATGATGACAACAGGTATCTGAGGACGTTGACTGACTAAAACAGATGCTATATGAGGTCC
>SNORD109_448
GATCTATGATGACAACAGGTCTCTGAGGACGTTGACTGACTGGAACAGATGCTGTCTGAGGTCC
>SNORD109_449
GATCTATGATGACAACAGGTCTCTGAGGACGTTGACTGACTTAAACAGATGCGATCTGAAGTCC
>SNORD109_450
GATCTATGATGACAACAGGTCTCTGAGGACGTTGACTGACTTAAACAGATGCGATCTGAAGTCC
>SNORD109_451
GATCTATGATGACAACAGGTATCTGAGGACGTTGATTGACTAGAACAGATGCGATATGAGGTCC
>SNORD109_452
GATCTATGATGACAACAGGTAATTGAGGACGTTGACCTACTTGAACAGATGCGATCTGAGGTCC
>SNORD109_453
GATCTATGATGACAACAGGTATCTGAGGACGTTGACTGACTTGAATATTTCAGATCTGAGGTC
>SNORD109_454
GACCTATGATGACAATTATCTGAGGACACTGACTGACTTGAACAGATGTGATTTTAGGTC
>SNORD109_455
GATCTATGATGACAACAGGTAAGTGATGACAATGACTGACTTAGAAGATGCGATCCGAGGTCC
