## Supplementary Figure 5 for "Evolutionary and biological insights into the paternally expressed Snord116-Ipw-Snord115 gene array at the imprinted Prader-Willi syndrome domain"

### snord64 sequences across mammals

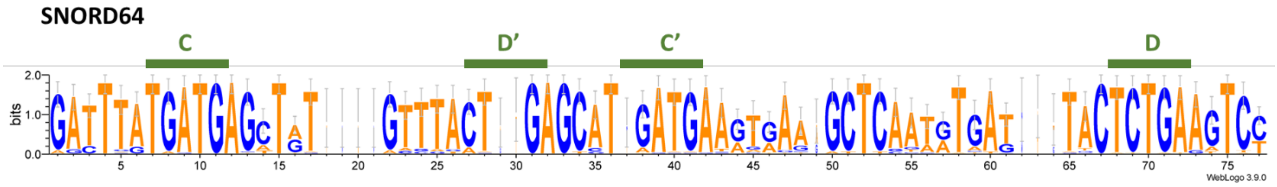


>sorex_araneus
aattcatgatgagctatgtctactgagcatgatgaagtaaagctcggtgtttgagtgctctgaagtcc
>erinaceus_europaeus
gatttatgatgagctatgtttactgagcatgatgaaatgaagcttgatatgagaactctgaggtcc
>talpa_occidentalis
gatttatgatgagctgtgtttactgagcatgatgatgtgaagctcaatgtgagtactctgaagtcc
>manis_pentadactyla
gatttatgatgagttatgtttgctgagcataagtgaagctcaatgtgagtgctctgaaatcc
>felis_catus
gatttatgatgagttatgtttactgagtatgatgaagtgaaactcagtgtgagtactctgaagtcc
>canis_lupus familiaris
agtttataatgatatgatgaggtgtactgagcatgatgaagtgtagctcaatgtaagtactctaaagtcc
>meles_meles
agtttgtgatgagttatgataactgagcatgatgaaatgaaactcagcatgagtactctgaagtat
>neomonachus_schauinslandi
ggtttatgatgagttatgattacagagcatgaagaagtgaagctcaatgtgagtacactgaagtcc
>equus_caballus
a
>tapirus_indicus
gatttatgatgagctatgtttaatgagcatgatgatgcaaagctccatgtcagtactctgaagtcc
>diceros_bicornis minor
gatttatgatgagctatatttaatgagcatgatgatgcaaagctcaatgtcagtactctgtagtcc
>mesoplodon_densirostris
a
>eubalaena_glacialis
a
>eschrichtius_robustus
a
>balaenoptera_acutorostrata
a
>hippopotamus_amphibius
a
>bos_taurus
a
>ovis_aries
a
>sus_scrofa
a
>desmodus_rotundus
gacttatgatgagctgtgttttctgagcatgaccgcgtgaagctcaatctgagtactctgaagtcc
>oryctolagus_cuniculus
gatttatgatgagctgtatttactgagcataatgaagggaggctccatgtgattactctgaagtcc
>marmota_monax
gatttatgatgagctatgtttactgagcatgatgaagtgaagctcaatatgtttactctgaggtcc
>microtus_oregoni
gatttatgatgagctatgtttactgagcatgatgaaatgaggctcaaagtgatgattactctgaaatct
>mesocricetus_auratus
gatttatgatgagctgtgattactgagcatgatgaaataaagctcaaaatgatgactctgaagtct
>rattus_norvegicus

```
gattaatgatgagctgtgtttactgagcataatgaaatgacgctcaaaataattactctgaagtct
>mus_musculus
gacttatgatgagctatgtttactgagcatgatgaaatgatgctcaaaatgattactctgaagtct
>apodemus_sylvaticus
gatttatgatgagttttgtttactgagcatgatgaaatgacgctcaaaatgattactctgaagtct
>eospalax_fontanierii
gatttatgatgagctatgtttactgagcatgatgaaatgaggctcaatatgattactctgaagtct
>nannospalax_galili
gatttatgatgagctatgtttactgagcatgatgaaatgaggctcaatatgattactctgaagtct
>jaculus_jaculus
gatttatgatgagctatgtttactgagcattgatgaaatgaggctcaataagattactctgaagtct
>dipodomys_spectabilis
gatttatgatgagctatatttactgagcgtgatgaggtgaggctcaatatgactactctgaagtct
>heterocephalus_glaber
ggctgatgatgagctgggtgttatgagcatgaatgaaatgaggctcgaaatgattactctgaggtcc
>cavia_porcellus
gactgatgatgagctttgtgtactgagtgtgaatgaacatgaggttcactgtgattactctgaagttc
>chionomys nivalis
gatttatgatgagctatgtttactgagcatgatgaaatgaagctcaaagtgatgattactctgaaatct
>meriones_unguiculatus
gatttatgatgagctgtgtttactgagcatgatgaaatgatgctcaaaataattactctgaagtct
>clethrionomys_rutilus
gatttatgatgagctgtgtttactgagcatgatgaaatgaggctcaaagtgatgattactctgaaatct
>cynocephalus_volans
gatttatgatgagctatgtttgctgagcatgataaagtgaagctcaatgtgattactctgaaatcc
>galeopterus_variegatus
gacttatgatgacctgtgtttactgagcatgatgaagtgaagcttaatgtgattactctgaagttc
>loris_tardigradus
gatttatgatgagctgtgtttaatgagcatgatgaagtaaagctcaatgtgattactctgaagtct
>nycticebus_coucang
gatttatgatgagctgtgtttaatgagcatgatgaagtaaagctcaatgtgattactctgaagtct
>lemur_catta
gatttatgatgagctgtgtttaatgagcatgatgaaataaggctcaatgtgattactctgaagtcc
>eulemur_rufifrons
gatttatggtgagctgtgtttactgagcataatgaagtgaagctcaaaattattactctgaagtcc
>callithrix_jacchus
gatttgtgatgagctgtgttaactgagcatgatgaagtgaagctcaatgtgattactctgaagtcc
>saimiri_boliviensis
gatttgtgatgagctgtgtttactgagcatgatgaagtaaagctcaatgtgattcctctgaagtcc
>nomascus_siki
gatttgtgatgagctgtgtttactgagcatgatgaagtaaagctcaacgtgattactctgaagtcc
>homo_sapiens
gatttataatgagctgtgtttactgagcattatgaagtaaagttcaatatgattactctgaagtct
>cercopithecus_mona
gatttgtgatgagctgtgtttactgagcatgatgaagtaaagctcaatgtgattactctgaagtcc
>macaca_fascicularis
gatttgtgatgagctgtgtttactgagcatgatgaagtaaagctcaatgtgattactctgaagtcc
>rhinopithecus_roxellana
gatttgtgatgagctgtgtttactgagcatgatgaagtaaagctcaatgtgattactctgaagtcc
>cephalopachus_bancanus
gatttatgatgagctgtgtctactgagcatggtgaagtaaagctcaatataattactctgaagtcc
>tupaia_tana
a
>heterohyrax_brucei
a
>trichechus_manatus
gacttatgatgagatatgcttactgtgagcatgatgaagtaaagctcaatatgatttctctgaattcc
>loxodonta_africana
gacttatgatgagatatgctgactatgagcatgatgaagtgaagctcaatatgatttctctgaattcc
>elephas_maximus
gacttatgatgagatatgctgactatgagcatgatgaagtgaagctcaatatgatttctctgaattcc
```

```
>chrysochloris_asiatica
a
>nesogale_talazaci
a
>echinops_telfairi
a
>rhynchocyon petersi
a
>elephantulus edwardii
a
>tolypeutes_matacus
gatttatgatgagatatgtttactgagcctgatgaagggaaagctcattgtgattactctgaagtcc
>dasypus_novemcinctus
gatttatgatgagatatgtttactgagcctgatgaaaagaaagctcactgtgattactctgaagtcc
>tamandua_tetradactyla
gatttatgatgagatatgtttaccgagcctgatgaagggaaagctcattgtaattactctgaagtcc
>choloepus_didactylus
gatttatgatgagatatgtttactgagcctgatgaagggaaagctcattgtgattactttgaagtcc
```

```
                        1       10        20        30        40        50        60        70     77
                        |--------+---------+---------+---------+---------+---------+---------+------|
        sorex_araneus   AATTCATGATGAGCTAT----GTCTACT--GAGCAT-GATGAAGTAAA-GCTCGGTGTTTG-AGTGCTCTGAAGTCC
  erinaceus_europaeus   GATTTATGATGAGCTAT----GTTTACT--GAGCAT-GATGAAATGAA-GCTTGATATGAG---AACTCTGAGGTCC
   talpa_occidentalis   GATTTATGATGAGCTGT----GTTTACT--GAGCAT-GATGATGTGAA-GCTCAATGTGAG---TACTCTGAAGTCC
   callithrix_jacchus   GATTTGTGATGAGCTGT----GTTAACT--GAGCAT-GATGAAGTGAA-GCTCAATGTGAT---TACTCTGAAGTCC
  saimiri_boliviensis   GATTTGTGATGAGCTGT----GTTTACT--GAGCAT-GATGAAGTAAA-GCTCAATGTGAT---TCCTCTGAAGTCC
   cercopithecus_mona   GATTTGTGATGAGCTGT----GTTTACT--GAGCAT-GATGAAGTAAA-GCTCAATGTGAT---TACTCTGAAGTCC
  macaca_fascicularis   GATTTGTGATGAGCTGT----GTTTACT--GAGCAT-GATGAAGTAAA-GCTCAATGTGAT---TACTCTGAAGTCC
 rhinopithecus_roxell   GATTTGTGATGAGCTGT----GTTTACT--GAGCAT-GATGAAGTAAA-GCTCAATGTGAT---TACTCTGAAGTCC
        nomascus_siki   GATTTGTGATGAGCTGT----GTTTACT--GAGCAT-GATGAAGTAAA-GCTCAACGTGAT---TACTCTGAAGTCC
    loris_tardigradus   GATTTATGATGAGCTGT----GTTTAAT--GAGCAT-GATGAAGTAAA-GCTCAATGTGAT---TACTCTGAAGTCT
   nycticebus_coucang   GATTTATGATGAGCTGT----GTTTAAT--GAGCAT-GATGAAGTAAA-GCTCAATGTGAT---TACTCTGAAGTCT
          lemur_catta   GATTTATGATGAGCTGT----GTTTAAT--GAGCAT-GATGAAATAAG-GCTCAATGTGAT---TACTCTGAAGTCC
  cephalopachus_bancan   GATTTATGATGAGCTGT----GTCTACT--GAGCAT-GGTGAAGTAAA-GCTCAATATAAT---TACTCTGAAGTCC
         marmota_monax   GATTTATGATGAGCTAT----GTTTACT--GAGCAT-GATGAAGTGAA-GCTCAATATGTT---TACTCTGAGGTCC
    cynocephalus_volans   GATTTATGATGAGCTAT----GTTTGCT--GAGCAT-GATAAAGTGAA-GCTCAATGTGAT---TACTCTGAAATCC
 oryctolagus_cuniculu   GATTTATGATGAGCTGT----ATTTACT--GAGCAT-AATGAAGGGAG-GCTCCATGTGAT---TACTCTGAAGTCC
     eulemur_rufifrons   GATTTATGGTGAGCTGT----GTTTACT--GAGCAT-AATGAAGTGAA-GCTCAAAATTAT---TACTCTGAAGTCC
          homo_sapiens   GATTTATAATGAGCTGT----GTTTACT--GAGCAT-TATGAAGTAAA-GTTCAATATGAT---TACTCTGAAGTCT
 mesocricetus_auratus   GATTTATGATGAGCTGT----GATTACT--GAGCAT-GATGAAATAAA-GCTCAAAATGAT---GACTCTGAAGTCT
    rattus_norvegicus   GATTAATGATGAGCTGT----GTTTACT--GAGCAT-AATGAAATGAC-GCTCAAAATAAT---TACTCTGAAGTCT
          mus_musculus   GACTTATGATGAGCTAT----GTTTACT--GAGCAT-GATGAAATGAT-GCTCAAAATGAT---TACTCTGAAGTCT
 meriones_unguiculatu   GATTTATGATGAGCTGT----GTTTACT--GAGCAT-GATGAAATGAT-GCTCAAAATAAT---TACTCTGAAGTCT
  apodemus_sylvaticus   GATTTATGATGAGTTTT----GTTTACT--GAGCAT-GATGAAATGAC-GCTCAAAATGAT---TACTCTGAAGTCT
 eospalax_fontanierii   GATTTATGATGAGCTAT----GTTTACT--GAGCAT-GATGAAATGAG-GCTCAATATGAT---TACTCTGAAGTCT
   nannospalax_galili   GATTTATGATGAGCTAT----GTTTACT--GAGCAT-GATGAAATGAG-GCTCAATATGAT---TACTCTGAAGTCT
      jaculus_jaculus   GATTTATGATGAGCTAT----GTTTACT--GAGCATTGATGAAATGAG-GCTCAATAAGAT---TACTCTGAAGTCT
 dipodomys_spectabili   GATTTATGATGAGCTAT----ATTTACT--GAGCGT-GATGAGGTGAG-GCTCAATATGAC---TACTCTGAAGTCT
 galeopterus_variegat   GACTTATGATGACCTGT----GTTTACT--GAGCAT-GATGAAGTGAA-GCTTAATGTGAT---TACTCTGAAGTTC
      microtus_oregoni   GATTTATGATGAGCTAT----GTTTACT--GAGCAT-GATGAAATGAG-GCTCAAAGTGATGATTACTCTGAAATCT
 clethrionomys_rutilu   GATTTATGATGAGCTGT----GTTTACT--GAGCAT-GATGAAATGAG-GCTCAAAGTGATGATTACTCTGAAATCT
            chionomys   GATTTATGATGAGCTAT----GTTTACT--GAGCAT-GATGAAATGAA-GCTCAAAGTGATGATTACTCTGAAATCT
           felis_catus   GATTTATGATGAGTTAT----GTTTACT--GAGTAT-GATGAAGTGAA-ACTCAGTGTGAG---TACTCTGAAGTCC
 neomonachus_schauins   GGTTTATGATGAGTTAT----GATTACA--GAGCAT-GAAGAAGTGAA-GCTCAATGTGAG---TACACTGAAGTCC
      tapirus_indicus   GATTTATGATGAGCTAT----GTTTAAT--GAGCAT-GATGATGCAAA-GCTCCATGTCAG---TACTCTGAAGTCC
    diceros_bicornis   GATTTATGATGAGCTAT----ATTTAAT--GAGCAT-GATGATGCAAA-GCTCAATGTCAG---TACTCTGTAGTCC
     desmodus_rotundus   GACTTATGATGAGCTGT----GTTTTCT--GAGCAT-GACCGCGTGAA-GCTCAATCTGAG---TACTCTGAAGTCC
    tolypeutes_matacus   GATTTATGATGAGATAT----GTTTACT--GAGCCT-GATGAAGGGAAAGCTCATTGTGAT---TACTCTGAAGTCC
  choloepus_didactylus   GATTTATGATGAGATAT----GTTTACT--GAGCCT-GATGAAGGGAAAGCTCATTGTGAT---TACTTTGAAGTCC
 tamandua_tetradactyl   GATTTATGATGAGATAT----GTTTACC--GAGCCT-GATGAAGGGAAAGCTCATTGTAAT---TACTCTGAAGTCC
  dasypus_novemcinctus   GATTTATGATGAGATAT----GTTTACT--GAGCCT-GATGAAAAGAAAGCTCACTGTGAT---TACTCTGAAGTCC
    trichechus_manatus   GACTTATGATGAGATAT----GCTTACTGTGAGCAT-GATGAAGTAAA-GCTCAATATGAT---TTCTCTGAATTCC
     loxodonta_africana   GACTTATGATGAGATAT----GCTGACTATGAGCAT-GATGAAGTGAA-GCTCAATATGAT---TTCTCTGAATTCC
       elephas_maximus   GACTTATGATGAGATAT----GCTGACTATGAGCAT-GATGAAGTGAA-GCTCAATATGAT---TTCTCTGAATTCC
           meles_meles   AGTTTGTGATGAGTTAT----GATAACT--GAGCAT-GATGAAATGAA-ACTCAGCATGAG---TACTCTGAAGTAT
           canis_lupus   AGTTTATAATGATATGATGAGGTGTACT--GAGCAT-GATGAAGTGTA-GCTCAATGTAAG---TACTCTAAAGTCC
    manis_pentadactyla   GATTTATGATGAGTTAT----GTTTGCT--GAGCAT-----AAGTGAA-GCTCAATGTGAG---TGCTCTGAAATCC
 heterocephalus_glabe   GGCTGATGATGAGCTGG----GTGTTAT--GAGCATGAATGAA-ATGAGGCTCGAAATGAT---TACTCTGAGGTCC
       cavia_porcellus   GACTGATGATGAGCTTT----GTGTACT--GAGTGTGAATGAACATGAGGTTCACTGTGAT---TACTCTGAAGTTC
             Consensus   gatTtaTGATGAGcTaT    GttTacT  GAGCaT.gAtGAagtgaa.GCTCaatgTgAt   TaCTCTGAAgTcc
```

### snord107 sequences across mammals

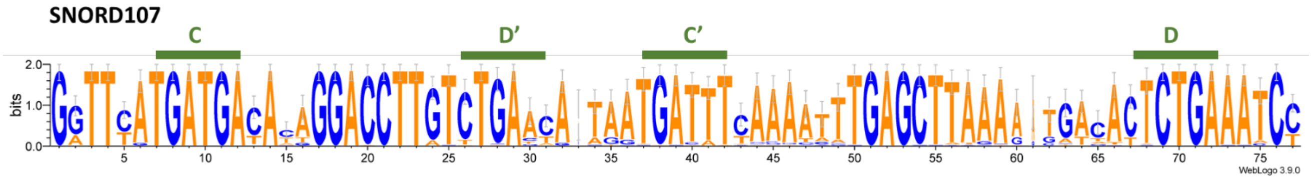


>sorex_araneus
gattaatgatgatacaggaccttgtctgaatatactgatttcaaaatttgagcttaaaataatattctgaaatcc
>erinaceus_europaeus
gattcatgatgacattggaccttgtatgactaaaatgacatcaaaatttgagcttagaattatactctgaaatcc
>talpa_occidentalis
ggttcatgatgacataggaccttgtttgaatataatgattttaaattttgagcttaaaatgacactctgaaatct
>manis_pentadactyla
ggttcatgatgacacaggaccttgtctgaacataatgattttgcaatttgagcttaaaatgacactgaaatcc
>felis_catus
ggttcatgatgacacaggaccttgtctgaacataatgattttaaaatttgagcttaaatgacactctgaaatct
>canis_lupus familiaris
ggttcatgatgacataggaccttgtctgaacataatgattcaaaatttgagcttaaatgacactctgaaatcc
>meles_meles
ggttcatgatgacaaaggaccttgtctgaacataatgatttcaaaacttgagcttaaatgacactctgaaatcc
>neomonachus_schauinslandi
ggttcatgatgacaaaggaccttgtctgaacataatgatttcaaaatttgagcttaaatgacactctgaaatcc
>equus_caballus
ggttcatgatgatacaggaccttgtctgaacaaagtgatttcaaaatttgagcttaaaatgacattctgaaatcc
>tapirus_indicus
ggttcatgatgacacaggaccttgtctgaatataatgatttcaaaatttgagcttaaaatgacactctgaaatcc
>diceros_bicornis minor
ggttcatgatgacacaggaccttgtctgaatataatgatttcaaaatttgagcttaaagtgacactctgaaatcc
>mesoplodon_densirostris
ggtttatgatgacacaggaccttgtctgaatatgatgatttcaaaatttgagcttaaaaggacactctgaaatcc
>eubalaena_glacialis
ggtttatgatgacacaggaccttgtctgaacatgatgatttcaaaatttgagcttaaaaggacactctgaaatcc
>eschrichtius_robustus
ggtttatgatgacacaggaccttgtctgaacatgatgatttcaaaatttgagcttaaaaggacactctgaaatcc
>balaenoptera_acutorostrata
ggtttatgatgacacaggaccttgtctgaacatgatgatttcaaaatttgagcttaaaaggacactctgaaatcc
>hippopotamus_amphibius
ggtttatgatgacacaggaccttgtctgaacatgatgatttcaaaatttgagcttaaagggatactctgaaaccc
>bos_taurus
gatttatgatgacataggaccttgtctgaacatgatgatttcaaagtttgagcttaaaaggacactctgaaatct
>ovis_aries
ggtttatgatgacataggaccttgtctgaacatgatgatttcaaaatttgagcttaaaaggacactctgaaatct
>sus_scrofa
ggtttatgatgacacaggaccttgtctgaacatggtgatttcacattttgagcttaaaaggataccctgaaatcc
>desmodus_rotundus
gatttatgatgagaaaggaccttgtctgaacataatgatttcaaattgtgagctttaaaatgtcactgaaatcc
>oryctolagus_cuniculus
ggttcatgatgacataggaccttgtctgaataataatgatttcaaattttgagctaaagtgactttctgaaatcc
>marmota_monax
ggttcatgatgacacaggaccttgtctgaatatgatgatttcaagatttgagcttaaattgacactctgaaatct
>microtus_oregoni
gatttatgatgatataggaccttgtctgatcaaaatgactttaaaaaatgagcttaaaataactctgaaatct
>mesocricetus_auratus
ggtttatgatgacataggaccttgcctgatcataatgatttaaagaaatgagcttgaaataattctgaaatct
>rattus_norvegicus
gatttatgatgacatgggaccttgtctgaacacaatgatttcaaaaattgagcttagagtgactctgaaatct

>mus_musculus
gatttatgatgatataggaccttgtctgaatataatgatttcaaaaattgagcttacagtgactgaaatct
>apodemus_sylvaticus
gatttatgatgacataggaccttgtctgaatataatgatttcagaaattgagcttagagtaactctgaaatct
>eospalax_fontanierii
ggtttatgatgaaataggaccttgtctgataaaaatgacttcaatatctgagcttaaagtgacactctgagatct
>nannospalax_galili
ggtttatgatgacataggaccttgtctgacaataatgatttcaaaatttgagcttaaagtgacactctgaaatct
>jaculus_jaculus
ggttcatgatgacacaggaacttgtctgaccataatgatttcaaaagtggagcttaaagtgatactctgaagtcc
>dipodomys_spectabilis
ggttcatgatgacactggaccttgtctgactacaaaatgatttcaaatttgagcttaaaaggatattctgaaaac
c
>heterocephalus_glaber
ggtttgtgatgacacaggaccttgtcttaacattatgattttgtattttgagctttatgtgacactgaaatcc
>cavia_porcellus
a
>chionomys nivalis
Gatttatgatgatataggaccttgtctgatgaaaatgattttaaataatgagcttaaaataactctgaaatct
>meriones_unguiculatus
gttttatgatgacataggaccttgtctcaccataaggatatcaaaagttgaggttagattaattctgaaatct
>clethrionomys_rutilus
gatttatgatgatataggaccttgtctgatcaaaatgattttaaaaaatgagcttaaaataattctgaaatct
>cynocephalus_volans
ggttcatgatgacatgggaccttgtctgagtatagtgatttgaaaatttgagcttaaagtgacactctgaaatcc
>galeopterus_variegatus
ggttcatgatgatatgggaccttgtctgagtatagtgatttgaaaatttgagcttaaagtgacactctgaaatcc
>loris_tardigradus
ggttcatgatgacataggaccttgtctgaacacaatgatttcaaaacttgagcttaaaatgacactgaaaacc
>nycticebus_coucang
ggttcatgatgacataggaccttgtctgaacataatgatttcaaaatgtgagcttaaaatgacactgaaaacc
>lemur_catta
ggttcatgatgacataggaccttgtctgaacaaaatgatttcaaaacttgagcttaaaacgatactgaaatcc
>eulemur_rufifrons
ggttcatgatgacataggaccttgtctgaacaaaatgatttcaaaacttgagcttaaaacgatactgaaatcc
>callithrix_jacchus
gattcatgatgatacaggaccttgtctgaacataatgatttcaaaacttgagcttaaaaatgacactctgaaatc
t
>saimiri_boliviensis
gattcatgatgatacaggaccttgtctgaacataatgatttcaaaacttgagcttaaaaatgacactctgaaatc
t
>nomascus_siki
ggttcatgatgacacaggaccttgtctgaacataatgatttcaaaatttgagcttaaaaatgacactctgaaatc
c
>homo_sapiens
ggttcatgatgacacaggaccttgtctgaacataatgatttcaaaatttgagcttaaaaatgacactctgaaatc
c
>cercopithecus_mona
ggttcatgatgacacaggaccttgtctgaacataatgatttcaaaatttgagcttaaaaatgacactctgaaatc
c
>macaca_fascicularis
ggttcatgatgacacaggaccttgtctgaacataatgatttcaaaatttgagcttaaaaatgacactctgaaatc
c
>rhinopithecus_roxellana
ggttcatgatgacacaggaccttgtctgaacataatgatttcaaaatttgagcttaaaaatgacactctgaaatc
c
>cephalopachus_bancanus
ggtttgtgatgatacaggaccttgtctgaacataatgatttcaaaatttgagcttaatatgacactgaaatcc
>tupaia_tana
ggttcatgatgacacgggaccttgtctgaaaaaagtgatttcaaaatttgagcttaaaataatactctgaaatcc
>heterohyrax_brucei

ggttcatgatgatagtggaccttgtctgaacataatgattttaaatttgagctaaaagtgactctgaaatcc
>trichechus_manatus
ggttcgtgatgacaatggaccttatctgaccattatgattttaaatttgagcttaaaatgactctgaaatcc
>loxodonta_africana
ggttcatgatgataatggaccttatctgaatgtaatgatttcaaactttgagcttaaaatgactttctgaaaccc
>elephas_maximus
ggttcatgatgataatggaccttatctgaatgtaatgatttcaaactttgagcttaaaatgactttctgaaaccc
>chrysochloris_asiatica
ggttcatgatgataagggaccttatctgaacataatgatttaaaatttgagcggaaaatggcgctctgaaatcc
>nesogale_talazaci
a
>echinops_telfairi
a
>rhynchocyon_petersi
ggttcatgatgacagtggaccttatctgaacataatgatttttaatttgagcttaaaagtacactctgaaatcc
>elephantulus_edwardii
ggttcatgatgacagtggaccttatttgagcataatgattaaaaaaactgagcttaaaatgacactgaaatcc
>tolypeutes_matacus
gattcatgatgacacaggaccttgtttgaacataatgatattagactttgagcttaaaccacactctgaaatcc
>dasypus_novemcinctus
gattcatgatgacacaggaccttgtttgaaaataatgatattaaactttgagcttaaaccacactctgaaatcc
>tamandua_tetradactyla
gattcatgatgacacaggaccttgtttgaacatagtgatattaaaatttgagcttaaatgacactctgaaatcc
>choloepus_didactylus
gatttatgatgacattggaccttgtttgaacgtgatgatattaaaatttgagcttaaatgacactctgaaatcc

```
                     1       10        20        30        40        50        60        70     77
                     |--------+---------+---------+---------+---------+---------+---------+------|
       sorex_araneus GATTAATGATGATACAGGACCTTGTCTGAATA-TACTGATTTCAAAATTTGAGCTTAAAA-TAATATTCTGAAATCC
  loxodonta_africana GGTTCATGATGATAATGGACCTTATCTGAATG-TAATGATTTCAAACTTTGAGCTTAAAA-TGACTTTCTGAAACCC
     elephas_maximus GGTTCATGATGATAATGGACCTTATCTGAATG-TAATGATTTCAAACTTTGAGCTTAAAA-TGACTTTCTGAAACCC
  talpa_occidentalis GGTTCATGATGACATAGGACCTTGTTTGAATA-TAATGATTTTAAATTTTGAGCTTAAAA-TGACACTCTGAAATCT
  cynocephalus_volans GGTTCATGATGACATGGGACCTTGTCTGAGTA-TAGTGATTTGAAAATTTGAGCTTAAAG-TGACACTCTGAAATCC
galeopterus_variegat GGTTCATGATGATATGGGACCTTGTCTGAGTA-TAGTGATTTGAAAATTTGAGCTTAAAG-TGACACTCTGAAATCC
         felis_catus GGTTCATGATGACACAGGACCTTGTCTGAACA-TAATGATTTTAAAATTTGAGCTTAAA--TGACACTCTGAAATCT
         meles_meles GGTTCATGATGACAAAGGACCTTGTCTGAACA-TAATGATTTCAAAACTTGAGCTTAAA--TGACACTCTGAAATCC
neomonachus_schauins GGTTCATGATGACAAAGGACCTTGTCTGAACA-TAATGATTTCAAAATTTGAGCTTAAA--TGACACTCTGAAATCC
      tapirus_indicus GGTTCATGATGACACAGGACCTTGTCTGAATA-TAATGATTTCAAAATTTGAGCTTAAAA-TGACACTCTGAAATCC
    diceros_bicornis GGTTCATGATGACACAGGACCTTGTCTGAATA-TAATGATTTCAAAATTTGAGCTTAAAG-TGACACTCTGAAATCC
       marmota_monax GGTTCATGATGACACAGGACCTTGTCTGAATA-TGATGATTTCAAGATTTGAGCTTAAAT-TGACACTCTGAAATCT
  callithrix_jacchus GATTCATGATGATACAGGACCTTGTCTGAACA-TAATGATTTCAAAACTTGAGCTTAAAAATGACACTCTGAAATCT
 saimiri_boliviensis GATTCATGATGATACAGGACCTTGTCTGAACA-TAATGATTTCAAAACTTGAGCTTAAAAATGACACTCTGAAATCT
       nomascus_siki GGTTCATGATGACACAGGACCTTGTCTGAACA-TAATGATTTCAAAATTTGAGCTTAAAAATGACACTCTGAAATCC
         homo_sapiens GGTTCATGATGACACAGGACCTTGTCTGAACA-TAATGATTTCAAAATTTGAGCTTAAAAATGACACTCTGAAATCC
  cercopithecus_mona GGTTCATGATGACACAGGACCTTGTCTGAACA-TAATGATTTCAAAATTTGAGCTTAAAAATGACACTCTGAAATCC
 macaca_fascicularis GGTTCATGATGACACAGGACCTTGTCTGAACA-TAATGATTTCAAAATTTGAGCTTAAAAATGACACTCTGAAATCC
rhinopithecus_roxell GGTTCATGATGACACAGGACCTTGTCTGAACA-TAATGATTTCAAAATTTGAGCTTAAAAATGACACTCTGAAATCC
      equus_caballus GGTTCATGATGATACAGGACCTTGTCTGAACA-AAGTGATTTCAAAATTTGAGCTTAAAA-TGACATTCTGAAATCC
         tupaia_tana GGTTCATGATGACACGGGACCTTGTCTGAAAA-AAGTGATTTCAAAATTTGAGCTTAAAA-TAATACTCTGAAATCC
 mesoplodon_densirost GGTTTATGATGACACAGGACCTTGTCTGAATA-TGATGATTTCAAAATTTGAGCTTAAAA-GGACACTCTGAAATCC
 eubalaena_glacialis GGTTTATGATGACACAGGACCTTGTCTGAACA-TGATGATTTCAAAATTTGAGCTTAAAA-GGACACTCTGAAATCC
eschrichtius_robustu GGTTTATGATGACACAGGACCTTGTCTGAACA-TGATGATTTCAAAATTTGAGCTTAAAA-GGACACTCTGAAATCC
balaenoptera_acutoro GGTTTATGATGACACAGGACCTTGTCTGAACA-TGATGATTTCAAAATTTGAGCTTAAAA-GGACACTCTGAAATCC
hippopotamus_amphibi GGTTTATGATGACACAGGACCTTGTCTGAACA-TGATGATTTCAAAATTTGAGCTTAAAG-GGATACTCTGAAACCC
          sus_scrofa GGTTTATGATGACACAGGACCTTGTCTGAACA-TGGTGATTTCACATTTTGAGCTTAAAA-GGATACCCTGAAATCC
          bos_taurus GATTTATGATGACATAGGACCTTGTCTGAACA-TGATGATTTCAAAGTTTGAGCTTAAAA-GGACACTCTGAAATCT
           ovis_aries GGTTTATGATGACATAGGACCTTGTCTGAACA-TGATGATTTCAAAATTTGAGCTTAAAA-GGACACTCTGAAATCT
     jaculus_jaculus GGTTCATGATGACACAGGAACTTGTCTGACCA-TAATGATTTCAAAAGTGGAGCTTAAAG-TGATACTCTGAAGTCC
         canis_lupus GGTTCATGATGACATAGGACCTTGTCTGAACA-TAATGATT-CAAAATTTGAGCTTAAA--TGACACTCTGAAATCC
eospalax_fontanierii GGTTTATGATGAATAGGACCTTGTCTGATAA-AAATGACTTCAATATCTGAGCTTAAAG-TGACACTCTGAGATCT
  nannospalax_galili GGTTTATGATGACATAGGACCTTGTCTGACAA-TAATGATTTCAAAATTTGAGCTTAAAG-TGACACTCTGAAATCT
  manis_pentadactyla GGTTCATGATGACACAGGACCTTGTCTGAACA-TAATGATTTTGCAATTTGAGCTTAAAA-TGACA--CTGAAATCC
cephalopachus_bancan GGTTTGTGATGATACAGGACCTTGTCTGAACA-TAATGATTTCAAAATTTGAGCTTAATA-TGACA--CTGAAATCC
   loris_tardigradus GGTTCATGATGACATAGGACCTTGTCTGAACA-CAATGATTTCAAAACTTGAGCTTAAAA-TGACA--CTGAAAACC
  nycticebus_coucang GGTTCATGATGACATAGGACCTTGTCTGAACA-TAATGATTTCAAAATGTGAGCTTAAAA-TGACA--CTGAAAACC
         lemur_catta GGTTCATGATGACATAGGACCTTGTCTGAACA-AAATGATTTCAAAACTTGAGCTTAAAA-CGATA--CTGAAATCC
    eulemur_rufifrons GGTTCATGATGACATAGGACCTTGTCTGAACA-AAATGATTTCAAAACTTGAGCTTAAAA-CGATA--CTGAAATCC
  tolypeutes_matacus GATTCATGATGACACAGGACCTTGTTTGAACA-TAATGATATTAGACTTTGAGCTTAAA--CCACACTCTGAAATCC
 dasypus_novemcinctus GATTCATGATGACACAGGACCTTGTTTGAAAA-TAATGATATTAAACTTTGAGCTTAAA--CCACACTCTGAAATCC
 tamandua_tetradactyl GATTCATGATGACACAGGACCTTGTTTGAACA-TAGTGATATTAAAATTTGAGCTTAAA--TGACACTCTGAAATCC
 choloepus_didactylus GATTTATGATGACATTGGACCTTGTTTGAACG-TGATGATATTAAAATTTGAGCTTAAA--TGACACTCTGAAATCC
 oryctolagus_cuniculu GGTTCATGATGACATAGGACCTTGTCTGAATAATAATGATTTCAAATTTTGAGCTAAAG--TGACTTTCTGAAATCC
  erinaceus_europaeus GATTCATGATGACATTGGACCTTGTATGACTA-AAATGACATCAAAATTTGAGCTTAGAA-TTATACTCTGAAATCC
 chrysochloris_asiati GGTTCATGATGATAAGGGACCTTATCTGAACA-TAATGATTT-AAAATTTGAGCGGAAAA-TGGCGCTCTGAAATCC
   heterohyrax_brucei GGTTCATGATGATAGTGGACCTTGTCTGAACA-TAATGATTTTAAAATTTGAGCTAAAG---TGACTCTGAAATCC
    trichechus_manatus GGTTCGTGATGACAATGGACCTTATCTGACCA-TTATGATTTTAAAATTTGAGCTTAAAA---TGACTCTGAAATCC
  rhynchocyon_petersi GGTTCATGATGACAGTGGACCTTATCTGAACA-TAATGATTTTTAA-TTTGAGCTTAAAGT-ACACTCTGAAATCC
 elephantulus_edwardi GGTTCATGATGACAGTGGACCTTATTTGAGCA-TAATGATTAAAAAACTGAGCTTAAAA-TGACA--CTGAAATCC
    desmodus_rotundus GATTTATGATGAGAAAGGACCTTGTCTGAACA-TAATGATTTCAAATTGTGAGCTTTAAAATGTCA--CTGAAATCC
 heterocephalus_glabe GGTTTGTGATGACACAGGACCTTGTCTTAACA-TTATGATTTTGTATTTTGAGCTTTA---TGTGACACTGAAATCC
 dipodomys_spectabili GGTTCATGATGACACTGGACCTTGTCTGACTACAAATGATTTCAAATTTGAGCTTAAAA-GGATATTCTGAAAACC
     microtus_oregoni GATTTATGATGATATAGGACCTTGTCTGATCA-AAATGACTTTAAAAAATGAGCTTAAA---ATAACTCTGAAATCT
 clethrionomys_rutilu GATTTATGATGATATAGGACCTTGTCTGATCA-AAATGATTTTAAAAAATGAGCTTAAA---ATAATTCTGAAATCT
            chionomys GATTTATGATGATATAGGACCTTGTCTGATGA-AAATGATTTTAAATAATGAGCTTAAA---ATAACTCTGAAATCT
  mesocricetus_auratus GGTTTATGATGACATAGGACCTTGCCTGATCA-TAATGATTTAAAGAAATGAGCTTGAA---ATAATTCTGAAATCT
    rattus_norvegicus GATTTATGATGACATGGGACCTTGTCTGAACA-CAATGATTTCAAAAATTGAGCTTAGA---GTGACTCTGAAATCT
  apodemus_sylvaticus GATTTATGATGACATAGGACCTTGTCTGAATA-TAATGATTTCAGAAATTGAGCTTAGA---GTAACTCTGAAATCT
         mus_musculus GATTTATGATGATATAGGACCTTGTCTGAATA-TAATGATTTCAAAAATTGAGCTTACA---GTGA--CTGAAATCT
 meriones_unguiculatu GTTTTATGATGACATAGGACCTTGTCTCACCA-TAAGGATATCAAAAGTTGAGGTTAGA---TTAATTCTGAAATCT
            Consensus GgTTcATGATGAcA.aGGACCTTgTcTGAacA taaTGATtTcaaAattTGAGCTTAaA..tgacActCTGAAATCc
```

### snord108 sequences across mammals

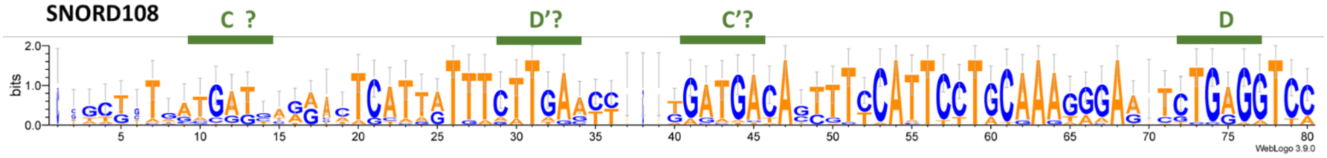


>sorex_araneus
a
>erinaceus_europaeus
a
>talpa_occidentalis
a
>manis_pentadactyla
ccccggcgggcagccagattcactgtttcttgaacttaatgacagtgttcattcctacaaggaaaatctgaggtc
c
>felis_catus
gctttatggtcagactccttgtttcttgactctgatgatagtgttcattcctacaaagggattctcaggtta
>canis_lupus familiaris
Gctctgatgattagactccttgtttattgaacctgatgacaacattcattcctacaaagagaatgcggggttc
>meles_meles
a
>neomonachus_schauinslandi
gcttgatgatcagactcattgtttcttgaacctgatgaaagtgctcattcctaaaagagaatgtggggttc
>equus_caballus
gctttatgatcagactcattgtttcttgagcctaatgacagctttcattcttgcaaagggaaatctgaggtcc
>tapirus_indicus
gctttatgatcagactcattgtttcttgagcctggtgacagctttcactcctgcaaatggaatctgaggtcc
>diceros_bicornis minor
gtgtggttttatgatctcattgtttcttgagcttgatgacagctttcattcctgcaaatggaatctgaagtcc
>mesoplodon_densirostris
a
>eubalaena_glacialis
a
>eschrichtius_robustus
a
>balaenoptera_acutorostrata
a
>hippopotamus_amphibius
a
>bos_taurus
a
>ovis_aries
a
>sus_scrofa
a
>desmodus_rotundus
a
>oryctolagus_cuniculus
a
>marmota_monax
a
>microtus_oregoni
a
>mesocricetus_auratus

a
>rattus_norvegicus
a
>mus_musculus
a
>apodemus_sylvaticus
a
>eospalax_fontanierii
a
>nannospalax_galili
a
>jaculus_jaculus
a
>dipodomys_spectabilis
a
>heterocephalus_glaber
a
>cavia_porcellus
a
>Chionomys nivalis
a
>Meriones_unguiculatus
a
>Clethrionomys_rutilus
a
>cynocephalus_volans
gcttgatgatgagactgatatttatttaccctggtgatattttccattcctcaaaggcaatttgaggtcc
>galeopterus_variegatus
gcttgatgatgagactcatatttatttaccctggtgatattttccattcctcaaagggaatttgaggtcc

>loris_tardigradus
ctatgttgcccagcctgaaactaatttcttgaactagaaggcagtttccatttctgcaaaggaaatctgagatca
>nycticebus_coucang
ccatgttgcccagcctgagactaatttcttgaaccagcagacagtttccatttctgcaaaggaaaactgagatca
>lemur_catta
tcttcatgatgagactcattatttcttgaaccagataatgacagtttccattcttgcaaatggaatctgaggtca
>eulemur_rufifrons
ttttcatgattaaactcattatttcttgaaccagatgatgacagtttccattcttgcaaatggaatctgaggtca
>callithrix_jacchus
gtaggaagagggttggcttgatgatgagatgaattggatgacaccttccattcctgcaaaaggagtctggggtcc
>saimiri_boliviensis
gtaggaggagggttgggttgatggtgagatgggttggatgacactttccattcctgcgaaaggagtctgaggtcc
>nomascus_siki
gcttaatgatgagaatcattatttcttgaattggatgacactttccattcctgcaaagggagtgtgaggtcc
>homo_sapiens
gcttaatgatgagaatcattatttcttgaattggatgacactttccattcctgcaaagggagcgtgaggtcc
>cercopithecus_mona
gtgtgttggtgtagaatcattatttcttaaattggatgacactttccattcctgcaaagagagcctggggtcc
>macaca_fascicularis
gcttaatgatgagaatcattatttcttaaattggatgacactttccattcctgcaaagagagcctggggtcc
>rhinopithecus_roxellana
gcttaatgatgagaatcattatttcttaaattggatgacactttccattcctgcaaagagagcctggggtcc
>cephalopachus_bancanus
ttgacttgagggtgagacttaatttcttgaattggataacagcttccaatcctgtaaggggaatctgaggtcc

>tupaia_tana
a

>heterohyrax_brucei
a
>trichechus_manatus
acttgatgatgagactcattatttcatggacctgacttcagttttaattcttgcgaagggaatctgaggtcc
>loxodonta_africana
a
>elephas_maximus
a
>chrysochloris_asiatica
a
>nesogale_talazaci
a
>echinops_telfairi
a
>rhynchocyon_petersi
a
>elephantulus_edwardii
a
>tolypeutes_matacus
a
>dasypus_novemcinctus
a
>tamandua_tetradactyla
a
>choloepus_didactylus
a

```
                       1       10        20        30        40        50        60        70        80
                       |--------+---------+---------+---------+---------+---------+---------+---------|
  manis_pentadactyla   CCCCGGCGGGCAGCCAGATTCACTGTTTCTTGAACT---TAATGACAGTGTTCATTCCTACAAGGAAAA-TCTGAGGTCC
    loris_tardigradus  CTATGTTGCCCAGCCTGAAACTAA-TTTCTTGAACT---AGAAGGCAGTTTCCATTTCTGCAAAGGAAA-TCTGAGATCA
   nycticebus_coucang  CCATGTTGCCCAGCCTGAGACTAA-TTTCTTGAACC---AGCAGACAGTTTCCATTTCTGCAAAGGAAA-ACTGAGATCA
   callithrix_jacchus   GTAGGAAGAGGGTTGGCTTGATGATGAGATGAATT---GGATGACACCTTCCATTCCTGCAAAAGGAG-TCTGGGGTCC
  saimiri_boliviensis   GTAGGAGGAGGGTTGGGTTGATGGTGAGATGGGTT---GGATGACACTTTCCATTCCTGCGAAAGGAG-TCTGAGGTCC
 cephalopachus_bancan   TTGACTTGAGGGTGAGACT--TAATTTCTTGAATT---GGATAACAGCTTCCAATCCTGTAAGGGGAA-TCTGAGGTCC
          felis_catus    GCT-TTATGGTCAGA-CTCCTTGTTTCTTGACTC---TGATGATAGTGTTCATTCCTACAAAGGGAT-TCTCAGGTTA
          canis_lupus    GCTCTGATGATTAGA-CTCCTTGTTTATTGAACC---TGATGACAACATTCATTCCTACAAAGAGAA-TGCGGGGTTC
neomonachus_schauins     GCT-TGATGATCAGA-CTCATTGTTTCTTGAACC---TGATGAAAGTGCTCATTCCTA-AAAGAGAA-TGTGGGGTTC
       equus_caballus    GCT-TTATGATCAGA-CTCATTGTTTCTTGAGCC---TAATGACAGCTTTCATTCTTGCAAAGGGAAATCTGAGGTCC
      tapirus_indicus    GCT-TTATGATCAGA-CTCATTGTTTCTTGAGCC---TGGTGACAGCTTTCACTCCTGCAAATGGAA-TCTGAGGTCC
     diceros_bicornis    GTG-TGGTTTTATGATCTCATTGTTTCTTGAGCT---TGATGACAGCTTTCATTCCTGCAAATGGAA-TCTGAAGTCC
          lemur_catta    TCT-TCATGATGAGA-CTCATTATTTCTTGAACCAGATAATGACAGTTTCCATTCTTGCAAATGGAA-TCTGAGGTCA
    eulemur_rufifrons    TTT-TCATGATTAAA-CTCATTATTTCTTGAACCAGATGATGACAGTTTCCATTCTTGCAAATGGAA-TCTGAGGTCA
    trichechus_manatus   ACT-TGATGATGAGA-CTCATTATTTCATGGACC---TGACTTCAGTTTTAATTCTTGCGAAGGGAA-TCTGAGGTCC
  cynocephalus_volans    GCT-TGATGATGAGA-CTGAT-ATTTATTTACCC---TGGTGATATTTTCCATTCCT-CAAAGGCAA-TTTGAGGTCC
 galeopterus_variegat    GCT-TGATGATGAGA-CTCAT-ATTTATTTACCC---TGGTGATATTTTCCATTCCT-CAAAGGGAA-TTTGAGGTCC
        nomascus_siki    GCT-TAATGATG-AGAATCATTATTTCTTGAATT---GGATGACACTTTCCATTCCTGCAAAGGGAG-TGTGAGGTCC
          homo_sapiens   GCT-TAATGATG-AGAATCATTATTTCTTGAATT---GGATGACACTTTCCATTCCTGCAAAGGGAG-CGTGAGGTCC
  macaca_fascicularis    GCT-TAATGATG-AGAATCATTATTTCTTAAATT---GGATGACACTTTCCATTCCTGCAAAGAGAG-CCTGGGGTCC
 rhinopithecus_roxell    GCT-TAATGATG-AGAATCATTATTTCTTAAATT---GGATGACACTTTCCATTCCTGCAAAGAGAG-CCTGGGGTCC
   cercopithecus_mona    GTG-TGTTGGTGTAGAATCATTATTTCTTAAATT---GGATGACACTTTCCATTCCTGCAAAGAGAG-CCTGGGGTCC
            Consensus  ..gct.tgatgat..ga.cTcattaTTTctTgaacc   tgatgAcAgttTcCAtTCcTgcaAaggAa tcTGaGGTcc
```

#### snord109 sequences across mammals

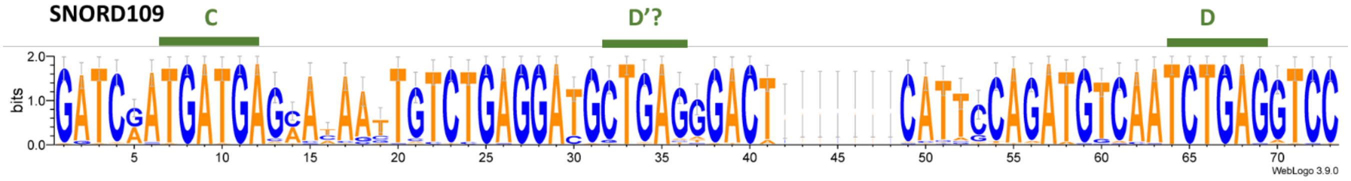


Note - To avoid introducing bias in the sequence alignment, only a randomly selected copy of SNORD109 was retained.

```
>sorex_araneus
gatcaatgatgagaaaaattgtctgaggacgctgagggactcattccagatgtcaatctgaggtcc
>erinaceus_europaeus
gatcgatgatgagaaaactgtctgaggatgctgagagacaaaacaagcactgcagatgtcaatctgaggtcc
>talpa_occidentalis
gatcaatgatgagaaaaattgtctgaggatgctgagggactcattccagatgtcaatctgaggtcc
>manis_pentadactyla
gatcaatgatgagcacaactgtctgaggatgctgagggactcattccagatgtcaatctgaggtcc
>felis_catus
gatcaatgatgagcacaattgtctgaggatgctgagggactcattccagatgtcaatctgaggtcc
>canis_lupus familiaris
gatcaatgatgagcacaattgtctgaggatgctgagggactcattccagatgtcaatctgaggtcc
>meles_meles
gatcaatgatgagcacaattgtctgaggatgctgagggactcattccagatgtcaatctgaggtcc
>neomonachus_schauinslandi
gatcaatgatgagcacaattgtctgaggatgctgagggactcattccagatgtcaatctgaggtcc
>equus_caballus
gatcaatgatgagcataattgtctgaggatgctgagggactcattccagatgtcaatctgaggtcc
>tapirus_indicus
gatcaatgatgagcataattgtctgaggatgctgagggactcattccagatgtcaatctgaggtcc
>diceros_bicornis minor
gatcaatgatgagcacaattgtctgaggatgctgagggactcattccagatgtcaatctgaggtcc
>mesoplodon_densirostris
gatcgatgatgagaaaaattgtctgaggatgctgagggactcattccagatgtcaatctgaggtcc
>eubalaena_glacialis
gatcgatgatgagaaaaattgtctgaggatgctgagggactcattccagatgtcaatctgaggtcc
>eschrichtius_robustus
gatcgatgatgagaaaaattgtctgaggatgctgagggactcattccagatgtcaatctgaggtcc
>balaenoptera_acutorostrata
gatcgatgatgagaaaaattgtctgaggatgctgagggactcattccagatgtcaatctgaggtcc
>hippopotamus_amphibius
gatcaatgatgagaaaaactgtctgaggatgctgagggactcattccagatgtcaatctgaggtcc
>bos_taurus
gatcgatgatgagaaaaactgtctgaggatgctgagggactcattccagatgtcaatctgaggtcc
>ovis_aries
gatcgatgatgagaaaaactgtctgaggatgctgagggactcattccagatgtcaatctgaggtcc
>sus_scrofa
gatcgatgatgagaaaaattgtctgaggatgctgagggactattccagatgtcaatctgaggtcc
>desmodus_rotundus
gatcagtgatgagcacagctgtctggggatgctgagggactcattgcagatgtcagtctgaggtcc
>oryctolagus_cuniculus
gatcgatgatgagaacaagtgtctgaggatgctgagggactcattccagatgtcaatctgaggtcc
>marmota_monax
gatcaatgatgagcacaattgtctgaggatgctgagagactcattccagatgtcaatctgaggtct
>microtus_oregoni (cluster)
gatcgatgatgagaaaccctgtctgaggatgctgagagactcatgtcaatgggaatctgaggtcc
>mesocricetus_auratus (cluster)
gatcaatgatgataacaagtctctgaggatgctgatcaactcatcgctatgtcaatctgagttcc
```

>rattus_norvegicus
a
>mus_musculus
a
>apodemus_sylvaticus
a
>eospalax_fontanierii (cluster)
gatcgatgatgagaacaactgtctgaggacgctgagggactcatacaaatggcaatctgaggtcc
>nannospalax_galili (cluster)
gatcgatgatgagaacaactacctgaggacgctgagggtctcataacaaatggcaatctgaggtcc
>jaculus_jaculus
gatcgatgatgagcacaactgtctgaggatgctgagggacacattccagatgtcaatctgaggtcc
>dipodomys_spectabilis
ggaagcagatgagcacaagggtctgaggacactgaaggactcaagtgcagaagtcaatctgagaagc
>heterocephalus_glaber
gatcgatgaagagaacaattccctgaggacgctgagggactgattttagatgtcaatctgaggtcc
>cavia_porcellus
gatcaatgatgagcacaattgtctgaggatgctgagggactcattccagatgtcaatctg
aggtcc
>meriones_unguiculatus (cluster)
gatctatgatgacaacaggtatctgaggatgttgactgacttgagcagatgtaatctgaggtcc
>clethrionomys_rutilus (cluster)
gatcgatgatgacaacaggtttctgaggatggtgatggactcacacataacccctctgaggtcc
>chionomys_nivalis (cluster)
aattgatgatgacaagaagtctctgaggatgctaactgactcatacaaatggcaatctgaggtcc
>cynocephalus_volans
gatcgatgatgagcaaaattgtctgaggatgctgagggactcattccagatgtcaatctgaggtcc
>galeopterus_variegatus
gatcaatgatgagcataattgtctgaggatgctgagggactcattccagatgtcaatctgaggtcc
>loris_tardigradus
gatcgatgatgagcaaaattgtctgaggacgctgagggactcattccagatgtcaatctgaggtcc
>nycticebus_coucang
gatcgatgatgagcaaaattgtctgaggacgctgagggactcattccagatgtcaatct
gaggtcc
>lemur_catta (cluster)
gatcgatgatgagcctaaatgtttgaggacggttaaggacacatggcagatgtacatctgacgtcc
>eulemur_rufifrons (cluster)
gatcgatgatgagcataagtgtctgaggatggtgagcgacacattgcagatgtcaatctgaggtcc
>callithrix_jacchus
gatcgatgatgagaatacttgtctgaggatgctgagggactcattccagatgtcaatctgaggtcc
>saimiri_boliviensis
gatcgatgatgagaatccttgtctgaggacgctgagggactcattccagatgtcaatctgaggtcc
>nomascus_siki
gatcgatgatgagaataattgtctgaggatgctgagggactcattccagatgtcaatctgaggtcc
>homo_sapiens
gatcgatgatgagaataattgtctgaggatgctgagggactcattccagatgtcaatctgaggtcc
>cercopithecus_mona
gatcgatgatgagaataattgtctgaggatgctgagggactcattccagatgtcaatctgaggtcc
>macaca_fascicularis
gatcgatgatgagaataattgtctgaggatgctgagggactcattccagatgtcaatctgaggtcc
>rhinopithecus_roxellana
gatcgatgatgagaataattgtctgaggatgctgagggactcattccagatgtcaatctgaggtcc
>cephalopachus_bancanus
gatcaatgatgagcataattgtctgaggatgctgagagactcattccagatgtcaatctgaggtcc
>tupaia_tana
gatcgatgatgagcattattgtctgaggatgctgagggactcattccagatgtcaatctgaggtcc
>heterohyrax_brucei
gatcaatgatgagtgtaattgtctgaggatgctgagggactccttccagatgtcaatctgaggtcc
>trichechus_manatus
gatcaatgatgagcataattgtctgaggatgctgagggactcattccagatgtcaatctgaggtcc
>loxodonta_africana

ggtcaatgatgaacataaatgtccgaggatgctgagtgactcattgcagatgtcaatctgagatcc
>elephas_maximus
ggtcaatgatgaacataaatgtccgaggatgctgagtgactcattgcagatgtcaatctgagatcc
>chrysochloris_asiatica
gatcaatgatgagcctagttgtctgaggatgctgagggactcattccagatgtcaatctgaggtcc
>nesogale_talazaci
a
>echinops_telfairi
a
>rhynchocyon_petersi
a
>elephantulus_edwardii
a
>tolypeutes_matacus
gatcaatgatgagcatagttgtctgaggatgctgagggactcattccagatgtcaatctgaggtcc
>dasypus_novemcinctus
gatcaatgatgagcataattgtctgaggatgctgagggactcattctagatgtcaatctgaggtcc
>tamandua_tetradactyla
gatcaatgatgagtataattgtctgaggatgctgagggactcattccagatgtcaatctgaggtcc
>choloepus_didactylus
gatcaatgatgagcataattgtctgaggatgctgagggactcattccagatgtcaatctgaggtcc

```
                         1        10        20        30        40        50        60        70 73
                         |--------+---------+---------+---------+---------+---------+---------+--|
       sorex_araneus     GATCAATGATGAGAAAAATTGTCTGAGGACGCTGAGGGACT-------CATTCCAGATGTCAATCTGAGGTCC
  talpa_occidentalis     GATCAATGATGAGAAAAATTGTCTGAGGATGCTGAGGGACT-------CATTCCAGATGTCAATCTGAGGTCC
mesoplodon_densirost     GATCGATGATGAGAAAAATTGTCTGAGGATGCTGAGGGACT-------CATTCCAGATGTCAATCTGAGGTCC
 eubalaena_glacialis     GATCGATGATGAGAAAAATTGTCTGAGGATGCTGAGGGACT-------CATTCCAGATGTCAATCTGAGGTCC
eschrichtius_robustu     GATCGATGATGAGAAAAATTGTCTGAGGATGCTGAGGGACT-------CATTCCAGATGTCAATCTGAGGTCC
balaenoptera_acutoro     GATCGATGATGAGAAAAATTGTCTGAGGATGCTGAGGGACT-------CATTCCAGATGTCAATCTGAGGTCC
  callithrix_jacchus     GATCGATGATGAGAATACTTGTCTGAGGATGCTGAGGGACT-------CATTCCAGATGTCAATCTGAGGTCC
        nomascus_siki    GATCGATGATGAGAATAATTGTCTGAGGATGCTGAGGGACT-------CATTCCAGATGTCAATCTGAGGTCC
         homo_sapiens    GATCGATGATGAGAATAATTGTCTGAGGATGCTGAGGGACT-------CATTCCAGATGTCAATCTGAGGTCC
  cercopithecus_mona     GATCGATGATGAGAATAATTGTCTGAGGATGCTGAGGGACT-------CATTCCAGATGTCAATCTGAGGTCC
 macaca_fascicularis     GATCGATGATGAGAATAATTGTCTGAGGATGCTGAGGGACT-------CATTCCAGATGTCAATCTGAGGTCC
rhinopithecus_roxell     GATCGATGATGAGAATAATTGTCTGAGGATGCTGAGGGACT-------CATTCCAGATGTCAATCTGAGGTCC
hippopotamus_amphibi     GATCAATGATGAGAAAAACTGTCTGAGGATGCTGAGGGACT-------CATTCCAGATGTCAATCTGAGGTCC
           bos_taurus    GATCGATGATGAGAAAAACTGTCTGAGGATGCTGAGGGACT-------CATTCCAGATGTCAATCTGAGGTCC
           ovis_aries    GATCGATGATGAGAAAAACTGTCTGAGGATGCTGAGGGACT-------CATTCCAGATGTCAATCTGAGGTCC
oryctolagus_cuniculu     GATCGATGATGAGAACAAGTGTCTGAGGATGCTGAGGGACT-------CATTCCAGATGTCAATCTGAGGTCC
  manis_pentadactyla     GATCAATGATGAGCACAACTGTCTGAGGATGCTGAGGGACT-------CATTCCAGATGTCAATCTGAGGTCC
      jaculus_jaculus    GATCGATGATGAGCACAACTGTCTGAGGATGCTGAGGGACA-------CATTCCAGATGTCAATCTGAGGTCC
          felis_catus    GATCAATGATGAGCACAATTGTCTGAGGATGCTGAGGGACT-------CATTCCAGATGTCAATCTGAGGTCC
          canis_lupus    GATCAATGATGAGCACAATTGTCTGAGGATGCTGAGGGACT-------CATTCCAGATGTCAATCTGAGGTCC
          meles_meles    GATCAATGATGAGCACAATTGTCTGAGGATGCTGAGGGACT-------CATTCCAGATGTCAATCTGAGGTCC
neomonachus_schauins     GATCAATGATGAGCACAATTGTCTGAGGATGCTGAGGGACT-------CATTCCAGATGTCAATCTGAGGTCC
    diceros_bicornis     GATCAATGATGAGCACAATTGTCTGAGGATGCTGAGGGACT-------CATTCCAGATGTCAATCTGAGGTCC
     cavia_porcellus     GATCAATGATGAGCACAATTGTCTGAGGATGCTGAGGGACT-------CATTCCAGATGTCAATCTGAGGTCC
      equus_caballus     GATCAATGATGAGCATAATTGTCTGAGGATGCTGAGGGACT-------CATTCCAGATGTCAATCTGAGGTCC
     tapirus_indicus     GATCAATGATGAGCATAATTGTCTGAGGATGCTGAGGGACT-------CATTCCAGATGTCAATCTGAGGTCC
  trichechus_manatus     GATCAATGATGAGCATAATTGTCTGAGGATGCTGAGGGACT-------CATTCCAGATGTCAATCTGAGGTCC
galeopterus_variegat     GATCAATGATGAGCATAATTGTCTGAGGATGCTGAGGGACT-------CATTCCAGATGTCAATCTGAGGTCC
choloepus_didactylus     GATCAATGATGAGCATAATTGTCTGAGGATGCTGAGGGACT-------CATTCCAGATGTCAATCTGAGGTCC
chrysochloris_asiati     GATCAATGATGAGCCTAGTTGTCTGAGGATGCTGAGGGACT-------CATTCCAGATGTCAATCTGAGGTCC
  tolypeutes_matacus     GATCAATGATGAGCATAGTTGTCTGAGGATGCTGAGGGACT-------CATTCCAGATGTCAATCTGAGGTCC
cephalopachus_bancan     GATCAATGATGAGCATAATTGTCTGAGGATGCTGAGAGACT-------CATTCCAGATGTCAATCTGAGGTCC
 cynocephalus_volans     GATCGATGATGAGCAAAATTGTCTGAGGATGCTGAGGGACT-------CATTCCAGATGTCAATCTGAGGTCC
   loris_tardigradus     GATCGATGATGAGCAAAATTGTCTGAGGACGCTGAGGGACT-------CATTCCAGATGTCAATCTGAGGTCC
  nycticebus_coucang     GATCGATGATGAGCAAAATTGTCTGAGGACGCTGAGGGACT-------CATTCCAGATGTCAATCTGAGGTCC
          tupaia_tana    GATCGATGATGAGCATTATTGTCTGAGGATGCTGAGGGACT-------CATTCCAGATGTCAATCTGAGGTCC
dasypus_novemcinctus     GATCAATGATGAGCATAATTGTCTGAGGATGCTGAGGGACT-------CATTCTAGATGTCAATCTGAGGTCC
  heterohyrax_brucei     GATCAATGATGAGTGTAATTGTCTGAGGATGCTGAGGGACT-------CCTTCCAGATGTCAATCTGAGGTCC
tamandua_tetradactyl     GATCAATGATGAGTATAATTGTCTGAGGATGCTGAGGGACT-------CATTCCAGATGTCAATCTGAGGTCC
 saimiri_boliviensis     GATCGATGATGAGAATCCTTGTCTGAGGACGCTGAGGGACT-------CATTCCAGATGTCAATCTGAGGTCC
        marmota_monax    GATCAATGATGAGCACAATTGTCTGAGGATGCTGAGAGACT-------CATTCCAGATGTCAATCTGAGGTCT
           sus_scrofa    GATCGATGATGAGAAAAATTGTCTGAGGATGCTGAGGGACT--------ATTCCAGATGTCAATCTGAGGTCC
    desmodus_rotundus    GATCAGTGATGAGCACAGCTGTCTGGGGATGCTGAGGGACT-------CATTGCAGATGTCAGTCTGAGGTCC
    eulemur_rufifrons    GATCGATGATGAGCATAAGTGTCTGAGGATGGTGAGCGACA-------CATTGCAGATGTCAATCTGAGGTCC
   loxodonta_africana    GGTCAATGATGAACATAAATGTCCGAGGATGCTGAGTGACT-------CATTGCAGATGTCAATCTGAGATCC
     elephas_maximus     GGTCAATGATGAACATAAATGTCCGAGGATGCTGAGTGACT-------CATTGCAGATGTCAATCTGAGATCC
eospalax_fontanierii     GATCGATGATGAGAACAACTGTCTGAGGACGCTGAGGGACT-------CATA-CAAATGGCAATCTGAGGTCC
  nannospalax_galili     GATCGATGATGAGAACAACTACCTGAGGACGCTGAGGGTCT-------CATAACAAATGGCAATCTGAGGTCC
heterocephalus_glabe     GATCGATGAAGAGAACAATTCCCTGAGGACGCTGAGGGACT-------GATTTTAGATGTCAATCTGAGGTCC
    microtus_oregoni     GATCGATGATGAGAAACCCTGTCTGAGGATGCTGAGAGACT-------CATGTCA-ATGGGAATCTGAGGTCC
 erinaceus_europaeus     GATCGATGATGAGAA-AACTGTCTGAGGATGCTGAGAGACAAAACAAGCACTGCAGATGTCAATCTGAGGTCC
          lemur_catta    GATCGATGATGAGCCTAAATGTTTGAGGACGGTTAAGGACA-------CATGGCAGATGTACATCTGACGTCC
mesocricetus_auratus     GATCAATGATGATAACAAGTCTCTGAGGATGCTGATCAACT-------CATCGCT-ATGTCAATCTGAGTTCC
   chionomys_nivalis     AATTGATGATGACAAGAAGTCTCTGAGGATGCTAACTGACT-------CATACAA-ATGGCAATCTGAGGTCC
dipodomys_spectabili     GGAAGCAGATGAGCACAAGGGTCTGAGGACACTGAAGGACTC------AAGTGCAGAAGTCAATCTGAGAAGC
merionnes_unguiculatu    GATCTATGATGACAACAGGTATCTGAGGATGTTGACTGACT--------TGAGCAGATGTAA-TCTGAGGTCC
clethrionomys_rutilu     GATCGATGATGACAACAGGTTTCTGAGGATGGTGATGGACT--------CACACATAACCCC-TCTGAGGTCC
            Consensus    GATCgATGATGAgaA.Aa.TgTCTGAGGAtGcTGAggGACt       cAttcCAgATGtcAaTCTGAGgTCC
```
